# Conserved and specific genomic features of endogenous polydnaviruses revealed by whole genome sequencing of two ichneumonid wasps

**DOI:** 10.1101/861310

**Authors:** Fabrice Legeai, Bernardo F. Santos, Stéphanie Robin, Anthony Bretaudeau, Rebecca B. Dikow, Claire Lemaitre, Véronique Jouan, Marc Ravallec, Jean-Michel Drezen, Denis Tagu, Gabor Gyapay, Xin Zhou, Shanlin Liu, Bruce A. Webb, Seán G. Brady, Anne-Nathalie Volkoff

**Author notes:** Co-first authors (Fabrice Legeai, Bernardo F. Santos, Stéphanie Robin). **Authors e-mail addresses** Fabrice LEGEAI, (co-first author), Bernardo F. SANTOS, (co-first author), Stéphanie ROBIN, (co-first author), Anthony BRETAUDEAU, Rebecca B. DIKOW, Claire LEMAITRE, Véronique JOUAN, Marc RAVALLEC, Jean-Michel DREZEN, Denis TAGU, Gabor GYAPAY, Xin ZHOU, Shanlin LIU, Bruce A. WEBB, Seán G. BRADY, Anne-Nathalie VOLKOFF, (corresponding author).

## Abstract

Polydnaviruses (PDVs) are mutualistic endogenous viruses associated with some lineages of parasitoid wasps that allow successful development of the wasps within their hosts. PDVs include two taxa resulting from independent virus acquisitions in braconid (bracoviruses) and ichneumonid wasps (ichnoviruses). PDV genomes are fully incorporated into the wasp genomes and comprise (1) virulence genes located on proviral segments that are packaged into the viral particle, and (2) genes involved in the production of the viral particles, which are not encapsidated. Whereas the genomic organization of bracoviruses within the wasp genome is relatively well known, the architecture of endogenous ichnoviruses remains poorly understood. We sequenced the genome of two ichnovirus-carrying wasp species, *Hyposoter didymator* and *Campoletis sonorensis*. Complete assemblies with long scaffold sizes allowed identification of the integrated ichnovirus, highlighting an extreme dispersion within the wasp genomes of the viral loci, *i.e.* isolated proviral segments and clusters of replication genes. Comparing the two wasp species, proviral segments harbor distinct gene content and variable genomic environment, whereas viral machinery clusters show conserved gene content and order, and can be inserted in collinear wasp genomic regions. This distinct architecture is consistent with the biological properties of the two viral elements: proviral segments producing virulence proteins allowing parasitism success are fine-tuned to the host physiology, while an ancestral viral architecture was likely maintained for the genes involved in virus particle production. Finding a distinct genomic architecture of ichnoviruses and bracoviruses highlights different evolutionary trajectories leading to virus domestication in the two wasp lineages.

## Background

Host-parasite interactions are one of the most fundamental ecological relationships, and yet one of the most complex from a mechanistic and evolutionary perspective. Hosts and parasites are involved in a continual coevolutionary arms race, with hosts evolving various defense mechanisms and parasites developing strategies to overcome them [1, 2]. Identifying the genomic basis of such adaptations is crucial to understand the dynamics of host-parasite interactions [3]. In fact, the cycle of adaptations and counter-adaptations involved in parasitism scenarios can result in complex biological strategies with far-reaching consequences at the genomic level. The use of endogenous viruses by parasitoid wasps represents a notable example of how complex host-parasite interactions can result in novel genomic adaptations.

Parasitoid wasps are among the most successful groups of parasitic organisms, potentially comprising several hundred thousand species and playing major ecological roles in terrestrial ecosystems [4]. While the adult wasps are free-living, at their immature stages they develop as parasites of other arthropods, eventually killing their host. Many groups of parasitoids are “koinobiont” parasitoids, which means that they develop inside a host that continues to develop after being parasitized, so that the wasp larva needs to deal with the immune system and physiology of the developing host. In order to cope with this biological constraint, some lineages of parasitic wasps have developed an astonishing strategy to manipulate their host by employing mutualistic viruses from the Polydnaviridae (PDVs) family.

PDVs are unusual viruses with a packaged genome composed of several circular molecules, or “segments”, of double-stranded DNA (hence the name “poly-dna virus”). They have been reported in the hyperdiverse wasp families Braconidae and Ichneumonidae. Ichnoviruses (IV) are associated with ichneumonid wasps from the subfamilies Campopleginae and Banchinae, whereas the bracoviruses (BVs) are associated with braconid wasps from the “microgastroid complex” [5, 6]. IVs and BVs differ in their morphology and gene content, but share a common life cycle [7]. The viral particles are produced exclusively within specialized cells located in the calyx region of the ovary during wasp female pupation. Mature virions are secreted into the oviduct lumen and transferred into the host during oviposition. Once the parasitoid’s host, usually a caterpillar, is infected, PDVs do not replicate but expressed genes induce profound physiological alterations in the parasitized host, such as impairment of the immune response or developmental alterations, which are required for successful development of the wasp larva [8–12].

The DNA segments enclosed in PDV particles have been sequenced from purified particles for several PDVs (see review in [9]). The segments consist in circular DNA molecules generated from template sequences integrated in the wasp genome [13, 14]. Currently, knowledge on how PDVs are organized into the wasp genome mainly concerns the bracoviruses [15]. It is known that most of the BV proviral segments are distributed in clusters or “macro-loci”, in which viral segments are organized in tandem arrays, separated by regions of intersegmental DNA that are not encapsidated [16–18]. In contrast, there is still very little information on the distribution and organization of ichnoviruses within the wasp genome.

PDV proviral sequences are amplified and circularized giving rise to the segments packaged in the particles by mechanisms still poorly understood. In bracoviruses, segment production involves direct repeated sequences present at the ends of each proviral segment [19, 20]. These “direct repeat junctions” (DRJs), also named “wasp integration motifs” (WIM), are conserved between BV segments [21]. Presence of direct repeats has also been reported at the extremities of IV segments [22, 23], but this finding was restricted to a few segments and so far there is no evidence of a conserved motif in IV DRJs, suggesting that segment excision may rely on different mechanisms in the two groups of PDV.

PDV insertions in the wasp genome do not consist only of proviral segments, but also include genes not packaged in the virion involved in the production of the virus particles [24–26]. These “viral machineries” derive from ancestral viruses endogenized by parasitic wasps, and they differ between BVs and IVs. BV-associated braconid wasps rely on a set of endogenous nudiviral genes to produce the BV particles [24]. The similarity of BV replication genes to nudivirus genes and their well-known baculovirus homologues made it possible to predict and test their function. It was thus shown that the nudivirus genes maintained in the wasps were mainly those encoding structural proteins; those involved in DNA replication have apparently been lost [24, 27]. These nudivirus-like insertions have also been shown to be more dispersed in the wasp genome compared to the proviral segments [27]. In contrast, the viral ancestor of IVs is not closely related to known pathogenic viruses: a series of conserved genes involved in IVs particle formation has been clearly identified but they show no similarity with known viral genes [26]. These genes are organized within the wasp genome in large clusters named “Ichnovirus Structural Proteins Encoding Regions” (IVSPER; [26]). So far, three IVSPERs enclosing approximately 40 genes have been identified in the campopleginae *Hyposoter didymator* [26] and in the banchinae *Glypta fumiferanae* [28], based on sequencing of corresponding regions from the wasp genomes (using a BAC approach). However, how IVSPERs are distributed within the wasp genome in not known and other IVSPERs might remain undisclosed.

Elucidating how viral insertions are distributed and organized in the wasp genomes is important to fully characterize the machinery that produces PDVs, a necessary step toward understanding the mechanisms that have driven the “domestication” of viruses in parasitic wasps. Yet, there is a dearth of information regarding the distribution of IV sequences in IV-carrying wasp genomes. For example, are IV proviral segments also clustered in replication units like the ones found in BVs? Are there conserved recombination motifs analogous to those seen in BVs? Is the position and gene composition of IVSPERs conserved across wasp species within the same lineage? Since IVs and BVs derive from the integration of unrelated viral ancestors, comparing their genomic characteristics can provide insights on whether similar selection forces have operated on the domestication of the two types of ancestral viruses.

To answer these questions, we sequenced the genome of two ichneumonid wasps from the subfamily Campopleginae, *Hyposoter didymator* and *Campoletis sonorensis*. Both species are parasitoids of larvae of owlet moths (Lepidoptera, Noctuidae) and are associated with endogenous ichnoviruses (respectively, HdIV and CsIV) putatively descending from a common viral ancestral integration. Both HdIV and CsIV genomes packaged in virus particles have been formerly Sanger sequenced ([29] for CsIV; [30] for HdIV) showing they share homologous genes. For both wasp species, we assembled high quality genomes that allowed us to decipher the genome architecture of the endogenous IVs, and to point out differences with that of BVs. In addition, comparison of the IV genome organization and gene content in the two campoplegine wasps indicated strong conservation of the virus-derived replicative machinery whereas the sequences packaged in the IV particles are far more divergent and species specific. These first data relative to genomic architecture of endogenous IVs represent a first crucial step in our understanding of IV evolution.

## Results

### Hyposoter didymator *and* Campoletis sonorensis *genomes share common features*

Whole genome sequencing was performed from haploid male wasp DNA using Illumina Hiseq technology. Assembly of the sequenced reads was conducted using either Supernova v.2.1.1 [31] or Platanus assembler v1.2.1 [32], depending on the species (see Methods section). The draft assembled genome of *H. didymator* consists of 199 Mb in 2,591 scaffolds ranging in size from 1 Kbp to 15.7 Mbp, with a scaffold N50 of 3.999 Mbp and a contig N50 of 151,312 bp (Table 1). The *C. sonorensis* assembled genome consists of 259 Mb in 11,756 scaffolds with sizes ranging from 400 bp to 6.1 Mbp, with an N50 of 725,399 bp and a contig N50 of 315,222 bp (Table 1). For both ichneumonid species, G+C content was similar to most other parasitoid species (between 33.6% and 39.5%) (Table 1).

**Table 1.** Summary statistics for *H. didymator* and *C. sonorensis* assembled genomes compared to other selected parasitoid genomes. Assemblathon2 [53] was used to calculate metrics of genome assemblies.

|  | *Hyposoter didymator* | *Campoletis sonorensis* | *Venturia canescens* | *Microplitis demolitor* | *Fopius arisanus* | *Diachasma alloeum* | *Nasonia vitripennis* |
| --- | --- | --- | --- | --- | --- | --- | --- |
| Number of scaffolds | 2,591 | 11,756 | 62,001 | 1,794 | 1,042 | 3,968 | 6,098 |
| Total length (Mbp) | 198.7 | 258.9 | 237.8 | 241.2 | 153.6 | 388.8 | 295.8 |
| Longest scaffold (Mbp) | 15.7 | 6.1 | 0.85 | 7.15 | 5.5 | 6.61 | 33.57 |
| Scaffold N50 (Mbp) | 4.00 | 0.73 | 0.114 | 1.14 | 0.98 | 0.65 | 0.90 |
| Median scaffold size (nt) | 1,941 | 3,200 | 233 | 2,621 | 12,305 | 4,372 | 2,037 |
| Contig N50 (bp) | 151,312 | 315,222 | 15,077 | 14,499 | 59,408 | 50,453 | 18,840 |
| %N | 1.35% | 0.40% | 1.70% | 14.65% | 8.24% | 6.16% | 19.33% |
| GC (%) | 39.5% | 37.2% | 39.33% | 25.99% | 35.42% | 36.09% | 33.65% |
| Reference | / | / | Pichon *et al.*, 2015 [62] | Burke *et al.*, 2018 [27] | Geib *et al.*, 2017 [59] | Tvedte *et al.*, 2019 [60] | Werren *et al.*, 2010 [61] |

Transposable elements (TE) represent 15.09 % of *H. didymator* and 17.38% of *C. sonorensis* genomes. The major TE groups (LTR, LINE, SINE retrotransposons, and DNA transposons) contribute to 54 % of the total TE coverage in *H. didymator* and up to 79% in *C. sonorensis* (Table 2). The two wasp species differ by the number of class 1 elements (retrotransposons), which was higher in *C. sonorensis* (46% of the TEs) compared to *H. didymator* genome (24% of the TEs).

**Table 2.** Transposable elements (TE) in ichneumonid genomes. Total amount and relative proportion of LINE, LTR, SINE retrotransposons and DNA transposons in the genomes of *Hyposoter didymator* and *Campoletis sonorensis*.

| TE classes | *Hyposoter didymator* |  | *Campoletis sonorensis* |  |
| --- | --- | --- | --- | --- |
|  | Number | Percentage of TE classes | Number | Percentage of TE classes |
| **LINE** | 98 | 10% | 189 | 20% |
| **LTR** | 106 | 11% | 193 | 20% |
| **SINE** | 33 | 3% | 56 | 6% |
| **DNA** | 299 | 30% | 314 | 33% |
| **Others** | 464 | 46% | 199 | 21% |
| **Total** | 1000 | / | 951 | / |

Automatic gene annotation for *H. didymator* (for which RNAseq datasets were available) and for *C. sonorensis* (for which no RNAseq dataset was available) yielded 18,119 and 21,915 transcripts, respectively (Table 3). Although different software packages were used for gene prediction, the two species have similar gene annotation statistics, except for the transcript size, which is longer in *H. didymator*, which also shows a higher predicted intron size (Table 3). BUSCO analyses indicate a high level of completeness of the two genome assemblies and annotations, with 99% of the BUSCO Insecta protein set (1,658 proteins) identified as complete sequences (Figure 1A).

**Figure 1.**
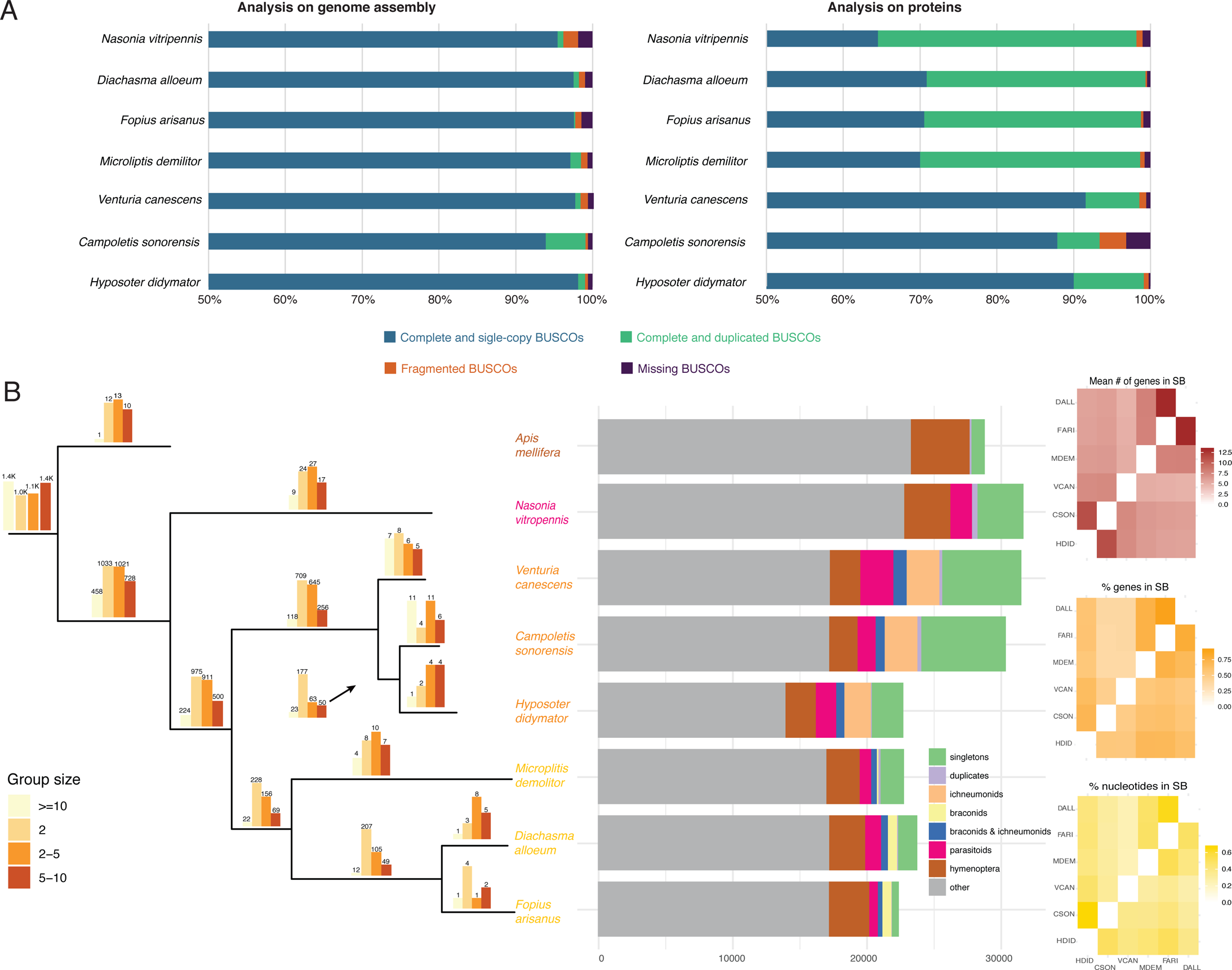
Genomic features of *Campoletis sonorensis* and *Hyposoter didymator* genomes. **A.** BUSCO analysis of parasitoid wasp genomes. BUSCO 3.0.2 analysis with BUSCO Insecta protein set (1,658 proteins). On the left, BUSCO analysis results using the genomes assemblies; on the right, BUSCO analysis results using the predicted protein set. X axis starts at 50% for better visualization. **B. Left panel:** Barplots above each branch of the phylogenic tree indicate the number of orthogroups specific to each species or group of species; the color of the bar indicates the size range of the corresponding orthogroups. **Mid panel:** For each of the hymenoptera species, are indicated the number of genes (i) specific to the species and present either as singletons or duplicates; (ii) present in ichneumonids; (iii) present in braconids; (iv) present in both ichneumonids and braconids; (v) present in all parasitoids and (vi) present in all hymenoptera. **Right panel:** Matrices (represented as heatmaps) indicating, for each couple of species, the mean number (#) of genes in synteny blocs (SB), the percentage (%) of genes in SBs, and the size of the genome (% nucleotides) in SBs. HDID, *Hyposoter didymator* (ichneumonid); CSON, *Campoletis sonorensis* (ichneumonid); VCAN, *Venturia canescens* (ichneumonid); MDEM, *Microplitis demolitor* (braconid); FARI, *Fopius arisanus* (braconid); DALL, *Diachasma alloeum* (braconid).

**Table 3.** Gene annotation statistics for *H. didymator* and *C. sonorensis* assembled genomes.

|  | *Hyposoter*<br>*didymator* | *Campoletis*<br>*sonorensis* |
| --- | --- | --- |
| Transcript number | 18,119 | 21,915 |
| Total transcript size (nt) | 95,868,518 | 68,669,265 |
| Mean transcript size (nt) | 5,291 | 3,133 |
| Median transcript size (nt) | 2,428 | 1,934 |
| Total exon number | 98,639 | 93,590 |
| Mean exon number | 5.4 | 4.3 |
| Median exon number | 4 | 3 |
| Total exon size (nt) | 28,900,964 | 27,144,418 |
| Mean exon size (nt) | 292 | 290 |
| Median exon size (nt) | 186 | 192 |
| Total intron size (nt) | 66,540,946 | 41,453,172 |
| Mean intron size (nt) | 826 | 578 |
| Median intron size (nt) | 245 | 296 |
| Total CDS size (nt) | 28,900,964 | 27,144,418 |
| Mean CDS size (nt) | 1,595 | 1,239 |
| Median CDS size (nt) | 1,090 | 806 |

Orthologous protein sequence families were calculated with Orthofinder by computing each pairs’ similarity among the genomes of different parasitoid wasps from the ichneumonid and braconid families, harboring polydnaviruses or not. For *H. didymator* and *C. sonorensis*, a total number of ∼10,000 orthogroups was identified (Figure 1B, Additional file 1A). The orthogroups included a large majority of the *H. didymator* (87.1%) and *C. sonorensis* (71.4%) genes. Amongst those, only a small portion corresponded to species-specific orthogroups: 11 orthogroups for *H. didymator* (69 genes) and 36 for *C. sonorensis* (288 genes). The number of shared orthogroups declines with the increasing evolutionary distance among the other species (from *Venturia canescens* to *Drosophila*, Additional file 1B). Amongst the orthogroups shared by *H. didymator* and *C. sonorensis* genes, 313 were specific to these two IV-carrying species (Figure 1B, Additional file 1C), representing 875 proteins for *C. sonorensis* and 509 proteins for *H. didymator*.

Global synteny analysis demonstrated the existence of a number of syntenic blocks between the two genomes, enabling the evaluation of the magnitude of the genomic reorganization between the two species even when using fragmented assemblies. When comparing the *C. sonorensis* and *H. didymator* genomes, the mean number of genes per synteny block obtained is 11.2, one of the highest pairwise values for the evaluated species, just below *F. arisanus* and *D. alloeum* (Figure 1B, right panel; Additional file 2). The percentage of regions in syntenic blocks shared between *C. sonorensis* and *H. didymator* compared to the complete genome size is respectively 67% for *H. didymator* and 50% for *C. sonorensis* (Figure 1B, right panel). Finally, the percentage of genes within syntenic blocks is respectively 71% for *H. didymator* and 54% for *C. sonorensis* (Figure 1B, right panel). These observed high pairwise values suggest that global collinearity of *H. didymator* and *C. sonorensis* genomes is well conserved.

### The two campoplegine genomes include numerous and dispersed ichnovirus loci

In the assembled *C. sonorensis* genome, a total of 35 scaffolds, ranging in size from 2.3 Kbp to more than 6 Mbp, contained CsIV sequences (Figure 2A, Additional file 3). Within these scaffolds, 40 viral loci were identified, corresponding either to CsIV segments or, for the first time in this species, to IVSPERs or IVSPER genes. A total of 31 proviral segments were recognized, with sizes varying from 6.4 to 23.2 Kbp (Additional file 3). Compared to the CsIV segments described in Webb et al., 2006, two segments were not found, and eight novel segments were identified (Table 4). Noteworthy, two short scaffolds each contained a repeat element gene (i.e. a member of a gene family encoded by IV segments), corresponding probably to additional viral segments (Table 4). Altogether, *C. sonorensis* genome contained 33 loci corresponding to CsIV proviral segments. In addition, we identified seven gene clusters corresponding to IVSPERs comprising 48 genes located in six different scaffolds; these IVSPERs varied in size from 8.6 Kbp to 33.3 Kbp (Additional files 3 and 4).

**Figure 2.**
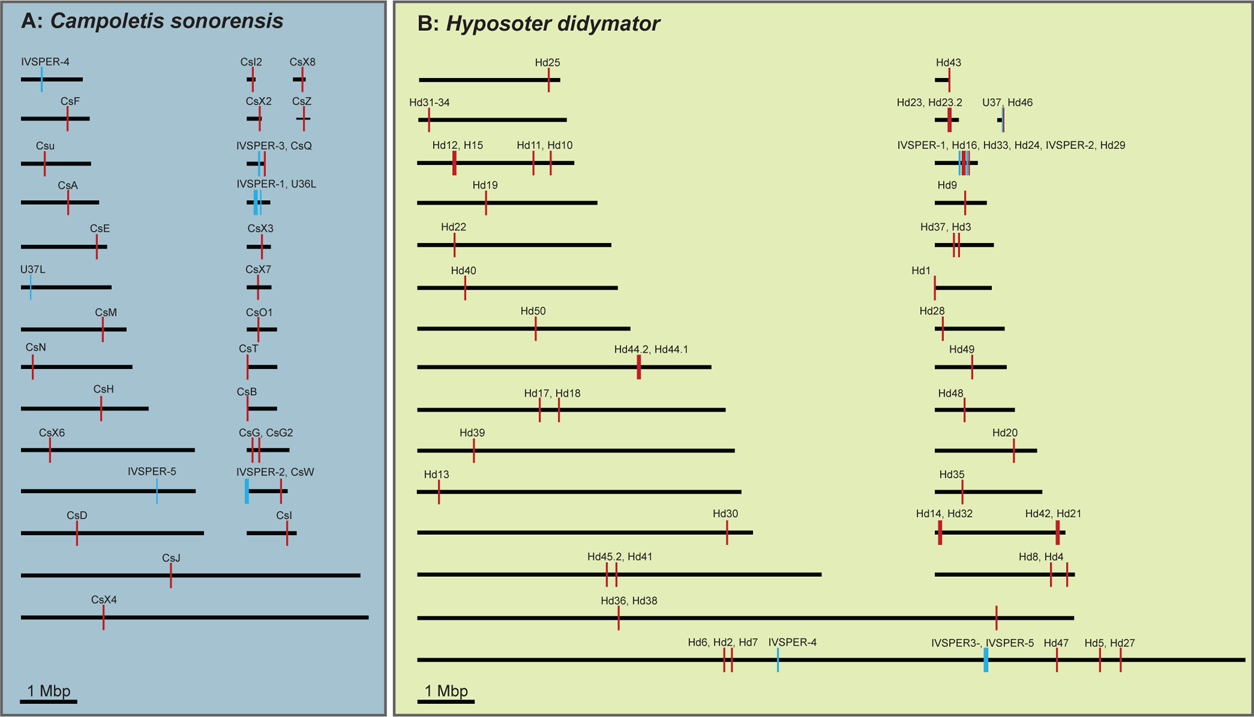
Distribution of ichnovirus sequences within *Campoletis sonorensis* and *Hyposoter didymator* genomes. **A.** Schematic representation of ichnovirus sequences within *C. sonorensis* scaffolds. Segments CsP and CsL, located in short scaffolds, are not shown. **B.** Schematic representation of ichnovirus sequences within *H. didymator* scaffolds. Segments Hd45.1 and Hd51, located in short scaffolds, are not shown. See Additional file 3 for more details.

**Table 4.** Summary of the number of viral loci identified in the genomes of *Campoletis sonorensis* (Cs) and *Hyposoter didymator* (Hd), in comparison with data available in NCBI database. For *H. didymator*, “merge segments” correspond to viral loci that included 2 segments former deposited in NCBI as distincts, “duplicated segments” those found in two copies in the wasp genome (see Figure 3 for more details). Number of IV segment loci are presented in the upper part of the table; number of IVSPERs are given in the lower part of the table. Number of segments and IVSPER in NCBI corresponds to the sequences deposited in NCBI that were available before this study.

|  | **Cs** | **Hd** |
| --- | --- | --- |
| **Number of segments in NCBI** | 25 | 50 |
| **Number of segments in genome** | 31 | 54 |
| **NCBI segments not found in genome** | 2 (CsA2, CsK) | 0 |
| **Merged segments (compared to NCBI)** | 0 | 6 (Hd2, Hd11, Hd17, Hd20, Hd26, Hd31-34) |
| **Duplicated segments (compared to NCBI)** | 0 | 3 (Hd23, Hd44, Hd45) |
| **Newly identified segments** | 8 (CsX1 to CsX8) | 6 (Hd46 to Hd51) |
| **"Isolated" segment genes (short scaffolds)** | 2 | 1 |
| **Total number of segment loci in genome** | **33** | **55** |
| **Number of IVSPERs in NCBI** | 0 | 3 |
| **Number of IVSPERs in genome** | 5 | 5 |
| **NCBI IVSPERs not found in genome** | na | 0 |
| **Newly identified IVSPERs** | 5 | 2 |
| **"Isolated" IVSPER genes** | 2 | 1 |
| **Total number of IVSPER loci in genome** | **7** | **6** |

In the *H. didymator* assembled genome, a total of 60 viral loci were identified (Figure 2B); they were located in 32 scaffolds ranging in size from 1.5 Kbp to over 15 Mbp (Additional file 3). A total of six IVSPERs were identified, which size varies from 1.6 Kbp to 26.6 Kbp; these *H. didymator* viral regions included three novel IVSPERs (more precisely two clusters and an isolated gene), for a total of 54 IVSPER predicted genes (Additional file 4). All the HdIV segments previously described in HdIV packaged genome [30] were identified in the wasp genome (Table 4). Some pairs of previously described segments actually co-localized at the same locus (in most cases, the segments shared part of their sequence, Figure 3A). Four segments were present in two copies (Figure 3B), three had copies in two different scaffolds, one was tandemly duplicated (Hd9). Finally, six totally new segments were identified in *H. didymator* genome. Altogether, 55 HdIV proviral segments were found, ranging in size from 2.0 to 17.9 Kbp (Additional file 3).

**Figure 3.**
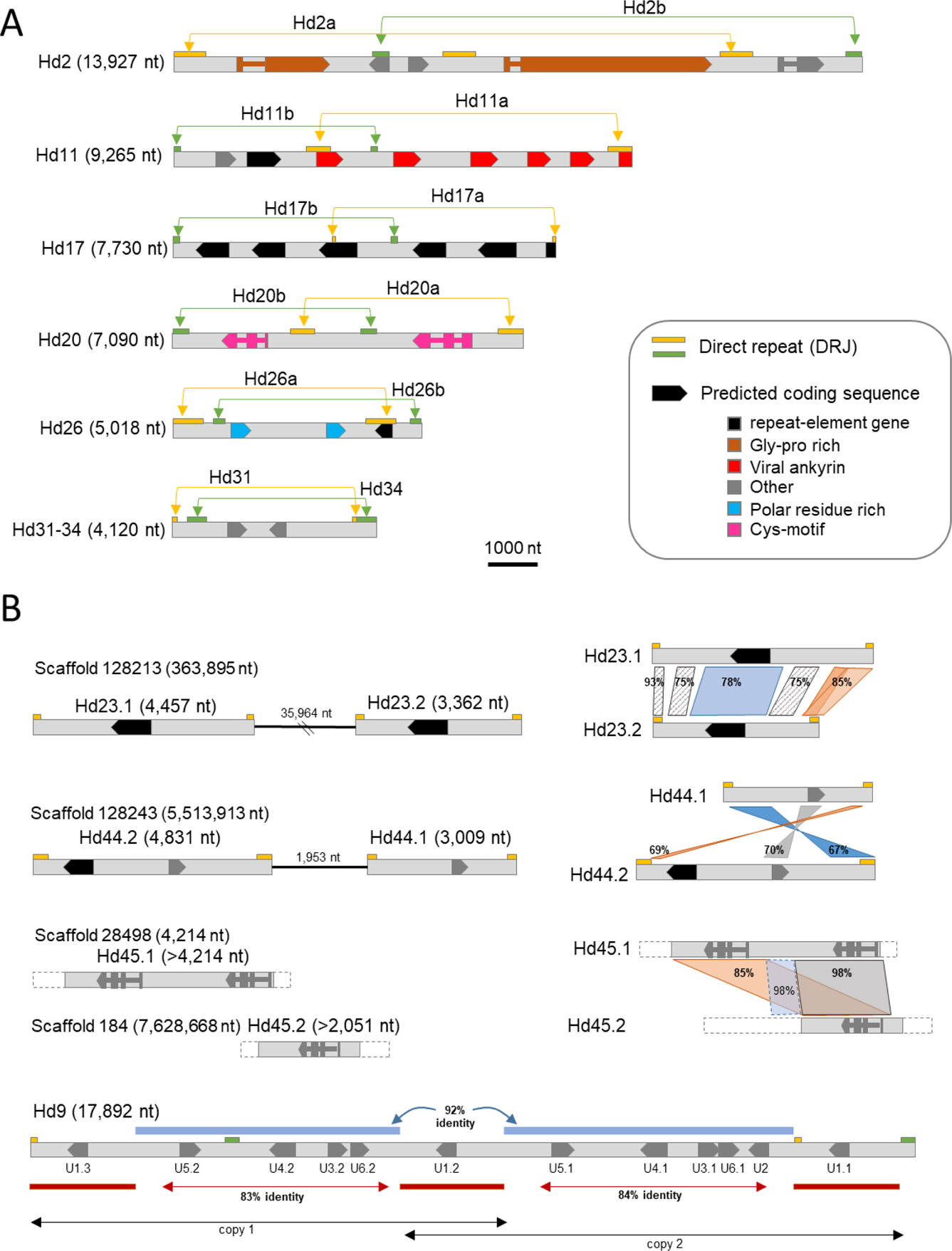
*Hyposoter didymator* nested and duplicated viral segments. **A.** Genomic architecture of the *H. didymator* ichnovirus (HdIV) segments described as distinct in Dorémus et al., 2014 but found located in a single wasp genomic locus in the present work. In Dorémus et al., 2014, segments with overlapping sequence were named Hd(n)a and Hd(n)b; they actually consist in a single locus here named Hd(n). **B.** Segments duplicated in *H. didymator* genome. Nucleotide percentage identity between the segment sequences is given on the right part of the figure. Hd23, Hd44 and Hd45 have two copies (named Hd(n).1 and Hd(n).2) that are either in the same scaffold but in different insertion sites or in two different scaffold; by contrast, Hd9 corresponds to a single viral loci containing an internal duplication (named “copy 1” and “copy 2” in the diagram).

Whole wasp genomes sequencing thus reveals a very large number of viral loci widely dispersed in these genomes. DNA fragments of viral origin are separated by large portions of wasp sequences, with a median size of 115.1 Kb between segments for those located on the same scaffold (Figure 4A). To confirm by an independent approach that IV proviral segment sequences were dispersed across the wasp genome, a FISH experiment was conducted for *H. didymator*, using genomic clones enclosing viral sequences as probes. Four probes were used, containing respectively segments Hd11, Hd6, Hd30 and Hd29, all in different genomic scaffolds. Results show that each of the probes hybridized with a different chromosome (Figure 4B), indicating that HdIV segments are indeed widely dispersed across the genome of *H. didymator*.

**Figure 4.**
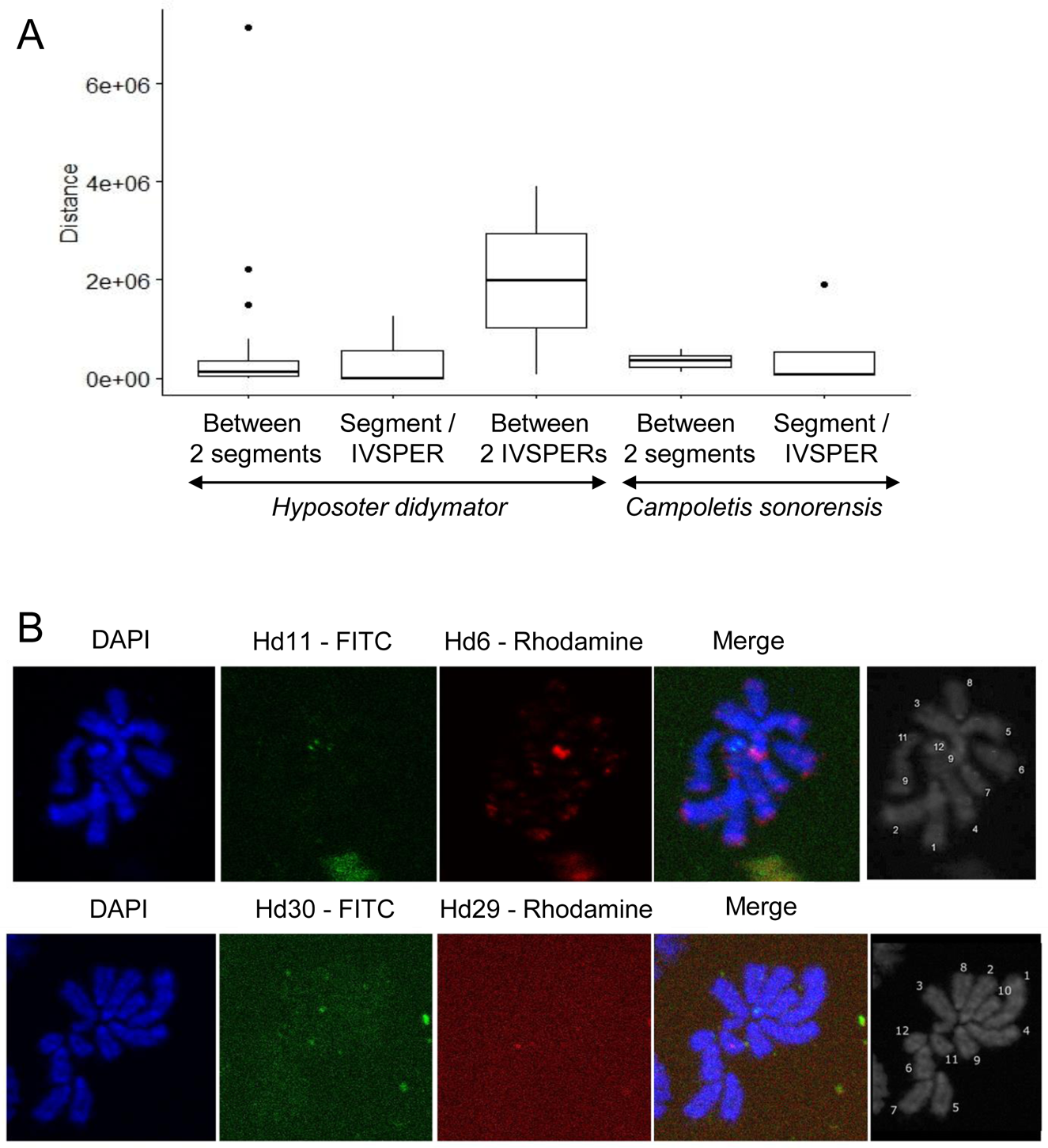
Dispersion of the viral segments in the wasp genome. **A.** Distance (in Kbp) between viral loci in *H. didymator* and *C. sonorensis* genomes. Data are given between 2 segments, between a segment and an IVSPER, and/or between 2 IVSPERs. **B.** FISH on *H. didymator* chromosomes using BAC genomic clones containing HdIV segments as probes (see material and methods part). Upper panels show hybridization using the probes containing viral segments Hd11 (labeled with FITC) and Hd6 (labeled with rhodamine); lower panel the probes containing viral segments Hd30 (labeled with FITC) and Hd29 (labeled with rhodamine). Each of the probes hybridized with a different *H. didymator* chromosome: Hd11 hybridized with the shortest chromosome (#12) whereas Hd6 mapped to a medium-sized chromosome (potentially #5); Hd30 hybridized with a large chromosome (#2) and Hd29 with a shorter chromosome (#11).

To assess whether dispersion of the viral loci could have been mediated during genome evolution by transposable elements, distribution of TEs was investigated in the regions surrounding the proviral segments. The analysis of the families of transposable elements in the regions surrounding the proviral segments (Additional file 5) did not reveal any particular enrichment that could suggest a role of TEs in the dispersion of the IV sequences in the wasp genomes.

### Ichnovirus DRJs show variable architecture and multiple excision sites

Repeated sequences flanking the proviral segment (or DRJ, for direct repeat junction) were found for all HdIV segment loci, except for Hd45.1 and Hd45.2. HdIV DRJs varied largely in size, ranging from 69 bp to 949 bp (Additional file 6). Similarly, most CsIV segments (25 of 32) were flanked by DRJs, which ranged in size from 99 bp to as much as 1,132 bp (Additional file 6). The number of direct repeats for a given proviral sequence was also variable (Figure 5A, Additional file 6). The majority of the HdIV (28) and CsIV (19) segments contained a single direct repeated sequence, one copy located on their right and left ends (named DRJ1R and DRJ1L, respectively; Figure 5A, a). A few HdIV and CsIV segments also contained internal repeats of the same sequence (hence named DRJ1int), potentially allowing the generation of more than one related circular molecules by recombination between the different DRJ1 copies (nested segments). Other IV proviral segments (21 HdIV segments, but only one CsIV segment) contained two different repeated sequences, named DRJ1 and DRJ2 (Figure 5A, b), combined or not with internal DRJs (Figure 5A, c; Additional file 6). Presence of several repeats differing in sequence and in position suggests the possibility that a mixture of overlapping and/or nested segments may be generated by homologous recombination in this context.

**Figure 5.**
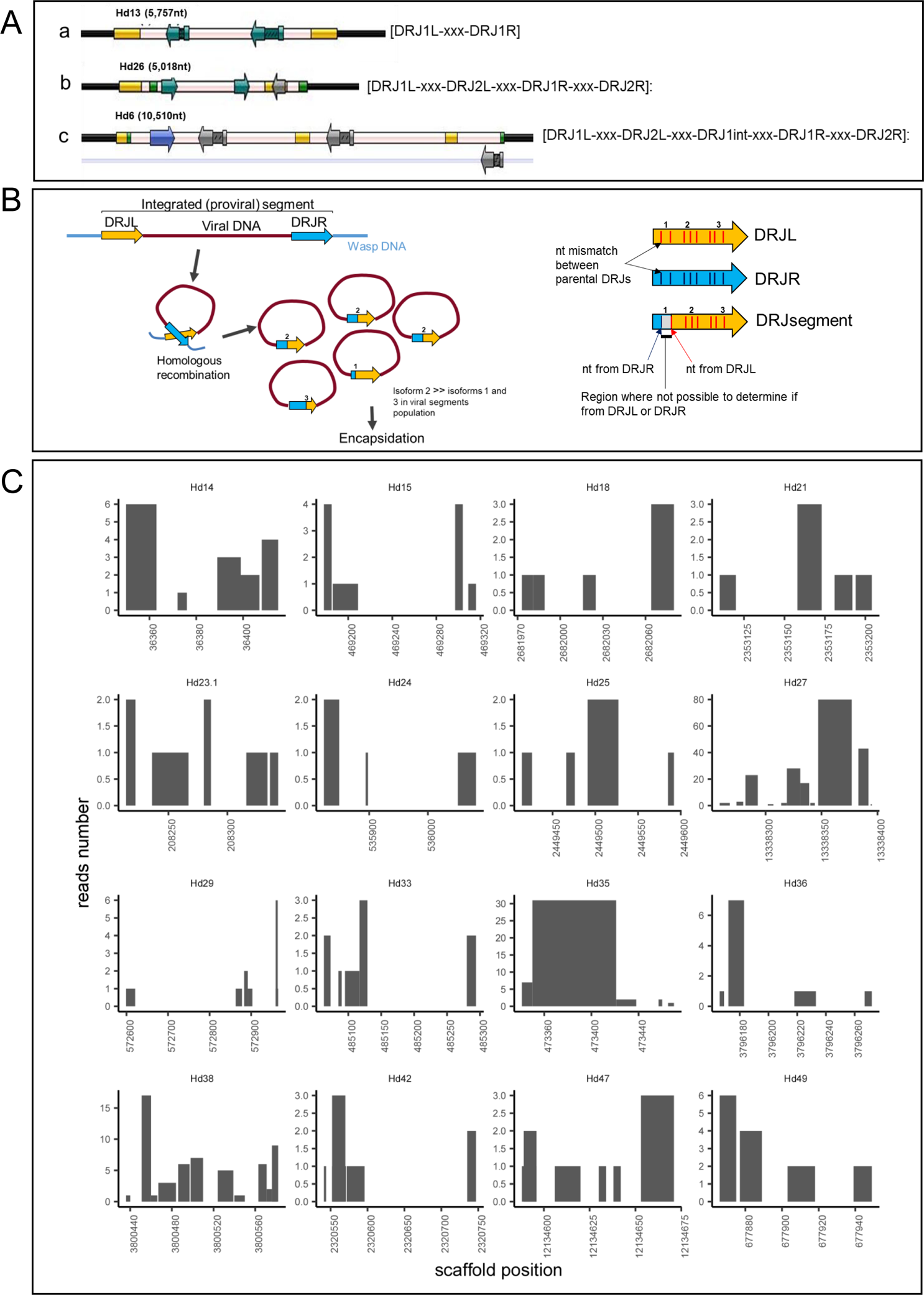
Segment DRJ variability in terms of number per segment, size and excision sites in *Hyposoter didymator*. **A.** Examples of the different types of DRJ position. **a.** Segment with two copies of a single direct repeat (DRJ1L and DRJ1R), each at one end of the segment. **b.** Segment with two distinct repeated sequences (DRJ1, in yellow and DRJ2, in green), each present in two copies (DRJ1L and DRJ1R, DRJ2L and DRJ2R). **c.** Segment with two repeated sequences, each present in two or more copies. DRJ1s in yellow, DRJ2s in green, HdIV genes represented by arrows. **B.** Schematic representation of the homologous recombination between the two DRJs flanking the proviral sequence (DRJL and DRJR) to produce the circular molecules (segment) containing one recombined DRJ sequence. The excision sites being located at different positions in the DRJ, segments differing in their recombined DRJ sequence are generated (isoform 1, 2 or 3 in the diagram). Excision occurs more frequently at some positions, resulting in different relative amounts of each isoform (isoform 2 more frequent than isoforms 1 and 3 in the diagram). The inset describes the rationale of the algorithm developed to identify the “breaking points”. Mapping of the segment sequence (DRJsegment) with the two parental DRJs - that differ in their sequences (nucleotide (nt) mismatches) - allows to identify the regions where occurred the switch from one parental DRJ to the other (in the diagram, the switch occurred between the first and second mismatch). **C.** Prediction of putative recombination breaking points in *H. didymator* DRJs. Each graph corresponds to the left copy of the DRJ for a given segment. The X axis is the position in the scaffold. The Y axis indicates the number of reads (obtained from sequencing of the packaged circular DNA molecules) confirming that the circle has been recombined at this position, based on the observed mismatches at both end of the segment for each reads.

We used the DMINDA webserver [33] to search for conserved excision site motifs embedded in IV DRJ sequences, using the all set of DRJs (99 DRJs) available for the two wasp species. Some motifs were found, in particular one that occurred at least once in almost all the analyzed DRJs (Additional file 7A). However, this motif may occur several times within a single DRJ; furthermore, a search across the whole *H. didymator* genome revealed that there was not a significantly higher chance for this motif to occur in the DRJ rather than in the rest of the wasp genome (Additional file 7B). Hence, circularization of IV segments does not seem to rely on the presence of a conserved nucleotide motif.

The two copies of the DRJ present at each end of the segment in the linear integrated form exhibit some punctual differences, so the excision site or breakpoint can be identified in a given recombined DRJ sequence with more or less resolution depending on the divergence between the parental DRJ copies. To identify potential excision sites in IV circular molecules, we analyzed two sets of *H. didymator* segment sequences using the DrjBreakpointFinder method developed for this purpose (see Methods section). The automatic analysis of a large set of recombined DRJ sequences revealed that excision could occur in different sites within a same DRJ (Figure 5B, 5C). This finding was confirmed by manual analysis of a subset of 8 segments sequenced using Sanger technology (Additional file 7C). Interestingly some positions of excision sites appeared more frequent than others for a given DRJ (Figure 5B, 5C).

### Proviral sequences serving as template for IV packaged genome show species-specific features

The number of proviral loci identified in *H. didymator* genome is higher (n=54) than in *C. sonorensis* genome (n=33). Altogether, IV segment loci represent a total size of 307.1 Kbp for HdIV and 314.1 Kbp for CsIV. The total sizes of the endogenous viruses are slightly underestimated since two HdIV segments (Hd1 and Hd45.1) and five CsIV segments (CsV, CsX3, CsX4, CsX5 and CsX7) were only partially identified because of the fragmentation of the genomes (see Additional file 3 for details). CsIV segments are globally longer compared to HdIV segments (Figure 6 A, B); they enclose 111 predicted genes whereas a total of 152 genes were predicted in the HdIV segments (Table 5, Additional file 4). Both encapsidated genomes contain a similar number of genes considering the IV-conserved multimembers families (repeat-element genes, vankyrins, vinnexins, cys-motif and N-genes (Table 5). HdIV contain more viral innexins, whereas CsIV more viral ankyrins and repeat-element genes (Table 5).

**Figure 6.**
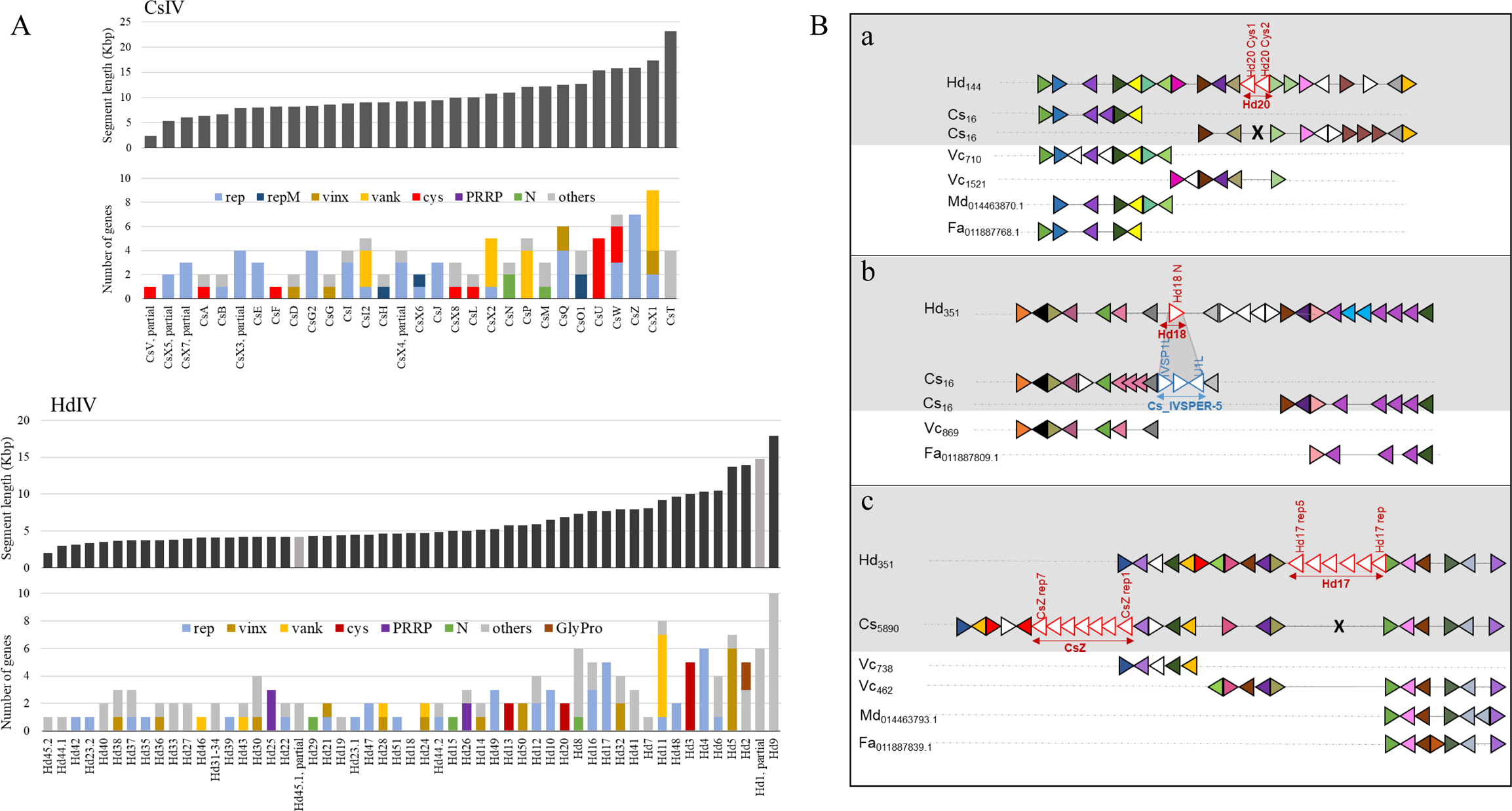
Comparative analysis of *Campoletis sonorensis* and *Hyposoter didymator* viral segments. **A.***C. sonorensis* ichnovirus (CsIV) segment size and gene content, i.e. the number of genes of each multigenic family found per segment. **B.** *H. didymator* ichnovirus (HdIV) segment size and gene content. For A and B: rep, repeat element genes; repM, repeat element genes with multiple repeated elements; vinx, viral innexin; vank, viral ankyrin; cys, cys-motif rich protein; PRRP, polar residue rich protein; N, N gene; Gly-Pro, glycine-proline rich protein. **C**. Synteny of *H. didymator* genomic regions where viral segments are inserted compared with *C. sonorensis* and other parasitoid genomes. **a.** Example of a syntenic region where only *H. didymator* genome presents a viral segment insertion. *H. didymator* genes from HD005010 to HD005030. **b.** The unique case found of a syntenic region where a viral segment in *H. didymator* and an IVSPER in *C. sonorensis* are inserted in the same position. *H. didymator* genes from HD010552 to HD010574. **c.** The unique case found of a syntenic region where a viral segment is inserted in both *H. didymator* and *C. sonorensis* genomes, but in two different positions. *H. didymator* genes from HD010503 to HD010526. Hd: *Hyposoter didymator*; Cs: *Campoletis sonorensis*; Vc: *Venturia canescens* (ichneumonid that has lost the ichnovirus ([62]); Md: *Microplitis demolitor* (braconid with a bracovirus); Fa: *Fopius arisanus* (braconid with virus-like particles). Numbers following the species name correspond to scaffold number for Hd, Cs and Vc, NCBI project codes for Md and Fa). Triangles within genomic regions correspond to predicted genes; triangles of the same color correspond to orthologs; white triangles are singletons or orphan genes. For better visualization, the name of the gene is indicated only for some viral (in red for segments, in blue for IVSPERs) genes. See additional file 8 for *H. didymator* genes list.

**Table 5.**
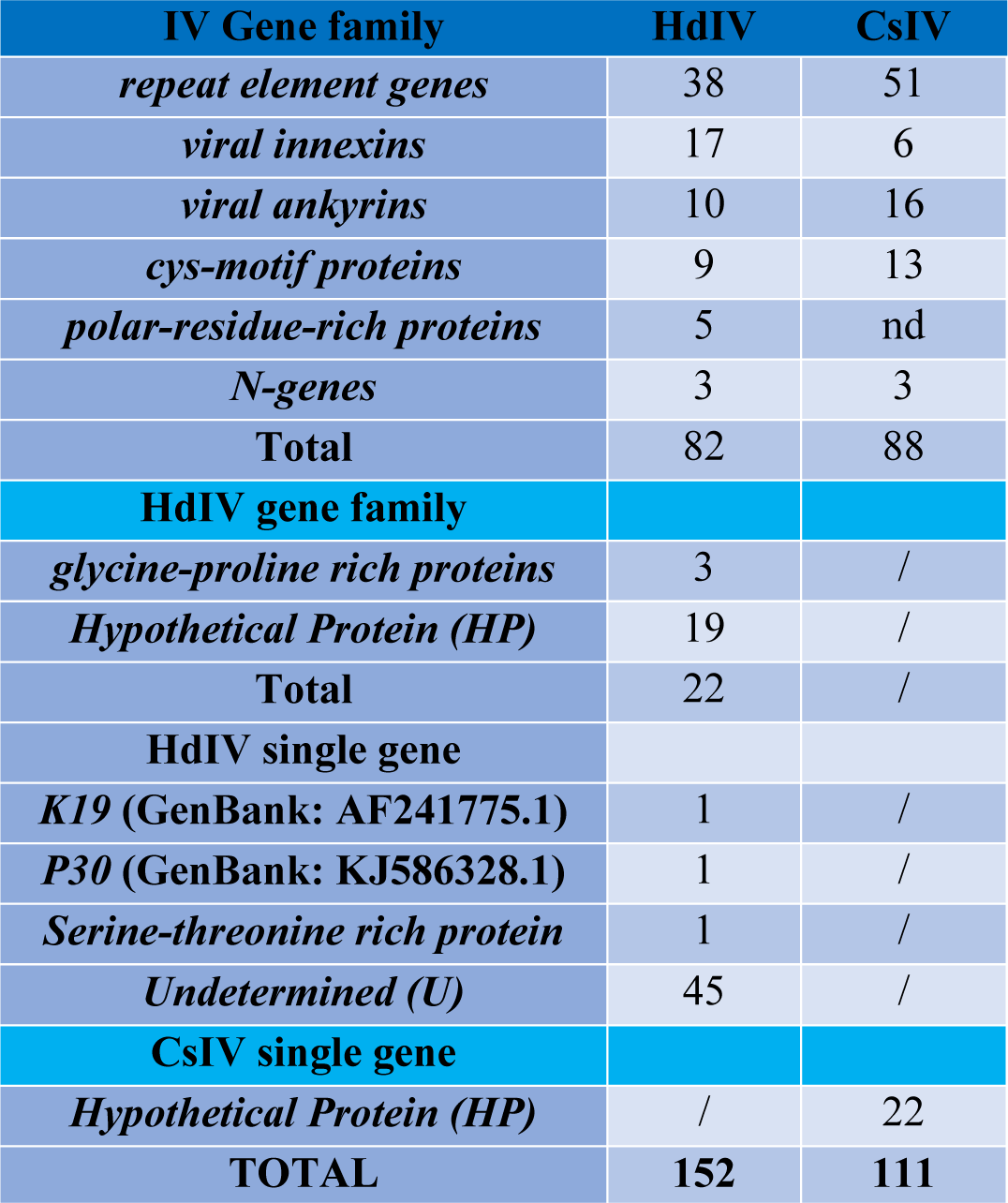
Comparative gene content of the IV segments in the ichnovirus carried by the campoplegine wasps *Hyposoter didymator* (HdIV) and *Campoletis sonorensis* (CsIV).

The high collinearity in gene order observed between the genomes of *H. didymator* and *C. sonorensis* made it possible to assess if the viral insertions were located in the same genomic environment for the two species. In order to accomplish this, the genomic regions containing viral insertions in *H. didymator* were compared to their syntenic genomic regions in *C. sonorensis* (Figure 6C). For the large majority of syntenic blocks containing an HdIV segment, there was no viral insertion in the corresponding *C. sonorensis* block (Figure 6C, a). Two exceptions were found. The first is the insertion site of *H. didymator* segment Hd18 which is located in the same genomic environment as the one where IVSPER-5 was inserted in *C. sonorensis* genome (Figure 6C, b). The second concerns *H. didymator* segment Hd17, inserted in a wasp genomic region that corresponded to the region, but not the insertion site, where *C. sonorensis* segment CsZ was inserted (Figure 6C, c).

### The ichnovirus machinery retained in wasp genomes (IVSPERs) is well conserved

A total of 45 different predicted IVSPER gene families were identified in the genomes of *H. didymator* and *C. sonorensis* (Figure 7A, Additional file 4). The majority (35, or 78%) are shared by both wasp species (Figure 7A) and 64% (29/45) are also shared with the banchine *Glypta fumiferanae* (Figure 7A). Amongst the 36 different genes/gene families (corresponding to a total of 48 genes) identified in the *C. sonorensis* genome, only one had no homolog in *H. didymator* genome. This gene, named Gf_U27L, has similarities (BlastP e-value 1E-31) with an IVSPER gene described in the banchine wasp *G. fumiferanae* [28]. On the other hand, amongst the 44 distinct genes/gene families identified in *H. didymator* genome (corresponding to 54 genes), nine were not detected in the *C. sonorensis* genome. Amongst them, three genes, U29, U32 and U33, are newly described for *H. didymator*. They were classified as IVSPER genes because of their localization in IVSPER-4 and, based on the transcriptome data generated for genome annotation, they are transcribed in *H. didymator* ovarian tissue. Note that among the homologous genes shared by *H. didymator* and *C. sonorensis*, the number of gene copies within some multigene families differs between the two wasp species (Figure 7A).

**Figure 7.**
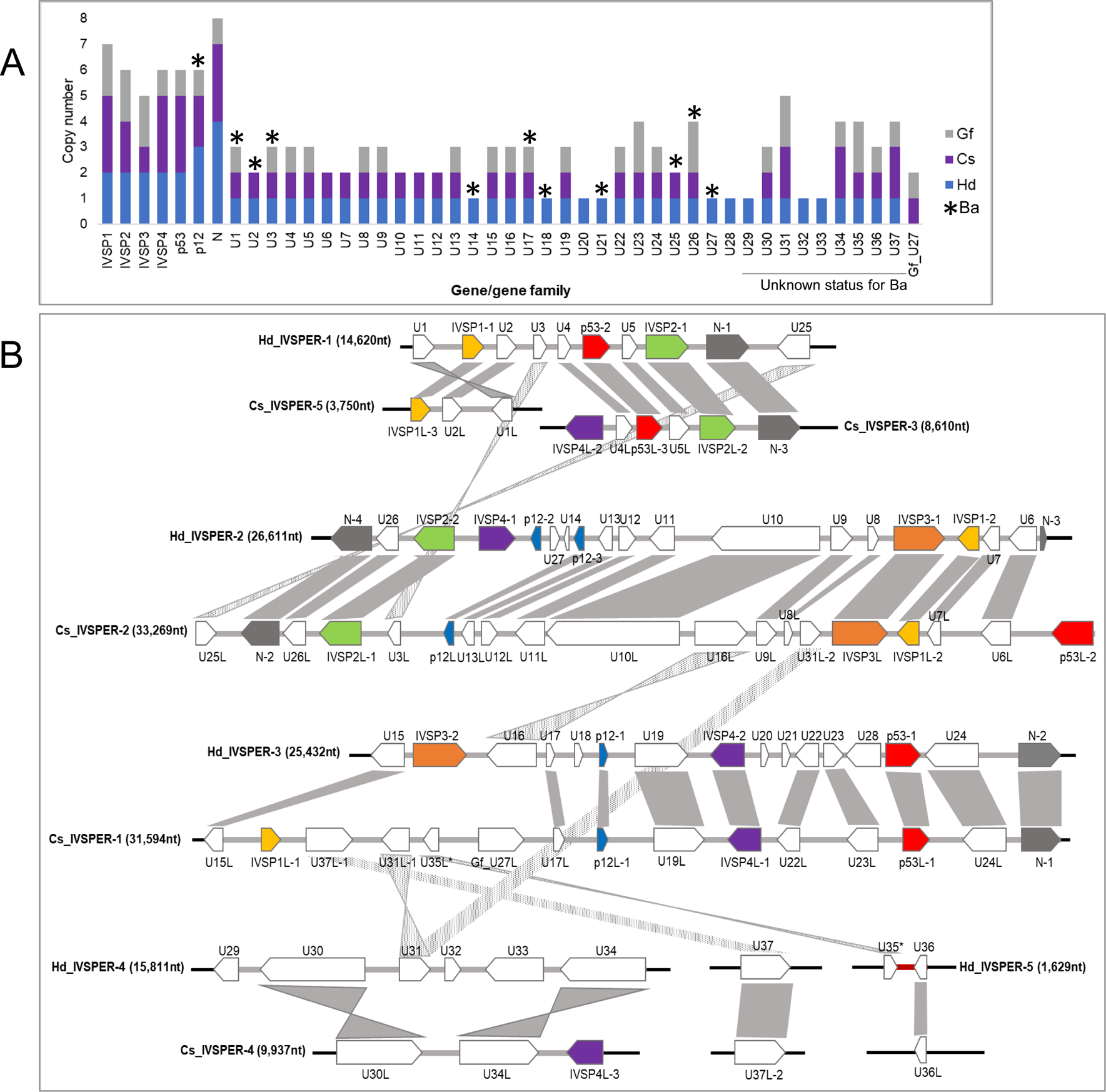
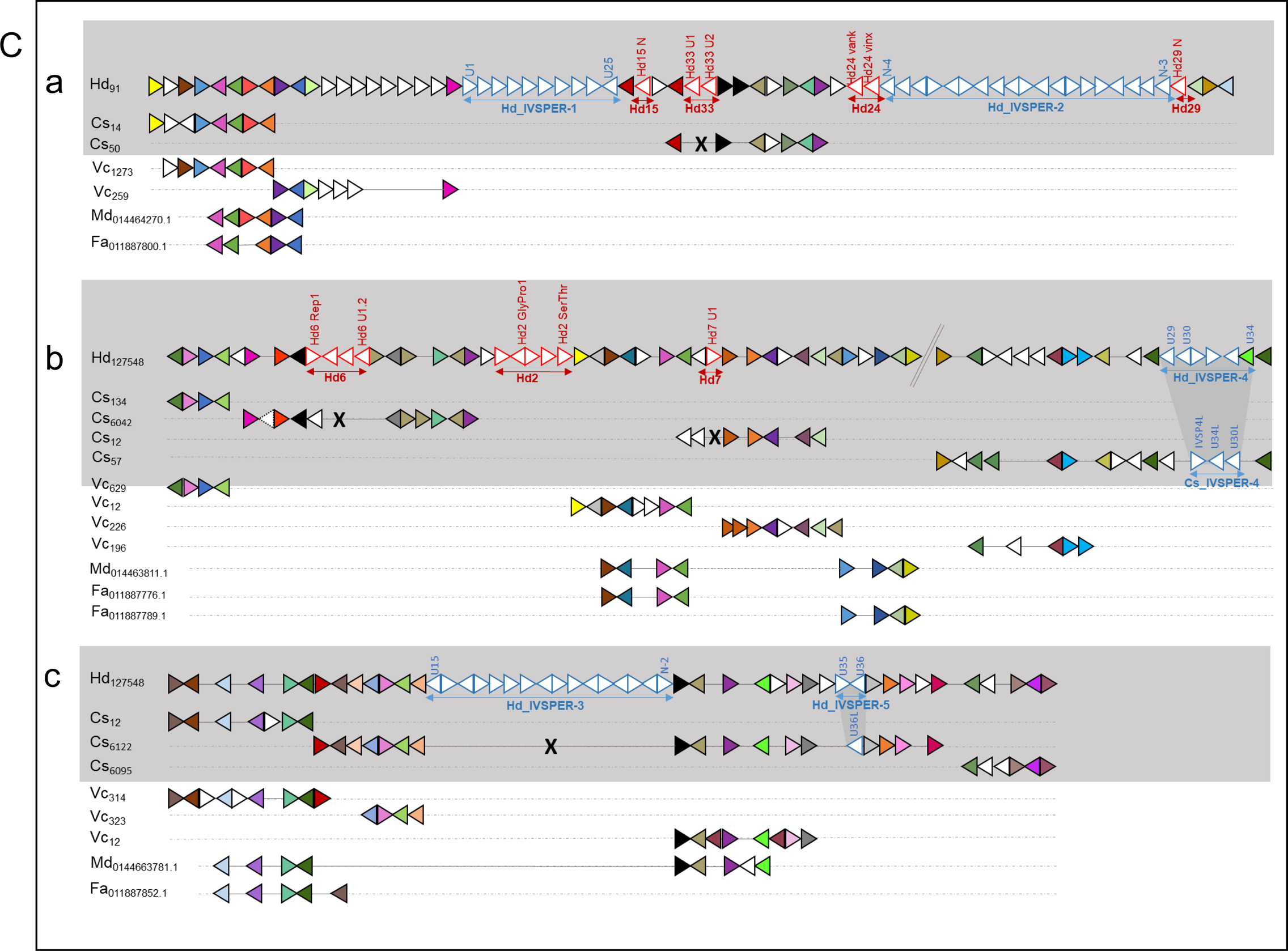
Comparative analysis of *Campoletis sonorensis* and *Hyposoter didymator* IVSPERs. **A.** List of IVSPER genes identified in four ichneumonid species. Hd, *Hyposoter didymator*; Cs, *Campoletis sonorensis*; Gf, *Glypta fumiferanae* (Banchinae) (from Béliveau et al., 2015); Ba, *Bathyplectes anurus* (Campopleginae) [74]. For the later, the number of gene copies in the genome is not known (presence of a gene copy indicated by an asterisk). **B**. Synteny between the IVSPERs identified in *H. didymator* (Hd) and *C. sonorensis* (Cs) genomes. *H. didymator* (Hd), where IVSPERs were first described is used as a reference. **C.** Synteny of *H. didymator* genomic regions containing IVSPERs compared with *C. sonorensis* and other parasitoid genomes. **a.** Synteny for *H. didymator* genomic region containing IVSPER-1 and −2 (genes from HD016092 to HD016153); no *C. sonorensis* scaffold corresponded to the *H. didymator* IVSPER insertion sites. **b.** Synteny for *H. didymator* genomic region containing IVSPER-4 (genes from HD001703 to HD001771); *H. didymator* IVSPER-4 and *C. sonorensis* IVSPER-4 are inserted in the same genomic environment. **c.** Synteny for *H. didymator* genomic region containing IVSPER-3 and −5 (genes from HD002066 to HD002111); in the region where *H. didymator* IVSPER-3 is inserted there is conservation in gene order compared to *C. sonorensis* but no viral insertion; conversely, *H. didymator* IVSPER-5 and *C. sonorensis* IVSPER-5 are inserted in the same genomic environment. Legend as in Figure 6. See additional file 8 for *H. didymator* genes list.

The IVSPERs of the two species show high synteny (Figure 7B). Synteny blocks with conserved gene order are shared for instance between Hd_IVSPER-2 and Cs_IVSPER-2 but also Cs_IVSPER-3 with part of Hd_IVSPER-1 and part of Cs_IVSPER-1 with Hd_IVSPER-3. Despite syntenic portions, there are some rearrangements, inversions and deletions between the two species. The highest number of rearrangements concerns the Cs_IVSPER-1 and Cs_IVSPER-4. The homologs of Cs_IVSPER-1 genes are dispersed in several different *H. didymator* IVSPERs (U15, IVSP1, U37, U31, U35). Similarly, *H. didymator* U3 and U25 have orthologs in Cs_IVSPER-2.

The genomic regions containing *H. didymator* IVSPER loci (Figure 7C) corresponds in most cases to synteny blocks where the gene order is well conserved compared with *C. sonorensis*; notably, this gene order is also quite well conserved in the IV-free campoplegine wasp *V. canescens*, and to some extent in the braconid wasps *M. demolitor* and *F. arisanus* (Figure 7C). For two of the three *H. didymator* genomic regions containing IVSPERs, some viral insertion sites were conserved between *H. didymator* and *C. sonorensis*. Regarding the insertion site of Hd_IVSPER-1 and −2 (Figure 6C, a), no corresponding *C. sonorensis* scaffold was found, making the comparison inconclusive. However, the comparison of the two other *H. didymator* genomic regions with their corresponding *C. sonorensis* syntenic blocks (Figure 7C b and c), revealed conservation of the insertion site for IVSPER-4 and IVSPER-5, whereas *H. didymator* and *C. sonorensis* IVSPER-3 are located in different wasp genomic regions.

## Discussion

The genome assemblies described here are of high quality with contig N50 over 150,000 bp and scaffold N50 from 1 to 4 Mbp. The annotated gene sets are similar in terms of numbers of genes and estimated completeness compared to genomes of other parasitic wasps. The assembled genomes allowed us to perform a comprehensive mapping of the viral inserts into the wasp genome.

The first major finding is the dispersion of the ichnovirus loci. This was particularly true for the IV proviral segments. All the 32 CsIV segments loci are located in different scaffolds, except CsG and CsG2, present in the same scaffold. For HdIV, half of the 54 viral segments are located in different scaffolds and for those located in the same scaffold, they are most frequently separated by relatively long portions of wasp genome. This dispersion was confirmed by FISH experiments, showing that viral loci are distributed across various chromosomes. We therefore did not find in the ichneumonid genomes any viral macro-locus as reported for bracoviruses [16]. Indeed, in braconid genomes, the BV segments are for the most part clustered in a unique locus gathering tens of segments, organized in replication units. For instance, the braconids *Glyptapanteles indiensis* and *G. flavicoxis* genomes contain 29 proviral segments, with 70% of them residing in a single large macrolocus [34]. Similarly, the *M. demolitor* genome contains 26 proviral segments; they are organized in 8 replication units located at 8 loci, loci 1 containing 14 segments [27]. In stark contrast, IV genomes consist of a series of single viral segments scattered within the wasp genome. In addition, as no enrichment of transposable elements in the surrounding of the IV segments have been observed, their dispersal in the wasp genome may result either from a diffuse integration of the virus ancestors or multiple genomic rearrangements events.

As previously described, the proviral segments are sequences that are packaged and transferred to the parasitized host whereas the IVSPER genes correspond to the “replication” genes, involved in the production of the virus particle. Until now, replication genes were known solely for one campoplegine wasp, *H. didymator* [26], and one banchine, *G. fumiferanae* [28]. Our study discovered replication genes in the *C. sonorensis* genome and showed a conserved IVSPER architecture for the two campoplegine wasps. In both *H. didymator* and *C. sonorensis* genomes, the majority of the replication genes are clustered. Indeed, only two isolated genes were identified in the *C. sonorensis* genome and only one in the *H. didymator* genome. In addition, *H. didymator* and *C. sonorensis* contain common genes arranged in a quite well conserved order. Noteworthy, most campoplegine IVSPER genes are also present in the banchine *G. fumiferanae*; however, the gene order is less conserved when comparing the two wasp subfamilies [9]. The high proportion of shared IVSPER genes between campoplegine and banchine wasps, including the D5 primase-like and DEDXhelicase-like first described in *G. fumiferanae* (corresponding to U37 and U34 respectively in *H. didymator*), tend to favor the hypothesis of a common virus ancestor for campoplegine and banchine IVs. This is quite notable, considering that the two subfamilies do not form a monophyletic group [35, 36] and IVs are not reported for other subfamilies in the same lineage. The divergences in IVSPER gene order between campoplegine and banchine reflects the phylogenetic distance between the species, with the viral loci evolving by genomic rearrangements specific to each of the ichneumonid subfamilies. Better understanding of the evolutionary trajectories of IVSPERs across ichneumonid lineages requires additional sequencing of banchine wasp genomes, as well as a thorough screening of species from other subfamilies for the presence of endogenous viruses.

Our study now provides the ability to make direct comparisons of viral composition between ichneumonid and braconid genomes. The genomes of campoplegine wasps associated with IVs contain numerous dispersed viral loci consisting of single viral segments and clusters of replication genes. By contrast, genomes of braconid wasps associated with BVs have clustered viral segments and more dispersed replication genes. For example, 76 nudiviral genes were identified in *M. demolitor*; half of these are located in a “nudiviral cluster” while the remainder are single genes located on different genome scaffolds [17]. This alternative genomic architecture probably reflects different regulatory mechanisms governing viral replication and particle production of the two PDV taxa. In braconids, the organization of the viral segments in replication units allow simultaneous co-amplification of several segments [21, 37] whereas in ichneumonids, each segment is individually amplified by a mechanism yet to be identified. IVSPERs DNA is also specifically amplified in the replicative ovarian tissue [26], what participates to increase gene copy number and thus to increase transcription levels. However, gene copy number is not the only mechanism involved in transcriptional control; IVSPER genes differ in their expression level in *H. didymator* calyx [38], suggesting that gene specific mechanisms are also involved in IVSPER gene expression regulation. In the case of bracoviruses, transcriptional control by nudiviral RNA polymerase allows expression of viral genes whatever their location, whereas amplification of the genes at the nudiviral locus allows achieving high levels of expression among nudiviral genes [25]. Although transcriptional control may not necessary require that genes are clustered in given regions, the clustering of replication genes in ichneumonids may facilitate coordinated regulation of their expression.

In *H. didymator* and *C. sonorensis* genomes, terminal and internal DRJs have been identified for most of the proviral segments. DRJs present at the extremities of the integrated form of the viral segment have been first described for CsIV [22] and the BVs associated with *Chelonus inanitus* [39] and *Cotesia congregata* [20]. The presence of internal repeats, which allows the generation of multiple circular molecules from the same proviral template in a process termed “segment nesting”, has also been reported, mainly for IV [22] and rarely for BVs [16].

In the case of bracoviruses, where segments are organized in a macro-locus, DRJs are thought to be involved downstream during replication. A first excision probably occurs thanks to sequences that limit the replication units in which there are also conserved sites [37]. Then the DRJs would serve to separate the different BV segments and generate the circular molecules. For ichnoviruses, where the proviral segments are individually amplified, there is no data presently on the limits of the replication units, making difficult to assess if DRJs are directly involved in the excision of the segment, or if there is also a two-step process involving other sequences.

Interestingly, some segments lack terminal repeats, mostly for CsIV (five segments of the 32 identified). This is particularly true for four CsIV segments newly identified; they are viral in nature since they all contain the IV characteristic repeat-element genes. The lack of one of their terminal repeats suggest they had lost their ability to be excised; this hypothesis would be consistent with the fact that they have not been revealed when the CsIV packaged genome was sequenced [29]. A similar finding was found in the braconid *Cotesia congregata* which harbors a pseudo-segment (CcBV pseudo-segment 34) mutated in the DRJ core which do not produce a packaged circle, whereas its homologous segment in *Cotesia sesamiae* produces a molecule packaged in the CsBV particle [16]. The fifth CsIV segment lacking DRJs is ShC, a segment that conversely to the four others, has been reported as part of the CsIV packaged genome [29]. In the present work, we found that ShC is embedded into IVSPER-2, i.e. corresponds to a region encoding replication genes which is, in addition, also conserved in *H. didymator* IVSPER-2 [26]. When we compared the ∼2 Kbp regions encompassing the pseudo ShC segment and flanking wasp sequences, no similarity was found (the only nucleotide motif repeated in IVSPER-2 was a 170 nt long motif located between U6L and U7L in one side, and within U25L on the other side). The absence of repeated sequence suggests that this sequence is not excised but part of an IVSPER.

Another important finding highlighted by our work is the documentation of inter-segment variability of DRJs in terms of sequence length, number and organization. Hence some HdIV segments harbored two different repeated sequences, and/or one or several internal repeats. This complexity suggests numerous possibilities to produce by intrachromosomal homologous recombination a series of related circular molecules that will be packaged in the particles. IV DRJs vary in size, ranging from less than 100 bp to more than 1 Kbp and in similarity rate, with identity between pairs of related DRJs varying from 70 to 98% (Additional file 6). The rate of homologous recombination between DNA fragments is affected by sequence length and divergence (reviewed in [40]); thus variability of IV segment DRJs length and homology rate suggests that homologous recombination efficacy may vary from one segment to the other. While for bracoviruses the DRJs contain a conserved tetramer AGCT embedded within a larger motif, which corresponds to the site of excision [17, 41], we did not identify a conserved motif specific to IV DRJs. On the other hand, an automatic search of the excision sites using the algorithm developed in the present study allowed us to posit various possible sites of excision and recombination within a given DRJ. Interestingly, some sites seemed to occur more frequently than others. However, for now, the mechanism and the proteins involved in DNA double-strand breaks and repair in IV sequences remains to be identified. In the case of bracoviruses, there are conserved DRJ sequences, and conserved members of the tyrosine recombinase family of nudiviral origin (vlf-1, int-1 and int-2) which seem involved in regulating excision [25]. In the case of ichnoviruses, there are neither conserved sites nor known virus-derived recombinases. For now, it is not possible to infer a function to many of the IVSPER genes, so it is not possible to determine if IV excision relies on the same mechanisms as bracoviruses; possibly, homologous recombination and generation of IV circular molecules rely on the wasp cellular machinery.

Two main conclusions arise from the comparison of the two genomes, giving insights on the evolutionary forces driving IV domestication in campoplegine wasps. At first, the genes potentially involved in producing the IV particles are very well conserved in terms of gene content and gene order (Figure 7). Hence, only nine of the 54 predicted replication genes in *H. didymator* are specific to this species; they correspond in majority to short sequences and may for some not be truly functional IVSPER genes, however they are all transcribed in calyx cells based on our transcriptome analyses. In both *H. didymator* and *C. sonorensis*, there is only a few clusters, and one or two isolated genes. Noteworthy, the isolated gene in *H. didymator* is U37, a homolog of the D5 primase gene described in the banchine *G. fumiferanae* where this gene is embedded within an IVSPER (Gf-IVSPER-1; [28]). One of the two U37 homologs in *C. sonorensis* is also isolated, but in a genomic context distinct from that in *H. didymator*, whereas the other is localized in Cs-IVSPER-1 (Figure 7). Gene order within the clusters is well conserved between *H. didymator* and *C. sonorensis*, even though these regions underwent rearrangements over evolution. Compared to the IVSPERs, the viral segments carrying virulence genes appear more divergent and species-specific, despite the existence of common gene families. The two components of the ichnoviral genome also differ in terms of conservation of their genomic localization. First, we did not find any viral segment inserted in a given block of synteny when comparing *H. didymator* and *C. sonorensis*. The situation of viral segments in ichneumonids therefore seems different from that described in braconids, where the viral segments remain in homologous positions, flanked by conserved wasp genes [16]. In ichneumonids, viral segment diversification and dispersion may result from transposition of each viral sequence in the wasp genome while for bracoviruses, segment multiplication occurs by duplication of large areas [16]). By contrast, two of the five IVSPERs were localized in the same genomic context in *H. didymator* and *C. sonorensis* (Figure 7C). These two IVSPER harbor related genes, what suggests a common ancestral origin. These loci may represent ancestral viral insertion sites that are still conserved in both wasp genomes. For the others, the lack of co-localization indicates that IVSPERs are able to move quite easily in the wasp recipient genomes. Note that we were not able to highlight transposable element (TE) enrichment close to viral insertions, suggesting that transposition of viral sequences may rely on another still unknown mechanism which might have been inherited from the ancestor virus.

The conservation of IVSPER genes is consistent with what would be expected from a functional point of view: they constitute the machinery allowing the wasp to produce the particles, i.e. the delivery systems on which the parasitoid relies for its survival. As such, mechanisms for viral replication and packaging are not expected to be subject to rapid change. Meanwhile, proviral segments carry virulence genes, which need to quickly respond to counter-adaptations arising in the immune system of the parasitoid host. Since campoplegines are koiniobiont endoparasitoids that often have a restricted host range [42, 43], proviral sequences are expected to evolve rapidly and in a species-specific manner. Finally, we did find that a same syntenic block contained a proviral segment in *H. didymator* and an IVSPER in *C. sonorensis*. In a scenario of a common origin between IVSPER and viral segments (i.e. the ancestral virus), this locus may represent an ancestral viral insertion site, which may have contained sequences corresponding to the complete ancestral virus genome before its separation in two components, the proviral segments and the replication gene clusters.

## Conclusions

Whole genome sequencing of two parasitoid wasps, *H. didymator* and *C. sonorensis*, which both harbor integrated ichnoviruses, is reported here for the first time. These are the first annotated full genomes available for the family Ichneumonidae. Most importantly, they allowed us to piece together a comprehensive picture of the architecture of ichnoviruses in the wasp genomes. Our results reveal an interesting duality between specific genomic features shown by the proviral elements of the polydnavirus genome and the conserved nature of the viral machinery within the wasp DNA. While this may be linked to the biological functions of these two types of genomic elements, this genomic organization is directly opposed to that observed in bracoviruses, highlighting how similar solutions to adaptive demands can convergently arise via very different evolutionary pathways. Tracing the steps that have led to the architecture of modern ichnoviruses from the domestication of an ancestral virus will require the sequencing of other ichneumonid wasps from multiple lineages.

## Methods

### Target species and insect rearing

Two species from the putatively monophyletic subfamily Campopleginae [35, 36] were chosen as target taxa for whole genome sequencing: *Hyposoter didymator* occurs in all Western Europe and mainly parasitizes *Helicoverpa armigera* [44] whereas *C. sonorensis* occurs from North to South America and parasitizes several noctuid species [45].

Specimens of *Hyposoter didymator* were reared on *Spodoptera frugiperda* larvae as previously described [30]. Male specimens of *Campoletis sonorensis* wasps were furnished from a colony maintained at the University of Kentucky and reared as described in Krell et al. [46].

#### C. sonorensis whole genome sequencing, assembly and automatic annotation

Genomic DNA was extracted from one single adult male using the Qiagen™ MagAttract HMW DNA kit, following the manufacturer’s guidelines. The resulting extraction was quantified using a Qubit™ dsDNA High Sensitivity assay, and DNA fragment size was assessed using an Agilent™ Genomic DNA ScreenTape. Sample libraries were prepared using 10X Genomics Chromium technology (10X Genomics, Pleasanton, CA), followed by paired-end (150 base pairs) sequencing using one lane on an Illumina HiSeqX sequencer at the New York Genome Center (Additional file 9A).

Assembly of the sequenced reads was conducted using Supernova v.2.1.1 [31]. Reads were mapped back to the assembled genome using Long Ranger (https://support.10xgenomics.com/genome-exome/software/pipelines/latest/what-is-long-ranger) and error correction was performed by running Pilon [47]. Note that Supernova recommends 56X total coverage and sequencing deeper than 56X reduces the assembly quality. A full lane of HiSeqX produced several times the sequencing data we needed to reconstruct the genome. Hence, raw reads were divided in four subsets and four separate assemblies were conducted in parallel, with the best one chosen for downstream analyses.

We used RepeatMasker [48], to identify repeat regions using Drosophila as the model species. *H. didymator* transcripts were aligned to the *C. sonorensis* genome using BLAT [49]. We created hints files for Augustus from the repeat-masked genome and the BLAT alignments. We also ran BUSCO v3 [50] with the –long option both to assess genome completeness and to generate a training set for Augustus. We then ran Augustus v3.3 [51] for gene prediction using the three lines of evidence, the RepeatMasker-generated hints, the BLAT-generated hints, and the BUSCO-generated training set.

#### H. didymator whole genome sequencing, assembly and automatic annotation

Genomic DNA was extracted from a batch of adult males (n=30). DNA extractions that passed sample quality tests were then used to construct 3 paired-end (inserts lengths = 250, 500 and 800 bp) and 2 mate pairs (insert length = 2,000 and 5,000 bp) libraries, and qualified libraries were used for sequencing using Illumina Hiseq 2500 technology (Additional file 9B) at the BGI. For genome assembly, the raw data was filtered to obtain high quality reads.

The reads were assembled with Platanus assembler v1.2.1 [32], in 2 steps (contigs assembly and scaffolding), then the scaffolds gaps were filled with SOAPdenovo GapCloser 1.12 [52]. Finally, only scaffolds longer than 1,000 bp were kept for further analyzes. Assemblathon2 [53] was used to calculate metrics of genome assemblies.

For annotation, EST reads from venom and ovaries, as well as reads obtained using GS FLX (Roche/454), Titanium chemistry from total insects [54] were mapped to the genome with GMAP [55], and Illumina reads published previously [38], or from the 1KITE consortium (http://1kite.org/subprojects.html) using STAR [56], and new calyx RNA-Seq (SRA accession: PRJNA590863) with TopHat2 v2.1.0 [57]. BRAKER1 v1.10 [58] was used to predict genes in the genome of *H. didymator* using default settings. Gene annotation was evaluated using Benchmarking Universal Single-Copy Orthologue (BUSCO) version 3.0.2 [50] with a reference set of 1,658 proteins (conserved in Insecta).

The other parasitoid genomes used in this work for comparison purposes were similarly analyzed (genomes available at NCBI for *Microplitis demolitor* [27] (PRJNA251518), *Fopius arisanus* [59] (PRJNA258104), *Diachasma alloeum* [60] (PRJNA306876) and *Nasonia vitripennis* [61] (PRJNA13660); *Venturia canescens* genome [62] available at https://bipaa.genouest.org/sp/venturia_canescens/).

#### Manual annotation of the viral loci

Manual annotation of viral regions was done using the genome annotation editor Apollo browser [63] available on the BIPAA platform (https://bipaa.genouest.org). The encapsidated forms of the HdIV and CsIV genomes were previously Sanger-sequenced by isolating DNA from virions [29, 30]. *H. didymator* IVSPER were also previously sequenced [26]. To identify the viral loci, sequences available at NCBI for campoplegine IV segments and IVSPER sequences from campoplegine and banchine species were used to search the *H. didymator* and *C. sonorensis* genome scaffolds using the Blastn tool implemented in the Apollo interface. To determine the limits of the proviral segments, we searched for direct repeats at the ends of the viral loci by aligning the two sequences located at each end using the Blastn suite at NCBI. The start or stop codons of the genes located at the ends of the IVSPER loci were considered as the borders of the IVSPER.

#### Transposable Element Detection

Transposable Elements (TEs) were identified in *H. didymator* and *C. sonorensis* genomes using the REPET pipeline [64, 65]. The enrichment analysis was performed using LOLA [66] comparing the observed number of each TE family member in a region encompassing IV segments and 10 kbp around, and 1,000 random segments of the same length extracted with bedtools shuffle [67].

#### Orthologous genes and syntenic regions

To identify homology relationships between sequences of *H. didymator*, *C. sonorensis* and other parasitoids with available genomes (*Venturia canescens*, *Microplitis demolitor*, *Fopius arisanus*, *Diachasma alloeum* and *Nasonia vitripennis*), as well as taxonomically restricted sequences (*Apis mellifera* and *Drosophila melanogaster*), a clustering was performed using the orthogroup inference algorithm OrthoFinder version 2.2.7 [68]. Sequences predicted by automatic annotation (Braker or Augustus) but also some resulting from manual annotations were used. Thus a total of 18,154 protein coding genes for *H. didymator* and 21,987 for *C. sonorensis* were included in the analysis (Table 4A). The syntenic blocks were reconstructed with Synchro [69] using the genomes and proteomes of the same species.

#### H. didymator genomic BAC library construction and sequencing

Genomic BAC clones were obtained as described in [26]. Briefly, high molecular weight DNA was extracted from *H. didymator* larval nuclei embedded in agarose plugs. The nuclei were lysed and the proteins degraded by proteinase K treatment. DNA was partially digested with *Hin*dIII. The size of the fragments obtained averaged 40 kbp as controlled by Pulse Field Gel Electrophoresis. Fragments were ligated into the pBeloBAC11 vector. High-density filters were spotted (18,432 clones spotted twice on nylon membranes) and screened using specific 35-mer oligonucleotides. Positive clones were analyzed by fingerprint and, for each probe, one genomic clone was selected and sequenced using Sanger technology (shotgun method) by the Génoscope, Evry, France. The sequences obtained were then submitted to a Blastn similarity search against NCBI nr database in order to confirm presence of HdIV sequences. Four BAC clones containing HdIV sequences were used as probes in FISH experiments (see below).

#### Fluorescent in situ hybridization (FISH) on H. didymator chromosomes

The *H. didymator* genome is composed of 12 chromosomes [70]. Karyotypes were prepared from male reproductive tracts from pupae and young adults. The testes were dissected in saline solution and placed in colchicine/colcemid solution (50ug / ml) during 10 minutes. After elimination of the liquid, a hypotonic solution (Na citrate 0.5%) was added during 10 minutes. The solution was then replaced by fixative (1 vol. acetic acid / 3 vol. methanol) and let to incubate during 40 minutes. The genitalia were then placed on a glass slide, a drop of acetic acid 60% was added to further shred the tissue and the slide was placed on a hot plate at 42°C until complete evaporation of the liquid. The samples were stained with DAPI and observed under a fluorescent microscope in order to select slides with sufficient and suitable caryotypes. Two genomic clones containing a viral sequence (CE-15P20 and CF-16G11) were used as probes. They were alternatively labeled using the Dig RNA labeling mix (Roche) or the biotin RNA labeling mix (Roche). For hybridization, the samples were rehydrated and denatured during 6 minutes by a 0,07 N NaOH treatment. The anti-digoxigenin antibody was labeled with rhodamine (Roche) (dilution 1/50) and the anti-biotin antibody with FITC (Vector laboratories) (dilution 1/200) overnight at 37°C. Images were captured on a Zeiss AxioImager Apotome microscope.

#### Re-sequencing of HdIV packaged genome

The viral DNA was extracted following the procedure described in Volkoff *et al*. [71]. Briefly, ovaries from about 100 female wasps were dissected in PBS and placed in a 1.5-ml microfuge tube. The final volume was adjusted to 500 μl using Tris-EDTA buffer and the ovaries were homogenized by several passages through a 23-gauge needle. The resulting suspension was passed through a 0.45-um pore-size cellulose acetate filter to recover the HdIV viral particles. For viral DNA extraction, the filtrate was submitted to proteinase K and Sarcosyl treatment overnight at 37°C, then to RNase A treatment 2 h at 37°C. DNA was further extracted with phenol/chloroform/isoamyl alcohol and precipitated with ethanol. The DNA pellet was re-suspended in ultra-pure water and was sequenced using GS FLX (Roche/454), Titanium chemistry (Eurofins Genomics). The obtained reads were used for DRJ excision site analyses (see below).

#### DRJs and breakpoint analysis in H. didymator

The proviral integrated segments are circularized and excised by homologous recombination between its extreme DRJs (at left and right extremities of the given segment). When the two copies of the DRJ exhibit some punctual differences, the excision site or breakpoint can be identified in a given recombined DRJ sequence with more or less resolution depending on the level of divergence between the DRJ copies.

In order to identify and analyze DRJ excision sites of a set of circularized IV sequences in an automatic fashion, we developed the following method, called DrjBreakpointFinder and freely distributed at http:// github.com/stephanierobin/DrjBreakpointFinder/. The method takes as input a set of circularized sequences (usually obtained by sequencing) and a reference genome. It is composed of two main steps. The first step consists in identifying triplets of sequences (read-DRJL-DRJR) representing the recombined DRJ and its two parental DRJs, by mapping the sequencing reads to the reference genome. In the second step, a precise multiple alignment is computed for each sequence triplet and a segmentation algorithm, inspired from the breakpoint refinement method Cassis [72, 73], is applied along the recombined DRJ sequence to identify in the best case scenario the excision site or more generally the breakpoint region.

To do so, the segmentation algorithm estimates the best partition of the recombined DRJ sequence into three distinct segments, corresponding to homology with DRJR, the breakpoint region and homology with DRJL respectively, given the repartition of punctual differences with the two parental DRJs. The segmentation algorithm is classically based on fitting a piecewise constant function with two changepoints to the punctual difference signal (see [73]). DrjBreakpointFinder further gathers breakpoint results by proviral segments or DRJ pairs, in order to obtain for each the distribution of potential excision sites observed in a given circular virus sequencing dataset. The output of DrjBreakpointFinder consists of breakpoint region coordinate files along with visual representations for each proviral segment or DRJ pair.

In this paper, DrjBreakpointFinder was applied to two circular viral DNA sequencing datasets. Circular DNA was extracted from HdIV particles and sequenced by 454 and Sanger technologies, resulting in 40,343 and 15,575 reads, respectively.

In addition, the DRJ copy was manually analyzed for a subset of 8 segments (Hd12, Hd16, Hd19, Hd22, Hd24, Hd28, Hd29 and Hd30) that presented only one right and left DRJs in their integrated form. Junctions were amplified by PCR using primers located within the viral sequence, downstream and upstream the DRJs. PCR products were cloned in pGEM and 3 to 5 plasmid clones were then sequenced using Sanger technology for each segment. The obtained recombined junction sequences were then aligned with the 2 parental DRJs in an attempt to localize the excision site, based on the nucleotides differing between the 2 DRJs (see Additional file 5, C).

## Declarations

### Ethics approval and consent to participate

Not applicable

### Consent for publication

Not applicable

### Availability of data and materials

The datasets supporting the conclusions in this article are available at the NCBI under the Bioproject accession numbers PRJNA589497 for *Hyposoter didymator* and PRJNA590982 for *Campoletis sonorensis*. The DrjBreakpointFinder method developed in this study is freely distributed at http:// github.com/stephanierobin/DrjBreakpointFinder. All other data are included within this article and its additional files.

### Author’s contributions

FL, SR and RD were responsible for the assembly of whole genomes and automatic annotations, and AB for database management. ANV annotated manually the endogenous viral sequences. BAW furnished *C. sonorensis* samples. BFS did the DNA extraction and QC for *C. sonorensis*. VJ did the *H. didymator* DNA extractions; XZ and SL supervised whole genome sequencing of this species. SR was responsible for the BUSCO and gene orthology analyses. FL was responsible for transposable element and synteny analyses. GG managed the *H. didymator* BACs sequencing. CL developed the pipeline for DRJ analyses and SR has performed the DRJ analyses. MR and VJ conducted the FISH experiments. ANV and BFS were responsible for the project conception, funding and management. ANV wrote the manuscript with large participation of BFS, FL and SR. DT, XZ, CL, SGB and JMD contributed to improve the manuscript. All authors read and approved the final manuscript.

## Acknowledgements

The authors thanks Drs Jing ZHAO and Ming TANG, from the BGI-Shenzhen, China, for their help with *H. didymator* genome sequencing. The Sackler Institute of Comparative Genomics (AMNH) funded the generation of preliminary data for *C. sonorensis* and made the final genome sequencing available at reduced cost through their partnership with the New York Genome Center. We thank Clotilde GIBARD and Gaétan CLABOTS from the DGIMI insectarium for the insects they have furnished, and the quarantine insect platform (PIQ), member of the Vectopole Sud network, for providing the infrastructure needed for experimentations on insects. The authors acknowledge Séverine CHAMBEYRON and Christine BRUN (IGH, Montpellier) for their great help in *H. didymator* FISH experiments. From the DGIMI lab, we also thank Isabelle DARBOUX for help with figure 4, Kiwoong NAM for constructive comments at the early stages of the MS writing, and Bertille PROVOST who performed the genomic DNA extraction for *H. didymator* BAC library preparation. We thank Mercer Brugler (NYC College of Technology) and Mark Siddall (AMNH) for their enthusiastic incentive and guidance at the early stages of this project.

## Table of content for Additional files

1. **Additional File 1 (word file)**. Orthogroups analyses.
2. **Additional File 2 (word file)**. Synteny blocks between pairwise comparisons of multiple parasitoid genomes.
3. **Additional File 3 (excell file)**. List of scaffolds in *Hyposoter didymator* and *Campoletis sonorensis* genomes containing at least one ichnovirus sequence.
4. **Additional File 4 (excell file)**. List of ichnoviral genes identified in *Hyposoter didymator* and *Campoletis sonorensis* genome scaffolds containing at least one ichnovirus sequence.
5. **Additional File 5 (excell file)**. Transposable elements (TE) found in *Hyposter didymator* segments, IVSPERs and neighboring regions.
6. **Additional File 6 (excell file)**. List of direct repeat junctions (DRJ) found in *Hyposoter didymator* and *Campoletis sonorensis* genome scaffolds.
7. **Additional File 7 (word file)**. DRJs analysis.
8. **Additional File 8 (excell file)**. List of the *Hyposoter didymator* genes present in the syntenic blocks represented in figures 6 and 7.
9. **Additional File 9 (word file)**. Characteristics of the libraries used for genome assembly.

**Additional File 1.** Orthogroups analyses. **A.** Orthofinder clustering metrics. G50: cluster size at which 50% of genes are in an orthogroup (OG) of that size or greater. O50: fewest number of orthogroups required to reach G50; G50 (assigned genes) = 16; G50 (all genes) = 14; O50 (assigned genes) = 3063; O50 (all genes) = 4112. **B.** Number of orthogroups shared by each species-pair (i.e. the number of orthogroups which contain at least one gene from each of the species-pairs). **C.** Number of species-specific orthogroups.

**Additional File 2.** Synteny blocks between pairwise comparisons of multiple parasitoid genomes. Synteny blocks were computed using SynChro (Drillon et al., 2014), a tool based on a simple algorithm that computes Reciprocal Best-Hits (RBH) to reconstruct the backbones of the synteny blocks.

**Additional File 3.** List of scaffolds in *Hyposoter didymator* and *Campoletis sonorensis* genomes containing at least on ichnovirus sequence. Are indicated the scaffold name and length, the name of the proviral segment or of the Ichnovirus structural protein encoding region (IVSPER) found in the scaffold, its length and position in the scaffold, the name of the direct repeats flanking the segment or within the segment, and the name of the genes predicted in each viral locus. DRJ, direct repeat junction; R, right; L, left; int, internal.

**Additional File 4.** List of ichnoviral genes identified in *Hyposoter didymator* and *Campoletis sonorensis* genome scaffolds containing at least on ichnovirus sequence. Are indicated the scaffold name, the name of the proviral segment or of the Ichnovirus structural protein encoding region (IVSPER) found in the scaffold, its length and position in the scaffold, the name of the gene, its position in the scaffold, if it contains or not an intron, the size of the predicted protein, then the NCBI blast P search results (NCBI accession number and ID of the best match, the blstP e-value and the percentage of identities).

**Additional File 5.** Transposable elements (TE) found in *Hyposter didymator* segments, IVSPERs and neighboring regions. The LOLA package (Sheffield and Bock, 2016) was used to assess if some particular TE were enriched close to viral circles or IVSPER. Genomics positions were enlarged to 10 kbp at each segments ends and sampled against 1,000 other similar regions from the genome, then used it a random reference. LOLA identifies overlaps and calculates enrichment for each TE. For each pairwise comparison, a series of columns describe the results of the statistical test (pvalueLog: -log10(pvalue) from the fisher’s exact result; oddsRatio: result from the fisher’s exact test; q-value transformation to provide false discovery rate (FDR) scores automatically). Some TE are enriched around viral locations, but after FDR correction, nothing was significant.

**Additional File 6.** List of direct repeat junctions (DRJ) found at the ends or within proviral segments genes identified in *Hyposoter didymator* and *Campoletis sonorensis* genome scaffolds. Are indicated the scaffold name, the name of the proviral segment, its length and position in the scaffold, the name of the DRJ, its length and position in the scaffold and the DRJ sequence. Nucleotide identities are indicated for each pair of DRJ.

**Additional File 7.** DRJs analysis. **A.** DNA motifs found in the direct repeated sequences flanking the IV segments inserted in wasp genomes. Analysis was performed using the DNAMINDA2 webserver (http://bmbl.sdstate.edu/DMINDA2/annotate.php); the input dataset was composed of 99 DRJ sequences (right junctions of HdIV and CsIV segments). A total of 89 motifs were obtained; only those whose occurrence exceed 70% of the DRJs are reported. **B.** Occurrence rate of motifs predicted with DMINDA 2.0 webserver in DRJs and whole genome sequences. Each of the two motifs was search among the 6 bp kmers present in the whole genome (201,969,604) and in the DRJs (33,930). The significance was evaluated using a Chi2 (taking into account the ratio of these motifs / all the other motifs in the DRJS and in the genome). **C.** Alignments of Sanger-sequenced DRJ regions from integrated and circular forms of seven *H. didymator* IV segments containing a putative excision site.

**Additional File 8.** List of the *Hyposoter didymator* genes present in the syntenic blocks represented in figures 6 and 7. Are indicated the *H. didymator* gene ID, its position in the scaffold, the number of the orthogroup to which it belongs and the result of the best match obtained following Blast similarity search. For each *H. didymator* gene, the corresponding *Campoletis sonorensis* gene ID, orthogroup number and position in *C. sonorensis* scaffold are indicated.

**Additional File 9.** Characteristics of the libraries used for genome assembly. **A.** *Campoletis sonorensis*. **B.** *Hyposoter didymator*.

## References

1. Brockhurst MA, Chapman T, King KC, Mank JE, Paterson S, Hurst GD. Running with the Red Queen: the role of biotic conflicts in evolution. Proc Biol Sci. 2014;281(1797).

2. Nuismer SL, Otto SP. Host-parasite interactions and the evolution of gene expression. PLoS Biol. 2005;3(7):e203.

3. Greenwood JM, Ezquerra AL, Behrens S, Branca A, Mallet L. Current analysis of host-parasite interactions with a focus on next generation sequencing data. Zoology (Jena). 2016;119(4):298–306.

4. Forbes AA, Bagley RK, Beer MA, Hippee AC, Widmayer HA. Quantifying the unquantifiable: why Hymenoptera, not Coleoptera, is the most speciose animal order. Bmc Ecol. 2018;18.

5. Whitfield JB. Molecular and morphological data suggest a single origin of the polydnaviruses among braconid wasps. Naturwissenschaften. 1997;84(11):502–7.

6. Whitfield JB. Estimating the age of the polydnavirus/braconid wasp symbiosis. P Natl Acad Sci USA. 2002;99(11):7508–13.

7. Strand MR, Burke GR. Polydnaviruses: Nature’s Genetic Engineers. Annu Rev Virol. 2014;1:333–54.

8. Beckage NE, Gelman DB. Wasp parasitoid disruption of host development: implications for new biologically based strategies for insect control. Annu Rev Entomol. 2004;49:299–330.

9. Darboux I, Cusson M, Volkoff AN. The dual life of ichnoviruses. Curr Opin Insect Sci. 2019;32:47–53.

10. Turnbull M, Webb B. Perspectives on polydnavirus origins and evolution. Adv Virus Res. 2002;58:203–54.

11. Webb B. Polydnavirus Biology, Genome Structure, and Evolution. In: Miller LK, Ball LA, editors. The Insect Viruses New York, USA: Springer; 1998. p. 105–39.

12. Webb B, Strand MR. The biology and genomics of polydnaviruses. In: Gilbert LI, Iatrou K, Gill SS, editors. Comprehensive Molecular Insect Science. San Diego, USA: Elsevier Press; 2005. p. 260–323.

13. Belle E, Beckage NE, Rousselet J, Poirie M, Lemeunier F, Drezen JM. Visualization of polydnavirus sequences in a parasitoid wasp chromosome. J Virol. 2002;76(11):5793–6.

14. Fleming JG, Summers MD. Polydnavirus DNA is integrated in the DNA of its parasitoid wasp host. Proc Natl Acad Sci U S A. 1991;88(21):9770–4.

15. Strand MR, Burke GR. Polydnaviruses: From discovery to current insights. Virology. 2015;479:393–402.

16. Bezier A, Louis F, Jancek S, Periquet G, Theze J, Gyapay G, et al. Functional endogenous viral elements in the genome of the parasitoid wasp Cotesia congregata: insights into the evolutionary dynamics of bracoviruses. Philos Trans R Soc Lond B Biol Sci. 2013;368(1626):20130047.

17. Burke GR, Walden KK, Whitfield JB, Robertson HM, Strand MR. Widespread genome reorganization of an obligate virus mutualist. PLoS Genet. 2014;10(9):e1004660.

18. Desjardins CA, Gundersen-Rindal DE, Hostetler JB, Tallon LJ, Fadrosh DW, Fuester RW, et al. Comparative genomics of mutualistic viruses of Glyptapanteles parasitic wasps. Genome Biol. 2008;9(12):R183.

19. Annaheim M, Lanzrein B. Genome organization of the Chelonus inanitus polydnavirus: excision sites, spacers and abundance of proviral and excised segments. J Gen Virol. 2007;88(Pt 2):450–7.

20. Savary S, Beckage N, Tan F, Periquet G, Drezen JM. Excision of the polydnavirus chromosomal integrated EP1 sequence of the parasitoid wasp Cotesia congregata (Braconidae, Microgastinae) at potential recombinase binding sites. J Gen Virol. 1997;78 (Pt 12):3125–34.

21. Burke GR, Simmonds TJ, Thomas SA, Strand MR. Microplitis demolitor Bracovirus Proviral Loci and Clustered Replication Genes Exhibit Distinct DNA Amplification Patterns during Replication. J Virol. 2015;89(18):9511–23.

22. Cui L, Webb BA. Homologous sequences in the Campoletis sonorensis polydnavirus genome are implicated in replication and nesting of the W segment family. J Virol. 1997;71(11):8504–13.

23. Rattanadechakul W, Webb BA. Characterization of Campoletis sonorensis ichnovirus unique segment B and excision locus structure. J Insect Physiol. 2003;49(5):523–32.

24. Bezier A, Annaheim M, Herbiniere J, Wetterwald C, Gyapay G, Bernard-Samain S, et al. Polydnaviruses of braconid wasps derive from an ancestral nudivirus. Science. 2009;323(5916):926–30.

25. Burke GR, Thomas SA, Eum JH, Strand MR. Mutualistic polydnaviruses share essential replication gene functions with pathogenic ancestors. PLoS Pathog. 2013;9(5):e1003348.

26. Volkoff AN, Jouan V, Urbach S, Samain S, Bergoin M, Wincker P, et al. Analysis of virion structural components reveals vestiges of the ancestral ichnovirus genome. PLoS Pathog. 2010;6(5):e1000923.

27. Burke GR, Walden KKO, Whitfield JB, Robertson HM, Strand MR. Whole Genome Sequence of the Parasitoid Wasp Microplitis demolitor That Harbors an Endogenous Virus Mutualist. G3 (Bethesda). 2018;8(9):2875–80.

28. Beliveau C, Cohen A, Stewart D, Periquet G, Djoumad A, Kuhn L, et al. Genomic and Proteomic Analyses Indicate that Banchine and Campoplegine Polydnaviruses Have Similar, if Not Identical, Viral Ancestors. J Virol. 2015;89(17):8909–21.

29. Webb BA, Strand MR, Dickey SE, Beck MH, Hilgarth RS, Barney WE, et al. Polydnavirus genomes reflect their dual roles as mutualists and pathogens. Virology. 2006;347(1):160–74.

30. Doremus T, Cousserans F, Gyapay G, Jouan V, Milano P, Wajnberg E, et al. Extensive Transcription Analysis of the Hyposoter didymator Ichnovirus Genome in Permissive and Non-Permissive Lepidopteran Host Species. Plos One. 2014;9(8).

31. Weisenfeld NI, Kumar V, Shah P, Church DM, Jaffe DB. Direct determination of diploid genome sequences. Genome Res. 2017;27(5):757–67.

32. Kajitani R, Toshimoto K, Noguchi H, Toyoda A, Ogura Y, Okuno M, et al. Efficient de novo assembly of highly heterozygous genomes from whole-genome shotgun short reads. Genome Res. 2014;24(8):1384–95.

33. Yang JY, Chen X, McDermaid A, Ma Q. DMINDA 2.0: integrated and systematic views of regulatory DNA motif identification and analyses. Bioinformatics. 2017;33(16):2586–8.

34. Gundersen-Rindal D, Dupuy C, Huguet E, Drezen JM. Parasitoid polydnaviruses: evolution, pathology and applications. Biocontrol Science and Technology. 2013;23:1–61.

35. Bennett AMR, Cardinal S, Gauld ID, Wahl DB. Phylogeny of the subfamilies of Ichneumonidae (Hymenoptera). J Hymenopt Res. 2019;71:1–156.

36. Quicke DLJ, Laurenne NM, Fitton MG, Broad GR. A thousand and one wasps: a 28S rDNA and morphological phylogeny of the Ichneumonidae (Insecta: Hymenoptera) with an investigation into alignment parameter space and elision. J Nat Hist. 2009;43(23-24):1305–421.

37. Louis F, Bezier A, Periquet G, Ferras C, Drezen JM, Dupuy C. The bracovirus genome of the parasitoid wasp Cotesia congregata is amplified within 13 replication units, including sequences not packaged in the particles. J Virol. 2013;87(17):9649–60.

38. Lorenzi A, Ravallec M, Eychenne M, Jouan V, Robin S, Darboux I, et al. RNA interference identifies domesticated viral genes involved in assembly and trafficking of virus-derived particles in ichneumonid wasps. Plos Pathogens. In Press.

39. Gruber A, Stettler P, Heiniger P, Schumperli D, Lanzrein B. Polydnavirus DNA of the braconid wasp Chelonus inanitus is integrated in the wasp’s genome and excised only in later pupal and adult stages of the female. J Gen Virol. 1996;77 (Pt 11):2873–9.

40. Opperman R, Emmanuel E, Levy AA. The effect of sequence divergence on recombination between direct repeats in Arabidopsis. Genetics. 2004;168(4):2207–15.

41. Desjardins CA, Gundersen-Rindal DE, Hostetler JB, Tallon LJ, Fuester RW, Schatz MC, et al. Structure and evolution of a proviral locus of Glyptapanteles indiensis bracovirus. BMC Microbiol. 2007;7:61.

42. Quicke DLJ. The Braconid and Ichneumonid Parasitoid Wasps: Biology, Systematics, Evolution and Ecology. Hoboken, USA: John Wiley & Sons; 2015.

43. Shaw MR, Horstmann K. An analysis of host range in the Diadegma nanus group of parasitoids in Western Europe, with a key to species. J Hymenopt Res. 1997;6:273–96.

44. Frayssinet M, Audiot P, Cusumano A, Pichon A, Malm LE, Jouan V, et al. Western European Populations of the Ichneumonid Wasp Hyposoter didymator Belong to a Single Taxon. Front Ecol Evol. 2019;7.

45. Pacheco HM, Vanlaerhoven SL, Garcia MAM, Hunt DW. Food web associations and effect of trophic resources and environmental factors on parasitoids expanding their host range into non-native hosts. Entomol Exp Appl. 2018;166(4):277–88.

46. Krell PJ, Summers MD, Vinson SB. Virus with a Multipartite Superhelical DNA Genome from the Ichneumonid Parasitoid Campoletis sonorensis. J Virol. 1982;43(3):859–70.

47. Walker BJ, Abeel T, Shea T, Priest M, Abouelliel A, Sakthikumar S, et al. Pilon: an integrated tool for comprehensive microbial variant detection and genome assembly improvement. PLoS One. 2014;9(11):e112963.

48. Smit AFA, Hubley R, Green P. RepeatMasker Open-4.0 2013-2015 [Available from: http://www.repeatmasker.org.

49. Kent WJ. BLAT--the BLAST-like alignment tool. Genome Res. 2002;12(4):656–64.

50. Simao FA, Waterhouse RM, Ioannidis P, Kriventseva EV, Zdobnov EM. BUSCO: assessing genome assembly and annotation completeness with single-copy orthologs. Bioinformatics. 2015;31(19):3210–2.

51. Stanke M, Tzvetkova A, Morgenstern B. AUGUSTUS at EGASP: using EST, protein and genomic alignments for improved gene prediction in the human genome. Genome Biol. 2006;7 Suppl 1:S111–8.

52. Luo R, Liu B, Xie Y, Li Z, Huang W, Yuan J, et al. SOAPdenovo2: an empirically improved memory-efficient short-read de novo assembler. Gigascience. 2012;1(1):18.

53. Bradnam KR, Fass JN, Alexandrov A, Baranay P, Bechner M, Birol I, et al. Assemblathon 2: evaluating de novo methods of genome assembly in three vertebrate species. Gigascience. 2013;2(1):10.

54. Doremus T, Urbach S, Jouan V, Cousserans F, Ravallec M, Demettre E, et al. Venom gland extract is not required for successful parasitism in the polydnavirus-associated endoparasitoid Hyposoter didymator (Hym. Ichneumonidae) despite the presence of numerous novel and conserved venom proteins. Insect Biochem Mol Biol. 2013;43(3):292–307.

55. Wu TD, Watanabe CK. GMAP: a genomic mapping and alignment program for mRNA and EST sequences. Bioinformatics. 2005;21(9):1859–75.

56. Dobin A, Davis CA, Schlesinger F, Drenkow J, Zaleski C, Jha S, et al. STAR: ultrafast universal RNA-seq aligner. Bioinformatics. 2013;29(1):15–21.

57. Kim D, Pertea G, Trapnell C, Pimentel H, Kelley R, Salzberg SL. TopHat2: accurate alignment of transcriptomes in the presence of insertions, deletions and gene fusions. Genome Biol. 2013;14(4):R36.

58. Hoff KJ, Lange S, Lomsadze A, Borodovsky M, Stanke M. BRAKER1: Unsupervised RNA-Seq-Based Genome Annotation with GeneMark-ET and AUGUSTUS. Bioinformatics. 2016;32(5):767–9.

59. Geib SM, Liang GH, Murphy TD, Sim SB. Whole Genome Sequencing of the Braconid Parasitoid Wasp Fopius arisanus, an Important Biocontrol Agent of Pest Tepritid Fruit Flies. G3 (Bethesda). 2017;7(8):2407–11.

60. Tvedte E, Walden KK, Mcelroy K, Werren JH, Forbes AA, Hood GR, et al. Genome of the parasitoid wasp Diachasma alloeum, an emerging model for ecological speciation and transitions to asexual reproduction. BioRxiv; 2019.

61. Werren JH, Richards S, Desjardins CA, Niehuis O, Gadau J, Colbourne JK, et al. Functional and evolutionary insights from the genomes of three parasitoid Nasonia species. Science. 2010;327(5963):343–8.

62. Pichon A, Bezier A, Urbach S, Aury JM, Jouan V, Ravallec M, et al. Recurrent DNA virus domestication leading to different parasite virulence strategies. Sci Adv. 2015;1(10):e1501150.

63. Dunn NA, Unni DR, Diesh C, Munoz-Torres M, Harris NL, Yao E, et al. Apollo: Democratizing genome annotation. PLoS Comput Biol. 2019;15(2):e1006790.

64. Flutre T, Duprat E, Feuillet C, Quesneville H. Considering transposable element diversification in de novo annotation approaches. PLoS One. 2011;6(1):e16526.

65. Quesneville H, Bergman CM, Andrieu O, Autard D, Nouaud D, Ashburner M, et al. Combined evidence annotation of transposable elements in genome sequences. PLoS Comput Biol. 2005;1(2):166–75.

66. Sheffield NC, Bock C. LOLA: enrichment analysis for genomic region sets and regulatory elements in R and Bioconductor. Bioinformatics. 2016;32(4):587–9.

67. Quinlan AR. BEDTools: The Swiss-Army Tool for Genome Feature Analysis. Curr Protoc Bioinformatics. 2014;47:11 21–34.

68. Emms DM, Kelly S. OrthoFinder: solving fundamental biases in whole genome comparisons dramatically improves orthogroup inference accuracy. Genome Biol. 2015;16:157.

69. Drillon G, Carbone A, Fischer G. SynChro: a fast and easy tool to reconstruct and visualize synteny blocks along eukaryotic chromosomes. PLoS One. 2014;9(3):e92621.

70. Rocher J, Ravallec M, Barry P, Volkoff AN, Ray D, Devauchelle G, et al. Establishment of cell lines from the wasp Hyposoter didymator (Hym., Ichneumonidae) containing the symbiotic polydnavirus H didymator ichnovirus. Journal of General Virology. 2004;85:863–8.

71. Volkoff AN, Cerutti P, Rocher J, Ohresser MCP, Devauchelle G, Duonor-Cerutti M. Related RNAs in lepidopteran cells after in vitro infection with Hyposoter didymator virus define a new polydnavirus gene family. Virology. 1999;263(2):349–63.

72. Baudet C, Lemaitre C, Dias Z, Gautier C, Tannier E, Sagot MF. Cassis: detection of genomic rearrangement breakpoints. Bioinformatics. 2010;26(15):1897–8.

73. Lemaitre C, Tannier E, Gautier C, Sagot MF. Precise detection of rearrangement breakpoints in mammalian chromosomes. BMC Bioinformatics. 2008;9:286.

74. Robin S, Ravallec M, Frayssinet M, Whitfield J, Jouan V, Legeai F, et al. Evidence for an ichnovirus machinery in parasitoids of coleopteran larvae. Virus Res. 2019;263:189–206.

