## Additional file 1 for "Conserved and specific genomic features of endogenous polydnaviruses revealed by whole genome sequencing of two ichneumonid wasps"

**Additional File 1. Orthogroups analyses**

**A**: Orthofinder clustering metrics. G50: cluster size at which 50% of genes are in an orthogroup (OG) of that size or greater. O50: fewest number of orthogroups required to reach G50 : G50 (assigned genes) = 16, G50 (all genes) = 14, O50 (assigned genes) = 3063, O50 (all genes) = 4112

| | | *Hymenoptera: Ichneumonidae* | | | *Hymenoptera: Braconidae* | | | *Hymenoptera: Pteromalidae* | *Hymenoptera: Apidae* | *Diptera: Drosophilidae* |
|---|---|---|---|---|---|---|---|---|---|---|
| | All | *Hyposoter didymator* | *Campoletis sonorensis* | *Venturia canescens* | *Microplitis demolitor* | *Fopius arisanus* | *Diachasma alloeum* | *Nasonia vitripennis* | *Apis mellifera* | *Drosophila melanogaster* |
| Number of genes | 199,523 | 18,154 | 21,987 | 23,401 | 18,586 | 18,906 | 19,692 | 24,846 | 23,471 | 30,480 |
| Number of genes in OGs | 168,268 | 15,808 | 15,691 | 17,511 | 16,843 | 18,392 | 18,270 | 21,417 | 22,483 | 21,853 |
| % of genes in OGs | 84.3 | 87.1 | 71.4 | 74.8 | 90.6 | 97.3 | 92.8 | 86.2 | 95.8 | 71.7 |
| Number of unassigned genes | 31,255 | 2,346 | 6,296 | 5,890 | 1,743 | 514 | 1,422 | 3,429 | 988 | 8,627 |
| Total number of OGs | 12,825 | 10,880 | 10,293 | 11,041 | 9,121 | 9,558 | 9,616 | 9,094 | 8,976 | 7,932 |
| Number species-specific OGs | 501 | 11 | 32 | 26 | 29 | 8 | 17 | 77 | 36 | 265 |
| Number genes in species-specific OGs | 2,773 | 69 | 288 | 216 | 163 | 36 | 76 | 406 | 141 | 1,378 |
| % genes in species-specific OGs | 1.4 | 0.4 | 1.3 | 0.9 | 0.9 | 0.2 | 0.4 | 1.6 | 0.6 | 4.5 |
| Number of single-copy OGs | 454 | | | | | | | | | |

**B.** Number of orthogroups shared by each species-pair (i.e. the number of orthogroups which contain at least one gene from each of the species-pairs)

| | *Hyposoter didymator* | *Campoletis sonorensis* | *Venturia canescens* | *Microplitis demolitor* | *Fopius arisanus* | *Diachasma alloeum* | *Nasonia vitripennis* | *Apis mellifera* | *Drosophila melanogaster* |
|---|---|---|---|---|---|---|---|---|---|
| *Hyposoter didymator* | 10,869 | 9,696 | 10,362 | 8,664 | 8,830 | 8,838 | 8,508 | 8,586 | 7,148 |
| *Campoletis sonorensis* | | 10,261 | 9,789 | 8,086 | 8,258 | 8,257 | 7,955 | 8,031 | 6,710 |
| *Venturia canescens* | | | 11,015 | 8,665 | 8,800 | 8,806 | 8,492 | 8,545 | 7,123 |
| *Microplitis demolitor* | | | | 9,092 | 8,592 | 8,597 | 8,159 | 8,207 | 6,991 |
| *Fopius arisanus* | | | | | 9,550 | 9,256 | 8,284 | 8,371 | 7,062 |
| *Diachasma alloeum* | | | | | | 9,599 | 8,312 | 8,357 | 7,068 |
| *Nasonia vitripennis* | | | | | | | 9,017 | 8,134 | 6,912 |
| *Apis mellifera* | | | | | | | | 8,940 | 7,022 |
| *Drosophila melanogaster* | | | | | | | | | 7,667 |

**C.** Number of species-specific orthogroups.

| | **Nb species-specific orthogroups** |
|---|---|
| ***Hyposoter didymator* (Hd)** | 11 |
| ***Campoletis sonorensis* (Cs)** | 32 |
| ***Venturia canescens* (Vc)** | 26 |
| **IV carrying Ichneumonids (Hd, Cs)** | 313 |
| **Ichneumonids (Hd, Cs, Vc)** | 1,728 |
| **Ichneumonids & Braconids (Hd, Cs, Vc, Md, Fa, Da)** | 2,610 |
| **Parasitic wasps** | 3,240 |
| **Hymenoptera** | 5,158 |
| **Hymenoptera + diptera** | 12,825 |
