## Additional file 2 for "Conserved and specific genomic features of endogenous polydnaviruses revealed by whole genome sequencing of two ichneumonid wasps"

**Additional File 2. Synteny blocks between pairwise comparisons of multiple parasitoid genomes**

Synteny blocks were computed using SynChro (Drillon et al., 2014; doi: 10.1371/journal.pone.0092621), a tool based on a simple algorithm that computes Reciprocal Best-Hits (RBH) to reconstruct the backbones of the synteny blocks.

| species 1 | species 2 | # genes in Species 1 | # genes in Species 2 | RBH | mean % similarity of RBH genes | # synteny blocks (SB) | mean number of genes in SB | Species 1 nucleotides In SB | Species 2 nucleotides in SB | # Species 1 genes in SB | # Species 2 genes in SB |
|---|---|---|---|---|---|---|---|---|---|---|---|
| ***Campoletis sonorensis*** | ***Hyposoter didymator*** | **21,987** | **18,154** | **9,201** | **82.2258** | **1,112** | **11.2334** | **130,986,569** | **136,689,043** | **11,816** | **12,828** |
| *Campoletis sonorensis* | *Diachasma alloeum* | 21,987 | 19,692 | 7,054 | 72.4531 | 1,581 | 6.1151 | 79,457,000 | 121,585,809 | 8,101 | 11,023 |
| *Campoletis sonorensis* | *Fopius arisanus* | 21,987 | 18,906 | 6,963 | 71.7424 | 1,583 | 6.2612 | 78,144,432 | 64,120,970 | 7,957 | 11,583 |
| *Campoletis sonorensis* | *Microplitis demolitor* | 21,987 | 18,586 | 6,889 | 71.3963 | 1,503 | 6.4112 | 79,951,790 | 87,227,954 | 8,066 | 10,937 |
| *Campoletis sonorensis* | *Venturia canescens* | 21,987 | 23,401 | 9,602 | 81.9265 | 1,501 | 7.3997 | 84,998,388 | 87,756,926 | 10,31 | 11,589 |
| *Diachasma alloeum* | *Fopius arisanus* | 19,692 | 18,906 | 10,85 | 84.4038 | 1,244 | 13.2826 | 188,094,018 | 98,011,380 | 15,707 | 17,083 |
| *Diachasma alloeum* | *Hyposoter didymator* | 19,692 | 18,154 | 8,784 | 71.9646 | 1,949 | 5.8489 | 147,155,684 | 85,519,660 | 12,100 | 10,426 |
| *Diachasma alloeum* | *Microplitis demolitor* | 19,692 | 18,586 | 9,227 | 75.5011 | 1,723 | 8.1993 | 156,017,415 | 115,880,628 | 13,263 | 13,987 |
| *Diachasma alloeum* | *Venturia canescens* | 19,692 | 23,401 | 8,435 | 71.8280 | 1,960 | 4.8980 | 101,537,848 | 57,440,014 | 10,261 | 8,663 |
| *Fopius arisanus* | *Hyposoter didymator* | 18,906 | 18,154 | 8,622 | 71.5998 | 2,004 | 5.9214 | 78,694,026 | 88,834,572 | 12,881 | 10,551 |
| *Fopius arisanus* | *Microplitis demolitor* | 18,906 | 18,586 | 9,151 | 74.8577 | 1,798 | 7.8715 | 82,643,504 | 117,622,728 | 14,057 | 13,906 |
| *Fopius arisanus* | *Venturia canescens* | 18,906 | 23,401 | 8,203 | 71.5408 | 1,957 | 5.0424 | 53,941,273 | 57,217,953 | 10,812 | 8,631 |
| *Hyposoter didymator* | *Microplitis demolitor* | 18,154 | 18,586 | 8,513 | 72.4243 | 1,844 | 6.2796 | 82,632,725 | 106,502,067 | 10,063 | 12,767 |
| *Hyposoter didymator* | *Venturia canescens* | 18,154 | 23,401 | 11,21 | 82.4090 | 1,695 | 7.2375 | 96,760,707 | 94,022,094 | 11,769 | 12,407 |
| *Microplitis demolitor* | *Venturia canescens* | 18,586 | 23,401 | 8,163 | 70.9083 | 1,848 | 5.1366 | 72,847,449 | 56,076,459 | 10,217 | 8,494 |
