## Additional file 3 for "Conserved and specific genomic features of endogenous polydnaviruses revealed by whole genome sequencing of two ichneumonid wasps"

| Scaffold | Scaffold length (nt) | Name of proviral segment /IVSPER | Proviral/IVSPER sequence length (nt) | Position in scaffold | Direct repeats | Genes present in proviral sequence |
|---|---|---|---|---|---|---|
| ***Hyposoter didymator*** | | | | | | |
| scaffold49647 | 1586 | similar to Hd2 | 1537 | 1-1537 (partial) | not applicable | Gly-Pro_P40-like, partial |
| scaffold28498 | 4159 | Hd45.1 | NA | 1-4214 (partial) | no repeat found at extremities | U1_Hd45.1, U2_Hd45.1 |
| scaffold82201 | 6109 | Hd51 | 4632 | 1077-5708 | Hd51_DRJ1R/L, Hd51_DRJ2R/L | Rep1_Hd51 |
| scaffold29771 | 25145 | IVSP_U37 | 1839 | 16848-18686 | not applicable | single CDS: U37 |
| scaffold29771 | 25145 | Hd46 | 4109 | 19344-23452 | Hd46_DRJ1R/L, Hd46_DRJ2R/L | Vank_Hd46 |
| scaffold1868 | 202142 | Hd43 | 4159 | 193639-197797 | Hd43_DRJ1R/L, Hd43_DRJ2R/L | U1_Hd43, Vank1_Hd43 |
| scaffold128213 | 363895 | Hd23.1 | 4457 | 208205-212661 | Hd23.1_DRJ1R/L | Rep1_Hd23.1 |
| scaffold128213 | 363895 | Hd23.2 | 3362 | 248624-251985 | Hd23.2_DRJ1R/L | Rep1_Hd23.2 |
| scaffold91 | 761975 | IVSPER-1 | 14020 | 453857-467876 | not applicable | U1, IVSP1-1, U2, U3, U4, p53-2, U5, IVSP2-1, N-1, U25 |
| scaffold91 | 761975 | Hd15 | 4987 | 469105-474091 | Hd15_DRJ1R/L | N1_Hd15 |
| scaffold91 | 761975 | Hd33 | 3835 | 485063-488897 | Hd33_DRJ1R/L | U1_Hd33, U2_Hd33 |
| scaffold91 | 761975 | Hd24 | 4697 | 535698-540394 | Hd24_DRJ1R/L | Vank1_Hd24, Vinx1_Hd24 |
| scaffold91 | 761975 | IVSPER-2 | 26611 | 541304-567914 | not applicable | N-4, U26, IVSP2-2, IVSP4-1, p12-2, U27, U14, p12-3, U13, U12, U11, U10, U9, U8, IVSP3-1, IVSP1-2, U7, U6, N-3 |
| scaffold91 | 761975 | Hd29 | 4356 | 572514-576869 | Hd29_DRJ1R/L | N1_Hd29 |
| scaffold128215 | 912714 | Hd9 | 17892 | 526917-544808 | Hd9_DRJ1R/int/L, Hd9_DRJ2R/L | U1.1_Hd9, U2_Hd9, U6.1_Hd9, U3.1_Hd9, U4.1_Hd9, U5.1_Hd9, U1.2_Hd9, U6.2_Hd9, U3.2_Hd9, U4.2_Hd9, U5.2_Hd9, U1.3_Hd9 |
| scaffold65 | 1016704 | Hd37 | 3708 | 342293-346000 | Hd37_DRJ1R/L | Rep1_Hd37, U1_Hd37, U2_Hd37 |
| scaffold65 | 1016704 | Hd3 | 10014 | 437082-447095 | Hd3_DRJ1R/int/L, Hd3_DRJ2R/int/L | Cys1_Hd3, Cys2_Hd3, Cys3_Hd3, Cys4_Hd3, Cys5_Hd3 |
| scaffold90 | 1092007 | Hd1 | NA | 1-14771 (partial) | Hd1_DRJ1R/int, Hd1_DRJ2R/int-1/int-2 | U1_Hd1, U2_Hd1, U3_Hd1, U4_Hd1, U5_Hd1, U6_Hd1 |
| scaffold264 | 1259582 | Hd28 | 4614 | 135485-140098 | Hd28_DRJ1R/L | Vinx1_Hd28, Vank1_Hd28 |
| scaffold128246 | 1305886 | Hd49 | 5265 | 677866-683130 | Hd49_DRJ1R/L | Rep1_Hd49, Rep2_Hd49, Rep3_Hd49 |
| scaffold429 | 1434187 | Hd48 | 9673 | 531833-541505 | Hd48_DRJ1R/L, Hd48_DRJ2R/L | Rep1_Hd48, Rep2_Hd48 |
| scaffold144 | 1844462 | Hd20 | 6864 | 1449468-1456331 | Hd20_DRJ1R/L, Hd20_DRJ2R/L | Cys1_Hd20, Cys2_Hd20 |
| scaffold116 | 1992701 | Hd35 | 3710 | 473341-477050 | Hd35_DRJ1R/L | Rep1_Hd35 |
| scaffold64 | 2416597 | Hd14 | 5196 | 36336-41531 | Hd14_DRJ1R/L | Vinx1_Hd14, U1_Hd14 |
| scaffold64 | 2416597 | Hd32 | 7916 | 88702-96617 | Hd32_DRJ1R/L, Hd32_DRJ2R/L | Vinx1_Hd32, Vinx2_Hd32, U1.1_Hd32, U1.2_Hd32 |
| scaffold64 | 2416597 | Hd42 | 3157 | 2320523-2323679 | Hd42_DRJ1R/L | Rep1_Hd42 |
| scaffold64 | 2416597 | Hd21 | 4368 | 2353107-2357474 | Hd21_DRJ1R/L | Rep1_Hd21, Vinx1_Hd21 |
| scaffold377 | 2592399 | Hd8 | 7356 | 2186417-2193772 | Hd8_DRJ1R/L, Hd8_DRJ2R/L | U1_Hd8, U2_Hd8, U3_Hd8, U4_Hd8, U5_Hd8 |
| scaffold377 | 2592399 | Hd4 | 10326 | 2459681-2470006 | Hd4_DRJ1R/int/L | Rep1_Hd4, Rep2_Hd4, Rep3_Hd4, Rep4_Hd4, Rep5_Hd4, Rep6_Hd4 |
| scaffold161 | 2602850 | Hd25 | 4174 | 2449400-2453573 | Hd25_DRJ1R/L | PRRP1_Hd25, PRRP2_Hd25, PRRP3_Hd25 |
| scaffold127549 | 2774194 | Hd31-34 | 4119 | 144445-148563 | Hd31-34_DRJ1R/L, Hd31-34_DRJ2R/L | U1_Hd31-34, U2_Hd31-34 |
| scaffold59 | 2934223 | Hd12 | 5902 | 674917-680818 | Hd12_DRJ1R/L | U1_Hd12, Rep1_Hd12, U2_Hd12, Rep2_Hd12 |
| scaffold59 | 2934223 | Hd16 | 7704 | 690205-697908 | Hd16_DRJ1R/L, Hd16_DRJ2R/L | Rep1_Hd16, Rep2_Hd16, Rep3_Hd16, U1_Hd16, U2_Hd16 |
| scaffold59 | 2934223 | Hd11 | 9190 | 2183512-2192701 | Hd11_DRJ1R/L, Hd11_DRJ2R/L | U1_Hd11, Rep1_Hd11, Vank1_Hd11, Vank1p_Hd11, Vank2_Hd11, Vank3_Hd11, Vank4_Hd11, Vank5_Hd11 |
| scaffold59 | 2934223 | Hd10 | 6507 | 2500377-2506883 | Hd10_DRJ1R/int/L, Hd10_DRJ2R/L | Rep1_Hd10, Rep2_Hd10, Rep3_Hd10 |
| scaffold198 | 3327987 | Hd19 | 4440 | 1265454-1269893 | Hd19_DRJ1R/L | U1_Hd19 |
| scaffold119 | 3623981 | Hd22 | 4178 | 698427-702604 | Hd22_DRJ1R/L | Rep1_Hd22, U1_Hd22 |
| scaffold127348 | 3747147 | Hd40 | 3494 | 926528-930021 | Hd40_DRJ1R/L | U1_Hd40, U2_Hd40 |
| scaffold357 | 4004213 | Hd50 | 5787 | 2218373-2224159 | Hd50_DRJ1R/L | Vinx1_Hd50, Vinx2_Hd50 |
| scaffold128243 | 5513913 | Hd44.2 | 4831 | 4197203-4202033 | Hd44.2_DRJ1R/L | Rep1_Hd44.2, U1_Hd44.2 |
| scaffold128243 | 5513913 | Hd44.1 | 3009 | 4203985-4206993 | Hd44.1_DRJ1R/L | U1_Hd44.1 |
| scaffold351 | 5773413 | Hd18 | 4696 | 2681961-2686656 | Hd18_DRJ1R/L | N1_Hd18 |
| scaffold351 | 5773413 | Hd17 | 7730 | 2329273-2337002 | Hd17_DRJ1R/L, Hd17_DRJ2R/L | Rep1_Hd17, Rep2_Hd17, Rep3_Hd17, Rep4_Hd17, Rep5_Hd17 |
| scaffold22 | 5910489 | Hd39 | 4122 | 1002620-1006741 | Hd39_DRJ1R/L, Hd39_DRJ2R/L | Rep1_Hd39 |

| | | | | | | |
|---|---|---|---|---|---|---|
| scaffold128241 | 6033897 | Hd13 | 5757 | 402487-408243 | Hd13_DRJ1R/L | Cys1_Hd13, Cys2_Hd13 |
| scaffold67 | 6283432 | Hd30 | 4164 | 5867182-5871345 | Hd30_DRJ1R/L, Hd30_DRJ2R/L | Vinx1_Hd30, U1_Hd30, U2_Hd30, U3_Hd30 |
| scaffold184 | 7628668 | Hd45.2 | 2051 | 3564791-3566841 | no repeat found at extremities | U1_Hd45.2 |
| scaffold184 | 7628668 | Hd41 | 7953 | 3768924-3776876 | Hd41_DRJ1R/L, Hd41_DRJ2R/L | U1_Hd41, U2_Hd41, U3_Hd41 |
| scaffold175 | 12456768 | Hd36 | 3738 | 3796140-3799877 | Hd36_DRJ1R/L | U1_Hd36, Vinx1_Hd36 |
| scaffold175 | 12456768 | Hd38 | 3664 | 3800393-3804056 | Hd38_DRJ1R/L | U1_Hd38, Vinx1_Hd38, U2_Hd38 |
| scaffold175 | 12456768 | Hd26 | 5018 | 10942034-10947051 | Hd26_DRJ1R/L, Hd26_DRJ2R/L | PRRP1_Hd26, PRRP2_Hd26, U1_Hd26 |
| scaffold127548 | 15678958 | Hd6 | 10461 | 5808388-5818848 | Hd6_DRJ1R/int/L, Hd6_DRJ2R/L | U1.1_Hd6, P30_Hd6, Rep1_Hd6, U1.2_Hd6 |
| scaffold127548 | 15678958 | Hd2 | 13937 | 5940296-5954232 | Hd2_DRJ1R/int/L, Hd2_DRJ2R/L | GlyPro1_Hd2, U1_Hd2, U2_Hd2, GlyPro2_Hd2, SerThr1_Hd2 |
| scaffold127548 | 15678958 | Hd7 | 8066 | 6062921-6070986 | Hd7_DRJ1R/L | U1_Hd7 |
| scaffold127548 | 15678958 | IVSPER-4 | 15811 | 6832835-6848646 | not applicable | U29, U30, U31, U32, U33, U34 |
| scaffold127548 | 15678958 | IVSPER-3 | 25432 | 10761570-10787001 | not applicable | U15, IVSP3-2, U16, U17, U18, p12-1, U19, IVSP4-2, U20, U21, U22, U23, U28, p53-1, U24, N-2 |
| scaffold127548 | 15678958 | IVSPER-5 | 1629 | 10860001-10861630 | not applicable | U35, U36 |
| scaffold127548 | 15678958 | Hd47 | 4503 | 12134587-12139089 | Hd47_DRJ1R/L | Rep1_Hd47, Rep2_Hd47 |
| scaffold127548 | 15678958 | Hd5 | 13713 | 12941247-12954959 | Hd5_DRJ1R/int-1/int-2/L | Vinx1_Hd5, Vinx2_Hd5, U1_Hd5, Vinx3_Hd5, Vinx4_Hd5, Vinx5_Hd5, Vinx6_Hd5 |
| scaffold127548 | 15678958 | Hd27 | 4002 | 13338255-13342256 | Hd27_DRJ1R/L | U1_Hd27, K19_Hd27 |
| ***Campoletis sonorensis*** | | | | | | |
| scaffold_8749 | 2269 | rep gene | NA | not applicable | not applicable | rep gene 1 |
| scaffold_8748 | 2287 | rep gene | NA | not applicable | not applicable | rep gene 2 |
| scaffold_7280 | 2380 | CsV | NA | not applicable | not applicable | cys_CsV, partial |
| scaffold_8362 | 5297 | CsX5, partial | >5297 | 1-5297 | no repeat searched | rep1_CsX5, rep2_CsX5 |
| scaffold_4391 | 18751 | CsL | 10024 | 8079-18102 | CsL_DRJ1R/L | cys_CsL |
| scaffold_5218 | 32996 | CsP | 12113 | 15720-27832 | CsP_DRJ1R/L | vank4_CsP, vank3_CsP, vank2_CsP, vank1_CsP |
| scaffold_5934 | 46742 | CsX1 | 17335 | 19391-36725 | CsX1_DRJ1R/int1/int2/L | vank1_CsX1, vnx1_CsX1, rep1_CsX1, vank2_CsX1, vank3_CsX1, vank4_CsX1, vnx2_CsX1, rep2_CsX1, vank5_CsX1 |
| scaffold_128 | 152063 | CsI2 | 9042 | 110016-119057 | CsI2_DRJ1R/L | vank1_CsI2, rep_CsI2, vank2_CsI2, vank3_CsI2 |
| scaffold_35 | 217784 | CsX8 | 9999 | 164467-174465 | CsX8_DRJ1R/L | HP1_CsX8, cys_CsX8, HP2_CsX8 |
| scaffold_5890 | 247009 | CsZ | 15871 | 134147-150017 | CsZ_DRJ1R/L | rep1_CsZ, rep2_CsZ, rep3_CsZ, rep4_CsZ, rep5_CsZ, rep6_CsZ, rep7_CsZ |
| scaffold_110 | 266287 | CsX2 | 10806 | 248068-258873 | CsX2_DRJ1R/L, CsX2_DRJ2R/L | vank1_CsX2, vank2_CsX2, vank3_CsX2, vank4_CsX2, rep_CsX2 |
| scaffold_50 | 323180 | Cs_IVSPER-3 | 8610 | 218627-227236 | not applicable | IVSP4L-2, U4L, p53L-3, U5L, IVSP2L-2, CsN-3 |
| scaffold_50 | 323180 | CsQ | 12543 | 290527-303069 | CsQ_DRJ1R/int/L | rep1_CsQ, vinx1_CsQ, vinx2_CsQ, rep2_CsQ, rep3_CsQ, rep4_CsQ |
| scaffold_6122 | 412025 | Cs_IVSPER-1 | 31594 | 122689-154282 | not applicable | U15L, IVSP1L-1, U37L-1, U31L-1, U35L, Gf_U27L, U17L, p12L-1, U19L, IVSP4L-1, U22L, U23L, p53L-1, U24L, CsN-1 |
| scaffold_6122 | 412025 | IVSP_U36L | 471 | 234803-235273 | not applicable | U36L |
| scaffold_60 | 424299 | CsX3, partial | >7876 | 304794-312669 | no repeat found at extremities | rep1_CsX3, rep2_CsX3, rep3_CsX3, rep4_CsX3 |
| scaffold_6095 | 433103 | CsX7, partial | >6041 | 183042-189082 | no repeat found at extremities | rep1_CsX7, rep2_CsX7, rep3_CsX7 |
| scaffold_6070 | 495171 | CsO1 | 12746 | 168701-181446 | no repeat found at extremities | 4rep_CsO1, 3rep_CsO1 |
| scaffold_116 | 506917 | CsT | 23217 | 7789-31005 | CsT_DRJ1R/L, CsT_DRJ2R/L | no gene found |
| scaffold_49 | 562263 | CsB | 6626 | 22030-28655 | CsB_DRJ1R/L | rep_CsB |
| scaffold_14 | 725399 | CsG2 | 8338 | 192247-200584 | CsG2_DRJ1R/L | rep1_CsG2, rep2_CsG2, rep3_CsG2, rep4_CsG2 |
| scaffold_14 | 725399 | CsG | 8656 | 76017-84672 | CsG_DRJ1R/L | vnx_CsG |
| scaffold_28 | 729583 | CsC | 7350 | 25280-32629 | no repeat found at extremities | overlap Cs_IVSPER-2 |
| scaffold_28 | 729583 | CsW | 15807 | 614005-629811 | CsW_DRJ1R/int1/int2/L | cys1_CsW, cys2_CsW, rep1_CsW, cys3_CsW, rep2_CsW, rep3_CsW |
| scaffold_28 | 729583 | Cs_IVSPER-2 | 33269 | 7310-40578 | not applicable | p53L-2, U6L, U7L, IVSP1L-2, IVSP3L, U31L-2, U8L, U9L, U16L, U10L, U11L, U12L, U13L, p12L-2, U3L, IVSP2L-1, U26L, CsN-2, U25L |
| scaffold_22 | 866858 | CsI | 8779 | 695663-704441 | CsI_DRJ1R/L | rep1_CsI, rep2_CsI, rep3_CsI |
| scaffold_57 | 1098263 | Cs_IVSPER-4 | 9937 | 383305-393241 | not applicable | U30L, U34L, IVSP4L-3 |
| scaffold_131 | 1180360 | CsF | 8155 | 808380-816534 | CsF_DRJ1R/L | cys_CsF |
| scaffold_149 | 1226226 | CsU | 15338 | 374074-389411 | CsU_DRJ1R/L | cys1_CsU, cys2_CsU, cys3_CsU, cys4_CsU, cys5_CsU |
| scaffold_11 | 1376756 | CsA | 6368 | 861628-867995 | CsA_DRJ1R/L | cys_CsA |
| scaffold_17 | 1497664 | CsE | 7990 | 1330025-1338014 | CsE_DRJ1R/L | rep1_CsE, rep2_CsE, rep3_CsE |
| scaffold_12 | 1572921 | IVSP_U37L | 1863 | 104145-106007 | not applicable | U37L-2 |

| | | | | | | |
|---|---|---|---|---|---|---|
| scaffold_23 | 1817383 | CsM | 12197 | 1422227-1434423 | CsM_DRJ1R/L | N_CsM |
| scaffold_5 | 1987530 | CsN | 10943 | 175169-186111 | CsN_DRJ1R/L | N1_CsN, N2_CsN |
| scaffold_38 | 2232507 | CsH | 9050 | 1398066-1407115 | CsH_DRJ1R/L | 5rep_CsH |
| scaffold_16 | 3063130 | Cs_IVSPER-5 | 3750 | 2424942-2428691 | not applicable | IVSP1L-3, U2L, U1L |
| scaffold_16 | 3063130 | CsX6 | 9213 | 504600-513812 | CsX6_DRJ1R/L | rep1_CsX6, rep2_CsX6 |
| scaffold_10 | 3211285 | CsD | 8168 | 961052-969219 | CsD_DRJ1R/L | vnx_CsD |
| scaffold_15 | 5987914 | CsJ | 9484 | 2621922-2631405 | CsJ_DRJ1R/int/L | rep1_CsJ, rep2_CsJ, rep3_CsJ |
| scaffold_13 | 6115246 | CsX4, partial | >9181 | 1407384-1416564 | no repeat found at extremities | rep1_CsX4, rep2_CsX4, rep3_CsX4 |
