## Additional file 4 for "Conserved and specific genomic features of endogenous polydnaviruses revealed by whole genome sequencing of two ichneumonid wasps"

| Scaffold | Proviral segment /IVSPER | Proviral/IVSPER sequence size (nt) | Proviral/IVSPER position in scaffold | Gene name | Gene position in scaffold | Introns in gene (yes/no) | Predicted protein size (aa) | NCBI accession | protein ID | Length NCBI seq. (nt) | BlastP e-value | Identities (%) | Comments |
|---|---|---|---|---|---|---|---|---|---|---|---|---|---|
| **Hyposoter didymator** | | | | | | | | | | | | | |
| **HdIV proviral segment** | | | | | | | | | | | | | |
| scaffold49647 | similar to Hd2 | 1537 | 1-1537 (partial) | Gly-Pro_P40-like, [ | 1-1235 - strand | yes | 315 | AAD40678.1 | P40 protein [Hyposoter didymator ichnovirus] | 397 | 2,00E-168 | 282/318(89%) | |
| scaffold184 | Hd45.2 | 2051 | 3564791-3566841 | U1_Hd45.2 | 3565610-3566474 - strand | yes | 135 | AIK25614.1 | D8 [Hyposoter didymator ichnovirus] | 138 | 4,00E-34 | 82/140(59%) | |
| scaffold128243 | Hd44.1 | 3009 | 4203985-4206993 | U1_Hd44.1 | 4205686-4205997 + strand | no | 104 | AIK25616.1 | U2 [Hyposoter didymator ichnovirus] | 104 | 1,00E-65 | 101/104(97%) | |
| scaffold64 | Hd42 | 3157 | 2320523-2323679 | Rep1_Hd42 | 2322099-2322881 - strand | no | 261 | AIK25619.1 | Rep1 [Hyposoter didymator ichnovirus] | 261 | 0,00E+00 | 248/261(95%) | |
| scaffold128213 | Hd23.2 | 3362 | 248624-251985 | Rep1_Hd23.2 | 249754-250470 - strand | no | 238 | AIK25664.1 | Rep1 [Hyposoter didymator ichnovirus] | 263 | 1,00E-124 | 180/239(75%) | |
| scaffold127348 | Hd40 | 3494 | 926528-930021 | U2_Hd40 | 926730-927180 - strand | no | 148 | AIK25622.1 | U2 [Hyposoter didymator ichnovirus] | 115 | 2,00E-69 | 101/104(97%) | pb (insertion) within CDS |
| scaffold127348 | Hd40 | 3494 | 926528-930021 | U1_Hd40 | 928473-928898 - strand | no | 142 | AIK25621.1 | U1 [Hyposoter didymator ichnovirus] | 143 | 5,00E-75 | 116/142(82%) | |
| scaffold175 | Hd38 | 3664 | 3800393-3804056 | U1_Hd38 | 3800677-3801003 - strand | no | 108 | AIK25624.1 | U1 [Hyposoter didymator ichnovirus] | 108 | 3,00E-69 | 105/108(97%) | insertion of a stop codon within CDS |
| scaffold175 | Hd38 | 3664 | 3800393-3804056 | Vinx1_Hd38 | 3801226-3802356 + strand | no | 377 | AIK25625.1 | Vinx1 [Hyposoter didymator ichnovirus] | 377 | 0,00E+00 | 362/377(96%) | |
| scaffold175 | Hd38 | 3664 | 3800393-3804056 | U2_Hd38 | 3802480-3802809 - strand | no | 109 | AIK25626.1 | F1U2 [Hyposoter didymator ichnovirus] | 107 | 5,00E-68 | 101/107(94%) | insertion of a stop codon within CDS |
| scaffold65 | Hd37 | 3708 | 342293-346000 | U1_Hd37 | 342555-342665 - strand | no | 36 | AIK25627.1 | U1 [Hyposoter didymator ichnovirus] | 143 | 2,00E-16 | 36/36(100%) | partial; corresponds to the N-term of the protein (aa 1-36) |
| scaffold65 | Hd37 | 3708 | 342293-346000 | Rep1_Hd37 | 343245-344048 + strand | no | 268 | AIK25629.1 | Rep1 [Hyposoter didymator ichnovirus] | 268 | 0,00E+00 | 262/268(98%) | |
| scaffold65 | Hd37 | 3708 | 342293-346000 | U2_Hd37 | 344748-344996 - strand | no | 82 | AIK25628.1 | U2 [Hyposoter didymator ichnovirus] | 146 | 1,00E-46 | 76/82(93%) | partial; corresponds to the C-term of the protein (aa 65-146) and one of the |
| scaffold116 | Hd35 | 3710 | 473341-477050 | Rep1_Hd35 | 475117-475821 + strand | no | 235 | AIK25632.1 | Rep1 [Hyposoter didymator ichnovirus] | 235 | 9,00E-177 | 234/235(99%) | |
| scaffold175 | Hd36 | 3738 | 3796140-3799877 | U1_Hd36 | 3797417-3797740 + strand | no | 107 | AIK25630.1 | F1U1 [Hyposoter didymator ichnovirus] | 107 | 2,00E-70 | 102/107(95%) | start codon modified (ATT instead of ATG in NCBI seq) |
| scaffold175 | Hd36 | 3738 | 3796140-3799877 | Vinx1_Hd36 | 3797918-3799006 - strand | no | 363 | AAO16963.1 | viral innexin [Hyposoter didymator ichnovirus] | 363 | 0,00E+00 | 335/363(92%) | |
| scaffold91 | Hd33 | 3835 | 485063-488897 | U1_Hd33 | 485760-486020 - strand | no | 86 | | No significant similarity found | | | | low transcription in parasitized lep larvae |
| scaffold91 | Hd33 | 3835 | 485063-488897 | U2_Hd33 | 487963-488427 - strand | no | 154 | | No significant similarity found | | | | |
| scaffold127548 | Hd27 | 4002 | 13338255-13342256 | U1_Hd27 | 13338842-13339126 + strand | no | 94 | AIK25654.1 | U1 [Hyposoter didymator ichnovirus] | 113 | 6,00E-62 | 92/93(99%) | modified start codon |
| scaffold127548 | Hd27 | 4002 | 13338255-13342256 | K19_Hd27 | 13340915-13341354 - strand | yes | 106 | AAF91314.1 | P12 [Hyposoter didymator ichnovirus] | 106 | 2,00E-71 | 105/106(99%) | |
| scaffold29771 | Hd46 | 4109 | 19344-23452 | Vank_Hd46 | 20304-20807 + strand | no | 167 | YP_00103123 | vankyrin-b1 [Hyposoter fugitivus ichnovirus] | 167 | 9,00E-86 | 127/165(77%) | |
| scaffold127549 | Hd31-34 | 4119 | 144445-148563 | U1_Hd31-34 | 145530-145901 + strand | no | 124 | AIK25633.1 | F4U1 [Hyposoter didymator ichnovirus] | 124 | 1,00E-71 | 121/124(98%) | |
| scaffold127549 | Hd31-34 | 4119 | 144445-148563 | U2_Hd31-34 | 146442-146729 - strand | no | 95 | AIK25634.1 | F5U2 [Hyposoter didymator ichnovirus] | 124 | 3,00E-64 | 94/95(99%) | shorter in C-term |
| scaffold22 | Hd39 | 4122 | 1002620-1006741 | Rep1_Hd39 | 1003991-1004692 - strand | no | 234 | AIK25623.1 | Rep1 [Hyposoter didymator ichnovirus] | 234 | 9,00E-170 | 227/234(97%) | |
| scaffold1868 | Hd43 | 4159 | 193639-197797 | U1_Hd43 | 194377-194676 - strand | no | 100 | AIK25617.1 | U1 [Hyposoter didymator ichnovirus] | 100 | 3,00E-67 | 97/100(97%) | |
| scaffold1868 | Hd43 | 4159 | 193639-197797 | Vank1_Hd43 | 196094-196618 - strand | no | 175 | AIK25618.1 | Vank1 [Hyposoter didymator ichnovirus] | 175 | 1,00E-125 | 171/175(98%) | |
| scaffold67 | Hd30 | 4164 | 5867182-5871345 | U1_Hd30 | 5867451-5867846 + strand | no | 131 | AIK25641.1 | U1 [Hyposoter didymator ichnovirus] | 104 | 3,00E-67 | 100/104(96%) | longuer in C-term |
| scaffold67 | Hd30 | 4164 | 5867182-5871345 | U2_Hd30 | 5867871-5868737 - strand | no | 288 | AIK25642.1 | U2 [Hyposoter didymator ichnovirus] | 288 | 0,00E+00 | 284/288(99%) | |
| scaffold67 | Hd30 | 4164 | 5867182-5871345 | Vinx1_Hd30 | 5869217-5870347 - strand | no | 376 | AIK25643.1 | Vinx1 [Hyposoter didymator ichnovirus] | 376 | 0,00E+00 | 375/376(99%) | |
| scaffold67 | Hd30 | 4164 | 5867182-5871345 | U3_Hd30 | 5870662-5870985 + strand | no | 107 | AIK25644.1 | U3 [Hyposoter didymator ichnovirus] | 107 | 2,00E-68 | 105/107(98%) | |
| scaffold161 | Hd25 | 4174 | 2449400-2453573 | PRRP3_Hd25 | 2450118-2450525 + strand | no | 136 | AIK25660.1 | PRRP2 [Hyposoter didymator ichnovirus] | 136 | 1,00E-83 | 136/136(100%) | |
| scaffold161 | Hd25 | 4174 | 2449400-2453573 | PRRP2_Hd25 | 2451908-2452147 - strand | no | 80 | AIK25661.1 | PRRP3 [Hyposoter didymator ichnovirus] | 80 | 7,00E-46 | 75/80(94%) | |
| scaffold161 | Hd25 | 4174 | 2449400-2453573 | PRRP1_Hd25 | 2452642-2452827 - strand | no | 61 | AIK25659.1 | PRRP1 [Hyposoter didymator ichnovirus] | 61 | 4,00E-09 | 58/61(95%) | |
| scaffold28498 | Hd45.1 | NA | 1-4214 (partial) | U1_Hd45.1 | 679-1541 - strand | yes | 138 | AAO33352.1 | unknown [Hyposoter didymator ichnovirus] | 138 | 3,00E-91 | 133/138(96%) | |
| scaffold28498 | Hd45.1 | NA | 1-4214 (partial) | U2_Hd45.1 | 3317-4180 - strand | yes | 135 | AIK25614.1 | D8 [Hyposoter didymator ichnovirus] | 138 | 3,00E-33 | 82/140(59%) | |
| scaffold119 | Hd22 | 4233 | 698410-702642 | U1_Hd22 | 698775-699101 - strand | no | 109 | AIK25666.1 | U1 [Hyposoter didymator ichnovirus] | 109 | 2,00E-72 | 106/109(97%) | |
| scaffold119 | Hd22 | 4233 | 698410-702642 | Rep1_Hd22 | 700424-701035 - strand | no | 204 | AAR89180.1 | repeat element protein 8 [Hyposoter didymator ichnovirus] | 204 | 9,00E-150 | 204/204(100%) | |
| scaffold91 | Hd29 | 4356 | 572514-576869 | N1_Hd29 | 573680-574969 - strand | no | 430 | AIK25650.1 | N-gene1 [Hyposoter didymator ichnovirus] | 430 | 0,00E+00 | 427/430(99%) | |
| scaffold64 | Hd21 | 4368 | 2353107-2357474 | Rep1_Hd21 | 2353545-2354333 - strand | no | 263 | AIK25667.1 | Rep1 [Hyposoter didymator ichnovirus] | 263 | 0,00E+00 | 258/263(98%) | |
| scaffold64 | Hd21 | 4368 | 2353107-2357474 | Vinx1_Hd21 | 2355781-2356860 - strand | no | 360 | AIK25668.1 | Vinx1 [Hyposoter didymator ichnovirus] | 360 | 0,00E+00 | 356/360(99%) | |
| scaffold198 | Hd19 | 4440 | 1265454-1269893 | U1_Hd19 | 1266250-1268833 - strand | yes | 618 | AIK25671.1 | U1 [Hyposoter didymator ichnovirus] | 618 | 0,00E+00 | 606/618(98%) | |
| scaffold128213 | Hd23.1 | 4457 | 208205-212661 | Rep1_Hd23.1 | 209796-210587 - strand | no | 263 | AIK25664.1 | Rep1 [Hyposoter didymator ichnovirus] | 263 | 0,00E+00 | 262/263(99%) | |
| scaffold127548 | Hd47 | 4503 | 12134587-12139089 | Rep1_Hd47 | 12135113-12135841 - strand | no | 242 | YP_00103131 | repeat element protein-d4.2 [Hyposoter fugitivus ichnovirus] | 248 | 2,00E-122 | 166/238(70%) | |
| scaffold127548 | Hd47 | 4503 | 12134587-12139089 | Rep2_Hd47 | 12136931-12137599 - strand | no | 222 | YP_00103131 | repeat element protein-d4.1 [Hyposoter fugitivus ichnovirus] | 255 | 4,00E-116 | 154/220(70%) | |
| scaffold264 | Hd28 | 4614 | 135485-140098 | Vinx1_Hd28 | 136611-137621 + strand | no | 336 | AIK25662.1 | Vinx1 [Hyposoter didymator ichnovirus] | 357 | 0,00E+00 | 301/334(90%) | |
| scaffold264 | Hd28 | 4614 | 135485-140098 | Vank1_Hd28 | 138475-138984 + strand | no | 170 | AFH35114.1 | vankyrin 1 [Hyposoter didymator ichnovirus] | 170 | 2,00E-122 | 168/170(99%) | |
| scaffold82201 | Hd51 | 4632 | 1077-5708 | Rep1_Hd51 | 2537-3115 - strand | no | 192 | AIK25629.1 | Rep1 [Hyposoter didymator ichnovirus] | 268 | 2,00E-89 | 129/190(68%) | no reads RNAseq |
| scaffold351 | Hd18 | 4696 | 2681961-2686656 | N1_Hd18 | 2683737-2685050 + strand | no | 438 | AIK25675.1 | N-gene1 [Hyposoter didymator ichnovirus] | 438 | 0,00E+00 | 434/438(99%) | |
| scaffold91 | Hd24 | 4697 | 535698-540394 | Vank1_Hd24 | 536918-537427 - strand | no | 170 | AFH35112.1 | vankyrin 1 [Hyposoter didymator ichnovirus] | 170 | 1,00E-123 | 170/170(100%) | |
| scaffold91 | Hd24 | 4697 | 535698-540394 | Vinx1_Hd24 | 538200-539270 - strand | no | 357 | AIK25662.1 | Vinx1 [Hyposoter didymator ichnovirus] | 357 | 0,00E+00 | 351/357(98%) | |
| scaffold128243 | Hd44.2 | 4831 | 4197203-4202033 | Rep1_Hd44.2 | 4197838-4198440 - strand | no | 200 | YP_00103128 | repeat element protein-c11.1 [Hyposoter fugitivus ichnovirus] | 201 | 3,00E-58 | 106/186(57%) | |
| scaffold128243 | Hd44.2 | 4831 | 4197203-4202033 | U1_Hd44.2 | 4199950-4200297 + strand | no | 115 | AIG88525.1 | hypothetical protein A7.1 [Diadegma fenestrale ichnovirus] | 76 | 1,00E-20 | 56/76(74%) | strand may be incorrect |
| scaffold91 | Hd15 | 4987 | 469105-474091 | N1_Hd15 | 470135-471607 - strand | no | 491 | AIK25681.1 | N-gene1 [Hyposoter didymator ichnovirus] | 491 | 0,00E+00 | 478/491(97%) | |
| scaffold175 | Hd26 | 5018 | 10942034-10947051 | PRRP1_Hd26 | 10943182-10943589 + strand | no | 136 | AIK25646.1 | PRRP1 [Hyposoter didymator ichnovirus] | 136 | 2,00E-91 | 136/136(100%) | |

| | | | | | | | | | | | | | |
|---|---|---|---|---|---|---|---|---|---|---|---|---|---|
| scaffold175 | Hd26 | 5018 | 10942034-10947051 | PRRP2_Hd26 | 10945126-10945518 + strand | no | 131 | AIK25645.1 | PRRP2 [Hyposoter didymator ichnovirus] | 131 | 4,00E-83 | 130/131(99%) | |
| scaffold175 | Hd26 | 5018 | 10942034-10947051 | U1_Hd26 | 10946124-10946471 - strand | no | 116 | AIK25658.1 | U1 [Hyposoter didymator ichnovirus] | 116 | 2,00E-80 | 116/116(100%) | |
| scaffold64 | Hd14 | 5196 | 36336-41531 | Vinx1_Hd14 | 38977-39405 + strand | no | 143 | AIK25682.1 | Vinx1 [Hyposoter didymator ichnovirus] | 143 | 5,00E-101 | 143/143(100%) | |
| scaffold64 | Hd14 | 5196 | 36336-41531 | U1_Hd14 | 41196-41501 + strand | no | 102 | AIK25683.1 | U1 [Hyposoter didymator ichnovirus] | 102 | 8,00E-70 | 102/102(100%) | |
| scaffold128246 | Hd49 | 5265 | 677866-683130 | Rep1_Hd49 | 678304-679005 + strand | no | 233 | YP_001031325.1 | repeat element protein-d7.2 [Hyposoter fugitivus ichnovirus] | 242 | 1,00E-119 | 173/232(75%) | |
| scaffold128246 | Hd49 | 5265 | 677866-683130 | Rep2_Hd49 | 679573-680190 + strand | no | 205 | YP_001031326.1 | repeat element protein-d7.3 [Hyposoter fugitivus ichnovirus] | 240 | 3,00E-114 | 160/205(78%) | |
| scaffold128246 | Hd49 | 5265 | 677866-683130 | Rep3_Hd49 | 681373-682041 + strand | no | 222 | YP_001031323.1 | repeat element protein-d7.1 [Hyposoter fugitivus ichnovirus] | 255 | 5,00E-93 | 156/257(61%) | |
| scaffold128241 | Hd13 | 5757 | 402487-408243 | Cys2_Hd13 | 403839-404424 - strand | yes | 157 | AIK25685.1 | Cys2 [Hyposoter didymator ichnovirus] | 157 | 5,00E-94 | 133/157(85%) | |
| scaffold128241 | Hd13 | 5757 | 402487-408243 | Cys1_Hd13 | 406027-406868 - strand | yes | 162 | AIK25684.1 | Cys1 [Hyposoter didymator ichnovirus] | 162 | 1,00E-107 | 149/162(92%) | |
| scaffold357 | Hd50 | 5787 | 2218373-2224159 | Vinx1_Hd50 | 2219976-2221058 + strand | no | 360 | YP_001031228.1 | innexin Vnx-d5.1 [Hyposoter fugitivus ichnovirus] | 375 | 0,00E+00 | 280/357(78%) | |
| scaffold357 | Hd50 | 5787 | 2218373-2224159 | Vinx2_Hd50 | 2221574-2222647 + strand | no | 357 | YP_001031229.1 | innexin Vnx-d5.2 [Hyposoter fugitivus ichnovirus] | 378 | 7,00E-170 | 241/378(64%) | |
| scaffold59 | Hd12 | 5902 | 674917-680818 | U1_Hd12 | 675013-675345 + strand | no | 111 | AIK25686.1 | U1 [Hyposoter didymator ichnovirus] | 111 | 9,00E-77 | 110/111(99%) | |
| scaffold59 | Hd12 | 5902 | 674917-680818 | Rep1_Hd12 | 675996-676679 - strand | no | 228 | AIK25689.1 | Rep1 [Hyposoter didymator ichnovirus] | 228 | 1,00E-169 | 226/228(99%) | |
| scaffold59 | Hd12 | 5902 | 674917-680818 | U2_Hd12 | 678859-679170 - strand | no | 104 | AIK25687.1 | U2 [Hyposoter didymator ichnovirus] | 104 | 2,00E-67 | 103/104(99%) | |
| scaffold59 | Hd12 | 5902 | 674917-680818 | Rep2_Hd12 | 679544-680233 - strand | no | 230 | AIK25688.1 | Rep2 [Hyposoter didymator ichnovirus] | 230 | 1,00E-173 | 229/230(99%) | |
| scaffold59 | Hd10 | 6507 | 2500377-2506883 | Rep3_Hd10 | 2501422-2502045 - strand | no | 208 | AIK25696.1 | Rep3 [Hyposoter didymator ichnovirus] | 208 | 9,00E-154 | 207/208(99%) | |
| scaffold59 | Hd10 | 6507 | 2500377-2506883 | Rep2_Hd10 | 2502434-2503066 - strand | no | 211 | AIK25695.1 | Rep2 [Hyposoter didymator ichnovirus] | 211 | 3,00E-155 | 211/211(100%) | |
| scaffold59 | Hd10 | 6507 | 2500377-2506883 | Rep1_Hd10 | 2504201-2504872 - strand | no | 224 | AAR89179.1 | repeat element protein 7 [Hyposoter didymator ichnovirus] | 224 | 4,00E-167 | 224/224(100%) | |
| scaffold144 | Hd20 | 6864 | 1449468-1456331 | Cys2_Hd20 | 1450485-1451362 - strand | yes | 145 | AIK25636.1 | Cys1 [Hyposoter didymator ichnovirus] | 145 | 3,00E-104 | 144/145(99%) | |
| scaffold144 | Hd20 | 6864 | 1449468-1456331 | Cys1_Hd20 | 1454299-1455511 - strand | yes | 254 | AIK25669.1 | Cys1 [Hyposoter didymator ichnovirus] | 254 | 0,00E+00 | 252/254(99%) | |
| scaffold377 | Hd8 | 7356 | 2186417-2193772 | U5_Hd8 | 2186651-2186968 + strand | no | 105 | AIK25705.1 | U4 [Hyposoter didymator ichnovirus] | 105 | 2,00E-71 | 103/105(98%) | |
| scaffold377 | Hd8 | 7356 | 2186417-2193772 | U4_Hd8 | 2188345-2188632 - strand | no | 95 | YP_00103136 | e1.1 [Hyposoter fugitivus ichnovirus] | 131 | 4,00E-07 | 46/131(35%) | |
| scaffold377 | Hd8 | 7356 | 2186417-2193772 | U3_Hd8 | 2190149-2190466 + strand | no | 105 | AIK25702.1 | F6U1 [Hyposoter didymator ichnovirus] | 105 | 5,00E-71 | 102/105(97%) | |
| scaffold377 | Hd8 | 7356 | 2186417-2193772 | U2_Hd8 | 2191793-2192095 - strand | no | 100 | AIK25703.1 | U2 [Hyposoter didymator ichnovirus] | 100 | 1,00E-64 | 98/100(98%) | |
| scaffold377 | Hd8 | 7356 | 2186417-2193772 | U1_Hd8 | 2192394-2192693 + strand | no | 99 | AIK25704.1 | F6U3 [Hyposoter didymator ichnovirus] | 106 | 6,00E-44 | 71/75(95%) | |
| scaffold59 | Hd16 | 7704 | 690205-697908 | Rep1_Hd16 | 690229-690951 - strand | no | 241 | AIK25680.1 | Rep1 [Hyposoter didymator ichnovirus] | 241 | 0,00E+00 | 241/241(100%) | |
| scaffold59 | Hd16 | 7704 | 690205-697908 | U2_Hd16 | 691425-691661 + strand | no | 78 | | No significant similarity found | | | | possibly not a CDS (low transcription in calyx) |
| scaffold59 | Hd16 | 7704 | 690205-697908 | U1_Hd16 | 693257-693817 + strand | no | 187 | AIK25679.1 | U1 [Hyposoter didymator ichnovirus] | 187 | 9,00E-122 | 186/187(99%) | |
| scaffold59 | Hd16 | 7704 | 690205-697908 | Rep2_Hd16 | 695154-695813 - strand | no | 219 | AAR89178.1 | repeat element protein 6 [Hyposoter didymator ichnovirus] | 219 | 1,00E-159 | 213/219(97%) | |
| scaffold59 | Hd16 | 7704 | 690205-697908 | Rep3_Hd16 | 696463-697149 - strand | no | 228 | AAR89177.1 | repeat element protein 5 [Hyposoter didymator ichnovirus] | 228 | 9,00E-170 | 228/228(100%) | |
| scaffold351 | Hd17 | 7730 | 2329273-2337002 | Rep5_Hd17 | 2329743-2330387 - strand | no | 214 | AHY22036.1 | repeat element 36 [Diadegma semiclausum ichnovirus] | 262 | 3,00E-124 | 174/214(81%) | |
| scaffold351 | Hd17 | 7730 | 2329273-2337002 | Rep4_Hd17 | 2330936-2331505 - strand | no | 189 | AIK25678.1 | Rep1 [Hyposoter didymator ichnovirus] | 230 | 1,00E-120 | 166/182(91%) | |
| scaffold351 | Hd17 | 7730 | 2329273-2337002 | Rep3_Hd17 | 2332279-2333010 - strand | no | 244 | AAO16957.1 | repeat element protein [Hyposoter didymator ichnovirus] | 244 | 0,00E+00 | 244/244(100%) | |
| scaffold351 | Hd17 | 7730 | 2329273-2337002 | Rep2_Hd17 | 2334085-2334759 - strand | no | 225 | AAO16959.1 | repeat element protein [Hyposoter didymator ichnovirus] | 225 | 5,00E-166 | 221/225(98%) | |
| scaffold351 | Hd17 | 7730 | 2329273-2337002 | Rep1_Hd17 | 2335459-2336166 - strand | no | 236 | AIK25673.1 | Rep1 [Hyposoter didymator ichnovirus] | 236 | 4,00E-173 | 232/236(98%) | |
| scaffold64 | Hd32 | 7916 | 88702-96617 | U1.2_Hd32 | 88842-89159 - strand | no | 105 | AIK25638.1 | U1 [Hyposoter didymator ichnovirus] | 105 | 5,00E-65 | 96/105(91%) | |
| scaffold64 | Hd32 | 7916 | 88702-96617 | Vinx2_Hd32 | 90142-91254 - strand | no | 370 | AIK25637.1 | Vinx1 [Hyposoter didymator ichnovirus] | 366 | 0,00E+00 | 279/353(79%) | |
| scaffold64 | Hd32 | 7916 | 88702-96617 | U1.1_Hd32 | 93443-93757 - strand | no | 105 | AIK25638.1 | U1 [Hyposoter didymator ichnovirus] | 105 | 6,00E-72 | 105/105(100%) | |
| scaffold64 | Hd32 | 7916 | 88702-96617 | Vinx1_Hd32 | 94429-95526 - strand | no | 366 | AIK25637.1 | Vinx1 [Hyposoter didymator ichnovirus] | 366 | 0,00E+00 | 366/366(100%) | |
| scaffold184 | Hd41 | 7953 | 3768924-3776876 | U1_Hd41 | 3769919-3770824 + strand | yes | 152 | AIK25620.1 | U1 [Hyposoter didymator ichnovirus] | 197 | 9,00E-64 | 112/152(74%) | N-term shorter |
| scaffold184 | Hd41 | 7953 | 3768924-3776876 | U2_Hd41 | 3772675-3773516 + strand | yes | 168 | AIK25614.1 | D8 [Hyposoter didymator ichnovirus] | 138 | 3,00E-22 | 72/170(42%) | |
| scaffold184 | Hd41 | 7953 | 3768924-3776876 | U3_Hd41 | 3775212-3776089 + strand | yes | 159 | AIK25620.1 | U1 [Hyposoter didymator ichnovirus] | 197 | 2,00E-22 | 59/96(61%) | |
| scaffold127548 | Hd7 | 8066 | 6062921-6070986 | U1_Hd7 | 6065301-6068372 + strand | no | 1024 | AIK25706.1 | U1 [Hyposoter didymator ichnovirus] | 1080 | 0,00E+00 | 916/1080(85%) | repeated motif missing from the CDS |
| scaffold59 | Hd11 | 9190 | 2183512-2192701 | U1_Hd11 | 2184311-2184694 + strand | no | 128 | AIK25649.1 | U1 [Hyposoter didymator ichnovirus] | 128 | 8,00E-87 | 125/128(98%) | |
| scaffold59 | Hd11 | 9190 | 2183512-2192701 | Rep1_Hd11 | 2184915-2185607 + strand | no | 231 | AIK25648.1 | Rep1 [Hyposoter didymator ichnovirus] | 231 | 7,00E-172 | 229/231(99%) | |
| scaffold59 | Hd11 | 9190 | 2183512-2192701 | Vank1_Hd11 | 2186326-2186802 + strand | no | 159 | AFH35113.1 | vankyrin 1 [Hyposoter didymator ichnovirus] | 159 | 2,00E-113 | 159/159(100%) | |
| scaffold59 | Hd11 | 9190 | 2183512-2192701 | Vank5_Hd11 | 2187892-2188395 + strand | no | 168 | AFH35119.1 | vankyrin 5 [Hyposoter didymator ichnovirus] | 168 | 2,00E-123 | 168/168(100%) | |
| scaffold59 | Hd11 | 9190 | 2183512-2192701 | Vank4_Hd11 | 2189456-2189962 + strand | no | 169 | AFH35118.1 | vankyrin 4 [Hyposoter didymator ichnovirus] | 169 | 4,00E-123 | 168/169(99%) | |
| scaffold59 | Hd11 | 9190 | 2183512-2192701 | Vank3_Hd11 | 2190561-2191067 + strand | no | 169 | AFH35117.1 | vankyrin 3 [Hyposoter didymator ichnovirus] | 169 | 1,00E-121 | 168/169(99%) | |
| scaffold59 | Hd11 | 9190 | 2183512-2192701 | Vank2_Hd11 | 2191436-2191942 + strand | no | 169 | AFH35116.1 | vankyrin 2 [Hyposoter didymator ichnovirus] | 169 | 2,00E-116 | 163/169(96%) | |
| scaffold59 | Hd11 | 9190 | 2183512-2192701 | Vank1p_Hd11 | 2192415-2192701 + strand | no | 95 | AFH35113.1 | vankyrin 1 [Hyposoter didymator ichnovirus] | 159 | 2,00E-58 | 88/95(93%) | partial; corresponds to the N-term of the protein (aa 1-95) |
| scaffold429 | Hd48 | 9673 | 531833-541505 | Rep1_Hd48 | 533753-534487 - strand | no | 244 | AHY22018.1 | repeat element 27 [Diadegma semiclausum ichnovirus] | 246 | 3,00E-127 | 175/245(71%) | |
| scaffold429 | Hd48 | 9673 | 531833-541505 | Rep2_Hd48 | 538812-539549 - strand | no | 245 | YP_001031283.1 | repeat element protein-c7.1 [Hyposoter fugitivus ichnovirus] | 282 | 4,00E-133 | 179/244(73%) | |

| | | | | | | | | | | | | | |
|---|---|---|---|---|---|---|---|---|---|---|---|---|---|
| scaffold65 | Hd3 | 10014 | 437082-447095 | Cys1_Hd3 | 438828-439558 - strand | yes | 181 | AIK25727.1 | Cys1 [Hyposoter didymator ichnovirus] | 181 | 4,00E-136 | 181/181(100%) | |
| scaffold65 | Hd3 | 10014 | 437082-447095 | Cys2_Hd3 | 440422-441241 - strand | yes | 200 | AIK25728.1 | Cys2 [Hyposoter didymator ichnovirus] | 200 | 1,00E-147 | 200/200(100%) | |
| scaffold65 | Hd3 | 10014 | 437082-447095 | Cys3_Hd3 | 442538-443060 - strand | yes | 147 | AIK25729.1 | Cys3 [Hyposoter didymator ichnovirus] | 147 | 3,00E-105 | 144/147(98%) | |
| scaffold65 | Hd3 | 10014 | 437082-447095 | Cys4_Hd3 | 443574-444657 - strand | yes | 267 | AIK25730.1 | Cys4 [Hyposoter didymator ichnovirus] | 241 | 3,00E-168 | 232/267(87%) | |
| scaffold65 | Hd3 | 10014 | 437082-447095 | Cys5_Hd3 | 445264-445755 - strand | yes | 125 | AIK25731.1 | Cys5 [Hyposoter didymator ichnovirus] | 125 | 3,00E-83 | 117/125(94%) | |
| scaffold377 | Hd4 | 10326 | 2459681-2470006 | Rep1_Hd4 | 2461072-2461818 - strand | no | 248 | AIK25721.1 | Rep1 [Hyposoter didymator ichnovirus] | 248 | 0,00E+00 | 248/248(100%) | |
| scaffold377 | Hd4 | 10326 | 2459681-2470006 | Rep2_Hd4 | 2462404-2463012 + strand | no | 202 | AIK25722.1 | Rep2 [Hyposoter didymator ichnovirus] | 202 | 8,00E-144 | 197/202(98%) | |
| scaffold377 | Hd4 | 10326 | 2459681-2470006 | Rep3_Hd4 | 2463770-2464366 + strand | no | 198 | AIK25723.1 | Rep3 [Hyposoter didymator ichnovirus] | 198 | 2,00E-145 | 198/198(100%) | |
| scaffold377 | Hd4 | 10326 | 2459681-2470006 | Rep4_Hd4 | 2465099-2465706 + strand | no | 202 | AIK25724.1 | Rep4 [Hyposoter didymator ichnovirus] | 202 | 6,00E-150 | 202/202(100%) | |
| scaffold377 | Hd4 | 10326 | 2459681-2470006 | Rep5_Hd4 | 2466854-2467603 - strand | no | 249 | AIK25725.1 | Rep5 [Hyposoter didymator ichnovirus] | 249 | 0,00E+00 | 246/249(99%) | |
| scaffold377 | Hd4 | 10326 | 2459681-2470006 | Rep6_Hd4 | 2468587-2469195 + strand | no | 202 | AIK25726.1 | Rep6 [Hyposoter didymator ichnovirus] | 202 | 3,00E-149 | 200/202(99%) | |
| scaffold127548 | Hd6 | 10461 | 5808388-5818848 | Rep1_Hd6 | 5809242-5809907 + strand | no | 221 | AAO33572.1 | rep protein [Hyposoter didymator ichnovirus] | 221 | 1,00E-164 | 221/221(100%) | |
| scaffold127548 | Hd6 | 10461 | 5808388-5818848 | P30_Hd6 | 5810559-5811315 - strand | yes | 159 | AIK25713.1 | P30 [Hyposoter didymator ichnovirus] | 414 | 2,00E-64 | 121/131(92%) | repeated motifs missing from the CDS |
| scaffold127548 | Hd6 | 10461 | 5808388-5818848 | U1.1_Hd6 | 5814030-5814758 - strand | yes | 151 | AIK25712.1 | U1 [Hyposoter didymator ichnovirus] | 159 | 2,00E-94 | 151/159(95%) | |
| scaffold127548 | Hd6 | 10461 | 5808388-5818848 | U1.2_Hd6 | 5818232-5818829 - strand | yes | 143 | AIK25712.1 | U1 [Hyposoter didymator ichnovirus] | 159 | 7,00E-12 | 53/129(41%) | |
| scaffold127548 | Hd5 | 10510 | 12941242-12951751 | Vinx1_Hd5 | 12941923-12942945 - strand | no | 340 | AIK25715.1 | Vinx1 [Hyposoter didymator ichnovirus] | 340 | 0,00E+00 | 336/340(99%) | |
| scaffold127548 | Hd5 | 10510 | 12941242-12951751 | Vinx2_Hd5 | 12943091-12943896 - strand | no | 267 | AIK25716.1 | Vinx2 [Hyposoter didymator ichnovirus] | 180 | 1,00E-99 | 141/143(99%) | pb within CDS |
| scaffold127548 | Hd5 | 10510 | 12941242-12951751 | U1_Hd5 | 12944615-12944923 - strand | no | 102 | AIK25717.1 | U1 [Hyposoter didymator ichnovirus] | 102 | 1,00E-65 | 98/102(96%) | |
| scaffold127548 | Hd5 | 10510 | 12941242-12951751 | Vinx3_Hd5 | 12945139-12946200 - strand | no | 353 | AIK25718.1 | Vinx3 [Hyposoter didymator ichnovirus] | 353 | 0,00E+00 | 352/353(99%) | |
| scaffold127548 | Hd5 | 10510 | 12941242-12951751 | Vinx4_Hd5 | 12947754-12948800 - strand | no | 348 | AIK25719.1 | Vinx4 [Hyposoter didymator ichnovirus] | 348 | 0,00E+00 | 346/348(99%) | |
| scaffold127548 | Hd5 | 10510 | 12941242-12951751 | Vinx5_Hd5 | 12950651-12951697 - strand | no | 348 | AAR82838.1 | innexin-like protein 2 [Hyposoter didymator ichnovirus] | 348 | 0,00E+00 | 346/348(99%) | |
| scaffold127548 | Hd5 | 10510 | 12941242-12951751 | Vinx6_Hd5 | 12952927-12953967 - strand | no | 346 | AAR82840.1 | innexin-like protein 4 [Hyposoter didymator ichnovirus] | 393 | 0,00E+00 | 334/343(97%) | |
| scaffold127548 | Hd2 | 13937 | 5940296-5954232 | GlyPro1_Hd2 | 5941600-5943451 + strand | yes | 452 | AAF08193.1 | glycine and proline-rich protein P45 precursor [Hyposoter didymator ichnovirus] | 452 | 0,00E+00 | 451/452(99%) | |
| scaffold127548 | Hd2 | 13937 | 5940296-5954232 | U1_Hd2 | 5944265-5944609 - strand | no | 115 | AIK25733.1 | F2U1 [Hyposoter didymator ichnovirus] | 115 | 5,00E-77 | 115/115(100%) | |
| scaffold127548 | Hd2 | 13937 | 5940296-5954232 | U2_Hd2 | 5945077-5945436 + strand | no | 120 | AAO33351.1 | unknown [Hyposoter didymator ichnovirus] | 120 | 8,00E-79 | 120/120(100%) | |
| scaffold127548 | Hd2 | 13937 | 5940296-5954232 | GlyPro2_Hd2 | 5946971-5951238 + strand | yes | 1357 | AAF08192.1 | glycine and proline-rich protein P69 precursor [Hyposoter didymator ichnovirus] | 683 | 6,00E-157 | 417/786(53%) | differences in the repeated region |
| scaffold127548 | Hd2 | 13937 | 5940296-5954232 | SerThr1_Hd2 | 5952553-5953461 + strand | yes | 216 | AIK25709.1 | SerThr [Hyposoter didymator ichnovirus] | 216 | 2,00E-153 | 211/216(98%) | |
| scaffold90 | Hd1 | NA | 1-14771 (partial) | U6_Hd1 | 3039-3506 + strand | no | 156 | AIK25750.1 | F9U13 [Hyposoter didymator ichnovirus] | 156 | 6,00E-109 | 153/156(98%) | |
| scaffold90 | Hd1 | NA | 1-14771 (partial) | U5_Hd1 | 4779-5282 - strand | no | 168 | AIK25739.1 | F8U2 [Hyposoter didymator ichnovirus] | 168 | 9,00E-119 | 168/168(100%) | |
| scaffold90 | Hd1 | NA | 1-14771 (partial) | U4_Hd1 | 6629-7054 + strand | no | 142 | AIK25738.1 | F7U1 [Hyposoter didymator ichnovirus] | 142 | 2,00E-99 | 142/142(100%) | |
| scaffold90 | Hd1 | NA | 1-14771 (partial) | U3_Hd1 | 8197-8511 - strand | no | 105 | AIK25745.1 | F11U8 [Hyposoter didymator ichnovirus] | 106 | 2,00E-46 | 82/106(77%) | |
| scaffold90 | Hd1 | NA | 1-14771 (partial) | U2_Hd1 | 8984-9325 - strand | no | 114 | AIK25746.1 | U9 [Hyposoter didymator ichnovirus] | 114 | 2,00E-79 | 114/114(100%) | |
| scaffold90 | Hd1 | NA | 1-14771 (partial) | U1_Hd1 | 12650-12967 - strand | no | 105 | AIK25745.1 | F11U8 [Hyposoter didymator ichnovirus] | 106 | 2,00E-46 | 82/106(77%) | |
| scaffold128215 | Hd9 | 17892 | 526917-544808 | U1.3_Hd9 | 527725-528057 - strand | no | 110 | AIK25698.1 | U1 [Hyposoter didymator ichnovirus] | 111 | 3,00E-32 | 81/110(74%) | |
| scaffold128215 | Hd9 | 17892 | 526917-544808 | U5.2_Hd9 | 529943-530425 + strand | no | 160 | | No significant similarity found | | | | |
| scaffold128215 | Hd9 | 17892 | 526917-544808 | U4.2_Hd9 | 531702-532277 - strand | no | 191 | AIK25701.1 | U4 [Hyposoter didymator ichnovirus] | 116 | 1,00E-48 | 99/116(85%) | |
| scaffold128215 | Hd9 | 17892 | 526917-544808 | U3.2_Hd9 | 532913-533263 + strand | no | 117 | AIK25700.1 | U3 [Hyposoter didymator ichnovirus] | 117 | 9,00E-76 | 112/117(96%) | |
| scaffold128215 | Hd9 | 17892 | 526917-544808 | U6.2_Hd9 | 533349-533735 + strand | no | 128 | AIK25748.1 | F7U11 [Hyposoter didymator ichnovirus] | 111 | 6,00E-18 | 56/126(44%) | |
| scaffold128215 | Hd9 | 17892 | 526917-544808 | U1.2_Hd9 | 535076-535489 - strand | no | 137 | AIK25698.1 | U1 [Hyposoter didymator ichnovirus] | 111 | 2,00E-30 | 76/110(69%) | |
| scaffold128215 | Hd9 | 17892 | 526917-544808 | U5.1_Hd9 | 537399-537881 + strand | no | 160 | | No significant similarity found | | | | |
| scaffold128215 | Hd9 | 17892 | 526917-544808 | U4.1_Hd9 | 539200-539775 - strand | no | 191 | AIK25701.1 | U4 [Hyposoter didymator ichnovirus] | 116 | 5,00E-48 | 99/116(85%) | |
| scaffold128215 | Hd9 | 17892 | 526917-544808 | U3.1_Hd9 | 540411-540941 + strand | no | 176 | AIK25700.1 | U3 [Hyposoter didymator ichnovirus] | 117 | 2,00E-72 | 110/117(94%) | |
| scaffold128215 | Hd9 | 17892 | 526917-544808 | U6.1_Hd9 | 540847-541233 + strand | no | 128 | AIK25748.1 | F7U11 [Hyposoter didymator ichnovirus] | 111 | 8,00E-18 | 60/128(47%) | |
| scaffold128215 | Hd9 | 17892 | 526917-544808 | U2_Hd9 | 541468-541782 - strand | no | 105 | AIK25699.1 | U2 [Hyposoter didymator ichnovirus] | 104 | 3,00E-67 | 102/105(97%) | |
| scaffold128215 | Hd9 | 17892 | 526917-544808 | U1.1_Hd9 | 543144-543476 - strand | no | 111 | AIK25698.1 | U1 [Hyposoter didymator ichnovirus] | 111 | 3,00E-70 | 110/111(99%) | |
| ***H. didymator* IVSPER** | | | | | | | | | | | | | |
| scaffold127548 | IVSPER-5 | 1629 | 10860001-10861630 | U35 | 10860001-10860537 + strand | no | 178 | AKD28058.1 | hypothetical protein [Glypta fumiferanae] gene="U26" | 180 | 1,00E-29 | 60/150(40%) | transcribed in calyx |
| scaffold127548 | IVSPER-5 | 1629 | 10860001-10861630 | U36 | 10861166-10861630 - strand | no | 154 | AKD28080.1 | hypothetical protein [Glypta fumiferanae] gene="U38" | 175 | 3,00E-14 | 45/144(31%) | transcribed in calyx |
| scaffold29771 | IVSP_U37 | 1839 | 16848-18686 | single CDS: U37 | 16848-18686 + strand | no | 612 | AKD28048.1 | helicase-primase domain [Glypta fumiferanae] | 721 | 0,00E+00 | 340/625(54%) | |
| scaffold91 | IVSPER-1 | 14020 | 453857-467876 | U1 | 453857-454591 + strand | no | 244 | ADI40452.1 | unknown [Hyposoter didymator] | 244 | 0,00E+00 | 241/243(99%) | |
| scaffold91 | IVSPER-1 | 14020 | 453857-467876 | IVSP1-1 | 455614-456375 + strand | no | 253 | ADI40453.1 | unknown [Hyposoter didymator] | 253 | 0,00E+00 | 245/253(97%) | |
| scaffold91 | IVSPER-1 | 14020 | 453857-467876 | U2 | 456897-457582 + strand | no | 228 | ADI40454.1 | unknown [Hyposoter didymator] | 228 | 3,00E-171 | 228/228(100%) | |
| scaffold91 | IVSPER-1 | 14020 | 453857-467876 | U3 | 458215-458709 + strand | no | 164 | ADI40455.1 | unknown [Hyposoter didymator] | 162 | 2,00E-119 | 162/162(100%) | |
| scaffold91 | IVSPER-1 | 14020 | 453857-467876 | U4 | 459123-459518 + strand | no | 131 | ADI40456.1 | unknown [Hyposoter didymator] | 131 | 5,00E-91 | 130/131(99%) | |
| scaffold91 | IVSPER-1 | 14020 | 453857-467876 | p53-2 | 460033-461034 + strand | no | 333 | ADI40457.1 | unknown [Hyposoter didymator] | 333 | 0,00E+00 | 329/333(99%) | |
| scaffold91 | IVSPER-1 | 14020 | 453857-467876 | U5 | 461470-462000 + strand | no | 176 | ADI40458.1 | unknown [Hyposoter didymator] | 176 | 6,00E-129 | 175/176(99%) | |
| scaffold91 | IVSPER-1 | 14020 | 453857-467876 | IVSP2-1 | 462381-463910 + strand | no | 509 | ADI40459.1 | unknown [Hyposoter didymator] | 509 | 0,00E+00 | 508/509(99%) | |
| scaffold91 | IVSPER-1 | 14020 | 453857-467876 | N-1 | 464563-466050 + strand | no | 495 | ADI40460.1 | unknown [Hyposoter didymator] | 470 | 0,00E+00 | 467/470(99%) | N-term longuer |
| scaffold91 | IVSPER-1 | 14020 | 453857-467876 | U25 | 467172-467876 - strand | no | 234 | AKD28083.1 | ring finger domain [Glypta fumiferanae] | 237 | 7,00E-14 | 37/118(31%) | putative IVSPER gene; may be wasp gene |

| | | | | | | | | | | | | | |
|---|---|---|---|---|---|---|---|---|---|---|---|---|---|
| scaffold127548 | IVSPER-4 | 15811 | 6832835-6848646 | U29 | 6832835-6833713 - strand | no | 292 | | No significant similarity found | | | | highly transcribed in calyx |
| scaffold127548 | IVSPER-4 | 15811 | 6832835-6848646 | U30 | 6834523-6838368 - strand | no | 1281 | AKD28060.1 | hypothetical protein [Glypta fumiferanae] gene="U28" | 1322 | 0,00E+00 | 616/1339(46%) | |
| scaffold127548 | IVSPER-4 | 15811 | 6832835-6848646 | U31 | 6839646-6840788 + strand | no | 380 | AKD28063.1 | hypothetical protein [Glypta fumiferanae] gene="U31" | 370 | 8,00E-27 | 93/323(29%) | transcribed in calyx |
| scaffold127548 | IVSPER-4 | 15811 | 6832835-6848646 | U32 | 6841304-6841912 + strand | no | 202 | | No significant similarity found | | | | transcribed in calyx |
| scaffold127548 | IVSPER-4 | 15811 | 6832835-6848646 | U33 | 6842790-6844940 - strand | no | 716 | | No significant similarity found | | | | transcribed in calyx |
| scaffold127548 | IVSPER-4 | 15811 | 6832835-6848646 | U34 | 6845548-6848646 - strand | no | 1032 | AKD28054.1 | helicase domain [Glypta fumiferanae] | 1012 | 0,00E+00 | 488/1040(47%) | |
| scaffold127548 | IVSPER-3 | 25432 | 10761570-10787001 | U15 | 10761570-10762802 - strand | no | 410 | ADI40477.1 | unknown [Hyposoter didymator] | 410 | 0,00E+00 | 407/410(99%) | |
| scaffold127548 | IVSPER-3 | 25432 | 10761570-10787001 | IVSP3-2 | 10763133-10765037 + strand | no | 634 | ADI40478.1 | unknown [Hyposoter didymator] | 527 | 0,00E+00 | 526/527(99%) | N-term 100aa longuer in genomic seq |
| scaffold127548 | IVSPER-3 | 25432 | 10761570-10787001 | U16 | 10765894-10767732 - strand | no | 612 | ADI40479.1 | unknown [Hyposoter didymator] | 612 | 0,00E+00 | 612/612(100%) | |
| scaffold127548 | IVSPER-3 | 25432 | 10761570-10787001 | U17 | 10768078-10768325 + strand | no | 82 | ADI40480.1 | unknown [Hyposoter didymator] | 82 | 1,00E-53 | 82/82(100%) | |
| scaffold127548 | IVSPER-3 | 25432 | 10761570-10787001 | U18 | 10769122-10769379 + strand | no | 85 | ADI40481.1 | unknown [Hyposoter didymator] | 85 | 7,00E-53 | 84/85(99%) | |
| scaffold127548 | IVSPER-3 | 25432 | 10761570-10787001 | p12-1 | 10770065-10770298 + strand | no | 77 | CAR31591.1 | p12-like1 protein [Hyposoter didymator] | 77 | 6,00E-45 | 77/77(100%) | |
| scaffold127548 | IVSPER-3 | 25432 | 10761570-10787001 | U19 | 10771340-10773322 + strand | no | 660 | ADI40483.1 | unknown [Hyposoter didymator] | 660 | 0,00E+00 | 658/660(99%) | |
| scaffold127548 | IVSPER-3 | 25432 | 10761570-10787001 | IVSP4-2 | 10774112-10775407 - strand | no | 431 | ADI40484.1 | unknown [Hyposoter didymator] | 431 | 0,00E+00 | 430/431(99%) | |
| scaffold127548 | IVSPER-3 | 25432 | 10761570-10787001 | U20 | 10776029-10776240 + strand | no | 70 | ADI40485.1 | unknown [Hyposoter didymator] | 70 | 6,00E-44 | 69/70(99%) | |
| scaffold127548 | IVSPER-3 | 25432 | 10761570-10787001 | U21 | 10776822-10777061 + strand | no | 79 | ADI40486.1 | unknown [Hyposoter didymator] | 79 | 4,00E-47 | 77/79(97%) | |
| scaffold127548 | IVSPER-3 | 25432 | 10761570-10787001 | U22 | 10777309-10778172 - strand | no | 287 | ADI40487.1 | unknown [Hyposoter didymator] | 257 | 0,00E+00 | 255/257(99%) | N-term longuer in genomic seq |
| scaffold127548 | IVSPER-3 | 25432 | 10761570-10787001 | U28 | 10778369-10779088 + strand | no | 239 | | No significant similarity found | | | | newly identified, putative (no reads in calyx_1 transcriptome dataset) |
| scaffold127548 | IVSPER-3 | 25432 | 10761570-10787001 | U23 | 10779153-10780400 - strand | no | 415 | ADI40488.1 | unknown [Hyposoter didymator] | 415 | 0,00E+00 | 408/415(98%) | |
| scaffold127548 | IVSPER-3 | 25432 | 10761570-10787001 | p53-1 | 10780635-10781822 + strand | no | 395 | CAR31590.1 | p53-like1 protein [Hyposoter didymator] | 395 | 0,00E+00 | 391/395(99%) | |
| scaffold127548 | IVSPER-3 | 25432 | 10761570-10787001 | U24 | 10782094-10783785 - strand | no | 563 | ADI40490.1 | unknown [Hyposoter didymator] | 563 | 0,00E+00 | 561/563(99%) | |
| scaffold127548 | IVSPER-3 | 25432 | 10761570-10787001 | N-2 | 10785523-10787001 + strand | no | 492 | ADI40491.1 | unknown [Hyposoter didymator] | 492 | 0,00E+00 | 490/492(99%) | |
| scaffold91 | IVSPER-2 | 26611 | 541304-567914 | N-4 | 541304-542734 - strand | no | 476 | ADI40491.1 | unknown [Hyposoter didymator] gene="N-2" | 492 | 0,00E+00 | 277/491(56%) | |
| scaffold91 | IVSPER-2 | 26611 | 541304-567914 | U26 | 542904-543740 - strand | no | 278 | AKD28081.1 | hypothetical protein [Glypta fumiferanae] Gf_U39 | 298 | 9,00E-30 | 59/139(42%) | |
| scaffold91 | IVSPER-2 | 26611 | 541304-567914 | IVSP2-2 | 544341-545867 - strand | no | 508 | ADI40476.1 | unknown, partial [Hyposoter didymator] | 143 | 5,00E-96 | 143/143(100%) | completed IVSP2-2 sequence |
| scaffold91 | IVSPER-2 | 26611 | 541304-567914 | IVSP4-1 | 546767-548107 + strand | no | 446 | ADI40475.1 | unknown [Hyposoter didymator] | 446 | 0,00E+00 | 443/446(99%) | |
| scaffold91 | IVSPER-2 | 26611 | 541304-567914 | p12-2 | 548725-549039 - strand | no | 104 | ADI40474.1 | unknown [Hyposoter didymator] | 104 | 6,00E-69 | 103/104(99%) | |
| scaffold91 | IVSPER-2 | 26611 | 541304-567914 | U27 | 549438-549740 + strand | no | 100 | | No significant similarity found | | | | |
| scaffold91 | IVSPER-2 | 26611 | 541304-567914 | U14 | 549917-550078 - strand | no | 53 | ADI40473.1 | unknown [Hyposoter didymator] | 53 | 3,00E-29 | 53/53(100%) | |
| scaffold91 | IVSPER-2 | 26611 | 541304-567914 | p12-3 | 550341-550643 - strand | no | 100 | ADI40472.1 | unknown [Hyposoter didymator] | 100 | 8,00E-68 | 100/100(100%) | |
| scaffold91 | IVSPER-2 | 26611 | 541304-567914 | U13 | 551219-551677 - strand | no | 152 | ADI40471.1 | unknown [Hyposoter didymator] | 152 | 1,00E-107 | 152/152(100%) | |
| scaffold91 | IVSPER-2 | 26611 | 541304-567914 | U12 | 551947-552558 + strand | no | 203 | ADI40470.1 | unknown [Hyposoter didymator] | 203 | 1,00E-147 | 203/203(100%) | |
| scaffold91 | IVSPER-2 | 26611 | 541304-567914 | U11 | 553154-554104 - strand | no | 316 | ADI40469.1 | unknown [Hyposoter didymator] | 316 | 0,00E+00 | 312/316(99%) | |
| scaffold91 | IVSPER-2 | 26611 | 541304-567914 | U10 | 555456-559523 - strand | no | 1355 | ADI40468.1 | unknown [Hyposoter didymator] | 1355 | 0,00E+00 | 1352/1355(99%) | |
| scaffold91 | IVSPER-2 | 26611 | 541304-567914 | U9 | 559893-560717 + strand | no | 274 | ADI40467.1 | unknown [Hyposoter didymator] | 274 | 0,00E+00 | 274/274(100%) | |
| scaffold91 | IVSPER-2 | 26611 | 541304-567914 | U8 | 561319-561552 + strand | no | 77 | ADI40466.1 | unknown [Hyposoter didymator] | 77 | 3,00E-47 | 77/77(100%) | |
| scaffold91 | IVSPER-2 | 26611 | 541304-567914 | IVSP3-1 | 562225-564117 - strand | no | 630 | ADI40465.1 | unknown [Hyposoter didymator] | 630 | 0,00E+00 | 625/630(99%) | |
| scaffold91 | IVSPER-2 | 26611 | 541304-567914 | IVSP1-2 | 564643-565380 - strand | no | 245 | ADI40464.1 | unknown [Hyposoter didymator] | 196 | 5,00E-141 | 194/196(99%) | N-term longuer |
| scaffold91 | IVSPER-2 | 26611 | 541304-567914 | U7 | 565507-566145 - strand | no | 212 | ADI40463.1 | unknown [Hyposoter didymator] | 212 | 4,00E-157 | 212/212(100%) | |
| scaffold91 | IVSPER-2 | 26611 | 541304-567914 | U6 | 566499-567548 - strand | no | 349 | ADI40462.1 | unknown [Hyposoter didymator] | 349 | 0,00E+00 | 348/349(99%) | |
| scaffold91 | IVSPER-2 | 26611 | 541304-567914 | N-3 | 567663-567944 - strand | no | 93 | ADI40461.1 | unknown [Hyposoter didymator] | 93 | 1,00E-60 | 90/93(97%) | |
| **Campoletis sonorensis** | | | | | | | | | | | | | |
| **CsIV proviral segment** | | | | | | | | | | | | | |
| scaffold_8749 | rep gene | 2269 | na | rep gene 1 | 45-674 + strand | no | 209 | AHY22033.1 | repeat element 33 [Diadegma semiclausum ic | 243 | 3,00E-64 | 101/213(47%) | |
| scaffold_8748 | rep gene | 2287 | na | rep gene 2 | TO CREATE (2285,,1713) | | there is no gene predicted in this scaffold; no possibility to click and drag feature | | | | | | |
| scaffold_7280 | CsV | 2380 | not applicable | cys_CsV, partial | 1546-nn + strand | yes | NA | YP_589077.1 | VHv1.4 protein [Campoletis sonorensis ichnov | 322 | | | |
| scaffold_11 | CsA | 6368 | 861628-867995 | HP_CsA | 861717-862004 - strand | no | 95 | | no significant similarity | | | | |
| scaffold_11 | CsA | 6368 | 861628-867995 | cys_CsA | 864776-865537 - strand | yes | 177 | AAO43443.1 | A'Hv0.8 cys-motif protein precursor [Campolet | 177 | 8,00E-127 | 173/177(98%) | |
| scaffold_49 | CsB | 6626 | 22030-28655 | rep_CsB | 23475-24182 - strand | no | 235 | AAA42923.1 | repeat element protein [Campoletis sonorensi: | 235 | 2,00E-174 | 235/235(100%) | |
| scaffold_49 | CsB | 6626 | 22030-28655 | HP_CsB | 26564-27903 + strand | yes | 220 | | no significant similarity | | | | may not be a gene |
| scaffold_28 | CsC | 7350 | 25280-32629 | overlap with Cs_IVSPER-2 | | | | | | | | | |
| scaffold_17 | CsE | 7990 | 1330025-1338014 | rep1_CsE | 1331076-1331756 - strand | no | 226 | YP_00103133 | repeat element protein-d11.1 [Hyposoter fugiti | 199 | 4,00E-06 | 42/175(24%) | |
| scaffold_17 | CsE | 7990 | 1330025-1338014 | rep2_CsE | 1333207-1333722 - strand | no | 171 | YP_00103133 | repeat element protein-d11.1 [Hyposoter fugiti | 199 | 0.004 | 33/133(25%) | |
| scaffold_17 | CsE | 7990 | 1330025-1338014 | rep3_CsE | 1336256-1336924 - strand | no | 222 | AAA42923.1 | repeat element protein [Campoletis sonorensi: | 235 | 8,00E-53 | 87/197(44%) | |
| scaffold_131 | CsF | 8155 | 808380-816534 | cys_CsF | 812667-814550 - strand | yes | 403 | AY197491 | FHv1.4 cys-motif protein precursor [Campoleti | 403 | 0.0 | 403/403 (100%) | |
| scaffold_10 | CsD | 8168 | 961052-969219 | vnx_CsD | 963294-964382 - strand | no | 362 | AAO45828.1 | innexin Vnx-d1 [Campoletis sonorensis ichnov | 362 | 0.0 | 361/362(99%) | |
| scaffold_10 | CsD | 8168 | 961052-969219 | HP_CsD | 964936-965238 - strand | no | 100 | | no significant similarity | | | | |
| scaffold_14 | CsG2 | 8338 | 192247-200584 | rep1_CsG2 | 192863-193606 + strand | no | 247 | BAF45598.1 | c7-1.1 [Tranosema rostrale ichnovirus] | 248 | 5,00E-48 | 98/237(41%) | |
| scaffold_14 | CsG2 | 8338 | 192247-200584 | rep2_CsG2 | 194788-195492 + strand | no | 234 | BAF45598.1 | c7-1.1 [Tranosema rostrale ichnovirus] | 248 | 2,00E-46 | 92/203(45%) | |
| scaffold_14 | CsG2 | 8338 | 192247-200584 | rep3_CsG2 | 196511-197119 + strand | no | 202 | BAF45598.1 | c7-1.1 [Tranosema rostrale ichnovirus] | 248 | 1,00E-59 | 95/215(44%) | |
| scaffold_14 | CsG2 | 8338 | 192247-200584 | rep4_CsG2 | 198753-199601 + strand | no | 282 | BAF45598.1 | c7-1.1 [Tranosema rostrale ichnovirus] | 248 | 1,00E-59 | 107/214(50%) | |
| scaffold_14 | CsG | 8656 | 76017-84672 | vnx_CsG | 80980-82059 - strand | no | 359 | AAO45829.1 | innexin Vnx-g1 [Campoletis sonorensis ichnov | 359 | 0.0 | 359/359(100%) | |

| | | | | | | | | | | | | | |
|---|---|---|---|---|---|---|---|---|---|---|---|---|---|
| scaffold_14 | CsG | 8656 | 76017-84672 | HP_CsG | 83584-83949 + strand | no | 121 | | no significant similarity | | | | |
| scaffold_22 | CsI | 8779 | 695663-704441 | rep1_CsI | 696515-697228 + strand | no | 237 | AAA42923.1 | repeat element protein [Campoletis sonorensi | 235 | 3,00E-54 | 91/197(46%) | |
| scaffold_22 | CsI | 8779 | 695663-704441 | rep2_CsI | 697713-698441 + strand | no | 242 | AAA42923.1 | repeat element protein [Campoletis sonorensi | 235 | 2,00E-62 | 98/199(49%) | |
| scaffold_22 | CsI | 8779 | 695663-704441 | HP_CsI | 700059-700382 - strand | no | 107 | | no significant similarity | | | | |
| scaffold_22 | CsI | 8779 | 695663-704441 | rep3_CsI | 700879-701784 + strand | no | 301 | AAA42923.1 | repeat element protein [Campoletis sonorensi | 235 | 3,00E-51 | 96/190(51%) | |
| scaffold_128 | CsI2 | 9042 | 110016-119057 | vank1_CsI2 | 110664-111179 - strand | no | 171 | AAX56959.1 | vankyrin 3 [Campoletis sonorensis ichnovirus] | 171 | 3,00E-125 | 171/171(100%) | |
| scaffold_128 | CsI2 | 9042 | 110016-119057 | rep_CsI2 | 112670-113302 - strand | no | 210 | BAF45598.1 | c7-1.1 [Tranosema rostrale ichnovirus] | 248 | 8,00E-61 | 98/187(52%) | |
| scaffold_128 | CsI2 | 9042 | 110016-119057 | vank2_CsI2 | 114140-114607 - strand | no | 155 | AAX56957.1 | vankyrin 1 [Campoletis sonorensis ichnovirus] | 155 | 2,00E-111 | 155/155(100%) | |
| scaffold_128 | CsI2 | 9042 | 110016-119057 | HP_CsI2 | 115736-116062 - strand | no | 108 | | no significant similarity | | | | |
| scaffold_128 | CsI2 | 9042 | 110016-119057 | vank3_CsI2 | 117111-117617 - strand | no | 168 | AAX56958.1 | vankyrin 2 [Campoletis sonorensis ichnovirus] | 168 | 8,00E-121 | 168/168(100%) | |
| scaffold_38 | CsH | 9050 | 1398066-1407115 | HP_CsH | 1400102-1400407 - strand | no | 101 | | no significant similarity | | | | |
| scaffold_38 | CsH | 9050 | 1398066-1407115 | 5rep_CsH | 1402633-1405449 + strand | no | 938 | AAA42923.1 | repeat element protein [Campoletis sonorensi | 235 | 2,00E-41 | 89/215(41%) | |
| scaffold_16 | CsX6 | 9213 | 504600-513812 | 2rep_CsX6 | 506770-507660 - strand | no | 296 | AAA42923.1 | repeat element protein [Campoletis sonorensi | 235 | 8,00E-45 | 84/175(48%) | |
| scaffold_16 | CsX6 | 9213 | 504600-513812 | rep_CsX6 | 512155-512874 - strand | no | 239 | AAA42923.1 | repeat element protein [Campoletis sonorensi | 235 | 8,00E-57 | 95/203(47%) | |
| scaffold_15 | CsJ | 9484 | 2621922-2631405 | rep1_CsJ | 2623490-2624194 + strand | no | 234 | BAF45598.1 | c7-1.1 [Tranosema rostrale ichnovirus] | 248 | 3,00E-59 | 98/216(45%) | |
| scaffold_15 | CsJ | 9484 | 2621922-2631405 | rep2_CsJ | 2625029-2625739 - strand | no | 236 | BAF45598.1 | c7-1.1 [Tranosema rostrale ichnovirus] | 248 | 6,00E-50 | 91/218(42%) | |
| scaffold_15 | CsJ | 9484 | 2621922-2631405 | rep3_CsJ | 2629669-2630322 + strand | no | 217 | BAF45598.1 | c7-1.1 [Tranosema rostrale ichnovirus] | 248 | 2,00E-37 | 80/187(43%) | |
| scaffold_35 | CsX8 | 9999 | 164467-174465 | HP1_CsX8 | 165205-165672 - strand | no | 155 | no significant similarity | | | | | |
| scaffold_35 | CsX8 | 9999 | 164467-174465 | cys_CsX8 | 168692-172561 - strand | yes | 581 | AAO43445.1 | LHv2.8 cys-motif protein precursor [Campoleti | 678 | 0.0 | 409/671(61%) | |
| scaffold_35 | CsX8 | 9999 | 164467-174465 | HP2_CsX8 | 172947-173360 - strand | no | 137 | no significant similarity | | | | | |
| scaffold_4391 | CsL | 10024 | 8079-18102 | cys_CsL | 10645-14521 - strand | yes | 678 | AAO43445.1 | LHv2.8 cys-motif protein precursor [Campoleti | 678 | 0.0 | 675/678(99%) | shorter |
| scaffold_4391 | CsL | 10024 | 8079-18102 | HP_CsL | 16908-17306 + strand | no | 132 | | no significant similarity | | | | |
| scaffold_110 | CsX2 | 10806 | 248068-258873 | vank1_CsX2 | 249108-249605 + strand | no | 165 | ABH10021.1 | vankyrin 2 [Campoletis chlorideae ichnovirus] | 168 | 2,00E-105 | 144/165(87%) | |
| scaffold_110 | CsX2 | 10806 | 248068-258873 | vank2_CsX2 | 251609-252118 + strand | no | 169 | AAX56956.1 | vankyrin 4 [Campoletis sonorensis ichnovirus] | 160 | 2,00E-65 | 100/151(66%) | short |
| scaffold_110 | CsX2 | 10806 | 248068-258873 | vank3_CsX2 | 252614-253099 + strand | no | 161 | AAX56955.1 | vankyrin 3 [Campoletis sonorensis ichnovirus] | 161 | 9,00E-81 | 122/149(82%) | |
| scaffold_110 | CsX2 | 10806 | 248068-258873 | vank4_CsX2 | 254294-254672 + strand | no | 125 | AAX56955.1 | vankyrin 3 [Campoletis sonorensis ichnovirus] | 161 | 2,00E-23 | 62/147(42%) | |
| scaffold_110 | CsX2 | 10806 | 248068-258873 | rep_CsX2 | 256559-257191 + strand | no | 210 | BAF45598.1 | c7-1.1 [Tranosema rostrale ichnovirus] | 248 | 2,00E-60 | 94/189(50%) | |
| scaffold_5 | CsN | 10943 | 175169-186111 | N1_CsN | 177111-178547 - strand | no | 478 | AAS79017.1 | NHv1.2 protein [Campoletis sonorensis ichnov | 400 | 0.0 | 305/404(75%) | |
| scaffold_5 | CsN | 10943 | 175169-186111 | N2_CsN | 180542-181744 - strand | no | 400 | AAS79017.1 | NHv1.2 protein [Campoletis sonorensis ichnov | 400 | 0.0 | 400/400(100%) | |
| scaffold_5 | CsN | 10943 | 175169-186111 | HP_CsN | 185333-185656 + strand | no | 107 | | no significant similarity | | | | |
| scaffold_5218 | CsP | 12113 | 15720-27832 | HP_CsP | 19074-19379 - strand | no | 101 | | no significant similarity | | | | |
| scaffold_5218 | CsP | 12113 | 15720-27832 | vank4_CsP | 19628-20110 + strand | no | 160 | AAX56956.1 | vankyrin 4 [Campoletis sonorensis ichnovirus] | 160 | 3,00E-114 | 159/160(99%) | |
| scaffold_5218 | CsP | 12113 | 15720-27832 | vank3_CsP | 20609-21094 + strand | no | 161 | AAX56955.1 | vankyrin 3 [Campoletis sonorensis ichnovirus] | 161 | 4,00E-115 | 161/161(100%) | |
| scaffold_5218 | CsP | 12113 | 15720-27832 | vank2_CsP | 23733-24215 + strand | no | 160 | AAX56954.1 | vankyrin 2 [Campoletis sonorensis ichnovirus] | 160 | 1,00E-114 | 160/160(100%) | |
| scaffold_5218 | CsP | 12113 | 15720-27832 | vank1_CsP | 26747-27262 + strand | no | 171 | AAX56953.1 | vankyrin 1 [Campoletis sonorensis ichnovirus] | 171 | 3,00E-124 | 171/171(100%) | |
| scaffold_23 | CsM | 12197 | 1422227-1434423 | HP1_CsM | 1425348-1425740 - strand | no | 130 | | no significant similarity | | | | |
| scaffold_23 | CsM | 12197 | 1422227-1434423 | N_CsM | 1428224-1429615 + strand | no | 463 | AAS79017.1 | NHv1.2 protein [Campoletis sonorensis ichnov | 400 | 3,00E-166 | 245/407(60%) | |
| scaffold_23 | CsM | 12197 | 1422227-1434423 | HP2_CsM | 1431734-1432099 - strand | no | 121 | | no significant similarity | | | | |
| scaffold_50 | CsQ | 12543 | 290527-303069 | rep1_CsQ | 291919-292629 - strand | no | 236 | AAA42923.1 | repeat element protein [Campoletis sonorensi | 235 | 2,00E-73 | 116/231(50%) | |
| scaffold_50 | CsQ | 12543 | 290527-303069 | vinx1_CsQ | 294036-295118 - strand | no | 360 | YP_589076.1 | innexin-like protein 1 [Campoletis sonorensis i | 369 | 0.0 | 323/328(98%) | |
| scaffold_50 | CsQ | 12543 | 290527-303069 | vinx2_CsQ | 296198-297295 + strand | no | 365 | YP_589075.1 | innexin-like protein 2 [Campoletis sonorensis i | 365 | 0.0 | 365/365(100%) | |
| scaffold_50 | CsQ | 12543 | 290527-303069 | rep2_CsQ | 299186-299929 - strand | no | 247 | AAA42923.1 | repeat element protein [Campoletis sonorensi | 235 | 9,00E-48 | 85/187(45%) | |
| scaffold_50 | CsQ | 12543 | 290527-303069 | rep3_CsQ | 300417-301001 - strand | | 194 | AAA42923.1 | repeat element protein [Campoletis sonorensi | 235 | 6,00E-45 | 83/191(43%) | |
| scaffold_50 | CsQ | 12543 | 290527-303069 | rep4_CsQ | 301383-302144 - strand | no | 253 | AAA42923.1 | repeat element protein [Campoletis sonorensi | 235 | 2,00E-53 | 94/210(45%) | |
| scaffold_6070 | CsO1 | 12746 | 168701-181446 | 4rep_CsO1 | 171805-173964 + strand | no | 719 | AAA42923.1 | repeat element protein [Campoletis sonorensi | 235 | 3,00E-43 | 83/184(45%) | |
| scaffold_6070 | CsO1 | 12746 | 168701-181446 | HP1_CsO1 | 174818-175135 + strand | no | 105 | | no significant similarity | | | | |
| scaffold_6070 | CsO1 | 12746 | 168701-181446 | HP2_CsO1 | 176713-177060 + strand | no | 115 | | no significant similarity | | | | |
| scaffold_6070 | CsO1 | 12746 | 168701-181446 | 3rep_CsO1 | 177630-179456 + strand | no | 608 | AAA42923.1 | repeat element protein [Campoletis sonorensi | 235 | 5,00E-33 | 81/223(36%) | |
| scaffold_149 | CsU | 15338 | 374074-389411 | cys1_CsU | 375123-375900 - strand | yes | 180 | AAO43446.1 | UHv0.8a cys-motif protein precursor [Campole | 180 | 8,00E-131 | 180/180(100%) | Number of Matches: 5 |
| scaffold_149 | CsU | 15338 | 374074-389411 | cys2_CsU | 377416-378140 - strand | yes | 161 | AAO43447.1 | UHv0.8 cys-motif protein precursor [Campolet | 152 | 6,00E-54 | 91/151(60%) | |
| scaffold_149 | CsU | 15338 | 374074-389411 | cys3_CsU | 383382-384157 - strand | yes | 178 | AAO43446.1 | UHv0.8a cys-motif protein precursor [Campole | 180 | 1,00E-62 | 96/164(59%) | |
| scaffold_149 | CsU | 15338 | 374074-389411 | cys4_CsU | 385182-385875 - strand | yes | 152 | AAO43447.1 | UHv0.8 cys-motif protein precursor [Campolet | 152 | 2,00E-108 | 152/152(100%) | |
| scaffold_149 | CsU | 15338 | 374074-389411 | cys5_CsU | 387355-388121 - strand | yes | 175 | AAO43447.1 | UHv0.8 cys-motif protein precursor [Campolet | 152 | 2,00E-66 | 104/152(68%) | NNN within the nt sequence |
| scaffold_28 | CsW | 15807 | 614005-629811 | cys1_CsW | 615213-616060 + strand | yes | 203 | YP_589078.1 | cysteine-rich protein [Campoletis sonorensis i | 203 | 3,00E-140 | 192/203(95%) | |
| scaffold_28 | CsW | 15807 | 614005-629811 | cys2_CsW | 617470-618890 + strand | yes | 261 | YP_589079.1 | cysteine-rich protein [Campoletis sonorensis i | 198 | 8,00E-43 | 71/120(59%) | |
| scaffold_28 | CsW | 15807 | 614005-629811 | rep1_CsW | 619946-620629 - strand | no | 227 | BAF73402.1 | f3.1 [Tranosema rostrale ichnovirus] | 226 | 8,00E-73 | 107/222(48%) | |
| scaffold_28 | CsW | 15807 | 614005-629811 | cys3_CsW | 622282-623280 + strand | yes | 198 | YP_589079.1 | cysteine-rich protein [Campoletis sonorensis i | 198 | 2,00E-145 | 198/198(100%) | |
| scaffold_28 | CsW | 15807 | 614005-629811 | rep2_CsW | 624629-625342 - strand | no | 237 | YP_00103126 | repeat element protein-b15.1 [Hyposoter fugiti | 272 | 5,00E-80 | 122/237(51%) | |
| scaffold_28 | CsW | 15807 | 614005-629811 | HP_CsW | 627004-627324 - strand | no | 106 | | no significant similarity | | | | |
| scaffold_28 | CsW | 15807 | 614005-629811 | rep3_CsW | 628358-629077 - strand | no | 239 | YP_00103131 | repeat element protein-d4.2 [Hyposoter fugitiv | 248 | 6,00E-98 | 132/226(58%) | |
| scaffold_5890 | CsZ | 15871 | 134147-150017 | rep1_CsZ | 134843-135562 + strand | no | 239 | AHY22033.1 | repeat element 33 [Diadegma semiclausum ic | 243 | 5,00E-69 | 103/219(47%) | |
| scaffold_5890 | CsZ | 15871 | 134147-150017 | rep2_CsZ | 137736-138422 + strand | no | 228 | AIK25648.1 | Rep1 [Hyposoter didymator ichnovirus] | 231 | 9,00E-72 | 105/213(49%) | |
| scaffold_5890 | CsZ | 15871 | 134147-150017 | rep3_CsZ | 139610-140290 + strand | no | 226 | BAF45626.1 | f3.2 [Tranosema rostrale ichnovirus] | 237 | 3,00E-62 | 93/189(49%) | |
| scaffold_5890 | CsZ | 15871 | 134147-150017 | rep4_CsZ | 140771-141505 + strand | no | 244 | AHY21950.1 | repeat element 11 [Diadegma semiclausum ic | 225 | 3,00E-55 | 87/202(43%) | |
| scaffold_5890 | CsZ | 15871 | 134147-150017 | rep5_CsZ | 142565-143128 - strand | no | 187 | YP_00103131 | repeat element protein-d3.2 [Hyposoter fugitiv | 230 | 4,00E-51 | 87/178(49%) | |
| scaffold_5890 | CsZ | 15871 | 134147-150017 | rep6_CsZ | 144223-144912 + strand | no | 229 | BAF45611.1 | d5.2 [Tranosema rostrale ichnovirus] | 218 | 4,00E-72 | 107/206(52%) | |

| | | | | | | | | | | | | | |
|---|---|---|---|---|---|---|---|---|---|---|---|---|---|
| scaffold_5890 | CsZ | 15871 | 134147-150017 | rep7_CsZ | 146321-147118 + strand | no | 265 | BAF45598.1 | c7-1.1 [Tranosema rostrale ichnovirus] | 248 | 3,00E-106 | 153/249(61%) | |
| scaffold_5934 | CsX1 | 17335 | 19391-36725 | vank1_CsX1 | 20620-21141 + strand | no | 173 | AFH35119.1 | vankyrin 5 [Hyposoter didymator ichnovirus] | 168 | 7,00E-43 | 75/164(46%) | |
| scaffold_5934 | CsX1 | 17335 | 19391-36725 | vnx1_CsX1 | 22756-23850 + strand | no | 364 | AHY21960.1 | viral innexin 3 [Diadegma semiclausum ichnov | 357 | 7,00E-131 | 182/351(52%) | |
| scaffold_5934 | CsX1 | 17335 | 19391-36725 | rep1_CsX1 | 24562-25275 + strand | no | 237 | BAF45598.1 | c7-1.1 [Tranosema rostrale ichnovirus] | 248 | 6,00E-65 | 108/216(50%) | NNN within the nt sequence |
| scaffold_5934 | CsX1 | 17335 | 19391-36725 | vank2_CsX1 | 26300-26770 + strand | no | 156 | AFH35115.1 | vankyrin 1 [Hyposoter didymator ichnovirus] | 159 | 3,00E-52 | 82/152(54%) | Number of Matches: 4 |
| scaffold_5934 | CsX1 | 17335 | 19391-36725 | vank3_CsX1 | 28742-29212 + strand | no | 156 | YP_00103123 | vankyrin-b17 [Hyposoter fugitivus ichnovirus] | 170 | 1,00E-46 | 78/156(50%) | Number of Matches: 3 |
| scaffold_5934 | CsX1 | 17335 | 19391-36725 | vank4_CsX1 | 30120-30608 + strand | no | 162 | YP_00103123 | vankyrin-b1 [Hyposoter fugitivus ichnovirus] | 167 | 8,00E-55 | 88/162(54%) | |
| scaffold_5934 | CsX1 | 17335 | 19391-36725 | vnx2_CsX1 | 31535-32626 + strand | no | 363 | BAF45609.1 | d4.1 [Tranosema rostrale ichnovirus] | 376 | 3,00E-141 | 198/360(55%) | |
| scaffold_5934 | CsX1 | 17335 | 19391-36725 | rep2_CsX1 | 34635-35156 + strand | no | 173 | BAF45626.1 | f3.2 [Tranosema rostrale ichnovirus] | 237 | 3,00E-55 | 93/180(52%) | |
| scaffold_5934 | CsX1 | 17335 | 19391-36725 | vank5_CsX1 | 35714-36208 + strand | no | 164 | ABH10021.1 | vankyrin 2 [Campoletis chlorideae ichnovirus] | 168 | 6,00E-50 | 80/163(49%) | |
| scaffold_116 | CsT | 23217 | 7789-31005 | HP1_CsT | 14790-15116 - strand | no | 108 | | no significant similarity | | | | |
| scaffold_116 | CsT | 23217 | 7789-31005 | HP2_CsT | 21883-22209 - strand | no | 108 | | no significant similarity | | | | |
| scaffold_116 | CsT | 23217 | 7789-31005 | HP3_CsT | 27023-27376 + strand | no | 117 | | no significant similarity | | | | |
| scaffold_116 | CsT | 23217 | 7789-31005 | HP4_CsT | 28660-29043 + strand | no | 127 | | no significant similarity | | | | |
| scaffold_8362 | CsX5, partial | >5297 | 1-5297 | rep1_CsX5 | 2289-3026 - strand | no | 245 | AAA42923.1 | repeat element protein [Campoletis sonorensi | 235 | 3,00E-50 | 95/240(40%) | |
| scaffold_8362 | CsX5, partial | >5297 | 1-5297 | rep2_CsX5 | 3571-4296 + strand | no | 241 | AAA42923.1 | repeat element protein [Campoletis sonorensi | 235 | 5,00E-57 | 104/233(45%) | |
| scaffold_6095 | CsX7, partial | >6041 | 183042-189082 | rep1_CsX7 | 183042-183731 - strand | no | 229 | BAF45626.1 | f3.2 [Tranosema rostrale ichnovirus] | 237 | 2,00E-76 | 112/202(55%) | |
| scaffold_6095 | CsX7, partial | >6041 | 183042-189082 | rep2_CsX7 | 185566-186273 - strand | no | 235 | BAF45626.1 | f3.2 [Tranosema rostrale ichnovirus] | 237 | 5,00E-72 | 106/201(53%) | |
| scaffold_6095 | CsX7, partial | >6041 | 183042-189082 | rep3_CsX7 | 188342-189082 - strand | no | 246 | BAF45626.1 | f3.2 [Tranosema rostrale ichnovirus] | 237 | 2,00E-74 | 112/201(56%) | |
| scaffold_60 | CsX3, partial | >7876 | 304794-312669 | rep1_CsX3 | 304794-305507 + strand | no | 237 | AHY22033.1 | repeat element 33 [Diadegma semiclausum ic | 243 | 1,00E-77 | 115/225(51%) | |
| scaffold_60 | CsX3, partial | >7876 | 304794-312669 | rep2_CsX3 | 306761-307570 + strand | no | 269 | BAF45598.1 | c7-1.1 [Tranosema rostrale ichnovirus] | 248 | 2,00E-119 | 165/247(67%) | Number of Matches: 2, NNN in 5'region on gene (position 507,725) |
| scaffold_60 | CsX3, partial | >7876 | 304794-312669 | rep3_CsX3 | 309164-309979 + strand | no | 271 | BAF45598.1 | c7-1.1 [Tranosema rostrale ichnovirus] | 248 | 2,00E-115 | 162/247(66%) | |
| scaffold_60 | CsX3, partial | >7876 | 304794-312669 | rep4_CsX3 | 311747-312586 + strand | no | 279 | BAF45598.1 | c7-1.1 [Tranosema rostrale ichnovirus] | 248 | 8,00E-116 | 158/237(67%) | |
| scaffold_13 | CsX4, partial | >9181 | 1407384-1416564 | rep1_CsX4 | 1409287-1410012 - strand | no | 241 | AAA42923.1 | repeat element protein [Campoletis sonorensi | 235 | 5,00E-57 | 104/233(45%) | |
| scaffold_13 | CsX4, partial | >9181 | 1407384-1416564 | rep2_CsX4 | 1410557-1411294 + strand | no | 245 | AAA42923.1 | repeat element protein [Campoletis sonorensi | 235 | 3,00E-50 | 95/240(40%) | |
| scaffold_13 | CsX4, partial | >9181 | 1407384-1416564 | HP_CsX4 | 1411909-1412361 + strand | no | 150 | no significant similarity | | | | | |
| scaffold_13 | CsX4, partial | >9181 | 1407384-1416564 | rep3_CsX4 | 1415602-1416564 + strand | no | 320 | AAA42923.1 | repeat element protein [Campoletis sonorensi | 235 | 6,00E-51 | 96/232(41%) | |
| ***C. sonorensis* IVSPER** | | | | | | | | | | | | | |
| scaffold_6122 | IVSP_U36L | 471 | 234803-235273 | U36L | 234803-235273 - strand | no | 156 | AKD28080.1 | hypothetical protein [Glypta fumiferanae] Gf_L | 175 | 7,00E-15 | 49/146(34%) | |
| scaffold_12 | IVSP_U37L | 1863 | 104145-106007 | U37L-2 | 104145-106007 - strand | no | 620 | AKD28048.1 | helicase-primase domain [Glypta fumiferanae] | 721 | 0.0 | 337/621(54%) | |
| scaffold_16 | Cs_IVSPER-5 | 3750 | 2424942-2428691 | IVSP1L-3 | 2424942-2425688 + strand | no | 248 | ADI40453.1 | unknown [Hyposoter didymator] Hd_IVSP1-1 | 253 | 6,00E-57 | 104/257(40%) | |
| scaffold_16 | Cs_IVSPER-5 | 3750 | 2424942-2428691 | U2L | 2426081-2426764 + strand | no | 227 | ADI40454.1 | unknown [Hyposoter didymator] Hd_U2 | 228 | 3,00E-120 | 165/226(73%) | |
| scaffold_16 | Cs_IVSPER-5 | 3750 | 2424942-2428691 | U1L | 2427951-2428691 - strand | no | 246 | ADI40452.1 | unknown [Hyposoter didymator] Hd_U1 | 244 | 1,00E-65 | 101/245(41%) | |
| scaffold_50 | Cs_IVSPER-3 | 8610 | 218627-227236 | IVSP4L-2 | 218627-219940 - strand | no | 437 | ADI40475.1 | unknown [Hyposoter didymator] Hd_IVSP4-1 | 446 | 7,00E-156 | 217/399(54%) | |
| scaffold_50 | Cs_IVSPER-3 | 8610 | 218627-227236 | U4L | 220560-221022 + strand | no | 138 | ADI40456.1 | unknown [Hyposoter didymator] Hd_U4 | 131 | 2,00E-29 | 52/118(44%) | |
| scaffold_50 | Cs_IVSPER-3 | 8610 | 218627-227236 | p53L-3 | 221232-222284 + strand | no | 350 | CAR31590.1 | p53-like1 protein [Hyposoter didymator] | 395 | 6,00E-67 | 139/337(41%) | |
| scaffold_50 | Cs_IVSPER-3 | 8610 | 218627-227236 | U5L | 222667-223197 + strand | no | 176 | ADI40458.1 | unknown [Hyposoter didymator] Hd_U5 | 176 | 1,00E-74 | 105/174(60%) | |
| scaffold_50 | Cs_IVSPER-3 | 8610 | 218627-227236 | IVSP2L-2 | 223589-224965 + strand | no | 458 | ADI40459.1 | unknown [Hyposoter didymator] Hd_IVSP2-1 | 509 | 0.0 | 303/459(66%) | |
| scaffold_50 | Cs_IVSPER-3 | 8610 | 218627-227236 | CsN-3 | 225761-227236 + strand | no | 491 | ADI40491.1 | unknown [Hyposoter didymator] Hd_N-2 | 492 | 0.0 | 346/492(70%) | |
| scaffold_57 | Cs_IVSPER-4 | 9937 | 383305-393241 | U30L | 383305-386526 + strand | no | 1073 | AKD28060.1 | hypothetical protein [Glypta fumiferanae] Gf_L | 1322 | 0.0 | 502/1044(48%) | |
| scaffold_57 | Cs_IVSPER-4 | 9937 | 383305-393241 | U34L | 387873-390950 + strand | no | 1025 | AKD28054.1 | helicase domain [Glypta fumiferanae] Hd_U34 | 1012 | 0.0 | 486/1009(48%) | |
| scaffold_57 | Cs_IVSPER-4 | 9937 | 383305-393241 | IVSP4L-3 | 391820-393241 - strand | no | 473 | ADI40484.1 | unknown [Hyposoter didymator] Hd_IVSP4-2 | 431 | 2,00E-169 | 230/405(57%) | |
| scaffold_6122 | Cs_IVSPER-1 | 31594 | 122689-154282 | U15L | 122689-123342 - strand | no | 217 | ADI40477.1 | unknown [Hyposoter didymator] Hd_U15 | 410 | 3,00E-97 | 130/212(61%) | |
| scaffold_6122 | Cs_IVSPER-1 | 31594 | 122689-154282 | IVSP1L-1 | 124770-125432 + strand | no | 220 | ADI40464.1 | unknown [Hyposoter didymator] Hd_IVSP1-2 | 196 | 6,00E-63 | 99/197(50%) | |
| scaffold_6122 | Cs_IVSPER-1 | 31594 | 122689-154282 | U37L-1 | 126353-128176 + strand | no | 607 | AKD28048.1 | helicase-primase domain [Glypta fumiferanae] | 721 | 0.0 | 338/624(54%) | |
| scaffold_6122 | Cs_IVSPER-1 | 31594 | 122689-154282 | U31L-1 | 129117-130226 - strand | no | 369 | AKD28063.1 | hypothetical protein [Glypta fumiferanae] Gf_L | 370 | 4,00E-32 | 95/341(28%) | |
| scaffold_6122 | Cs_IVSPER-1 | 31594 | 122689-154282 | U35L | 130724-131257 - strand | no | 177 | AKD28058.1 | hypothetical protein [Glypta fumiferanae] Gf_L | 180 | 5,00E-37 | 70/156(45%) | |
| scaffold_6122 | Cs_IVSPER-1 | 31594 | 122689-154282 | Gf_U27L | 132692-134518 + strand | no | 608 | AKD28059.1 | hypothetical protein [Glypta fumiferanae] Gf_L | 325 | 1,00E-31 | 97/312(31%) | |
| scaffold_6122 | Cs_IVSPER-1 | 31594 | 122689-154282 | U17L | 135604-135929 + strand | no | 79 | ADI40480.1 | unknown [Hyposoter didymator] Hd_U17 | 82 | 1,00E-14 | 31/66(47%) | |
| scaffold_6122 | Cs_IVSPER-1 | 31594 | 122689-154282 | p12L-1 | 137295-137555 + strand | no | 86 | AAD01200.1 | p12 [Campoletis sonorensis ichnovirus] | 92 | 8,00E-17 | 43/92(47%) | |
| scaffold_6122 | Cs_IVSPER-1 | 31594 | 122689-154282 | U19L | 139263-141209 + strand | no | 648 | ADI40483.1 | unknown [Hyposoter didymator] Hd_U19 | 660 | 0.0 | 480/649(74%) | |
| scaffold_6122 | Cs_IVSPER-1 | 31594 | 122689-154282 | IVSP4L-1 | 141938-143242 - strand | no | 434 | ADI40484.1 | unknown [Hyposoter didymator] Hd_IVSP4-2 | 431 | 5,00E-175 | 232/434(53%) | |
| scaffold_6122 | Cs_IVSPER-1 | 31594 | 122689-154282 | U22L | 143868-144761 - strand | no | 297 | ADI40487.1 | unknown [Hyposoter didymator] Hd_U22 | 257 | 4,00E-59 | 95/249(38%) | |
| scaffold_6122 | Cs_IVSPER-1 | 31594 | 122689-154282 | U23L | 146427-147608 - strand | no | 393 | ADI40488.1 | unknown [Hyposoter didymator] Hd_U23 | 415 | 2,00E-170 | 245/389(63%) | |
| scaffold_6122 | Cs_IVSPER-1 | 31594 | 122689-154282 | p53L-1 | 148451-149548 + strand | no | 365 | AAD01199.1 | p53 [Campoletis sonorensis ichnovirus] | 364 | 0.0 | 306/365(84%) | |
| scaffold_6122 | Cs_IVSPER-1 | 31594 | 122689-154282 | U24L | 150683-152365 - strand | no | 560 | ADI40490.1 | unknown [Hyposoter didymator] Hd_U24 | 563 | 0.0 | 387/558(69%) | |
| scaffold_6122 | Cs_IVSPER-1 | 31594 | 122689-154282 | CsN-1 | 152861-154282 + strand | no | 473 | ADI40491.1 | unknown [Hyposoter didymator] Hd_N-2 | 492 | 0.0 | 307/491(63%) | |
| scaffold_28 | Cs_IVSPER-2 | 33269 | 7310-40578 | U6L | 10522-11574 + strand | no | 350 | ADI40462.1 | unknown [Hyposoter didymator] Hd_U6 | 349 | 0.0 | 297/350(85%) | |
| scaffold_28 | Cs_IVSPER-2 | 33269 | 7310-40578 | U7L | 13049-13684 + strand | no | 211 | ADI40463.1 | unknown [Hyposoter didymator] Hd_U7 | 212 | 5,00E-61 | 119/211(56%) | |
| scaffold_28 | Cs_IVSPER-2 | 33269 | 7310-40578 | IVSP1L-2 | 13781-14701 + strand | no | 306 | ADI40464 | unknown [Hyposoter didymator] Hd_IVSP1-2 | 196 | 9,00E-58 | 93/195(48%) | |
| scaffold_28 | Cs_IVSPER-2 | 33269 | 7310-40578 | IVSP3L | 15113-17002 + strand | no | 629 | ADI40465.1 | unknown [Hyposoter didymator] Hd_IVSP3-1 | 630 | 0.0 | 423/628(67%) | |
| scaffold_28 | Cs_IVSPER-2 | 33269 | 7310-40578 | U31L-2 | 17484-18179 - strand | no | 231 | AKD28063.1 | hypothetical protein [Glypta fumiferanae] Gf_L | 370 | 0.029 | 40/136(29%) | |
| scaffold_28 | Cs_IVSPER-2 | 33269 | 7310-40578 | U8L | 18541-18768 - strand | no | 75 | ADI40466.1 | unknown [Hyposoter didymator] Hd_U8 | 77 | 4,00E-14 | 36/75(48%) | |
| scaffold_28 | Cs_IVSPER-2 | 33269 | 7310-40578 | U9L | 19075-19890 - strand | no | 271 | ADI40467.1 | unknown [Hyposoter didymator] Hd_U9 | 274 | 2,00E-112 | 170/269(63%) | |
| scaffold_28 | Cs_IVSPER-2 | 33269 | 7310-40578 | U16L | 20258-22096 - strand | no | 612 | ADI40479.1 | unknown [Hyposoter didymator] Hd_U16 | 612 | 0.0 | 491/605(81%) | |
| scaffold_28 | Cs_IVSPER-2 | 33269 | 7310-40578 | U10L | 22789-27768 + strand | no | 1344 | ADI40468.1 | unknown [Hyposoter didymator] Hd_U10 | 1355 | 0.0 | 1035/1353(76%) | partial? |

| | | | | | | | | | | | | | |
|---|---|---|---|---|---|---|---|---|---|---|---|---|---|
| scaffold_28 | Cs_IVSPER-2 | 33269 | 7310-40578 | U11L | 27768-28709 + strand | no | 313 | ADI40469.1 | unknown [Hyposoter didymator] Hd_U11 | 316 | 3,00E-180 | 244/317(77%) | |
| scaffold_28 | Cs_IVSPER-2 | 33269 | 7310-40578 | U12L | 29360-29971 - strand | no | 203 | ADI40470.1 | unknown [Hyposoter didymator] Hd_U12 | 203 | 1,00E-121 | 168/203(83%) | |
| scaffold_28 | Cs_IVSPER-2 | 33269 | 7310-40578 | U13L | 30286-30768 + strand | no | 160 | ADI40471.1 | unknown [Hyposoter didymator] Hd_U13 | 152 | 5,00E-29 | 66/160(41%) | |
| scaffold_28 | Cs_IVSPER-2 | 33269 | 7310-40578 | p12L-2 | 31075-31353 + strand | no | 92 | AAD01200.1 | p12 [Campoletis sonorensis ichnovirus] | 92 | 5,00E-59 | 92/92(100%) | |
| scaffold_28 | Cs_IVSPER-2 | 33269 | 7310-40578 | U3L | 32951-33436 + strand | no | 161 | ADI40455.1 | unknown [Hyposoter didymator] Hd_U3 | 162 | 3,00E-67 | 93/155(60%) | frameshifts, partial? |
| scaffold_28 | Cs_IVSPER-2 | 33269 | 7310-40578 | IVSP2L-1 | 34504-36036 + strand | no | 510 | ADI40459.1 | unknown [Hyposoter didymator] Hd_IVSP2 | 509 | 0.0 | 325/510(64%) | |
| scaffold_28 | Cs_IVSPER-2 | 33269 | 7310-40578 | U26L | 36497-37327 + strand | no | 276 | AKD28081.1 | hypothetical protein [Glypta fumiferanae] Gf_U | 298 | 3,00E-29 | 66/165(40%) | |
| scaffold_28 | Cs_IVSPER-2 | 33269 | 7310-40578 | CsN-2 | 37470-38912 + strand | no | 480 | ADI40491.1 | unknown [Hyposoter didymator] Hd_N-2 | 492 | 0.0 | 323/491(66%) | |
| scaffold_28 | Cs_IVSPER-2 | 33269 | 7310-40578 | U25L | 39883-40578 - strand | no | 231 | AKD28083.1 | ring finger domain [Glypta fumiferanae] Hd_U: | 237 | 2,00E-16 | 41/128(32%) | |
| scaffold_28 | Cs_IVSPER-2 | 33269 | 7310-40578 | p53L-2 | 7310-8944 + strand | no | 544 | CAR31590.1 | p53-like1 protein [Hyposoter didymator] | 395 | 6,00E-57 | 155/247(62%) | |

repeated motifs found within NCBI seq
