## Additional file 5 for "Conserved and specific genomic features of endogenous polydnaviruses revealed by whole genome sequencing of two ichneumonid wasps"

| TE family | pValueLog | oddsRatio | qValue |
|---|---|---|---|
| RSX-incomp_MCL345_Hd-B-P240.102-Map4 | 3.0918611809 | 17.7821472342 | 0.3349924897 |
| noCat_MCL24_Hd-B-R551-Map20 | 2.8996118151 | 4.7330642255 | 0.3349924897 |
| RIX-comp_MCL591_Hd-B-R5563-Map3_reversed | 2.8512836938 | 39.1567664587 | 0.3349924897 |
| noCat_MCL6_Hd-B-R206-Map20 | 2.7428462035 | 6.23463961 | 0.3349924897 |
| RXX-TRIM_MCL42_Hd-B-G1267-Map5 | 2.7210184405 | 33.4299844597 | 0.3349924897 |
| RLX-incomp_MCL488_Hd-B-R4280-Map3_reversed | 2.6946367599 | 32.3691115766 | 0.3349924897 |
| DTX-incomp_MCL29_Hd-B-R2332-Map20 | 2.4301167503 | 23.5485060582 | 0.5279688384 |
| noCat_MCL674_Hd-B-R817-Map20 | 2.3493809132 | 21.361553524 | 0.5563552372 |
| noCat_MCL32_Hd-B-P0.268-Map20 | 2.2436635223 | 3.5674867377 | 0.630836842 |
| DMX-incomp_MCL506_Hd-B-R4459-Map4 | 2.1531036649 | 16.833546992 | 0.699389996 |
| noCat_MCL1_Hd-B-R3553-Map20 | 2.1112339162 | 5.4812696426 | 0.7001586835 |
| RXX-TRIM_MCL286_Hd-B-G414-Map4 | 2.0474614404 | 126.9993162993 | 0.7433278323 |
| DTX-incomp_MCL26_Hd-B-P123.194-Map8 | 1.9685388055 | 13.4457664706 | 0.8117307217 |
| RXX-TRIM_MCL276_Hd-B-G345-Map4 | 1.942283062 | 3.9690366006 | 0.8117307217 |
| noCat_MCL370_Hd-B-R1303-Map20 | 1.9072778818 | 12.4661076982 | 0.821209453 |
| RIX-incomp_MCL588_Hd-B-R5512-Map3 | 1.8271666032 | 2.724421063 | 0.9258411828 |
| RLX-incomp_MCL663_Hd-B-R7854-Map3_reversed | 1.6685508923 | 9.2725494301 | 1 |
| RXX-TRIM_MCL72_Hd-B-G1239-Map12 | 1.6543524573 | 5.270349224 | 1 |
| noCat_MCL62_Hd-B-G863-Map20 | 1.5630543946 | 4.8496542888 | 1 |
| RIX-incomp_MCL101_Hd-B-G2185-Map3_reversed | 1.5150174868 | 33.879581236 | 1 |
| RXX-TRIM_MCL117_Hd-B-G570-Map4 | 1.5150174868 | 33.879581236 | 1 |
| RXX-TRIM_MCL24_Hd-B-G1465-Map5 | 1.3754542738 | 24.213201794 | 1 |
| RXX-TRIM_MCL42_Hd-B-G1488-Map4 | 1.3656818056 | 23.649915417 | 1 |
| RIX-incomp_MCL374_Hd-B-R1487-Map3 | 1.2967139065 | 3.7766989824 | 1 |
| RIX-incomp_MCL402_Hd-B-R2054-Map3 | 1.2764334714 | 2.2091797975 | 1 |
| DXX-MITE_MCL158_Hd-B-P88.216-Map6 | 1.2753500619 | 5.6025026049 | 1 |
| RXX-TRIM_MCL252_Hd-B-G2042-Map5 | 1.2609500026 | 5.4979815512 | 1 |
| noCat_MCL51_Hd-B-G1882-Map20 | 1.2205488211 | 16.6762866175 | 1 |
| noCat_MCL214_Hd-B-G1561-Map20 | 1.2006604748 | 15.8928215465 | 1 |
| RIX-comp_MCL538_Hd-B-R4997-Map9 | 1.1879210812 | 15.4042477348 | 1 |
| RXX_MCL495_Hd-B-R4392-Map3 | 1.1879210812 | 15.4042477348 | 1 |
| noCat_MCL280_Hd-B-G363-Map11 | 1.1816979489 | 15.1742302343 | 1 |
| noCat_MCL39_Hd-B-G539-Map17 | 1.1462403296 | 13.9282132115 | 1 |
| DTX-incomp_MCL577_Hd-B-R5370-Map3 | 1.141547324 | 3.2421357768 | 1 |
| RIX-comp_MCL383_Hd-B-R1664-Map4 | 1.1350799278 | 13.5578466416 | 1 |
| DTX-incomp_MCL230_Hd-B-G1755-Map4 | 1.103343304 | 12.5477356701 | 1 |
| DXX-MITE_MCL108_Hd-B-G829-Map6 | 1.0279382182 | 2.4282562686 | 1 |
| noCat_MCL24_Hd-B-R2328-Map14 | 1.0277971244 | 2.8868187124 | 1 |
| DHX-comp_MCL1_Hd-B-R539-Map20_reversed | 1.0259762345 | 2.881362401 | 1 |
| DXX-MITE_MCL122_Hd-B-R2760-Map15 | 1.0250959422 | 10.3704731009 | 1 |
| DXX-MITE_MCL74_Hd-B-R215-Map14 | 1.0087334898 | 9.9587101772 | 1 |
| DTX-incomp_MCL361_Hd-B-P76.225-Map8 | 0.9705889493 | 9.0725947733 | 1 |
| noCat_MCL16_Hd-B-R1527-Map20 | 0.9475342594 | 2.6527528238 | 1 |
| RXX-TRIM_MCL188_Hd-B-G1290-Map3 | 0.9455850846 | 1.7435664586 | 1 |
| noCat_MCL13_Hd-B-G1765-Map20 | 0.9358557215 | 8.3256437109 | 1 |
| noCat_MCL6_Hd-B-R237-Map20 | 0.929967673 | 2.2284565677 | 1 |
| DXX-MITE_MCL25_Hd-B-G1524-Map9 | 0.9216897078 | 3.4561597075 | 1 |
| RYX-comp_MCL688_Hd-B-R9118-Map3 | 0.9164256318 | 7.9345346081 | 1 |
| RXX-TRIM_MCL73_Hd-B-G2038-Map8 | 0.8937733659 | 3.3204097 | 1 |
| RLX-incomp_MCL31_Hd-B-R547-Map6_reversed | 0.8868566945 | 1.6341481554 | 1 |
| noCat_MCL7_Hd-B-R2178-Map20 | 0.8511231227 | 3.1209573276 | 1 |
| DXX-MITE_MCL352_Hd-B-P313.38-Map3 | 0.8473302785 | 6.6785777778 | 1 |
| DXX-MITE_MCL32_Hd-B-R139-Map20 | 0.846925586 | 3.101831095 | 1 |
| RXX-TRIM_MCL234_Hd-B-G181-Map3 | 0.8344052643 | 6.4654600637 | 1 |
| DTX-incomp_MCL236_Hd-B-G1825-Map6 | 0.8219145777 | 6.2652977175 | 1 |

| | | | |
|---|---|---|---|
| noCat_MCL41_Hd-B-R236-Map20 | 0.8208340445 | 1.7446647698 | 1 |
| DTX-incomp_MCL25_Hd-B-G2324-Map6 | 0.8194662029 | 6.2267382269 | 1 |
| DTX-incomp_MCL153_Hd-B-G2301-Map3 | 0.8170339431 | 6.1886488699 | 1 |
| RXX-TRIM_MCL603_Hd-B-R5997-Map7 | 0.8170339431 | 6.1886488699 | 1 |
| DTX-incomp_MCL584_Hd-B-R548-Map13 | 0.8130360085 | 2.0013230859 | 1 |
| RXX-TRIM_MCL7_Hd-B-G1097-Map8 | 0.7922308672 | 2.8605725867 | 1 |
| DXX-MITE_MCL70_Hd-B-P337.14-Map3 | 0.7663758131 | 2.7513763818 | 1 |
| RLX-incomp_MCL54_Hd-B-R5062-Map20 | 0.7663758131 | 2.7513763818 | 1 |
| RSX-incomp_MCL319_Hd-B-G733-Map3 | 0.763034416 | 5.3971063058 | 1 |
| RXX-TRIM_MCL28_Hd-B-R7888-Map3 | 0.7588768696 | 5.3401248076 | 1 |
| RXX-TRIM_MCL655_Hd-B-R7621-Map3 | 0.7568159678 | 5.3120798255 | 1 |
| DTX-incomp_MCL160_Hd-B-R228-Map20 | 0.7535795485 | 1.8901977721 | 1 |
| DXX-MITE_MCL155_Hd-B-P127.190-Map3 | 0.7387823134 | 5.0720909652 | 1 |
| DYX-comp_MCL658_Hd-B-R7678-Map3_reversed | 0.7253607958 | 4.8999404933 | 1 |
| RXX-TRIM_MCL249_Hd-B-G2026-Map3 | 0.7234842518 | 4.8749826179 | 1 |
| noCat_MCL8_Hd-B-G2371-Map13 | 0.7070302453 | 4.6725752544 | 1 |
| RXX-TRIM_MCL452_Hd-B-R3514-Map4 | 0.7052487061 | 4.6510751114 | 1 |
| noCat_MCL6_Hd-B-R194-Map20 | 0.6964747246 | 4.5464170259 | 1 |
| RXX-TRIM_MCL481_Hd-B-R395-Map3 | 0.6896112481 | 4.466550903 | 1 |
| RLX-incomp_MCL23_Hd-B-R962-Map20_reversed | 0.6719703402 | 2.3778316076 | 1 |
| RIX-incomp_MCL622_Hd-B-R636-Map6 | 0.6714097285 | 4.2587377697 | 1 |
| RLX-incomp_MCL68_Hd-B-R4393-Map20 | 0.6682013519 | 4.223076421 | 1 |
| DXX-MITE_MCL357_Hd-B-P48.242-Map20 | 0.6650227558 | 4.1880116123 | 1 |
| DTX-incomp_MCL478_Hd-B-R3763-Map7 | 0.6525965454 | 4.0533816299 | 1 |
| DXX-MITE_MCL58_Hd-B-R511-Map20 | 0.6510747343 | 4.0370651819 | 1 |
| noCat_MCL131_Hd-B-R1669-Map20 | 0.6478961799 | 2.2884714862 | 1 |
| RXX-TRIM_MCL180_Hd-B-G1165-Map3 | 0.6429418281 | 2.2703511169 | 1 |
| RXX-TRIM_MCL238_Hd-B-G1844-Map7 | 0.6391408207 | 3.9118814922 | 1 |
| DTX-incomp_MCL253_Hd-B-G2043-Map6 | 0.6318915448 | 3.8374522737 | 1 |
| RXX-TRIM_MCL195_Hd-B-G1379-Map3 | 0.6262031527 | 3.7799252801 | 1 |
| RXX-TRIM_MCL411_Hd-B-R235-Map16 | 0.6247960905 | 3.7658138208 | 1 |
| DXX-MITE_MCL693_Hd-B-R9264-Map5 | 0.6234890207 | 1.6555564319 | 1 |
| DTX-incomp_MCL684_Hd-B-R902-Map10 | 0.6233949533 | 3.7518082086 | 1 |
| RXX-TRIM_MCL548_Hd-B-R507-Map4 | 0.6233949533 | 3.7518082086 | 1 |
| noCat_MCL62_Hd-B-R2380-Map20 | 0.5904100882 | 2.0838032092 | 1 |
| RXX-TRIM_MCL73_Hd-B-G113-Map3 | 0.5864027323 | 3.3974193202 | 1 |
| RXX-TRIM_MCL113_Hd-B-G390-Map6 | 0.5753530305 | 3.2972497971 | 1 |
| RIX-incomp_MCL676_Hd-B-R825-Map3 | 0.5646716699 | 3.2027818585 | 1 |
| DMX-incomp_MCL578_Hd-B-R5378-Map4_reversed | 0.5520867271 | 3.094379232 | 1 |
| noCat_MCL32_Hd-B-G30-Map20 | 0.5263988761 | 1.2730907769 | 1 |
| noCat_MCL5_Hd-B-P15.85-Map20 | 0.5174311102 | 1.471408109 | 1 |
| RXX-TRIM_MCL95_Hd-B-G194-Map3 | 0.5131384268 | 2.7779119951 | 1 |
| RXX-TRIM-chim_MCL120_Hd-B-G775-Map3 | 0.5009853554 | 1.3784244978 | 1 |
| DMX-incomp_MCL512_Hd-B-R4711-Map5 | 0.4877238088 | 1.4206701288 | 1 |
| DTX-incomp_MCL7_Hd-B-R1930-Map20 | 0.4830592436 | 1.5149809545 | 1 |
| noCat_MCL22_Hd-B-G1253-Map20 | 0.4725339059 | 2.4763142152 | 1 |
| noCat_MCL34_Hd-B-R3658-Map20 | 0.4674077527 | 2.4401506036 | 1 |
| noCat_MCL123_Hd-B-P10.263-Map20 | 0.4657190782 | 2.4283406034 | 1 |
| RXX-TRIM_MCL339_Hd-B-G968-Map4 | 0.4648784443 | 2.4225032309 | 1 |
| RXX-TRIM_MCL665_Hd-B-R789-Map3 | 0.4644963818 | 1.475399453 | 1 |
| RXX-TRIM_MCL224_Hd-B-G1719-Map3 | 0.4640402633 | 2.4166909532 | 1 |
| noCat_MCL5_Hd-B-P66.85-Map19 | 0.4615026965 | 1.6654762654 | 1 |
| RXX-TRIM_MCL184_Hd-B-G1227-Map4 | 0.45363783 | 1.1805146205 | 1 |
| RXX-TRIM_MCL28_Hd-B-R502-Map3 | 0.4454128834 | 2.289563753 | 1 |
| RXX-TRIM_MCL451_Hd-B-R341-Map5 | 0.4377664947 | 1.4189352099 | 1 |
| noCat_MCL10_Hd-B-G74-Map20 | 0.4331461681 | 2.2086567971 | 1 |

| | | | |
|---|---|---|---|
| RIX-incomp_MCL140_Hd-B-R6366-Map4 | 0.4203029107 | 1.5421197468 | 1 |
| DTX-incomp_MCL32_Hd-B-R4819-Map4 | 0.406251768 | 1.5010596817 | 1 |
| DTX-incomp_MCL256_Hd-B-G2078-Map3 | 0.4042905298 | 1.4953679096 | 1 |
| RXX-TRIM-chim_MCL348_Hd-B-P274.73-Map5 | 0.4019518834 | 1.4885931129 | 1 |
| RXX-TRIM_MCL637_Hd-B-R7246-Map4 | 0.4013326849 | 2.0089937354 | 1 |
| noCat_MCL50_Hd-B-R1740-Map15 | 0.3986796423 | 1.9930080883 | 1 |
| noCat_MCL364_Hd-B-R1253-Map14 | 0.3912479847 | 1.4577529779 | 1 |
| DMX-incomp_MCL405_Hd-B-R222-Map20 | 0.3822734279 | 1.3034327869 | 1 |
| RIX-comp_MCL544_Hd-B-R5026-Map5 | 0.3783754578 | 1.8733576764 | 1 |
| RXX-TRIM_MCL46_Hd-B-G2097-Map8 | 0.3621193776 | 1.3751644089 | 1 |
| RXX-TRIM_MCL318_Hd-B-G73-Map3 | 0.3436850473 | 1.6805912316 | 1 |
| DTX-incomp_MCL225_Hd-B-G1737-Map3 | 0.3410486502 | 1.6665185354 | 1 |
| noCat_MCL161_Hd-B-R815-Map11 | 0.3391169761 | 1.2148008391 | 1 |
| DTX-incomp_MCL127_Hd-B-P302.46-Map3 | 0.3368902142 | 1.6444801303 | 1 |
| noCat_MCL97_Hd-B-R6288-Map15 | 0.3312893466 | 1.615100795 | 1 |
| RXX-TRIM_MCL28_Hd-B-G2282-Map3 | 0.3219210779 | 1.5667245447 | 1 |
| RXX-TRIM_MCL114_Hd-B-G483-Map6 | 0.3130805068 | 1.2401068364 | 1 |
| noCat_MCL1_Hd-B-R963-Map20 | 0.299843028 | 1.4563906564 | 1 |
| noCat_MCL306_Hd-B-G583-Map20 | 0.2980761169 | 1.1996520693 | 1 |
| RSX-incomp_MCL5_Hd-B-G228-Map20 | 0.2959376017 | 1.4373933238 | 1 |
| RIX-comp_MCL653_Hd-B-R7590-Map3 | 0.2921013288 | 1.4188792931 | 1 |
| RSX-incomp_MCL82_Hd-B-G1278-Map10 | 0.2891642091 | 1.4048018915 | 1 |
| DTX-incomp_MCL259_Hd-B-G2134-Map5 | 0.2694245101 | 1.1233524744 | 1 |
| noCat_MCL1_Hd-B-R4412-Map11 | 0.2640654804 | 1.2878349252 | 1 |
| DXX-MITE_MCL14_Hd-B-P47.118-Map20 | 0.2536426545 | 1.2409239579 | 1 |
| DTX-incomp_MCL394_Hd-B-R188-Map20 | 0.2450851275 | 1.203105246 | 1 |
| RXX-TRIM-chim_MCL192_Hd-B-G1334-Map10 | 0.2364315959 | 0.9922069894 | 1 |
| RIX-incomp_MCL598_Hd-B-R5917-Map3 | 0.2355418741 | 1.0031928146 | 1 |
| DTX-incomp_MCL461_Hd-B-R3637-Map20 | 0.2353767113 | 0.9903427048 | 1 |
| noCat_MCL412_Hd-B-R2362-Map11 | 0.2346684256 | 1.1578873661 | 1 |
| RXX-TRIM_MCL71_Hd-B-G2274-Map5 | 0.2260147519 | 1.1209801583 | 1 |
| noCat_MCL466_Hd-B-R3699-Map14 | 0.2206649633 | 1.0984520824 | 1 |
| RXX-TRIM_MCL10_Hd-B-R1696-Map20 | 0.2109875594 | 1.0582066404 | 1 |
| RLX-incomp-chim_MCL11_Hd-B-R22-Map20_reversed | 0.2104350094 | 1.0559972371 | 1 |
| RXX-LARD_MCL384_Hd-B-R1675-Map3 | 0.2090615066 | 1.0503449303 | 1 |
| RXX-TRIM_MCL464_Hd-B-R3667-Map3 | 0.2065598622 | 0.959031564 | 1 |
| RXX-TRIM_MCL110_Hd-B-R5307-Map5 | 0.1987212447 | 1.0082286258 | 1 |
| RXX-TRIM_MCL377_Hd-B-R1609-Map4 | 0.1979522999 | 1.005127167 | 1 |
| RLX-incomp_MCL54_Hd-B-P318.33-Map3_reversed | 0.1919304422 | 0.9809773784 | 1 |
| RXX-TRIM_MCL79_Hd-B-G2126-Map3 | 0.1723189789 | 0.903900026 | 1 |
| noCat_MCL93_Hd-B-R2089-Map20 | 0.1674326606 | 0.8850441704 | 1 |
| RXX-TRIM_MCL564_Hd-B-R5172-Map15 | 0.1657437361 | 0.8745686984 | 1 |
| RIX-incomp_MCL678_Hd-B-R8483-Map3 | 0.1635201187 | 0.8700389473 | 1 |
| DXX-MITE_MCL117_Hd-B-R1594-Map4 | 0.1462109344 | 0.8045676088 | 1 |
| noCat_MCL5_Hd-B-G78-Map20 | 0.1453236875 | 0.8012485537 | 1 |
| DXX-MITE_MCL297_Hd-B-G48-Map3 | 0.1359950272 | 0.7747208501 | 1 |
| noCat_MCL315_Hd-B-G669-Map20 | 0.1356370048 | 0.7652186037 | 1 |
| RXX-LARD-chim_MCL594_Hd-B-R5678-Map3 | 0.1345078989 | 0.7878228382 | 1 |
| RIX-incomp_MCL15_Hd-B-G1297-Map3 | 0.1305252237 | 0.7463447871 | 1 |
| DXX-MITE_MCL96_Hd-B-G196-Map3 | 0.1263827996 | 0.7311123679 | 1 |
| DMX-incomp_MCL66_Hd-B-R3066-Map17 | 0.1244417631 | 0.7239923695 | 1 |
| DTX-incomp_MCL113_Hd-B-R5878-Map4 | 0.1220663735 | 0.759390942 | 1 |
| noCat_MCL19_Hd-B-R2562-Map17 | 0.1192660006 | 0.7528902472 | 1 |
| RXX-TRIM_MCL95_Hd-B-R6534-Map5 | 0.1147221127 | 0.7174703903 | 1 |
| RXX-TRIM_MCL28_Hd-B-G178-Map4 | 0.1112394297 | 0.7079398792 | 1 |
| noCat_MCL9_Hd-B-R3260-Map16 | 0.1085343855 | 0.6659751842 | 1 |

| | | | |
|---|---|---|---|
| RIX-incomp_MCL491_Hd-B-R4342-Map4 | 0.1044343647 | 0.6510892298 | 1 |
| noCat_MCL128_Hd-B-R1465-Map20 | 0.1000402666 | 0.6351497453 | 1 |
| RXX-TRIM_MCL667_Hd-B-R795-Map5 | 0.0975245311 | 0.6260265601 | 1 |
| DTX-incomp-chim_MCL612_Hd-B-R6025-Map5_reversed | 0.0931414881 | 0.6899358077 | 1 |
| RSX-incomp-chim_MCL208_Hd-B-G1541-Map3 | 0.0891490312 | 0.5956339155 | 1 |
| RXX-TRIM_MCL388_Hd-B-R1756-Map9 | 0.0863366673 | 0.5854104599 | 1 |
| noCat_MCL15_Hd-B-R3885-Map13 | 0.0848698051 | 0.580072218 | 1 |
| RSX-incomp_MCL217_Hd-B-G1583-Map3 | 0.0791764514 | 0.5593007656 | 1 |
| DXX-MITE_MCL358_Hd-B-P5.265-Map20 | 0.0773128808 | 0.5524791622 | 1 |
| RXX-TRIM_MCL385_Hd-B-R1706-Map13 | 0.0746684208 | 0.6892940088 | 1 |
| RXX-TRIM_MCL90_Hd-B-G1523-Map3 | 0.068366169 | 0.5195071398 | 1 |
| DTX-comp_MCL265_Hd-B-G2378-Map3_reversed | 0.0619437942 | 0.5640107097 | 1 |
| DTX-incomp-chim_MCL537_Hd-B-R4985-Map3_reversed | 0.0615340652 | 0.493981838 | 1 |
| noCat_MCL18_Hd-B-R4871-Map16 | 0.061070211 | 0.5612192237 | 1 |
| RXX-TRIM_MCL390_Hd-B-R1837-Map3 | 0.0568658509 | 0.5914017165 | 1 |
| DXX-MITE_MCL355_Hd-B-P340.11-Map3 | 0.055172682 | 0.4698285649 | 1 |
| RXX-TRIM_MCL340_Hd-B-P119.195-Map3 | 0.0545491057 | 0.5399454565 | 1 |
| RXX-TRIM_MCL21_Hd-B-R3610-Map20 | 0.0540213362 | 0.465405888 | 1 |
| RXX-TRIM_MCL333_Hd-B-G880-Map4 | 0.0523701959 | 0.4590321415 | 1 |
| RXX-TRIM_MCL313_Hd-B-G651-Map4 | 0.0513366438 | 0.4550225203 | 1 |
| RXX-TRIM_MCL389_Hd-B-R1786-Map3 | 0.047883421 | 0.4415031979 | 1 |
| noCat_MCL5_Hd-B-G50-Map20 | 0.0438426823 | 0.4254078205 | 1 |
| RXX-TRIM_MCL335_Hd-B-G889-Map4 | 0.0414978164 | 0.4159070724 | 1 |
| noCat_MCL48_Hd-B-R3271-Map12 | 0.0408626668 | 0.4920686584 | 1 |
| noCat_MCL85_Hd-B-R4856-Map16 | 0.0313926664 | 0.3731481726 | 1 |
| RXX-TRIM_MCL310_Hd-B-G615-Map4 | 0.0311198844 | 0.371942253 | 1 |
| DTX-incomp_MCL77_Hd-B-R904-Map5 | 0.0299219006 | 0.3666060922 | 1 |
| noCat_MCL509_Hd-B-R4481-Map11 | 0.0286037895 | 0.493747686 | 1 |
| RXX-TRIM_MCL626_Hd-B-R6590-Map4 | 0.0255797328 | 0.346641977 | 1 |
| RXX-TRIM_MCL128_Hd-B-P38.249-Map3 | 0.0243581546 | 0.3408201367 | 1 |
| DTX-incomp_MCL7_Hd-B-G850-Map20 | 0.0222815205 | 0.3306744379 | 1 |
| noCat_MCL140_Hd-B-R390-Map12 | 0.0186776674 | 0.5517706473 | 1 |
| RIX-incomp_MCL489_Hd-B-R4307-Map5 | 0.014071373 | 0.3702452214 | 1 |
| RXX-TRIM_MCL386_Hd-B-R1716-Map20 | 0.0113445143 | 0.352553533 | 1 |
| DMX-incomp_MCL545_Hd-B-R5047-Map8_reversed | 0.0083618304 | 0.419379345 | 1 |
| RIX-incomp_MCL483_Hd-B-R4195-Map4 | 0.0067427716 | 0.367743497 | 1 |
| DXX-MITE_MCL70_Hd-B-G1085-Map6 | 0.0061251295 | 0.2292382524 | 1 |
| RIX-incomp_MCL593_Hd-B-R5611-Map4 | 0.0024254658 | 0.1868463071 | 1 |
| DXX-MITE_MCL112_Hd-B-G39-Map20 | 0.0023776273 | 0.1860982987 | 1 |
| DXX-MITE_MCL144_Hd-B-R9074-Map7 | 0.0019474929 | 0.178906413 | 1 |
| RXX-TRIM_MCL18_Hd-B-R1776-Map20 | 0.001677916 | 0.1738712735 | 1 |
| RXX-TRIM_MCL444_Hd-B-R3261-Map12 | 0.0008579917 | 0.1541730841 | 1 |
| noCat_MCL524_Hd-B-R4846-Map15 | 0.0001496817 | 0.3543836046 | 1 |
| noCat_MCL459_Hd-B-R3584-Map11 | 2.6742164455795 | 0.2187300696 | 1 |
