## Additional file 6 for "Conserved and specific genomic features of endogenous polydnaviruses revealed by whole genome sequencing of two ichneumonid wasps"

| Scaffold | Proviral segment | Proviral segment size (nt) | Segment position in scaffold | DRJ name | DRJ size (nt) | DRJ position in scaffold | sequence | Identity DRJs (%) |
|---|---|---|---|---|---|---|---|---|
| **Hyposoter didymator** | | | | | | | | |
| scaffold128243 | **Hd44.1** | 3009 | 4203985-4206993 | Hd44.1_DRJ1R | 192 | 4203985-4204176 | ATGGTTGAGATTTGTAGTTCACGTTTCATCCGTTGATTGCAGCCGTGCTTCCGATGCATCAGATTGACTCAAACGTACCATCTTTCAAACCACGCATGGCTTCTAATGGTGAAGGCAGTGACCAGAACGCTGATTGTTTTGTTCCCTTTCGTAATTCCTTTGCCGTGAACTCTGCACTAGACATACAAAGTC | 73% |
| scaffold128243 | **Hd44.1** | 3009 | 4203985-4206993 | Hd44.1_DRJ1L | 192 | 4206802-4206993 | ATGGTTGAGCTCTGCACGTCAAGTTCCACGCGATGATTGCAGCCGAGCTTCCGATATATCAGATTGACTCAAACGTGTGGGTTTTCGAACCACGTAAGGCGTTCAATGGTGGAGATAGTCTCCATAACGTTGGTCCTATTGTTACTTTCTGTATTTTCCTTGCTGTGACAACTTCACGAGACTCACAGAGTC | |
| scaffold64 | **Hd42** | 3157 | 2320523-2323679 | Hd42_DRJ1R | 214 | 2323466-2323679 | GCTGCACGTCGTACGGCAGGCTTTCATGGGTCTACCGTGCCTGCTGCCAGGATCGTTGTCCAACTGTTATCTGATTCGCCTGCTGACAGAAACTCGTAGCGGCGCAGTCTTACTCCTCTTGGTAATGGCTCAAGAAAATATAAAAATAATAGCTTAGGTTGCATGCCCGCGAACGTACTGACGTGCATTAGGGTTCGCTGAACGCGTTATCAAT | 87% |
| scaffold64 | **Hd42** | 3157 | 2320523-2323679 | Hd42_DRJ1L | 223 | 2320523-2320745 | GCCGCACCTCGAACGGGGCATGCTCTCACGGGTCTACCGTGCCTGCTCTGCCAGGATCGTTGTCCAACTGTTGTCTGATTCACAGCTGCTGACAGAAACTCGTAGCAGCGCGGTCTTACTCCTCTTGGTAATGGCATGAGAAAAGATAAGAAAATAATAGCTTAGGTTCCATGCCCGCGAACGTGCCGACGTGCATCAGGGTTCGCTGTAAAGCGTTATCAAT | |
| scaffold128213 | **Hd23.2** | 3362 | 248624-251985 | Hd23.2_DRJ1R | 208 | 248624-248831 | GTGCTCCGCCGTTGATGCTGCTCCACGGTCATGTGCGTCGATGTTCGAAAGCCAGGTCAAGAAGTTTGTGAGAAATATGTGATTTTTTCTCCAACGATCACACTCTCTCCTCCGCTTGTTCCTTTGAGCGATTGCGAGCTTTATGCGGCGTCGAATAGTCACGTAGACAATGACATGCTCCGTGATGGATCCAAGACGGAACGATGCG | 74% |
| scaffold128213 | **Hd23.2** | 3362 | 248624-251985 | Hd23.2_DRJ1L | 204 | 251782-251985 | GTGCGCGCCCGTTGATGCTGCAGCGTCGTGTACGTCGCGATGCGACGGTCTGGTCAACAGGTTCGCGAGAAAGAAGTGAATTTCTCTTCAACGGGGCATCCTCATAGCTTCTCGTTCCGTTGGCCGATGGCAAGCTTTATGCAGCATCGAATAGTCACGTAGACAATGACATGCTCGATGGTTGATCCACGACGGGAAGCTGCG | |
| scaffold127348 | **Hd40** | 3494 | 926528-930021 | Hd40_DRJ1R | 175 | 926528-926702 | TCCTCGCCATTCTTCGTTCACGTTTCTGTGAACAAACAATAACAAAACGTTGAATACAGTGGTGGACGAGCTCACCTGTGTTGCGGAGATCAAGAAGGAATCTTTTTTTTTCACTTTACCTCAAGAAGGACTGATGCTTCCGCCACGGTTTCTAAAAAGCTGCCGCAGGTGGTGC | 89% |
| scaffold127348 | **Hd40** | 3494 | 926528-930021 | Hd40_DRJ1L | 176 | 929846-930021 | TCCTCGCCATTCTTCGTTCACGTTTCTGTGAACAACGAGAAGAAAACGTTGAGTACAGTGGTAAACGAGCTCACCTGTGTTGCGGAGATCAAGGGGAATCTTTTTTTTTCTCTTTACCTCAAGAAGGACTGATGCTTCTTCCGCTACGGTTTCGGAAAAGCTTCTGCAGGCGGTGC | |
| scaffold175 | **Hd38** | 3664 | 3800393-3804056 | Hd38_DRJ1R | 188 | 3803869-3804056 | AGCATGCCTCGCTGTACTGGACGTAGATATTCTAATAGATTACCGCAGCAATGGCTACGTGACAGACTCGTAAGACCAGTAGCACGGCTATAAGAGTGCAAACGATACCAAGCTCTCGGGAACCAGTCGTCGGCGAGTCATTGCACTTGCGACTTACACTGCAGAGCAGGCCAGACGCCGATGTCACT | 77% |
| scaffold175 | **Hd38** | 3664 | 3800393-3804056 | Hd38_DRJ1L | 188 | 3800393-3800580 | AGCAGGTCCCGTTGTACTGGACAACGACATTCAACAGCCTACGGCAACAGAGACTGCATGACAGACCCGTAAAGCCAGTAGCACGGCTATTGAAGTGCAAACGCCACCAAGCTCTCAATATTTATTTGCCAGCGAGTCATTGCACCAGCGACTTACCGCTGCAGAGCAGACCAGACGTCGATATCACT | |
| scaffold65 | **Hd37** | 3708 | 342293-346000 | Hd37_DRJ1R | 211 | 345790-346000 | TTGATAACGAATTACGGTAAATGCTGATGCACGTCGATGCGTTCGCGAGCATGCAACCCAAGCTATTCTTTTGTTCTTTTATCATGCCGGCACCAAGAGATAAGACCGCGCCGCTACGAGTTTCTGTCAGCAGACGAATCGGAGAATGGTGTGACAACGCTCTTGCAGCAGCAGGCACGGTAGACCCGTGAAAGCATGCCGTACAACGTGC | 82% |
| scaffold65 | **Hd37** | 3708 | 342293-346000 | Hd37_DRJ1L | 202 | 342293-342494 | TTGATAACGCATTACGTCAAACGCTAATGCACGTCGACGTGTTCACGGGCATGCAGCCTGAGCTATTCATCTTTTCTCATGCCATTACCATGAGGTGAGACCGCCGCTACGAGTTTCTGTCAGCTGAGGAACCGGAGAACGGTTGGACAACGCTCTTGACAGCAGGCACGGTAGACCCGTGAGAGCATGCCGTACAGCGTGC | |
| scaffold116 | **Hd35** | 3710 | 473341-477050 | Hd35_DRJ1R | 132 | 473341-473472 | ATCACCGACACGCTGACGTGCCCCGCAGCAGGATGGAGCGCAGCTTCACGTCGTGGATCAACCATCGAGCACGTCATTGTTTACGTGACTATTCGATGCTGCATAAATCTTGCCAACGGCCAACGGAACGAG | 91% |
| scaffold116 | **Hd35** | 3710 | 473341-477050 | Hd35_DRJ1L | 132 | 476919-477050 | ATCACCGAAACGCTGACGTGCCCCGCAGCAGGATGGAGCGCAGCTTCACGTCGTGGATCAACCATCGAGCACGTCATTGCTTACGTGACTATTCGACGTCGCATAAAGCTCGCCATCGCTCAAAGGAATGAG | |
| scaffold175 | **Hd36** | 3738 | 3796140-3799877 | Hd36_DRJ1R | 208 | 3799670-3799877 | ATTGTAAAGTTTTTCTGGTTGCCTATTGACTGTGGAGTGACATTAGGGTGTGGTCTGCTCTGCCGCCGTAAGTCGCAGGTGCAATGACTCGCTGGTGAATGAATGTTGAGAGCTTGGTGGCGTATGCACTTCAATAGTCGTGCCACTGGCCTCACGGGTCTGTCATACAGTCACCGTTGCCGTAGGCTGTTCAGTGTCTTCGTCCAGT | 75% |
| scaffold175 | **Hd36** | 3738 | 3796140-3799877 | Hd36_DRJ1L | 209 | 3796140-3796348 | ATTGTAAAATTCTCTTGTTTGCCCGATGGCTTTGGAGTGACAGCGGCGTCTGGTCTGCTCTGCAGCGGTAAGTCGCAAGTGCAATGACTCGCCGACGAATGGTTACCGAGAGCTTGGTATCGTATGAACTTTCACAGCCGTGCTGCTGGTCTTCCGAGTCTGTCACGTAGCCACTGTTGCGGTAATCTATTAAAATATCTACGTCCAGT | |
| scaffold91 | **Hd33** | 3835 | 485063-488897 | Hd33_DRJ1R | 233 | 488665-488897 | AATCAACATGTCCGACGTTCTAGCTTCCTCAAGCAGTTGCATGAAGCTTCAGTCGGTTTGCAACAACGATCGTTTGTTTCGTTTGCGCTGAAAGCACGTTCCAACCAATAATGGATGCGAGACTCGTCCACGGTGGCTCGTGGGTTTAAATTCCGCTTTGGACATGCATTCATCGTCCTGATTGACGTTCCAACTGGACAGATACGATTCCTTCAAGCTCGAAGAGTCTACTA | 81% |
| scaffold91 | **Hd33** | 3835 | 485063-488897 | Hd33_DRJ1L | 234 | 485063-485296 | AATCAACATATCCAACGTGTTCGCTTTCTCAGGCAGTTGAATGAAGCTTCAGTAGGTTTGCAACACCGATTATTTATTTCGTTTGCGATGAAAGCACGTTCAACCGAATAACGTATGCGAGACTCGTCCGCGTTAGCTTCGCGGTTCTGAATTCCTTTTAAAACCTGCATCCAACGTCGTGCTTGTCGTTGCAACTGTCCAGATACGATCCCTCCATGCTCGAAGAGTCTGCTA | |
| scaffold127548 | **Hd27** | 4002 | 13338255-13342256 | Hd27_DRJ1R | 140 | 13342117-13342256 | CCATCGCGAGCTACCGATAGCACCGCTCTGCACGTGACTATCCGAGGCTGCATCGAGATCTTGGTAACACAATAGAACGAGGTGCGCCGAGTGCGCTATCTTTGAAGAAAAAACCATGTATTCCGCTCGAGCGTATCAAC | 81% |
| scaffold127548 | **Hd27** | 4002 | 13338255-13342256 | Hd27_DRJ1L | 138 | 13338255-13338392 | CCATCGAGCTACTGATAAGACCGCGACGCACGTGACTCTTCACCGCCGCATGGAGTTCGTGGTCACACAATAGAATGAGGTGCAGCGACTGTGCTATCTTTGAAGAAAAAACCATGTATTCTACTCAGGCGTATCACC | |
| scaffold29771 | **Hd46** | 4109 | 19344-23452 | Hd46_DRJ1R | 555 | 22508-23062 | ACTTTGGCATGATGCCCGAGTTCAGCCGATAAAATCCTAACATACCATTTCTCGAAGCCAACCGCTCCCACGTATACACTACCCGGCCACAGCACGAAGCGACGATTTAGGAGGACTCGTCAACGCAACATACCGGCATTTCCCTAACTTCCGATCGATCATCTATGTACTGCATCGAAAGCAAGTTTTTTCTCGGGAACCGATGTGCTCGGTCCGAGCTTTGAAGAATTGTTTGCAGATAGCAGGGTAGTTAAGTACAGGTATGTGAAGTCTACAATCGCGGTTTTGAAGTCTACAATAGGGATTCTTTACAGTACTCGAGAAGGTCCCTTTGAGTGTGAAATCGTCAAGAGAATGCACAGAGTTCAATGATAAACGAGAGTCGTTCGGAGTCTGCATCATAACTAATGATTTTACGGCAACCTGATGACTTGTGCAATCTTGCCGGAGATGGAGTGAACTGCATATCACGTATGTACTAAACCGAATAGCAAGTGGTGATAAACAGACTATGTTCTGGGGGTTCCGAAAGACTTTGCGACTTAGCAAGGCCCG | 82% |
| scaffold29771 | **Hd46** | 4109 | 19344-23452 | Hd46_DRJ1L | 558 | 19344-19901 | ACTTTGGCACGATGCCCGTGTTGACTGGATAAATGCCTAAAATAACTTTTTTTTCAAGCCACCAGCTTCCAATGGGACACTGCTCGGTCACAGCACGAAGCTACGATGCGGGAGGACTCGTAATCGCAACGTACCGGCTTTTCCCCAACTGCCGATCGATCATCTATGTAATGCGTCGAACTCAAATTTTTCTTCTCGGGAACCAACGTGCTCGGTCCAAGCTTCGAAAAATTGTTTGCAGATATCAGGGTAGTTAAGTAAAGGTATGTAAAGTCCACAACCGCGGTTTTGAAGCCCGTCACTACGGATTCTTTTCAGTGCTTGAGAAGGTCTCTTTGAATGTGGAATCGTTAAGAGAATGCACACAGTTCAATGATGAACGAGAATGGTTCGGATTCTGCATCGTGACTAATGATTTGTCGGCAACCCCATGGCTGATTTAATCTTGCGGGAAATGGAGTGAACTGCATATCCCGTATATACTAAACCGAGTAGTGAGAGTGGTAATAAACAGACCATGTTCTGGGACTTAGTAAGACTGCGACATAGCAAGGCCCG | |
| scaffold29771 | **Hd46** | 4109 | 19344-23452 | Hd46_DRJ2R | 372 | 23081-23452 | ATACGCAATAGAACAACGTGGTGGTCTGGATCTGTTCAAGCTTGAAAACTGATGTTCTTTTCACGGCAACAAGCCTTCGTTAATTCAGATGAGCAATGTAAACGTGATCAAGCACAGGCACTGGACCACTTGACGTACTTCCAATGCGCTTCTTTAGTTGCGATAAATAAGCCAGTCACTGTGAGGTCAGGGTCATACATCCTCACACTGAAAACCATGGATGAAATGCAAAGACAATAATCACCGAACCACCTAATGTCACTGTAAAATAGTACTATATGAATGAAATAGTCGAATGCATAATTCATCACTAGTCGGCCGTCTATAATGACGATGGCCGTAACCTCAATTGACCTGTTTCACAATTCGGTA | 88% |
| scaffold29771 | **Hd46** | 4109 | 19344-23452 | Hd46_DRJ2L | 371 | 21450-21820 | ATATGCAATAGAACAACGTGGTGGCGTGGAACTGTTGAAGCTTGAAAACTGACGTTCTTTTCACGGCAACAAGCCTTTGCTAATTCAGATAAGCAAAGTGAACGTGTTCAAGCACGGGCATTGGACCACTCAACGTACTTCCAATGCGCTTCCTCAGCTGCGATAAATAGGCCAGTCACTGTGAGGTTAGGGGTATACATCCTCACACTGAAAACCATGACGAAATGCAAAGACAATAATCACCGAACCACCTAGTGTCACTGCGAAACAGTACTATATGAATCAAATAGTGAAATGTATATTTCATCATCTATCAGCCGTCTATAATGACAATGGCCGAGACCTCAATTGAACTGCTTAACAATTCAGTA | |
| scaffold127549 | **Hd31-34** | 4119 | 144445-148563 | Hd31-34_DRJ1R | 129 | 148037-148165 | GGTATCAATAAGTTGCAAGTCAGCAGGGCATCTACAAAATGCAGTTGGAGCACAGCCAACAAATACCATATTAAGTAATCACCTACTTAAGCGATAAGCACTCCGGCACGCGTGATCAGAGCGCATACT | 86% |
| scaffold127549 | **Hd31-34** | 4119 | 144445-148564 | Hd31-34_DRJ1L | 129 | 144445-144573 | GGTATCAATAAGTTACAAGTTAGGAGGGCATCTATAAAATGCAAGTCGGGGACAGCCAACAAACATCATATAGAGTAATCACCCACTTAAGCGATAAGCACTCCGGCACACGTGATCGAGGCGCATACT | |
| scaffold127549 | **Hd31-34** | 4119 | 144445-148564 | Hd31-34_DRJ2R | 396 | 148166-148563 | CGTGCTGAGCAATGAGCTCAAAATGCACGTATGGTCATGTGCATTGATAACGTACGTCGTAGTGGCCGGTCTGGAAGGCCGATTTTAGCACAATCATCACGAACTTGGAGAATAGGTCGATCGAACCGTGTTACATCCGTCAGTGCATGCGGATGCCATCAACTGCTTCAGAACCCGATGGTACACTCCATGATCTCTAACTCTCGCGGAATAATCAGTAAATCTTTGAAGTGACCAATCAACACTCTCCCGACACAATGATGCGGAGCATGCGGAGGTTTGCTTTATGACAACTTTGAGACCCCGTCGATGCCATAGAGGTATGGCTTGTCGAACGGAGACTACGAGCGCAGTGCATCGTCACAAGTTCCTGTCCTGTTTCATCAACTAGAAATG | 89% |

| | | | | | | | | |
|---|---|---|---|---|---|---|---|---|
| scaffold127549 | **Hd31-34** | 4119 | 144445-148564 | Hd31-34_DRJ2L | 394 | 144723-145116 | CGTGCCGAGCAATGAGCTTAAAATGCACGTATGGTCACGTGCATTGATAACGTACGTCGTAGTGGCCGGTCTGGAAAACCGATTTTAGCACAATTATCACGAACTCAAAGAATAGGTCGATCGAACCGCGTTACATCCGTCAGTGCATGCGGATGCCATCAACTACCCCAGAACCCGTTGGTACACTCCACGATCTCTAACTCTCGCGGAATAATCAGTAAGTCTTTAAATTGATTAATCAATACTCTTTCGACATAATGATGCGGAGCATGCGGAGGTTTCCTTTAAAACAACTTCGAGGCCTCGTCGATGCGATAGAGGTACAGCGCGTCGAACGAGGGTTACGAGCGAAGTGCACCGCCAAGTTCCTGTCCTGTTTCATCAACTGGAAATG | 89% |
| scaffold22 | **Hd39** | 4122 | 1002620-1006741 | Hd39_DRJ1R | 423 | 1002686-1003108 | AGCTGAATACGACAGAGTGAGGTGCTATCAAGAATGAACTAGAAAACATTTCCAAAGTTGCCGTGAGTTTCCGAAACCACTACGATCATGGGAAGCCTCTTTTGGCAAGCACGCTCTACAGGAGCGAGACAGAGACTTTGAGCTGCTCAGAACGTTCAGTGGCGTCCGATGATCTACGGCACTAGTCGAATCGCTTTAGTAGTTGTGCTGGCTGACAGCGTTCTCAATCAAATGGTCTTTCTGCACGTAGAACAACAAGGTGTACTGTTGAATGTTTGTAGGAAAATCTAGTTTCCTAACGCAGATAGATGCTGGGATGTCGTGCAGCGACATAGATTTGAAGTAGTTTGAGCGACACTCCGCTACACGTTATTCGTCACAATGCTACACAGTGCTGCAAGATTTTCAAGCGATGACGTTGCG | 94% |
| scaffold22 | **Hd39** | 4122 | 1002620-1006741 | Hd39_DRJ1L | 423 | 1006319-1006741 | AGCTGAATACCACACACAGAGTGAGGTGCTATCAAGAATGAACTAGAGAACATTTCCAAGTCGGCCGTGAGTTTGCGAAACCACTACGATCATGGGAAGCCTCTTTTGGAAAGCACGCTCTACAGGAGCGAGACAGAGACTTTGAGCTGCTCAGAACGTTCAGTGGCGTCCGATGATCTACGGCACTAGTCGAATCGCTTTAGTAGTTGTGCTGACAGCGTTCTCAATCAAATGGTCTTTCTGCACGTAGAACAACAAGGTGTACTGTTGAAAGTTTGTAGGAAAATCTAGTTTCCCAACGCAGATAGATGCTGGGATGCCGTGCAGCGACATAGATTTGAAGTAGTTTGAGCGACGCTCCGCCACACGTTCTTCGTCACGATGCTACACAATGCTGAAAGATTTCCAAGCGGTGACGTTGCG | |
| scaffold22 | **Hd39** | 4122 | 1002620-1006741 | Hd39_DRJ2R | 307 | 1002620-1002926 | GGCATATGTGCAAGTCAAAGGCGAATGAAGGGAATCATACAATTTGTCATTGCGGGCTGTTAACGTAGCTGAATACGACAGAGTGAGGTGCTATCAAGAATGAACTAGAAAACATTTCCAAAGTTGCCGTGAGTTTCCGAAACCACTACGATCATGGGAAGCCTCTTTTGGCAAGCACGCTCTACAGGAGCGAGACAGAGACTTTGAGCTGCTCAGAACGTTCAGTGGCGTCCGATGATCTACGGCACTAGTCGAATCGCTTTAGTAGTTGTGCTGGCTGACAGCGTTCTCAATCAAATGGTCTTTC | 84% |
| scaffold22 | **Hd39** | 4122 | 1002620-1006741 | Hd39_DRJ2L | 332 | 1005941-1006272 | GGCTTATGAGCAAGTTGAAGGCGAATAAAGAGAATCATACTGTTTGTTATCTCTGGCTGTTAACGTGGGGCCATGTCTCGTTGAGTCACTGAGCTGAATACCACGTACGGAGTGAGGTGCTATCAAGAATGAACTAGAAAACATTTCCAAGTCGGCCGTGAGTTTGCGAAACCACTACGATCATGGGAAGCCTCTTTTGGCAAGCACGCTCTACAGGAGCGAGACAGAGACTTTGAGCTGCTCGGAACGTTCAGTGGCGTCCGATGATCTACGGCACTAGTCGAATCGCTTTAGTAGTTGTGCTGACAGCGTTCTCAATCAAATGGTCTTTC | |
| scaffold1868 | **Hd43** | 4159 | 193639-197797 | Hd43_DRJ1R | 69 | 195404-195472 | GCGGAGATAAGAAAGTTAGTTTCGAAGCTCAAACAGATTCAGACCACGACGTTGTTCTATGGCATACGG | 83% |
| scaffold1868 | **Hd43** | 4159 | 193639-197797 | Hd43_DRJ1L | 69 | 197729-197797 | GCGGAGAAAAGAATGTTAGTTTCGAAGCGTAACGGTTCCAGACCACCGCGTTGTCCTTTGGCATACGGA | |
| scaffold1868 | **Hd43** | 4159 | 193639-197797 | Hd43_DRJ2R | 949 | 193639-194587 | ATGGAACAGATGTTTCAATGCACAGTCCTGCATTAGTCGGAAGTATTTGTTACGGTCGTCCATCTCGAAAGAAGTTCGTGAACGTAATGAGTCATGTTCTCCGATTCGCATCTATTTATAGATAACACACTGTATACGAGTCGCGAACGGTTCGCTATGGTTGGGATTCCCCGCGTTTTATTTCGTCGCGATTTTATTGACCCGGCTATTTTAAATATTCAGACGGTTGGTTATATATCTATCTGCGGTTTCTCACCGGGTTGAACCCGGAAGCCAATCGGGCCTTGCTAATTTCTTCAGGAATATGTTGATCATAGGTTGTTCGAAAGATATCCGAAAGGTATGAATAACATTGAAGTCTTCCCGAAGTATCAGAACATCGTTTGTTCATTACGACATAGTACTCAGTTTGTTTCAGTATATACGTGATATGCAGTTCGCCCAATCTCCGGTAAGGTTGGAGGAGCCATGAGCCAACAAATCATTAGTCATAAGCCAGACTCCGAACAACTCTCGCTCATTACTGAACTGTCTATATTCTCCTGACAATTGCACACTCGAACGAATCTCCTTACGCATTGAAATGTATCCGTAGGGTGAAGAGTAACTCATGTTGTAGCTTTGTACGTTCATATACTTATATACCCTGTTATCTGCAGACAATCCTTCGAAGCTCGGACCGAGCAGGTTTCGGAGGAGGAACACTTGTGTTCGACGCACCACGTAGTTGATCGATCAGTGGTCGGAAAGGAACTGGTACGGTGTGATTACCACCCATGTCTCGAGTCGTTGCTTTGTGCTGTGACCGAACAGCGTGTAGTAAGTTGGAGCGGTGTCTCGACAAATGGTATTTCGGGATTTTATTAACCGAGCTTGGGCATCGTCCCGCAGTTGACAGAAGCTCATGTTCGTTCTCAAATTTACGGAGTGATGTTCCTTCTTCCACCAT | 92% |
| scaffold1868 | **Hd43** | 4159 | 193639-197797 | Hd43_DRJ2L | 945 | 196556-197500 | ATGGAACAGATGTTTCAATGCACACTTCTGCATTAGTTGGAAGTATTTGTTACGGTCTGCCATCTCGAAAGAAGTTCGTGAACGTAATGAATCATGTTCTCCGATTCGCATCTGTTTATAGATAAGACACTGTATACGTGTCGCGAACGGTTCGCTATGGTTGGGATTCCCCGCGTTTTGTTTCGTCGCGATTTTATTAGCCCGGCTATTTTAAATATTCAGACGGTTGGTTATATATCTATCTGCGGTTTCTCACCGGATTGAACGCGGAAGCCAATCGGTCCATGCTAATTTGTTCAGGAATATAATGATCATAGGTTGTTCGAAAGATATCCGAAAGGTATGAATAACATTGAAGTCTTCCCGAAGTATCAGAACATCGTTGGTCCATTACGACATAGTACTCAGTTTGTTTCAGTATATAGGTGATATGCAGTTCACCCAATCTCCGGTAAAGTTGGACGAGCCATGAGCCAACAAATCATTAGTCATAAGCCAGCCTCCAAACAACTCTCGCTCATTACTGAACTGTCTATATTTTTCTGACAATTGCACACTCGAACGAATCTCCTTACGCATTGAAATGTATCCGTAGGGCGAAGAGTAACTCATGTTGTAGCTTTGTACGTTCATATACTTATCGACCTTGTTATCTGCAGACAAACCTTCGAAGCTCGGACCGAGCGGGTCGGTTTCAGAGGAGGAACACTTGTGTTCGACGCACCACGTAGTTGATCGATCAGTGTTCGGGAAGAAACCAGTACGTTGCGATTACCACACACGTCCCAAATCGTTGCTTCGTGCTGTGACCGAGCAGCGTGTAAGTTGGAGCGGTGCCTGGAAAAATGGTATTTCGGGATTCAATCAGGTGAGCTTGCGCATCGTCCCGAAGTTAGAAGCTCATGTTCGCTCTCAAATTTCCGGAGTGATGTTCCTTCTTACCAT | |
| scaffold67 | **Hd30** | 4164 | 5867182-5871345 | Hd30_DRJ1R | 176 | 5870745-5870920 | ACATGGTGTATGTTCAGACCAAGTCTCCCTCGCTACGAGGCTAGTTGGTTGGTCCATCCGTGCATTGCACGCTCGCTGCGGGTTGGCACCCAGCATCGTGCCACTACTGCGACTCCGGCACCTTACGAGCTGCAGCGATAAGACACTTCGTGCTGTGGAGCACGGTGTCAACGGTG | 89% |
| scaffold67 | **Hd30** | 4164 | 5867182-5871345 | Hd30_DRJ1L | 179 | 5867182-5867360 | ACGTGATATATGCTGAAAACAGGTCTGACTCGCTACGAGGCTAGTTGGTTGGTCCATCAGTGCATTGCACGCTCGCTACGGGTTGGCACGCAGCATCGTGCCACTGCTGCAACTCCAGCACCTTACAAACTGCAGCGATAATACACTTCGTGCTGTGGAGCACGGTGTCAACGGTGT | |
| scaffold67 | **Hd30** | 4164 | 5867182-5871345 | Hd30_DRJ2R | 98 | 5871248-5871345 | TCACAGTCGCCACTAGTGCTGAATTAAAAGATATTTTTCCTACGCGAGGCATCATTATCGCTGTGACCTTATCCGACAAGCTGTTGCACATAATAATG | 77% |
| scaffold67 | **Hd30** | 4164 | 5867182-5871345 | Hd30_DRJ2L | 107 | 5867359-5867465 | TCCCAGCTGCCGCTGGTGCTGCATATATTAAAAGATATTTTTTCTACAAGAGGCATCCTTATCGCTGTAACCTTTTTCGAAACGCTGTAAATATGCACATAGTAATC | |
| scaffold161 | **Hd25** | 4174 | 2449400-2453573 | Hd25_DRJ1R | 193 | 2453381-2453573 | ATTCACGACTCGTTGAGAACGGATCGCAGTTGTCGTAATAGGCATATGCAAGCCCAGCTATCATTGTGTCATTTGGAGGCAGACAGAATTATGTCGGAGTCATAATCCGTCTTCAGCTCTCTCGAGCTCTGTGTACCATTTTAGTGAACCGATAGGAATCGTGGAAAAGCGATGAGAATGAACAACCAGTCCA | 85% |
| scaffold161 | **Hd25** | 4174 | 2449400-2453573 | Hd25_DRJ1L | 191 | 2449400-2449590 | ATTGACAACTCGTAGAGAACGGATCTCAGTTGTCAAATAAGCATATGCAAGGAAAGCCATCATTGCGTCATTTGGGGGTAGACAGGCTTACGTCGGAGTCATAATCCGTCTTCAGCTCTCTCGAGCTGTGTAACACCATAGAGAACTGATAGGAAACGTGGAAATGCGATGGGAATGAACAGCCGAGTCCA | |
| scaffold119 | **Hd22** | 4178 | 698427-702604 | Hd22_DRJ1R | 105 | 698427-698531 | CAATGGCAGGAGCTTGACAAGGCAGATGATATCGAGTGAGTGCGTAGGTCTCTGGAACCGCGGGTCGTGAACAATATCGTGACCATTGATCGCCACAGCGACTTG | 90% |
| scaffold119 | **Hd22** | 4178 | 698427-702604 | Hd22_DRJ1L | 105 | 702500-702604 | CAATGGCAGGAGCTTGGCAGGACAGATGACATCGAGTGAATGCGTAGGAGTCTGAAACAGCGCGTCGTGAACAATATCGTGACCATTGATCGCCATAGCGACTTG | |
| scaffold91 | **Hd29** | 4356 | 572514-576869 | Hd29_DRJ1R | 458 | 576412-576869 | ATTTCGTGGCAAGCGACTTTTCATCGTAGAACGGATGGCCACCGCTGTGCGTTGTTGAACCAATGATCCAACTTTGTGTTCTTCAGGAACGGTGAACATTCGTGCAATCAACTTGACACGTCGTCATGCGGCAAGAAACTCGATAGACCATGAGGAATATCGTAACGTCATTCATCGTGCACGCATTCCTAGGACAAGTCTCTGCGCTCGAATCGTTGCGGATGCATTTGGAAGCAATGTGTCTCCAATTATGTTGGGAAAAAGTGTCCAAAAATTTATTAGATTGACCGAATGTTAATTTGAAGCTCTTGAGAAATTCACAGCTCGAGGTTAACGAGAGTGTGAAGGGATAAGCGTCACGTTCTTCATGGACGGTGCGGCTCGCACACGTCGCATGGAGCTTGCGAGTTCCAAATAAAACAACTGCGTGGGGTATTGTTTATTTCTTGAGAGCACCG | 82% |
| scaffold91 | **Hd29** | 4356 | 572514-576869 | Hd29_DRJ1L | 450 | 572514-572963 | ATTTCGTGGCAGGCTTAGTTTTGATGGAGGGACGTAGCGATCGCAGTGCGTCGCTGCATCAATGATGCAGCTTTGTGTCCTTCTCGAACGGTGAACATTCGTGCAACCAACCTGACACGTCGCCATTCGGAGAGAGGTTCGATAGACCATGAGGAATATCCAAACGTCATTCATTGTGCGCACTTCTAGGCTATTGCAATCGAATCATTGCAGATACGCATTTTGAAGCAACGCGGTTCAAATGATGTTGGGGAAAAGTGTGCAAACATTTATTAGATTGACCGAATGTTCATTTGAACCTCTTGAAAAACTCACAGCTCGAGGTTGAGAGTTTGACGGGATAGACGTGACGTTCTTCATGGAAGGTGTGGCTCGCAAACGTCGCATGCAGCTTGCGAGTTCCAAATGAAAACAACTGCGCGGGGTATTGTTTATTTCTGGAGAACACCG | |
| scaffold64 | **Hd21** | 4368 | 2353107-2357474 | Hd21_DRJ1R | 116 | 2357359-2357474 | TCCATTTTCGATCCTCGTGAGCTGCGGGGGCTGTGAGTGACGTTCCTGGAAACGTTGAGCAGCTTCTTCATCCGTGACAAAGATGGTCTCGAATTGAGGAGCTTATCATACGGCAG | 81% |
| scaffold64 | **Hd21** | 4368 | 2353107-2357474 | Hd21_DRJ1L | 114 | 2353107-2353220 | TCAATTTTTGATGCTCGTGAGCTGCGGGAGTCGTGAGCGTCGTTCCTGGTCACGTTGAGCAGCTTATTCACCCCTGACAAGGATTGCCTCGAATTGGAGGCTTTGCAACGGCAG | |
| scaffold198 | **Hd19** | 4440 | 1265454-1269893 | Hd19_DRJ1R | 105 | 1269789-1269893 | TTTTTGCTGCTTCATGTCTCAGAGCAGCTTGTCTACATGATACACGCTGATTTGTCTGTGACGGCACGATAATTCGCCTCATGCTGCACAAGCTTTGTGAGCTTT | 86% |
| scaffold198 | **Hd19** | 4440 | 1265454-1269893 | Hd19_DRJ1L | 106 | 1265454-1265559 | TTTTTGCTGGTTCATGTTTCAGAGCAGCCGGTCTATGTGATACACGCTGCTTAGCCTGGTGACGGCACGATGGTTCCCCTCATGCTGCACGAGCTCTGTGAGCTTT | |
| scaffold128213 | **Hd23.1** | 4457 | 208205-212661 | Hd23.1_DRJ1R | 137 | 208205-208341 | TCCTTTGAGCGATGGCGAGCTTTATGCGGCGTCGAATAGTCACGTAGACAATGACATGCTCCGTGGTTAATCCAAGACGGAACGATGCGTATCCGCCATATTGTGGGGCATGTCAGCGTTTCGGTGATAAGGTTGCA | 81% |
| scaffold128213 | **Hd23.1** | 4457 | 208205-212661 | Hd23.1_DRJ1L | 136 | 212526-212661 | TCCGTTGGCCGATGGCAAGCTTTATGCAGCATCGAATAGTCACGTAGACAATGACATGCTCGATGGTTGATCCACGACGGGCAGCTGCGCTCCGTCTTGCTACGGGGCACGTCAGCGTTTCGGTGATTGCGTTGCA | |
| scaffold127548 | **Hd47** | 4503 | 12134587-12139089 | Hd47_DRJ1R | 85 | 12139005-12139089 | CTGCTGGCCAGCACGTTCCACGAACAGTCTGAGATCTTGAACTGCGGAGAACGTTCAGTGGTCTCACACGATCTACGGCAGCACT | 87% |
| scaffold127548 | **Hd47** | 4503 | 12134587-12139089 | Hd47_DRJ1L | 85 | 12134587-12134671 | CTGCTGGCAAGCACGTTCTACGAACAGTCTGAAACCTTGAATAGCGTAGACCGTACAGTGTTCTCGCACGATCTACGGCAGCAGT | |

| | | | | | | | | |
|---|---|---|---|---|---|---|---|---|
| scaffold264 | **Hd28** | 4614 | 135485-140098 | Hd28_DRJ1R | 639 | 139460-140098 | GCCTTTGGAGCTAAACTTGCACGAGATTTCGGTGTCCGTGATAACTGAATGGCATTGTCAGGCGTAATATTTATGACGGCGGTGATGTTAACGTAGGTTCAGGCGTTAGACTGAGAGCCGGAAGGTATGGTAGTCGTTCGACCGTCTCCGAATCACGCTGACGAGAACTCTTCGTTCATCGCTTAAGCTCTGAGGACTTGACGCAATCAAGTTCTGGATGATAAACGCCAGACGCAAAGTTTTGTTTCCATAGATCACAAGTAAGAACAGTTCTTACGGACCTGACAGTCCAACATTACTCTCACGACACTGTAAAACCTATGGACGTACTTCGCACGTACCCGAGTCCACGAAACACTCGTGCTACATCCATTTGACGTTGGCCTTCTGACAAGGCCCACTATAGAGTCCCAGCGTGTGCCCGTGTCCCGGAAGGAGCTTTTACTCCCAACCGACTTTTTAGTGTGACCTTTGTCGTTACAGTTGCGTTACGTAACTACTACTGTGGAACTTTTCACTACCACAGTTACAACAGCTTATCGGCAACAGTTCCTTCAGTATCAAGAAGTAATTATTCAAGTGGAGAATGGCCGGGGTCGGAGTTGCATGAAAAGCACGGAAGTGTGTACAATACAGACG | 78% |
| scaffold264 | **Hd28** | 4614 | 135485-140098 | Hd28_DRJ1L | 664 | 135485-136148 | GCCTTTGCAGCTGAGCTGCATGAGGTTTTCATGTCCTCGATAACTGGATTCCATTGTCAGGCGTAAGCATTATGACGGCGGTGGTGTTAGGCGTTGCTTGGGAAGGTTGGTGAGCCACGTCGACGTAGATTCAGGCGTGAAACGGAGAGCTGAAAGGTATAGTAGTCGTTCGACCAACTCGGAATCACACCGACGAAAACTATTTGTCCATTGCTCAGGCTATGAGGACTTGAGGCAATCAAGTTCCGGACGCAAAGTTTTGTTTCCGGCCATAGATCACAAGTAAGAACAGTTCTGGCAGACTGGACAGTCCACCGATACCCTCACAAGACTGTAAAACCTCTGGACGTACTGCGCACGTACCCAAGTCCACGAAACACTCGTGCCACATCCATTTGATGTTGGCTTCTGCCAGGGCATGTCGCTATAGATTCCCAGCGTGTGCCCGTGTCCCAGAAAAAGCTTTGTTCACAACCGCGACTTTTGACTGTCACCTTTGTTGTTACAGTTGCGTTACGTAACTACTACTGTGGAACTTCTCACTGCCACAGTCACAACAGCCTATCGACAACTGTTCTTTCAGTATCAAGAAGTCATTATGCAAGTAGAGAATGGCCGGGGTTTTAAGTTGCATGAAAGCACTCAATTGTCTACAATGCAGACG | |
| scaffold82201 | **Hd51** | 4632 | 1077-5708 | Hd51_DRJ1R | 221 | 5368-5588 | GCACGTTGTACGGCATGCACTCACAGGTCTACCGTGCCTGCTGTCAAGAGCGTTGTCCAACTGTTCTCCGATTCATGTGCTGACAGAAACTCGTAGCGGCGGTCTTATCTCTTGGTGCTGGCATGATAATAGAACAAAAGAATATAGCTTAGGTTGCATGCCCGCGAGCGCATCGACGCGCATCAGCGTTTGCCGTAATGCGTTTATCAATGACGAGCCAT | 89% |
| scaffold82201 | **Hd51** | 4632 | 1077-5708 | Hd51_DRJ1L | 222 | 1077-1298 | GCACGTTATACCGCATGCTTTCACGGGTCTACCGTGCCTGCTGCTGCAAGAGCCTTGTCCAACCATTCTCCGATTCGTCTGCTGACAGAAACTCGTAGCGGCGGTCTTATCTCTTGGTGCTGGCATGATAAAAGAACAAAAGAATAGCTTAGGTTGCATATGCCCGTGAACGCAGCGACGTGCATTAGCGTTTGCCGTAATGCGTTATCAATGACGAGCCAT | |
| scaffold82201 | **Hd51** | 4632 | 1077-5708 | Hd51_DRJ2R | 122 | 5587-5708 | ATTTGTTTCTGTTTGCTTCGGTTTATAGCAACGCGAAGAACGGCCACCGTTGTTATCACTCTCGAGCGACATACACACGTGAGAGTGATGATGATTGTTGAGGCGGCCGATACAAAAGGCAG | 95% |
| scaffold82201 | **Hd51** | 4632 | 1077-5708 | Hd51_DRJ2L | 120 | 1473-1592 | ATTTGTTTCTGTTTGCTTCGGGTTAGCAACGTGAAGAACGGCCACCATTGTTATCACTCTCGAGCGACATACACACGTGAGACTGATGATGATTGTTGAGGCGGCCGATACAAAAGGCAG | |
| scaffold351 | **Hd18** | 4696 | 2681961-2686656 | Hd18_DRJ1R | 118 | 2686539-2686656 | AGCGACGAATGCTCGTCATACAGAAACGATGTCAGCAGGTCGTCCGACTGAAACGCTTGCAGAGAAGACTCAAACTCAACAGCTAATGGTTCGAGAAAAACTGATTGAGCTGGTAGAA | 81% |
| scaffold351 | **Hd18** | 4696 | 2681961-2686656 | Hd18_DRJ1L | 118 | 2681961-2682078 | AGCGACGGCGGATCGTCATTCAGAAACAATGTGAGCAGGTCGTTCGCCTGAAACACTTGCAGACAATGTTTAAACCCAACGGCTCATGTTTCACGAAAAACCAATTGAGCTGGTAGAA | |
| scaffold91 | **Hd24** | 4697 | 535698-540394 | Hd24_DRJ1R | 662 | 539733-540394 | CGTCTGCATTGTAGACAATTGAGTGCTTTCATGCAAATTAAGCCCCGGTCATTCTCTACTTGTATAATGACTTCTTGATACTGAAAGAACAGTTGTCGATAGGCTGTTGTGACTGTGGCAGTGAGAAGTTCCACAGTAGTAGTTACGTAACGCAACTGTAACAACAAAGGTGACTGTCAAAAGTCGGTTGGGAATAAAGCTTTTTCTGGGACACGGACACACGCTGGGAATCCATAGTGACATGCCCTGGCAGAAGCCAACGTCAAATGGATGTGGCACGAGTGTTTCGTGGACTCGGGTACGTGCGCAGTACGTCCATAGGGTTTCACAGTCTTGTGAGGGTAATGGTGGACTGTCAAGTCTGTCAGAACTGTTCTTACTTGTGATCTATGGCCGGAAACAAAACTTTGCGTCCGGAACTTGATTGCCTCAAGTCCTCGGAGCCTGAGCAATGGACAAATAGTTCTCGTCGGCGTGATTCCGAGGCGGTCGAACGACTACTATACCTTTCAGCTCTCCGTTTCGCGCCTGAATCTACGTCGACGTAGCTCACCAACCTTCCCAACGAACGCCTAACACCACCGCCGTCATGATGCTTACGCCTGACAATGGAATCCAGTTATCGCAGATATGAAAGCCTCGTGCAGCTCAGCTGCAAAGGC | 77% |
| scaffold91 | **Hd24** | 4697 | 535698-540394 | Hd24_DRJ1L | 628 | 535698-536325 | CGTCTGTATTGTATACACTTCCGTGCTTTTCATGCAACTGAGACCCCGGCCATTCTCTACTTGAATAATTACTTCTTAATACTGAAGAAACTTGTTGGGTCTGTGGCAGTTAAAAGTTGCGCAGCAGTAGTTACGTAACGCAACTGTAACGACAAAGGTGACAGTCAAAAGTCGGTTGGGAATAAAAGCTCTATCCGGGGCACGGGCACACGCTGGGACTCTATAGTGCGACATGCCCTGTCAGAAGCCAACGTCAAATGGATGTGGCACGAGTGTTTCGTGGACTCGGGTACGTGCGCAGTACGTCCAGAGGTTTTACAGTAACGTGAGGGTAATGTTGGACTGTCAGGTCCGTCAGAACTGTTCTTACTTGTGATCTATGGAAGCAAGACTTTGCGTCTGGCGTTTGTCGTCCAGAATTTGATTTCGTCAAGTCCTCAGAGCTTAAGCGATGGACGAAGAGTTTTCGTCAGCGTGGTTCGGAGACGGTCGAACGACTACCGTACCTTCCAGCCTCAGTCTAACGCCTGAACCTACGTTGATATCACCGCCGTCATAAATATTACGCCTGACAATGCCATTCAGTTATCACGGACACCGAAATCTCGTGCAGTTTAGCTCCAAAGGC | |
| scaffold128243 | **Hd44.2** | 4831 | 4197203-4202033 | Hd44.2_DRJ1R | 287 | 4197203-4197489 | CAGTTCGTCAGTGGGATGAACAGTGCTACCGAGCCCTGCTTTCAACTTTGCAACAGATTTGGAACCTGCTACTCACACTTCGTCTATCTTATGTATTAGCTACTTGGACAATGACGTCTATCCATCAGGACCAGTTTGTATTGGTATATTATATCTTACCAGAGGTATGCAAGTTTGTGGCAACGTCTTAAGCACCAAATGAGGATTTCTAATGTTTCGGACACTGTACGTCCCGTGCAGTTGTCACGGCTAGAAAATCATGAAAAGCAACCAGAGGACAAGCGTTC | 80% |
| scaffold128243 | **Hd44.2** | 4831 | 4197203-4202033 | Hd44.2_DRJ1L | 288 | 4201746-4202033 | CAGTTCGTCAGTGGGCTAAACAGTGCTACCGAGCCCTGCTTTCAACTTTGCAATATATTTGATACCTGCTAACTACCACTCTTCTATCACATTTATTGGGTACTTGGACGATGCCATTTATCCATCACGAACGATTTGCCATTCGCATCTTATACCTCATCGGATGTATGCGAGTTTGTGGCTACGTCTTTTGAACCGAATGTACATTTCTAATGTTTCGGACTCTGTACGTCCTGTGCAGTCGTCACGGCAAGAGAATTATGGAAAGGAAGGAGACGATCAACGTTC | |
| scaffold91 | **Hd15** | 4987 | 469105-474091 | Hd15_DRJ1R | 210 | 473882-474091 | GCTCTGTAATCTCGAATAAAACCAAAACATTCAGTTCATGCCTTGATATGAGCACCGTGAACGTTCATCAAAGACGTTCTGCCGGGTCGTGCCGATTACACTCAGTCTTTAAGATCTGGAAGACTCCCTGAATAGTAATGGGTGATAAGATGGCTGGAGTACGAATAGCAGTTGATCGAACAGCTTGTGGGAGATTACCAAATAGTCAAG | 78% |
| scaffold91 | **Hd15** | 4987 | 469105-474091 | Hd15_DRJ1L | 210 | 469105-469314 | GCTCTGTAGTTTCGAATGAAAGGAGAACGAGTTGTTCATTTTTTGTGATATCCACTGTGAACATTATCCGACAACGTTCCACCGGGTCGTGCCGATTACACTCGGTCCTTAAGCTCTGGAAGACTACCTGAATAGCAACGTATGATAAGATTGCTGGAGGACGGATACCAGTTGACCGAACAGCCCGTGGGTGATTACTAAATGGTCAAG | |
| scaffold175 | **Hd26** | 5018 | 10942034-10947051 | Hd26_DRJ1R | 592 | 10945930-10946521 | ATCATGCTTGGTCATGTCGATCATTGCTGTGATGCGGTTATTGTGGTTGATGAATCAACGATCAATGTCGAAACATGTGTGACGAAACTCTTGACAAGGCAAAGTGCTGACTGTTGCGGTATCAATAAGTTACAAGTTGGCAGGGCATCTACAAAATTTTGGTTTCTGTAGTTGATGTATGCAAGTCAACCTCAGCTAACAAACACAAGATAGGGTAATCGGCCACCAAGTGCTAAACGCCCCGACAGCCGTGATTGGAGCGCATACTGGCCGTTGATGCTATCATTTGTGTAATTCTCACACTGCCGACTTATGTTACCAGCAGATTTATCAGTTACTGCAGAAACAAAATAGAGTTTGTACCTTCCAGCTGACGCTTGTGTGCCCCGACATGCATGCAAATGAACGTACCTAGCAATGAGCATAAAATGCACGTATGGTCGGGTGCAATGATAACGTGCGTCGTAGTGGCTAGTCTTGAAGACCGATAGTAGCACAATCATCACGAACTTGAAGAATGAGAGTATCGAACCCTGTTGCATCTCTCAGCTCATATGGATGCGATCAGCTACCCCAGAACCCGATGGTTCAC | 91% |
| scaffold175 | **Hd26** | 5018 | 10942034-10947051 | Hd26_DRJ1L | 592 | 10942034-10942625 | ATCACGGTTAGGCATGTCGATTATTGCTGTATTGCGGTTATTGTGGTTGATGAATCAACGATCAATGTCGAAACATGTGTGACGAAACTCCTGACAAGGTAAAGTGCTGACTGTTGCGGTATCAATAAGTTACAAGTTGGCAGGGCATCTACAAAATTTTGGTTTCTGTAGTTGATATATGCAAGTCAACCTCAGCTAACAAACACAAGATAGGGTAATCGGCCACCAAGTGCTAAACGCCCCGACAGCCGTGATCGGAGCGCATACTGGCCGTCGATGCTATCATTTGTGCAATTCTCATACTGCCGACTTGGGTTACCAGCAGATTTATCAGTTACTGCAGAAACAAGATTGAGTTTGTACCTTCCAGCTGACGTTCGTGTGTCCCAACATGCATGCAAATGAACGTGCCGAGCAATGAGCTTTAAATGCACGTTTGGTCGGTTGCATTAATAACGTACGTCGTCGTGGCCAGTCTTGAAGACCGATCGTGGCACAATCATCACGAACTTGTAGAATAGGTGGACCAAACCGTGCTGTATCCGTCAGCGCATGCGGATGCGATCAGCTACCCCAGAACCCGATGGTACAC | |
| scaffold175 | **Hd26** | 5018 | 10942034-10947051 | Hd26_DRJ2R | 208 | 10946844-10947051 | AGCAAATCTGTAAACGCTGTCTTTGCTCAGTCGTGACGCTTGGCATTATTTATGACTCAAAAAGACCTGCAACAAGTAATGACGTCGCGATATGGAGAATGACAAAATGCCGATGAGACAAGGATCGACCCAGCTAATGCATACTTCGTTTATGCGCTAACTATGAGTGTAGCAATTTTCCTTCTCGGGACAAGAGAAAGAGTAGTGT | 80% |
| scaffold175 | **Hd26** | 5018 | 10942034-10947051 | Hd26_DRJ2L | 208 | 10942843-10943050 | AGCAAATCTGTAGAAGCTATCATTGTTTAGTCGTGTCGCCTCACATAATTCATAACTCAAAAAGACTTGCAACAAGTAATGGCGTTGCGATATGGAGAATGACAAAACGCCGTTAACACGGGGACCGATCTGACTAATACGTACTTCGTTTATGTGCTCACTATGAGTGTAGCAATTATCCGAATAGGCAAGAGAGAGGCAGTAGAGT | |
| scaffold64 | **Hd14** | 5196 | 36336-41531 | Hd14_DRJ1R | 78 | 41454-41531 | CGTGCTGCGTATCCACTGTTCCACGTCAAACGACCTTCGCAGCAAGCCTGACTAGTGCCAAGTATCTACCGCGTCAGC | 85% |
| scaffold64 | **Hd14** | 5196 | 36336-41531 | Hd14_DRJ1L | 78 | 36336-36413 | CGTGCGTCGTATCAACTGTTCCACGTAAAACGGCCGTCGTAGCAAGCCAGACCAGTGCCAAGCATCTACCTTGTCAGC | |
| scaffold128246 | **Hd49** | 5265 | 677866-683130 | Hd49_DRJ1R | 107 | 683024-683130 | CGCGCAGACAACTTAAAACCATGGTTCGTCGATAATGTCGTCGTACACGTAATGCAGGTTCAACTCGTGATACGCGAACGTCCCCAATACGAAGCCGTTCCCGTT | 83% |
| scaffold128246 | **Hd49** | 5265 | 677866-683130 | Hd49_DRJ1L | 104 | 677866-677969 | CGCGCAGAGACCTTAAAACCATCGTTCGCCGTTTATATCGTCGTACACGTAGTGCAGGTTGAGCCCGTGATAAGCGAACGTCAGCAATGCGAAGCCGCCGTTAT | |

| | | | | | | | | | | |
|---|---|---|---|---|---|---|---|---|---|---|
| scaffold128241 | **Hd13** | 5757 | 402487-408243 | Hd13_DRJ1R | 673 | 407571-408243 | TCGTAAACGGGCAGCCTGTTCAGATGTTATGGTCGACCATGGGTCCTGAGGCTTATGTACCACATCGCTTAGACACTTTGGGACTCTGGAAGTTTTTTCTCTAC AAGAAGTTGACACCCGTGTGCAAAGTACCCAAGTGCATCCGCTGAAAAGCTAACGATTCATCGAACCGTCTTACGGGAACACACGTCACCACTGACGTGTACT AGAGGTATAGGATCGGACGGCCATCGAGAGTACATGGCAAAGCGTCGCTAACGTCTCTGCGAAAGTACGCTACCATGGTGTAGTAGTGTCTATCCGCGTCAT GAAAAGTTTTCCCGACGAACCTATGAAAGTCGCCGACATACTCTCTCGTGAGCCAAAGCCACTGTCGACCCGCTCGTGTGAAAAATATGTTTTTTTCTTTAACA ACAACACCCTCGTGACACATCGTTTCATTGTACGACGATGAGCACGATGGTCATGTTGAATGGTCACGTACGCAGCGATGGTGTGACGACAACACTGAATCAC GCTTACTGTACATCGATTACGTTTCAGAAAAAATGCATCCACGCGAACTTGAAATGTAGATGTATGACAACGTCAGTTGAAGTGACCGATATGCATTTAGCTTC TTTTACCATCCCACACGGTCAGCTCAAAATATACACGTGTATCCCCTATGATA | 92% | | |
| scaffold128241 | **Hd13** | 5757 | 402487-408243 | Hd13_DRJ1L | 675 | 402487-403161 | TCGTAAGCGGACAGCCTGTTCAGATGTTATGGTCGACCATGGGTCCTGAGGCTTATGTACCACATCGCTTAGACACTTTGGGACTCTGGAAGTTTTTTCTCTAC AAGAAGTTGACACCCGTGTGCAAAGTACCCAAGTGCATCCGCTGAAAAGCTAACGATTCATCGAACCGTCTTACGGGAACACACGTCACCACTGACGTATACT AGAGGTTAAAATCGGACGGCAAGCGAAAGTGCATGGCAAAGCGTCACTAACGTCTCTGCGACAGTGCGCTACCATGGTGTAGTAGTGTCGATCCACGTCAAG CAAAGTTTTCCCGACGAACCTATGAAAATCGCCGACATACTCTCTCGTGAACCAAAGCCACTGTCGACCTGCTCGCGTGAAAAATATGTTTTTCTTGAACAAC ACCACTCTCCTCACACGTCGTTTCATTGTACGGCGATAAGCACGATGGTCAACGTCGAATGGTCACGTACGTCGCAATGGTGTGACGACAACACTGAATCACG CTTACTGTACATCGATTGCGTATCAGAAAAAATGCCTGCACCCGAGATTGAAGTGCAGATATATTACAACGTCTGCAGTTGTAGTGACCGACATGCATTTAGCT TCTTTTACAATCCCACACGGTCAGCTTAAAATATTCTCGTGTATCCCCTATGATA | | | |
| scaffold357 | **Hd50** | 5787 | 2218373-2224159 | Hd50_DRJ1R | 902 | 2223258-2224159 | CTTGCTGTGTTGATGAGGTACCATACCCGTGCCGGGGGCAAACACGTAAGCATCCAGCGTGGCGTTGAACCAGCTCTTGCAATAAGTATTCAACGGTTACGC AGGCTACTGTAAGAACTTTCAGATTTTAACATGTACGATTAGCGCATGCAATGAAGGATGACTCGACATCAGTTCTTTCAACGATTCTTTCATGAGATTGCATGG TGCAGTACTCCTATCGATCCGTTGAAAGAGCTAGTTATTAGTGTTATCTATATGATATGCCGAGTGTACTACGGCCCCCTGAACATGAAAACTCTAGTACATTCT TCCAATGGTAGAGGTACGAAGCGTTTTTCACATGAAGCATTAATACGTTCCAAACGTCGATGCAGTCTGGAGCTGAACGGTATTCGTCAGACGAGCACCAACA ATCCTCGCCATAGCATGGCATCAATAATGCATGAGCGTAAGCAGAGAGGTTGCGACTGTGCTACAACAATGTAAGTATAAGTTCTATCTTACGGAACTTGTCAA CGAGTAACAAGTCCTAATAACTCACACATCCAGAGCCAACTATGATCATTGACTCTGAGGATACTGTAAGGCTTGTCGAACTGTGCGAGAAGAATCTTGAAACC AAATGCAGGGCGGGTCCTTAGGAAATCATGAAAGCCCGCTTGTAGACTCGTAAATGGTTTTCAGAAAAAGTCTTTCATGATGAATCTGCACAATTTGAATGATG TCATTGCGGTTAAACGTAAATATCATTACGGCTATTTTTGCAATGTTTTTAACCGTAATATAATCCCATACCTACATCCATGACAAGCGGTATTGTTATAGAACTC CGCTTTTTCCTTGTTGCTTATCATGATTCAAATCAGTACTAACTCTATGATAGTGCACACATAGCGTTC | 95% | | |
| scaffold357 | **Hd50** | 5787 | 2218373-2224159 | Hd50_DRJ1L | 890 | 2218373-2219262 | CTTGCTGTGTTGATGAGGTACCATACCTGTGCCGGGGACAAACGCGTAAGCATCCAGCGTGGCGTTAAACCAGCTCTTGCAATAAGTATTCAACGGTTACGCA GGCTACTGTAAGAACTTTCAGTTTTTAACATGTACGACTAGCGCATGCAATGAAGGATGACTCGACATCCGTTCTTTCATGAGATTGCATGGTGCAGTACTCCC ATCGATCCGTTGAAAGAGCTAGTTATTAGTGTTATCTATATGATATGCAGAGTGTACTACGGCCCCCTGAACATGAAAACTCTAGTACATTCTTCCAATGGTAGA GGTACGAAGCGTTTTTCACATGAAGCAGTAATATGTTCCAAACGTCGATGCAGTCTGGAGCTGAACGGTATTCGTCAGACGAGCACCAACAATCCTTGCCATA GCATGGCATCAATAATGCATGAGCGTAAGCAGAGAGATTGCGACTGTGCTACAACAATGTGAGTATAAGTTCTATCTTATGGAACTTGTTAACGAGTAACAAGT CCTAATAACACACATATAGAGAGCCAACTATGATCATTGACTCTGAGGATACTGTAAGGCTTGTCGAACTGTGCAAGAAGAATCTTGAAACCAAATGCAGGGG GGGTCCTCAGAAAATCATGAAAGCCCGCTTGTAGACTCGTAAATGGTTTTCAGAAAAAGTCTTTCATGATGAATCTGCACAATTTGAATGATGTCATTGCGGTT AAACGTAAATACCATTACGGCTATTTTTGCAATGTTTTTAACCACAATATAATCTGATACCTACATCCATGACAAGCGGTATTGTTATAGAACTCCGCTTTTTCCT TGTTGCTTATCATGATCTAAATCAGGACTAACTCTATGATGGTACACCCATAGCGTTC | | | |
| scaffold59 | **Hd12** | 5902 | 674917-680818 | Hd12_DRJ1R/L | 411 | 680408-680818 | ATAGAACGCTTATCACCACCAATCTTTGGCGTCATACTGTTTGCGGCTTATCTGCAGTTGGTACTTATGGATCAAGTTGAGGAAAACCTTATCCAGGCGTCTCA TACCCTTGTATGGTCTGCTCCGTAACTGAACATCGCAGGTACACATACTCAGAGAGTAATCCACACTCAGACCTTACTCAAGACATATGATGGGCATATTCATA AGCATAGTTTGTGATATATAACGGAGCTTTGCATCCCATGTGCACATAACTCTAACAGAGATAAGTAGGGAGATCTCCAATGCTATAGGGCTCGAGTGGTTCCT ACAGAGCGGCGCTTTTTCTTATCGGTCAACTAATAAAGCTGTCTTAGATGTTTTTTTTGCTCGGTCGTGCTCGCACAAATCCATCCTGCTCGGCAGAGA | 84% | | |
| scaffold59 | **Hd12** | 5902 | 674917-680818 | Hd12_DRJ1R/L | 412 | 674917-675328 | ATAGTACGATAATTGCCGCCAGTCTTTGGCGTACTACTGTTTACGGCTTATCTGGAGTTGTCACTCATGAGTCAAGTTAAGAAAAACCTCATTTAGATGTCTCAT ATTCCTGTATGGTCTGCTCCGTACCTAAACGTCGCGGGTACACGTACTCAGAAAGGAATGCACACCCAGGCCTTGCTCAAGACATATGATGAGCATATTCGTA AGCATAGTTTGCGATATACAACGGAGCTTTGCATCCCATGTGCACATAACTCCATCAGAGATGAACGGGGAGATCTCCAATGCTATAAGGCCCTAGTGGTTCC TACAAAGCGACGGTTTTACGTATCGGTCAACTGCTCCAGCTGTCATAGGTGTTTGTTGTTGCTCGGTCGTGCTTGCAGTAATCCATCCTGCTCGGCAGAGA | | | |
| scaffold59 | **Hd10** | 6507 | 2500377-2506883 | Hd10_DRJ1R | 251 | 2500584-2500834 | AAAATTACGACCAAGGTGGATGCCGTGATGATGTAGTGGCATAGTGAAGGTGTAGCGCCTGCTAAGCCCAACAACTGTATACCCTGGACTACAATCCGTCAC GGCGCTTGGAGCATACCCCAGTAACAACGTGACTTATGTGGTAAAAAGCAGATTTCAGTACCAGAGCAAATACCTGTGCTATCTGTATGCACCAGATTTACAAT CTGTTAGCGTCAGTGTCTACACAATGATGTGGCATACGATAGATG | 78% | | |
| scaffold59 | **Hd10** | 6507 | 2500377-2506883 | Hd10_DRJ1int | 234 | 2503546-2503779 | AAAGTTACGACGAAGGTGGATGCCGTGATGATGTAGTGGCATAGCAAAGGTGTAGCGGCTGCTGGGACCAACTACTGTGTACCCTGTGATACAATCCGTCAC GGTGCTTCGAGTATGGTCTCGAGCACAGTCGTTCCTACTTCCATGTCAGAACAAATACCTGTGCTACCTGTGCACCAGATTTTCAATCTGGTAGCGTCAGTGT CTACACAATGGTGTGGCATACGACAGGTG | | 84% | 70% |
| scaffold59 | **Hd10** | 6507 | 2500377-2506883 | Hd10_DRJ1L | 288 | 2506596-2506883 | AAAATTACGACCAAGGTGGATGCCGTGATGATGTAGCGGCATAGTGAAGGTGTAGCACCTGCTAAGCCCAACAACTGTATACCCTGGACTACAATCCGTCAC GGCGCTTGGAGCATACCCCAGTAGCAACGTGACCTATGTGGTAAAAAACAGATTTCAGTTACCACTGGACTTGAGCGGAGTTGTTCTTACTTTCATGCCAGAG CAAATACCTGTGCTATCTGTATGCACCAGATTTACAATCTGTTAGCGTCAGAGTCGACACAATGATGTGGCATACGATAGGTG | | | |
| scaffold59 | **Hd10** | 6507 | 2500377-2506883 | Hd10_DRJ2R | 206 | 2500377-2500582 | CGTTGCTGTGCCGTAGTGGACATAGCTTTGCAGCAGCCTCGATGAGGTCATGATCATCTGCATGCGGCTCTCCAAAACTGCAGTGCTAAAAGTGGTTCGCAAA AGAATGTGACAACATACCGTTCCCTCTAGATTATATTCTGTCATTGCGAACAATAAGTGTGAACAAGAACTTAGTTAGATGTCTCAGGTTCTTTTGCGGCAAC | 87% | | |
| scaffold59 | **Hd10** | 6507 | 2500377-2506883 | Hd10_DRJ2L | 205 | 2506390-2506594 | CGCTGCGATGCTGTAGTAGCCGTAGCGTTGCAGTTGCATTGACGAGGTTCATATCGTCTGCATGCGGCTCTCCAAAACTGCAGTGCTAAAGGTGGTTCGCAA AAAAATGTGACAACATTCCGTTCCCTCTAGATTATCTTCTGTCATTGCGAACAGTAAGTGTGAACTAGAACTTATTAGATGTCTCAGGTTATTTCACGGCGAC | | | |
| scaffold144 | **Hd20** | 6864 | 1449468-1456331 | Hd20_DRJ1R | 215 | 1451905-1452119 | GAACGTGTTCCCGCATTAAAGGAGTTTCAGGGAAAAATAAATTTACTCTGAGTCGGTAATTCAACGTTTCACAATGTTGGACCGGGCGAGATGCATTCAGCTG CTTCTGCCTTGGATTCGATGCTGGAATCCTGTGACCACGTCGTCGTCAATCGGAGAACTGCATTGCTCACAGTGCTATCGACCTATTCGCTGTGGCTACCGTA AACCTCTAC | 84% | | |
| scaffold144 | **Hd20** | 6864 | 1449468-1456331 | Hd20_DRJ1L | 214 | 1456118-1456331 | GAACAGTGTTCCCCCATTGATGGGGTTTCAGGGTAATGAATGTACTCTGAGTCGGTATGTCAATGTTTCACAATGTTGGACCGGGCGTGATGCATTCAGCTAC TTCTGCATCGGATTTGATGCTGCGGCATCATGTGAACACGTCGTTAATTGTAGAACTGCATTTGCTCACAGTCCCATTGACCTATTCGCTATGGCTGCCATAAA CCTCTAC | | | |
| scaffold144 | **Hd20** | 6864 | 1449468-1456331 | Hd20_DRJ2R | 308 | 1449468-1449775 | GAATAGACCGTTAAGAATGACCGAAGCCGCAATGAATGTGCGCTTCTTGATGGTGCCCTTTCAGGGAGATTTAGAGCAATTTCTATAGTAGCTTTCCACAACTT TGACATAACATAAAATGTTTACATCGATACTTAACACGTGCCCATGCGCTCGCCAGAAAGAGTCGTATCTCCGCTGCAAGCGACAGATATGCAGGTTCGCATT GCAGTCGCAGCCGCGGGCATGACCCTAGCGAGATCCTGAACAGAAATCATACTTGGTCTGACATTCTTATACCAGCACTCCACCGAGATTGCAGCTTCCCA | 86% | | |
| scaffold144 | **Hd20** | 6864 | 1449468-1456331 | Hd20_DRJ2L | 300 | 1453262-1453561 | GAATAGGACCGTTAGGTATGACCGAAGCCGCAATGAACGTGCGCTCCTTGATGGTGCCCCTTCAGGAAGGTTTGGAGTAATTTCTATAGTAGATTTTCACAAC TTTGACATAACATAAAATGTTTACATCGATACTTAACACGTGTCCATGCGCTCGCCAGAAAGAGTCGTATCTCTGACAGATCTGCAGGTTCGCGTTGCAGTCTC AACCGTGGGCATGACCCTAGCGAGATCCTAGGTATGAAACATACTTGACCTAACATTCTTATACTAGAACGCCACCAATATTGCGGCTTCCCA | | | |
| scaffold377 | **Hd8** | 7356 | 2186417-2193772 | Hd8_DRJ1R | 719 | 2189916-2190634 | AAAGCAACAGCGATTGCATTTCAAATGAAGATAAGTACATGCAATGATAAATATCACAAGGTCACGTAACTCATAGGGTATTGGGTAGTTTTTCGAGATACGCA GCGCTTCATAAAGCACGTTCAACTTGGACGTCCAAAGCGTTGATAGCGTAACTTAGCGTTCGACTTGGAGCTGTAAACGCTGACGAGCGTTCAGACGGATGC GTCCACGGCACGTGCAGTCCAAGTCGAATGCGTTCGACAAACGCTTCCAACGCTTTCAACGTTGTGAGCGTCCAAGTTGAACGTGCCGATAAGCTGATGGAT ATAACTACGAGGATGCTGGTGATGATATCAGGAGAAGCAGAGGCAGCTCTCACGAATGACTACTTGCTGTGTTCGAGCAGCTTTTTCTATTCTATATGCAATTT CGTTGGCTCCGGTCAACACTCTAAAGTGATGTGGCCATACGGCTATGAGATGGATGCGCTTGACGACAGTGTCAAAGGTCAACGCCCTCGAGTAGCACATGG TCCAAAAATTCGTCTCATTCATTCGGACGCTAATTAAAGTGAGACAATGAATCCTGTTAATATATTCATGCACATGATCCAACGTACACTGAAAAACCTGGTGTG TTTGAACGATAGCGTAGACTTGTCCATCATGCTGTTGTTTTTCCAATCATTGCATCCGGCACTGCAGCAGGAATGCGTGCACCCTGTTGCCCCTTTTTTC | 95% | | |

| | | | | | | | | |
|---|---|---|---|---|---|---|---|---|
| scaffold377 | **Hd8** | 7356 | 2186417-2193772 | Hd8_DRJ1L | 720 | 2186417-2187136 | AAAGCAACAGCGATTGCATTCGGAATGAAAACAAGTACATGCAATGATAAATATCACAAGGTCACGTAACTCATAGGGTATTGGGTAGTTCTTCGAATTACGCATCGCTTCATAAGGCACGTCCAACTTGGACGTCCAAAGCGTTGAAAAGCATTACTTAGCGTTCGACTTGGAGCTGTAAACGCTGACGAGCGTTCAGACGGATGCGTCTACGGCACGTGTAGTCCAAGTCGAATGCGTTCGACAAACGCTTCCAACGCTTTCAACGTTGTGAGCGTCCAAGTTGAACGTGCCGATAAGCTGATGGATATAACTATGAGGATGCTGGTGATGATATCAGGAGAAGCAGAGGCAGCTCTCACGAATGACCGCTTGCTGTGTTCGAGTAGCTTTTTCTATTCTATATGCAATTTCGATGGCTCCGGTCAACACTCTGAAGTGATGTGGCCATACGGCTATGAGATGGATGCGCTTGACGACAGCGTCAGAGGTCAACGCCCTCGAGTAGCACATGGTCCAAACATTCGTCTCATTCATTCGAACGCTAATTAATGTCAGTCAATGAATCCTGTTAATATATTCATGCACGTGATCCAACGTACACTGAAAAACCTGGTGTGTTTGAACGATAGGGTAGACTTGTCCATCGTGCTGTTGCTTTACCAATCATTGCATCCGGCACTGCAGCAGGAATGCGTGCACCCTGCTGCGCTTTTTTTC | 9[illegible]% |
| scaffold377 | **Hd8** | 7356 | 2186417-2193772 | Hd8_DRJ2R | 292 | 2193481-2193772 | ATAACGTGTTATGGCGACGTTTTACCAAATCCAAGAAACATTGCTTAACTATCAGTAATTACCAGACACCAGTAACTACTCTAAAAGCAACAGCGATTGTATTCGGAATGAAAATAAGTACATGCAATGATAAATATCACAAGGTCACGTAACTCATAGGGTATTGGGTAGTTTTTCGAAATACGCACCGCTTCATAAAGCACGTTCAACTTGGACGTCCAAAGCGTTGAAAAGCGTTACTTAGCGTTCGACTTGGAGCTGTAAACGCTGACGAGCATTCAGATGGATGCGT | 91% |
| scaffold377 | **Hd8** | 7356 | 2186417-2193772 | Hd8_DRJ2L | 291 | 2189833-2190123 | ATGACATGTTATGGCGGTGTTTTATCAAATCCAAGACACATTGCTCAGTATCAGTAATTATCAGACTACCAGGAGCTACACTAAAAGCAACAGCGATTGCATTTCAAATGAAGATAAGTACATGCAATGATAAATATCACAAGGTCACGTAACTCATAGGGTATTGGGTAGTTTTTCGAGATACGCAGCGCTTCATAAAGCACGTTCAACTTGGACGTCCAAAGCGTTGATAGCGTAACTTAGCGTTCGACTTGGAGCTGTAAACGCTGACGAGCGTTCAGACGGATGCGT | |
| scaffold59 | **Hd16** | 7704 | 690205-697908 | Hd16_DRJ1R | 65 | 694965-695029 | TTGTCATGAGCGTAACTACATTTAAACTTGAAGCACTGTAAGAAGGAACGGTCTCACAGTACCTT | 85% |
| scaffold59 | **Hd16** | 7704 | 690205-697908 | Hd16_DRJ1L | 65 | 690205-690269 | TTGTCATGAGAGGAACTAAATTTAAACTTGTACCAACGTAAGAGGAAACGCTCTCACAGTACCTT | |
| scaffold59 | **Hd16** | 7704 | 690205-697908 | Hd16_DRJ2R | 172 | 697723-697908 | TAGATTATGCAAACGTGGCAAAGCGGTGTATGGCTTACGAGATGCGGCTGACCGCTGCATGTCAACGATAAGCAGATCATTCTACAAGAAGTATGTTTGTTCTAGGCGTTCTAAGTGTCGTTCTATGATAACGTGAGCTGTAGGTTTCTGATCATTCGGCCCCAGCTTGGAT | 94% |
| scaffold59 | **Hd16** | 7704 | 690205-697908 | Hd16_DRJ2L | 172 | 691583-691754 | TACATTATGCAAAAGTGGCGAAACGGTGTATGGCCTACGAGATGCGGGTGACCGCTGCATGTCAACGATAAGCAGATAGTTCTACAAGAAGTATGTTTGTTTTAGACGTTCTAAGCGTCGTTCTATGATAACGTGAGCTGTAGGTTTCTGATCATTCGGCCCCAGCTTGGAT | |
| scaffold351 | **Hd17** | 7730 | 2329273-2337002 | Hd17_DRJ1R | 70 | 2332468-2332537 | GTGTTGCGAGCAATAATGGTGGAAACACCTGTCATGGCACGCGTCCACTGCAGGCTGTACAAACGCTCTA | 77% |
| scaffold351 | **Hd17** | 7730 | 2329273-2337002 | Hd17_DRJ1L | 73 | 2336930-2337002 | GTGTTGCTTAGGAATAATGGTGGAAGCATCCGCGTACTTACACGTGTTCACTGGAGGCTGTACGAGCGCTCTA | |
| scaffold351 | **Hd17** | 7730 | 2329273-2337002 | Hd17_DRJ2R | 161 | 2329273-2329433 | CGTGGAGCGTGGAATGTTGACGAGTCATAGAATTTCGGCTGGCACGAGCACGTGCAAGTTGAAGTTGTACGCCTGCGATGACGAGAAGTTATCGACAGGAGTTGACAACTTTGGCTGCGACTTCCGAACGCCAGATGGCCTGATGGTGGCTGTCCCGTAAG | 79% |
| scaffold351 | **Hd17** | 7730 | 2329273-2337002 | Hd17_DRJ2L | 161 | 2333675-2333835 | CGTGCATCATGGCATCTTCACGAGTCATAGACATTCGGCAGGCACGAGCGAGAACAAGTTGAAGTTGTAGGCGTGCACAGACAAGGAGTTATAGACGGGCGTCGACAACTTCGGCTGTGACTTTTGAATGCCAGATTGCCTGATGGTAGCTGACCAATAAG | |
| scaffold64 | **Hd32** | 7916 | 88702-96617 | Hd32_DRJ1R | 613 | 92195-92807 | AGGCCTGACCTCAAGCCCGATAACCTACCGCTTATTTACCGAGTACCTACGAGAGTCGCTCAAAAACTGTTTATACATAAAGTTTCTACTGCATGCTTGAGGCCTATCTACAAATACTGCAATGTAGTTGAAAGTGCAATCCATCGCAATGCTTACTTTAGCACCTTATTATGTAAGAATTCTGCATTTGCGATACTGCTTGCACTTTACAGTGAGTGCATGAACGATTCTCCATGGTTAACCACCCACGGTACTTCATTTCATGTTTTCGCCATGACGTTGGAGATGCTCTGCTCGAACGCTAATCATGGTGACAATGCAATAAGCTGGGTGGAATGCTGACGTTAAGAATGAAATCATTACCGTTGATAGGAATACCTAAATATGCACATTTTCCTGTTCATTCAATTAGTCCCACCGCAAATAATGTTCTTTACCTACATACATTAGGTATTGTAATAATACATTGTCGCGATCTTATGCCGATGCATACTACCTGACAAGACGCTTCGCTCTGTACGTAAGGCGTCAGCTACATCTACAACGAGCGCTCTGTGGGTTGACTATCTTAAATCACTCGTCCTCGCCGCAAATATCAAGCCGTAACAAGTCT | 91% |
| scaffold64 | **Hd32** | 7916 | 88702-96617 | Hd32_DRJ1L | 626 | 95992-96617 | AGGGCTGACTTCATGCCGAATAACCTACTGCTCATTCACTGAGTACCTACGGGAGTCGCTCAAAAACTGCAATACATATGGCTTCTACTGCATGTGCATGCTTGACGCCGATCTACAGCCCAAAATACTGCAATGTAGTTGAAAGTGCAATCCATCGCATAATGCTTACTTTAGCACCTCGTTATGTAAGAACTCTGCATTCACGATACTACTTGCACGTTTCAGTAAGTGCATGAAAGATTCTCCGTGGTTAACCACCCACGGTACGTCATTTCATGTTTACGCCATGACGTTGGAGATGCTCTGCTCGAACGCTAATCATGGTGACAATGCAATAAGCTGGGTGGAATGCTGACGTTAAGAATGAAATCATTACCGTTGACAGGAATACCTAAATATGCACATTTTCCTGTTTATTCAATTAGTCCCAGCGCAAATAATGTTGTTTACCTACATACATTAGGTATTGTAATAATACATTGTCGCGATCTTATGCCGATGCATACTACTTGACAAGACGCTTCGCTCTGTACGTAAGGCGTCAGCTATATCTACAACGAGCGCTCTGTGGGTTGACTATCTTAAATCACTCGCCCTCGCGGCAAATCTCAAGCCGTAACAAATCT | |
| scaffold64 | **Hd32** | 7916 | 88702-96617 | Hd32_DRJ2R | 757 | 88702-89458 | TGAAGGAGATGATGAAGGAGCTGTGGGGCTGCAAACAAAGCGAGCGTCAGGGAGGAACCTGGAAGTAATGGAAGTGAAGTGTTGAACGAAAACCGCCGCACCAGTTATGTACGCAGCTGTCCCAACTAGCATGTAAGGTTTCAACTCTGGCTGTGTTCAGCGGTTGTGTGTATTAAGTGGTGATAGTGTTGTAAGCTTCTAAAATAAATAATTCGACTCCATTCGTATTTCTTTCCTTTAGAGACCGTGTGAGCTCGGCAAAATCTGCTCAGCTCACATGCATCTTTTCCCAGCAGTAGGACAAACATTTTCTTCAACTTGATCCTCTTGAGGCAGGCACTCATACCGAGGACAGACATACACACTTCTATTACTTCCAAGAAGCTCCTCCAATAGTTGGCATAGTCCGAGCTTGTACGGTGTCCACCCAAACAGTATAGCTCAAGAGGTATATGCATTCAACACTTAACACTACTTATTTCTACACCTATGCTAGGAATAATCATTTGTTTTATCTTACTGCTATCTTCAGGGCTACAGGTGCTAACGACAGTTCGTGCCAGCATCAGCTCACCTGAGTTTATGCTTATAACAGCTGCAGCACCCAAGGCTTTGCATTTCGTGTAGATAACACTTATCATTATTTACAGCTAAGTCTAGCTTCGCATATCTTGCCGTTGGGGACTTGGTGCAAGAGCTGGTCAAATGCAGGATACATGTTCACATGTTCATTGGTTAAGGGTACAGTTGACCATCT | 90% |
| scaffold64 | **Hd32** | 7916 | 88702-96617 | Hd32_DRJ2L | 750 | 93300-94049 | TGAAGGAGATGGTGAAGGAGCTGTGGGGCCGCAAACAAAGCGAGCGTCAGGGAGGAACCTGGAAGTAATGGAAGTGAAGTGTTGAAAGAAAACCGCTGCACCAGTTATGTACGCAGCTATCCCAACTAGCATGTAAGGTTTCAACTCTGGCTGTGTTCAGCGGTTGTGTGTATTAAGTGGTGATATTGTTGTAAGCTTCTAAAATAAATAATTTGACTCCATTCATGTTTCTTTCCTTTAGAGACCGTGTGAGCTCGGCAAAATCTGCTCAGCTCACATGCATCTTTTTCCAGCAGTAGGAAAAACATTTTCTTCAACTTGATCCTCTTTAGGCAGGCACTTATACCGAGGACAGACATACACACTTCTATTACTTCCAAGAAGCTTCTCCAATAGTTGGCATGGTCCGCGCTTGTACGGTGTCCATCCAAACAGTATAGCTCAAGAGGTATATGCATTCAACACTTAACACTACTTATTTCTACACCTATGCTAGGAATAATCATTTGTTTTATCTTACTGCTATCTTCAAGGCTACTGACGCTAACGACGGTTCGTGCCAACATCAGCTTACCTAAGTACACTTATGACACTGCAACACAATCATTTGCAATGCGTGTAGATAGCACTTACTACTACAAATAAGTCTAGCTTGGTAAATGTTGCTATTGCAGACTTGGTGCAAGAGCTGGTTATGCAGCACACGACGGGACGCATGCTCGCATTCGTTAAGGGTACGATTGAGCCTCT | |
| scaffold184 | **Hd41** | 7953 | 3768924-3776876 | Hd41_DRJ1R | 551 | 3768924-3769474 | AAGTATAACTCATAGGATGTTTGGCAATTCTCTGCATAAGCGCGGCTTTTTATTTCAAACAAGACGATGACTCTCGCATTTACAGTGACGTACTAAGGGGCGTGGGAAACTTGTGTCGGCTGGGTATGCCGGGTCGTATACTAAGTTTCCCACAACGCACGTAAACCAAAAAACTTCACTTAGACACAAACTATAGGAACGGACCGAACGTTTTCTCACTAACTGCACGTAACGTGAATGCTGCCATGTAATGCATTGCCGATCGCTGTCAAGCACCATTACTCACGTCAGCTTCGTCGCGATGATATCGTTATCACCGTGAGAACTGGTACTAAAGACTATTCAAAAGCAACGTCTGAGAGAGCACGCAGGATTTTCGAAGACCAACAAAGGTAAAACCGTCGCGTCAGAGTTGCATGGAGACTCGCAAACTGATCACAAACGACGTCTACCTTTCACACTCGTGAAAATGTTACATTGAGCTGTCAGTGGTACTTCATTCCACGCCGGCGAGAGTCCCTTTTGGTAGAAAAATATGCACGCATTTCTGT | 85% |
| scaffold184 | **Hd41** | 7953 | 3768924-3776876 | Hd41_DRJ1L | 540 | 3771924-3772463 | AAGTGTAACTTATAGGATGTTAGGCAATTCTCTGAATGAGCGCGGCTTTTCATATCAAACAATATGCTACCGTATTTATAGCGACACTAAGAGGCGTGGGAAACTTGTGTCGGCTGGGTATGCCGGGTCGTATCCAAGTTTCCCCCGACGCACGTAAACCAAAAAACTTCACTTGGACACAAACTATAGGAACACACCTAACGTTTTCTCACTAACTGCACGTAACGTGAATGCTGCCAGGTAATGCATTGCCGATCGCTGTCAAGCACCATTACTCACGTCAGCTTCGTCCCGATGATATCGTTATCACCGTGAGAACTGGTACTGAAGACTATTCAGAAGCGGTGTGTGAGAGCAGGCCAGATTTTCAAAGATGAACAAGGGTAGATCCGTCGTGTCATAGTTACATAGAGCAACTTTGCAACTGATCACAAGCGACGTCCACGTGTTAAAGTCGTGAAAATATTAAATTGAGTTGTCTGTACTATTTAATTCCATATCGGCGAGTGTCTAGGTAGGAAAATATGCGAGGATTTCTGT | |
| scaffold184 | **Hd41** | 7953 | 3768924-3776876 | Hd41_DRJ2R | 305 | 3771631-3771935 | AACTTGTAGCATCTAGTAGACTCAAAATACCAATCATGTCTGAGAGATCTATGTATCCGGTGGTGGCGTAGTTATCTGAGAAGCTAACAAAGCTCGTGCAGTATGAGGGGAACAATCATGCCGTCACCAGGCGAACCATGTGTATCACGTAGATAAGCTAGTCAGAGTAATGAACCAGCTATACATGTTACTTGCTCCCAACGATTACCAGAACGCCGGTAGTTCTACAGCAACCAAAAGAATTTGCTTTGCGATCCAAAGTAGGTCTATCCACCTATAGCTTTTACAAGCTGAAGTGTAACTTA | 81% |
| scaffold184 | **Hd41** | 7953 | 3768924-3776876 | Hd41_DRJ2L | 306 | 3776571-3776876 | AACTTGTAGCATCTGGTAGACTCGAAATAGCAATCATGTCTAAAGGATATATGTACCCGGTGGTGGCGTAGTTATCTGGGAAACTCACAAAGCTCGTGCAGCATGAGGAGAACCATCGTGCCGTCACCAGGCAAAGCAGCACGTATCACGTAAACAAGCTGCTCAGAGACATGAACGAGCTATGCATGCTATTGGCTACCAACGGTTACCAGACCGCCAGCAGTTCCGCAGCAAACAAAGACATTTGCTTTCCGACTTCAAGTGGGCTTATCCACCGATAGGTATCACAAGGTAAAGTATAAGTTA | |
| scaffold127548 | **Hd7** | 8066 | 6062921-6070986 | Hd7_DRJ1R | 362 | 6062921-6063280 | GATTTCGATTCCCTTCCGATCTCGTTTCACCTATCCTCGTCACTCTTACTCGACACCGACCAGAAACAACCCATTGGGTATATTGGTGGAACGAAGCGTGATTTACATCGAAATCTCCTGCGCAGCATTACGACCGGTTTATTGGTAAAACGATACGAAGTGAGAACGGAAGTGATCAAAATAAAAAGATGAAGGATTTATCGATTTGTTGCCTGACCATTTTCCTCAACGTTCAAATCCCTCCTTCTAGTCAGTTTCTTTTTATCGTGTAACCGAAACATCGCTTGGACAGCTGAATTTTGTTTCGTTGCATCTTTGGTGATATCTTTTTTTTGACGTCCTCGCGTGTATACTCGAGCC | 79% |
| scaffold127548 | **Hd7** | 8066 | 6062921-6070986 | Hd7_DRJ1L | 366 | 6070623-6070986 | GATTTCGATATCCTTCCCATTTTGTTTTACCTGTCGCGTCGTAACTCTTATTCGACACTCACCAGAAACACCCATTGGCAGGGTATATTGTTGGAACGAGGCGTCACTTACACCGAAATCTTCTACGCAGCGCCACGACCGATTTACTGGTAAAACGATGTGAAGATAGAACGAGAGTGATGAAAGGGAAAAGATGAAGGATTTTATCGATTTGTCACTTCACAAATTTCCTTACCGTTCGAGCCGCTCTTTCTAATCAGTTTCTTTATTGTATAACCGAAACATCACTTAGGCAGCTGAATTTTGTTTCGTTTAATCTTTTATCACATGTCTTTTCAGACGTCCACGCGTGTATGCTCCAGCC | |

| | | | | | | | | | | | |
|---|---|---|---|---|---|---|---|---|---|---|---|
| scaffold59 | **Hd11** | 9190 | 2183512-2192701 | Hd11_DRJ1R | 495 | 2186118-2186612 | ACCTACATCTGGGTTTTCAAAACATTCGTCGGACCGACCGAGAGTGTGAGATAATACGTCACTTGAGGACGATATTCTGATGACCTGTCACCTATCACTCATTGTTTTGAGATAATTTGATAAAATGTTGTATAAATACTCATTCCAACAACAAAAAGTCTCTTCTGTCAACATCCTCTGCTGAGATTGTTTCAATTATTTGGCAACAATGGCGACTTCTCAAATCGACAAATTGTTCGACAAGGATCCTTTCACCGGGAATACAATTTTTCACGAGCTCGCCTACATTGGATCACTGACGCTGCTCAAGCGATTCCGGAACAACCTCGACGAGCCATGTACCTTCATCCTGCAACAGTTTAACTCCGATGGTGAGTACAGTATCCACGTTGCAGCAAACGTCCACAGAGATCTACATGCAATTCAAGTGATCGAGCTACTGCGGGATTTGGGAGCAGATCTGGATGCAAGGGACGATCAATTAGGCATTACCGT | 97% | | | |
| scaffold59 | **Hd11** | 9190 | 2183512-2192701 | Hd11_DRJ1L | 493 | 2192209-2192701 | ACCTCCATGAGTTTCCAAAACATTCGTCGGACCGACCGAGAGTGAGAGATAATACGTCACTTGAGGACGATATTCTGATGACCTGTCACCTATCACTCATTGTTTTGAGATAATTTGATAAAATGTTGTATAAATACTCATTCCAACAACAAAAAGTCTCTTTTGTCAGCATCCTATGCTGAGATTTTTTCAATTATTTGTCAACAATGGCGACTTCTCAAATCGACAAATTGTTCGACAAGGATCCTATCACCGGGAATACAATTTTTCACGAGCTCGCCTACATTGGATCACTGACGCTGCTCAAGCGATTCCGGAACAACCTCGACGAGCCATGTACCTTCATCCTGCAACAGTTTAACTCCGATGGTGAGTTCAGTATCCACGTTGCAGCAAACGTCCACAGAGATCTACATGCAATTCAAGTGATCAAGGTATTGCGGGATTTGGCAGTAGATCCGGATGCAAGAGACGATCAATTAGGCATTACCGT | | | | |
| scaffold59 | **Hd11** | 9190 | 2183512-2192701 | Hd11_DRJ2R | 95 | 2183512-2183606 | CATCATAAGTCGAAGTAACTGACGGGGTGTTGGACGAATCGCCGGAATACCCACGGCTCAAAATATCGACAAACGTAATTGTGATGGTCGTGATG | 80% | | | |
| scaffold59 | **Hd11** | 9190 | 2183512-2192701 | Hd11_DRJ2L | 95 | 2187479-2187575 | CATCATAAGTTAAAGTAACTGCTGGGGTGTAGGACGACTTGCCGGAATAAACACGGCAGTGAATGTCGACAAACATAACTTTGATGGTGGTGCTG | | | | |
| scaffold429 | **Hd48** | 9673 | 531833-541505 | Hd48_DRJ1R | 616 | 536626-537241 | ATTTCTTTATGTTTCAGATTGCACTTGAAACGTGCGGGATACATGCGACCACTTTTTATCCATTCGCGATTACCTACATTTGAATCGTCAACCAATAAGAAGGCTTGATGCAGCCGTATCTCAATATCAAGGTGCCAGAGGTTCGTGACGTTCGTCGCTGTGTGATCATCGAGGGGCTACCAACCGTTGAAGTCGCGACTGCGTACGTGAATGCACGGACGACTCAGAGCAGACCATCAACTCGATCAAGCACAGATCCAAGTGAAAGATTTTACAACTGCTAGAGGGAACCACAAGGAGTCAGGATCGCGAACGAGAAGGATGCTGGATGTTCACAAGTATCAAGTAGAACGAGAAGCCTGATAGCTTAGAGATATGACTTCAAGCACCAGTTGCAACAGCAGGCACGCGCCACGTGCCGTAAAGTCTTTATGTTCTGATGATAACCCCAAGTACGTGATGCAAACCGATTTTATATGCAATCGCTACTTGGATACGGCGGACAACACGGTGTCGTGACGGAGTCTTGGAATCTGTAATTGCAACGCTGAGATTGCATGCTGGTTGCAATGCCAAGAAGCACGAAGTAGCTGCAGCACATGCTTGGACTCGGACA | 84% | | | |
| scaffold429 | **Hd48** | 9673 | 531833-541505 | Hd48_DRJ1L | 610 | 531833-532442 | ATTTCTTCATGTTTCTGATTGCACACAAAACGTGTGAGACACATGTGCGACAACTTTTTATCCAATCACGATTCCCTACACTTAGTTCGGCAACCAAGAGTGCTTGGGTAGCCGTTTTTCAATATCAAGATGCCAGAGGCTCGTGACGTTCGTCACTGTGTGATCATCGAGGTGCTATCAACCTTTGAAGTCCGTCGCGACTGCGTACGTGAGTGCACGGACGACTCAGAGCAGACCATCAACTCGATCGAGCACAGATCCAAGTGAAAGATTTTACAATCGCTCGAGGGAACCAGAAGGAGTTGCGATCATCAACGAGAAGGATGCTGGATTATTACAAGTCTCAAGCAGAACGAGAAGCCGTGATAACTTGGCGAGATAACTTCAGGCGCCAGTTGCAACAGCAGGCACGCGCCATGCGCTGTAAAGTCTGTATGTTCTGCTGGTAACCCCATACGTGGTGCGGACCGATTTTATATGCAATCTCTACTTGGATACTGCCGACAACACGGTTTGGCGCTGGAATCTTGGGCTGTGTGATTGCAACACTTGGATTGGATACTGGTTGTAATGCCAAGAAGCACAGAGTAGATACACCGCATGCTTGGAC | | | | |
| scaffold429 | **Hd48** | 9673 | 531833-541505 | Hd48_DRJ2R | 205 | 541301-541505 | GTCTTTGATTCTCCAGACCGACCACCAAGACCTACCTACCACTTAAATCGGACCGTCAACTCGATCAAGCACGAGATCGAAGAGAAAGATGACAGTCGCTAAAAAGGAAGTAATAGTAGTGACAATCGTGGAAGAGGAGGATGCTGGGATCTTACGAGTCCCCAAGCAGAGCGAGGGGAAAGTGACAGCGCAGAGATCGTAGCTC | 91% | | | |
| scaffold429 | **Hd48** | 9673 | 531833-541505 | Hd48_DRJ2L | 205 | 536223-536427 | GTCTTTGATTCTCCAGACTGACCTCTGAGACCCACCGACGACTCAAATCGGACCGTCAACTCGATCAAGCACGAGATCGAAGAGAAAGATGACAGTCGCTGAAAAGGAAGTAATAGTAGCTACAATCGTGAAAGAGTAGGATGCTGTAATCTTACGAGCCCCCAAGCAGAGGGAGGGGAAAGTGACAGCGCAGAGAGTGTAGCTC | | | | |
| scaffold65 | **Hd3** | 10014 | 437082-447095 | Hd3_DRJ1R | 156 | 441738-441893 | ACGTCCGCGAGAATGCGGACGATAACGTTCTCGAAAAGTGCATAGTGCAGATACTTTTCAGAGGCGAGCTCAGAGGTAAGAGAACGGTAGTTTAGACAGCGTAAGTGTCATACATGCAAGAAGTCGTTTGGATGCAGGTTTTATGCTATCGCTTCG | 76% | | 93% | |
| scaffold65 | **Hd3** | 10014 | 437082-447095 | Hd3_DRJ1int | 149 | 437664-437812 | ACGTCTGCGAGAATGCGGATGATAACGTTCTCGCAAAGTGCATAGTCCGGATACTTGGCAGAAGCAAGCCGCGAGGTTCCGTTTGGTCCGCGTAAGTGTCATACACGCGACAAGTCGTATCGATGCTAATTTTGTTCAATCGCTTCGTA | | 80% | | |
| scaffold65 | **Hd3** | 10014 | 437082-447095 | Hd3_DRJ1L | 159 | 437082-437240 | ACGTCTGCAAGAATGCGGACGATAACGTTCTTGAAAAGTGCATAGTGCAGATACTTGGCAGAGGCGAGCTCCGAGGTAAGAGTTACGGTAGTTTGGACAGCGTAAGTGTCATACATGCAAGAAGTCGTATCGATGCAGGTTTTATGCTATCGCTTCGTA | | | | |
| scaffold65 | **Hd3** | 10014 | 437082-447095 | Hd3_DRJ2R | 75 | 447021-447095 | GAGGGTCGGCGGCGTTATCCGAGAGGACGCTTGCAAACTTGACGAGCAGACGCATCATCGAAGAGCTCAGCCACG | 80% | | 85% | |
| scaffold65 | **Hd3** | 10014 | 437082-447095 | Hd3_DRJ2int | 77 | 441661-441737 | CTCGCAGGTCGGCGGCGTTAGCCGGAGGGCTCTTGCAGTCATGACGAGTGGACTTACAATCGAAGAACTCAGCCATG | | 89% | | |
| scaffold65 | **Hd3** | 10014 | 437082-447095 | Hd3_DRJ2L | 77 | 437587-437663 | CTCGAGGGTCGGCGGCGTTAGCCGGAGGACTCTTGCAGAAATGACGAGTCGACGTACAATCGAAGAGCTCAGCCACG | | | | |
| scaffold377 | **Hd4** | 10326 | 2459681-2470006 | Hd4_DRJ1R | 345 | 2469662-2470006 | TACGGACGGGCCATGATCCAACGGCATCGGCGCGGCCACGATATTGCAGTGCAGACTCTCTTATCAGCAGCAGGTTCTGCATCATGGTGTCGGCGGCCGTGACAGCGTCAATAGGTTTGAAAATTAGTTACTGATTACATGGGTCTCGAAGATTATACGGGTTTACACTTTGAAGACCCGTGCATTGTAGATCTTTAGAGCCTAACGTGGAGTTTCAGAGGGTTTGGTTCACAACCGTTTTTAATTCCACAATGCTTCGAGACGTTCGACACTACGTGACGAGGTGCTCAGCCGTTGATGCTGCGCTACATCGTGTCCACGAATGGATCACGCGATGCGATAACA | on 289nt 84% | | (3 fractions) on 160nt 88% | |
| scaffold377 | **Hd4** | 10326 | 2459681-2470006 | Hd4_DRJ1int | 433 | 2466097-2466529 | TACGGACGGGCCATGTTCCAACGGCGTCGGCGCGGCCACGATATTGCAGTGCAGACTCCCTTATCAGCAGCAGCTTCTGCATCATGATGTCGGCGACCGTGACAGTGTCAATAGGTTTGAAAATGAGTTACTGATTCCATGGGTCTCGAAGATCGTACAGGTTAACACTTCCAAGACCCGTGCATCGTAGATCGTTAGAGCCTAACGTAGTGGCTCAGAGCTAGGGTTTGGGGTGACCGTTTTGAATAGTACAGTGCTTGTTGCAATGTTCGTTACTACGTGATGAGTGCCAGTACGTGATGAACGTTGGAACCTTATACTTACGATCTCCGCAACGCGTTGTTATATGTACACCATAACGCCTTTAGCGAAGCTACTCTGCCGTTGAGGCTGCATTGTATCGCGTTTGCACGAGTGTCACGATGGGATATCA | | 72% | | |
| scaffold377 | **Hd4** | 10326 | 2459681-2470006 | Hd4_DRJ1L | 385 | 2459681-2460065 | TACGGACGAGCCATGTTCCTACGGCGTTGGCGCGGCCTCGATATTGCAGTGCAGCACTCCCTTATGATCAGCAGCAGGTTCTGCATCATGGATGTCGGCGACCGTGACAGTGTCAATAGGTTTGAAAATTAGTTACTTATTTATCATGGGTCTCGGAGATCTGCATGGTGTAAATCTTTAGAGAGAGTGCAGCAACCGTTTTTAATTGCACGACGCTTCGAGACGTTTGTCACTACGTGATGGATGTCAGTACGTGACGAACGTGGCGAGCTAATGACGATCTCTGCAACAGGTTGCGATATGTAGACCAACGCGTTTAGCGGACGTGCTCAGCCGTGGATGCTGCGCTACATCGTGTTCGCACGAATGTCACGATGCGATAACA | | | | |
| scaffold127548 | **Hd6** | 10461 | 5808388-5818848 | Hd6_DRJ1R | 311 | 5808388-5808698 | ACCGGCTAATGCTAGGATGGTCGTTGATTACGGGCAACCACGTTGACACATGTTGGCACCCAACGAGGGATCTTCACCCTGGGCCCTGGGCCTGAGATCTTTTGCCCGAGACTGCACCCGGCATCGGCAAACTATTCTGCCTCGGCAATTTTTCCTCGGTCGATCACTCTACTGTATCGTGATCCACCCCTTTGGAAACAACATGTTCGACACATTATCTAAATGAGTAGAACGTTTCTGTTTACAGCGTCATAGAGATGCGTTGCGCTTGTAAAAAATGGTCTGCATGGCAATTGCAGTCTCGCTCCGTG | 89% | on 270nt: 74% | | |
| scaffold127548 | **Hd6** | 10461 | 5808388-5818848 | Hd6_DRJ1int | 323 | 5813253-5813575 | ACCGGCTAATGCTAGGATGGTCGTTGATCACGGGCAACCACGTTGACACATGTTGGCACCCGACGACGGATCTGCACCCTGGGCCCTGGGCCTGAGATCATTCTCCCAAGACTGCACCCGGCATGGCTAACTATTCTGCCTCGGCAATTTTTCCCCAGTCGATCCACTCTACTGTATCGTGATCCACCCCCTTGGAAACAATACGTTCGACACATTATGTAAATGAGTAGAACGCGAGGTAGAACGTTTCTATTGACAGCATCATAGAGATGCGTTGTGCTTGTAAAAAATGGTCAACATGGCAATTGCAGTCTCGCTGCGTG | | | on 282nt: 71% | |
| scaffold127548 | **Hd6** | 10461 | 5808388-5818848 | Hd6_DRJ1L | 269 | 5817294-5817562 | GCAACTATGTTGACACGTGTTGGCGTCCAACGACGAATCTGCATCCTGAGTTTCATATCATTTGCTCAAGACTGCACCAAGCATGGCCAGCTATCCTGTCTCGGCAATTTTTTCACCGTCCGTCGTTCTACTGTGTCGTGATCTGCCCTCTTGAGAGTAATTAATTTAACACATCATCTGAATAAATAGAACCTCTTGGTTCAGAGCGTCATGGAGAGGCGTTGTGCTTATAAAAAATGGTTTCTACGGCAATTGCAGTCTCCGTTGTG | | | | |
| scaffold127548 | **Hd6** | 10461 | 5808388-5818848 | Hd6_DRJ2R | 101 | 5808626-5808726 | TGTTTACAGCGTCATAGAGATGCGTTGCGCTTGTAAAAAATGGTCTGCATGGCAATTGCAGTCTCGCTCCGTGGTGCAGCATGATACAGCAGAGCAAACCT | 75% | | | |
| scaffold127548 | **Hd6** | 10461 | 5808388-5818848 | Hd6_DRJ2L | 97 | 5818752-5818848 | TGTTCGCAGCGACATCGAGATGCGTTCGGCTTGAATAAAATGGTCTGCACAGCAATTGTACTCTCCATGGCGATGCATGGTACAGCGAAGTAAAACT | | | | |
| scaffold127548 | **Hd5** | 10510 | 12941247-12954959 | Hd5_DRJ1L | 209 | 12941247-12941455 | CACGTTCTAAACTCCACTCGTCCTTCGTTCACGGCATCGCCGTGCAGCACCCATTCCTCAATTCTTGCGCTGCACGAATTATAGTATAGCCTCGTTGTGAATACCTGAAGTAACGCATAACCCACTTCGGGAATGACGACCAAATGGTTGCCGATGGCTCGCACTTCCATCCCCGTTCGGTTATGCATTCACTGACGATAATCAGAGCT | on 209nt: 86% | | on 209nt: 70% | |
| scaffold127548 | **Hd5** | 10510 | 12941247-12954959 | Hd5_DRJ1int1 | 491 | 12946722-12947212 | CACGTTCTAAACTTCACTCGTCCTTCGTTCGCGGCATCGCCATGCAGCACCCGTTCCTCAATTCTTGCGCTGGGCAAGTTATAGTATAACCTTGTTGCGAATGCCTGAAGGAACGCAGAATTCACTTTGGGAATGACGGCAAATTGCTTGCCGACGGCTCACACTTTCATCCCCGTTCGGTTATGCATTCACTGCCAGTAAGCAAGCTCATCATAGACAACCCAATCTCATAAGGCCGAGAATCGTCCTCGATAACACGGGCGTTATCAAGAGCTGAGTCCCAGAACGATCTGCCATTTTTGTTATTGCGAGATGATAAAACAGCACATGTTCTTAACGCTTGTTTCTTATTACACACTTGTAAATTGCAAATGCGGGAATGAGAACTTAGCGTAGAGAATTGCTGCAACAATGCAGGCATCAGAAAATGAAGTTGGACGCCAGCAGAAATTCAGCGGTAGAATCTACAGATCTGTGTAAACACCCTCACA | | on 369nt: 84 % | | on 383nt: 86% |
| scaffold127548 | **Hd5** | 10510 | 12941247-12954959 | Hd5_DRJ1int2 | 369 | 12949974-12950342 | ACTCGTCCTTCATTCGCGGCATCGCCGTGCAGCGCCCGTTCCTCAATTCTCGCGCTGGGCACGCTATACATAGTGTTGTCTCGTTGTGAAACCTGAAGTAATGCATAACCCACTCGGGATTGACGGTACATTGCTTCACGACGACTCGCACTTTCATTCCCGTTCGGTTATGCATTCGCTGACGATGAGCGAGCTCATCTTAGACAACTTAATCTCGTAAGGCCGAGAATCGTCCTCTATAACATGGGCGTTATCAAGGGCTTGAGTCCAAGAACGATCTGCCATTTTTGCTATTGCGACATGATCGAACAGCTCATGTTCTTAACGCTTGTTGCTTATTGCATCCTTGTAAATTCCAAGTTCGGGAAT | on 254nt: 88% | | | |
| scaffold127548 | **Hd5** | 10510 | 12941247-12954959 | Hd5_DRJ1R | 464 | 12954496-12954959 | GTACTGAACTCCATTCGTCCTCCGTTCACGGCATCGCCGTGCAGCACCCACTCTGTTCCTCAATTCTTCTCCACTATAGCTCTTCGGGAATGACGGCGAATTGCTTGCCGACGGCTCGCACTTCCATCCCCGTTCGGTTATGCATTCACTGCCGATAAGCGAGCTCATCCTAGACAACCCAATCTCATAAGGCCGAGAATCGTCCTCGATAACATGAGCGTTATCAAGAGCTGAGTCCAAGAACGATCTGCCATTTGTGCTATTGCGACATGATGGAACAGCTCATGTTCTTAACACTTGTTCCTTATTCCACACTTGTCGATTTCAAGTGCGGGATTGAGAGCTTGCAGTATACAAAAGCGTAAAGAATGTTGCTGCAGCAATGTATGCATGAGAAGGTGAAGTTGGACGCCGGCAGAAATTGAGCGGAAGGATCTACGGATCTGTGTAAACAATTCCTCACA | | | | |

| | | | | | | | | | | |
|---|---|---|---|---|---|---|---|---|---|---|
| scaffold127548 | **Hd2** | 13937 | 5940296-5954232 | Hd2_DRJ1R | 637 | 5951349-5951985 | CATTACTGTCGGCCTCGAACATAGCAGCTGGATCGCGAGGAGACGTGTTCGTGAGAAGGTCTAGCGATCGGCTGTTATCACAGAGCCACCGTCCACGTTTACTGCACATGTTCTGCCCACACTTTGGAAATTCCGCATTTGCGGGATATGATAAACAAAAGTCGCGTACGCATACGCTGAAATACAGCATGCGTCACACGACATGTATAGGTATGGGGAATTACAAAATATCAAAACAAGAATAGAAACAGCGGAATGTACTCGAGCGCCATGATGTCATCCAGATTACACGGTTTCAGTTTGCGTTCTACATATAGTTTATGAACAAAGAAATCGACTGTGGCAGGTACACATGCAAAAAGGTTCTAGCAGATCTTTCCAGTTGCATTCTGGAATGCCAAAAAATCTGTCGCCAGCGTGCTTAGTTGACGCCGTGACTGCCGCTCCATGCAGTGCTAACCCAACTGATTGTTTGTCACGTACGTGACGGTCGCAGTGATGCTACTATGGAGATAACAAGTTGTGCACTGATTAACGTCCGGCAGGGCTTCACCTGTCGTACACCGACAACTTATTTCTTTATCGTATCTGGACTATACTGTTTACTGGAGCGGTCCCCTTTCCTGTAAAATAGTTT | 96% | | |
| scaffold127548 | **Hd2** | 13937 | 5940296-5954232 | Hd2_DRJ1int | 640 | 5945753-5946392 | CATTACTGTCGGCCTGAAACATAGTAGCTGGATCGCGAGCAGACATGTTCGTGAGAAGGTCTAGTGATCGGCTGTTATCACAGAGCCACCGTCCACGTTTACTGCACATGTTCTGCCCACACGTTGGAAATTCCGCATTTGCGGGATATGATAAACAAAAGTCGCCTACGCATACGCTGAAATACAGCATGCGTCACACGACATGTATAGGTATGGGGAATTACAAAATATCAAAACAAGAATAGAAATAGCGGAATGTACTCGAGCGCCATGATGTCATCCAGATTACACGGTTTCAGTTTGCGTTCTACATATAGTTTATGAACAAAGAAATCGACTGTGGCAGGTACACATGCAAAAAGGTTCTACCAGATCTTTCCAGTTGCATTCTGGAATGCCAAAAAATCTGTCGCCAGCGTGCTTAGTTGACGCCGTGACTGCCGCTCCATGCAGTGCTAACCCAACTGATTGTTTGTCACGTACGTGACGGTCGCAGTGATGCTACTATGGAGATAACAAGTTGTGCACTGATTAACGTCCAGCAGGGCTTCACCTGTCGTACACCGACATTTCATGTCTTTATTATCGCATCTGCACTACACCGTTTGCTGGAGCGGTCTCCTGTTCTGTAAAATAGTAT | | 86% | 86% |
| scaffold127548 | **Hd2** | 13937 | 5940296-5954232 | Hd2_DRJ1L | 651 | 5940296-5940946 | CATTATTGTCGGCCGCAAACATAGGAACTGGATCGCGAACAGACATGTTCGTGAGAAGGTCTAACGATCGGCTGTTATCACAGGGTCACCGTCCACGTTTACTGCCCACGTTTACTGCACATGTTCTGCCCACATGTTGGAAATTCCGCATTTGCGGGATATGATAAACAAAAGTCGCCTTGGCATACGCTGAAGTTCAGCATGCGTCACACAACATGTGTAGGTATGGGGAATTACAAAATATCAAAACAAGGAAAGAAACAGCGGAATGTACTGGAGTGCCGTGATGACATGCAGACTACACGGTTCCAGTTTGCGTTCTACATATAGTTCATGAACAAAGAAATCGAATACGGTGAGTACACATGCAAAATGTTCCAGCAGATCTTTCCGGTTGCATTCTGGAGCACCAAAAAATCTGTTGCCAATGGGCTGAGGTGACGCCGTGACTGCAGCTCCATGCAACGCTAACCTAAGTAATTGTTTGTCACGTACGTGACGGTTACCGTGATGTTGCTATGGAGATAAGAAGTTGTGCACTGATTATAGTCGAGCATGGTTCACCTGTCGTACGGCGACAACTTATGTTCTTATTATCGCATCTGCACTACACCGTTTGCTGGAGCGGTCTCCTGTTCTGGAAAATAGTAT | | | |
| scaffold127548 | **Hd2** | 13937 | 5940296-5954232 | Hd2_DRJ2R | 294 | 5944329-5944622 | TTCTGTATCATGCTGCAGCACGCAGCTAGACTGCAATTGCCATGCAGACCGTTTTTTACAAGCGCAATGCATCTTCTTAACGCTCTAAACAGAAACGTTCTACTTATGTAGAAAATGTGTCCAACAATGAGTTCCCGAGAGAGCGGATCGCGATACAAGAGAACGACGGTCCGTAAAGAAATTGCCGAGGCGCAATAGTTGACCAATGCACGGTGCAGTCTTGAGCAAATGATCTCAAGCCCGGGATGCAGATCTCTCATTGAGCGCCGACGTGTCAACATGGTTGCCTACCAT | 84% | | |
| scaffold127548 | **Hd2** | 13937 | 5940296-5954232 | Hd2_DRJ2L | 296 | 5953937-5954232 | TTCTGTATCATGCTGCACCACGGAGCGTGACTGCCATTGCCATGCAGACCATTTTATACAAGCGTAACGTATTTCCATGACGCTCTGAACATGAACGTTCTACGTATTTAGAAAACGTGAGCAACTGATTGTTCCCGAGAGAGCGGATCGCGATACACGAGAACGACGGTCCGTGAAAAAATTGCCGAGTCACAATAATTCACCAATGCACGGTGCAGTCTTGAGCGGAAAATCTCAAGCCCGATATGCAGATCCGTCATTGGGCGCCGACATGTGTCAACATGGTTCCTGACCAT | | | |
| scaffold90 | **Hd1** | NA | 1-14771 (partial) | Hd1_DRJ1R | 203 | 14569-14771 | GCGCCTCTCACGTACGCTCCTAAACGGTGATGAAGCTCGGTAATCATCTTCACTTCGACATGATCGACGACTCAATTCAGTCAATTCAATCGCATGTTTTCATTCTCGAATGTCAAAGGCCGATAAAGCCGGCAGTGCGCAGGAAGAGTGAACGGAAATGAAAACTGAGATTTCTCTATATTATTAAAAATTCAAAAAATTTA | 96% | | |
| scaffold90 | **Hd1** | NA | 1-14771 (partial) | Hd1_DRJ1int | 203 | 1-203 | GCGCCTCTCACGTACGCTCCTAAACGGTGATGAAGCTCAGAAATCATCTTTACTTCGACATGATCGATGAATCAATTTAGTCAATTCAATGGCATGTTTTCATTTTCGAATGTCAAAGGCCGATAAAGCCGGCAGTGCGCAGGAAGAGTGAATGGAAATGAAAACTGAGATTTCTCTATATTATTAAAAATTCAAAAAATTTA | | | |
| scaffold90 | **Hd1** | NA | 1-14771 (partial) | Hd1_DRJ2R | 255 | 12006-12260 | CAGCTAAGACACTATATGTAGTTTTAGAAGTACGCCATATCTACAGTTGCGCTATGAAGCACAATATTCTCCAGAACCTGAGTCTCTAATGTTGTATTGCATTTGTTGACGGTGGCCACAGCCCGCGCAGGAGCATGTCGTTCCATTCATATGACATTAGCCGAACCGCAAGCAAGCGGTTACACCCGATACGGTCATAGCAGATAGTGAGCTTTATAGTTTTTGTTACGACACATGTGTGTTTATATATATGTA | 89% | | |
| scaffold90 | **Hd1** | NA | 1-14771 (partial) | Hd1_DRJ2int-1 | 252 | 7539-7790 | CAGCTAATACACTGTGTAGTTTTAGAGGTACGTCGTATTTACAGGTGCGCTATGAAGCACATCATTCTCCCGGACCTGAGTCTCTAATGTTGTATTGCATTAGTTGACGGTGGCCTCAGCCCGCGCAGGAGCAGGTCGTTCCATTCATATGACATTAGCCAAACCGCAAGCAAGCGGTCACACCCGATACAGACATAGCAGATAGTGAGCTTTATAGGTTTTGTTACTACATATGTGTCTTATATGTGCGTA | | 74% | 76% |
| scaffold90 | **Hd1** | NA | 1-14771 (partial) | Hd1_DRJ2int-2 | 249 | 1114-1362 | CATCTAATACTCTATATGTAGTTTTAGCAGGACGTCGTGTGCTTAGTTTTGCACTTGCAAGTAGTATATTCTTCCGAACCTGACTCTCTGACGTCGCATTAGTTGACGGTGGCCACAGCCCGCGTAGGAGCATGTCATTCGATCCATACGTTACCCGAACCGAAAGCAAGCGTTTCCACCTGATACGGACTTACAGATAGTGAGCTTTATAGTCTATGAATTCGTGCAATATGTGTGTATATGTGTGTA | | | |
| scaffold128215 | **Hd9** | 17892 | 526917-544808 | Hd9_DRJ1R | 157 | 526917-527073 | TCCCAAAATCGTCAACTTGTGATAAAACATCCAACGAAGGCATCTCTCAGAGAATCCGGTGCAGCTCCAGGATGGTCCGCCGTATCGGATTCCACCACAACCCGATACAGCTGAACATCTGGGACATCTGACGGGAACTCTCTGCTCAGTTTCAATT | 90% | | |
| scaffold128215 | **Hd9** | 17892 | 526917-544808 | Hd9_DRJ1L | 156 | 542329-542484 | TCCCTAAATCGTCAACTTTTGATAAAACATCCAACGGAAGCATCTTTCAGAGAATCCACGGAGCTGCAGGATGGTCCACCGGCTCGGATTCCACCACAACCCGATACAGCTGAATATCTGGGACATCTGACGGGAACTCTCTGTTGAGTTTCAATT | | | |
| scaffold128215 | **Hd9** | 17892 | 526917-544808 | Hd9_DRJ2R | 305 | 530838-531142 | ATGCTAGGTTTCACATGCATTTGTGGGTTACGGCTGTACCTGTATGTACATGTAAGTGTCGTTTTGCACGTGCTATCAATCAGACAAAAAAGAAGGCGATAGTATAGACTCTCGTCCGTTCGGCTGCAGCTGCACTAGCGAGATCCACTCTGTGTCTCCTGTCTATAGCTTACTTATTACTTGCAATTGTAAAAGTATGGGAAAACTGAGGGCTAGGAACACAGTAGTTTTTATGTCGTCTTTTAATAGATCCCGTCAGCAAGAGATGAGATGGTTAATAAGTTAGCGAACTAGAGAGAGGCGGA | 85% | | |
| scaffold128215 | **Hd9** | 17892 | 526917-544808 | Hd9_DRJ2L | 308 | 544501-544808 | ATGCTAGGTTTCATATGCATTTGTGGATCACGGCTGTACCTGTATGTACTTGTAAGTGTCGTTTTGCACGTGCTATCAATCAGACAAAAAAAGAGGGCGATAAAGGAGAGACTACTCTCGTCCGTTCGGCTGTAGCTGCACCAGCGAAATACCCTCTCTGTCTCCTGTCTATAACTTACTCATTGCCGACAATTGTAAACGTATGGGAAAACTGAGGGCTAGGAAGACATTGGTGGTTGTGCCGTCTTTCAATCGATCTCTTCGGCACGAGATGAGATAGTTAAGTTAGCGAACTAGGAAAAGGCGGA | | | |
| **Campoletis sonorensis** | | | | | | | | | | |
| scaffold_11 | CsA | 6368 | 861628-867995 | CsA_DRJ1L | 244 | 861628-861871 | AATTGACGGCAAGCAGATCATGTGCCGTGCATTCCCCAGAGGTAATCGCAATTTGTGTTATCAAAACTGAGTCATGACTTGGCGACTCACTAGGCTGGATGCCAAGGATTCTGATCGCTGTCCACCGGCGGTAATGCACCGTCGTACCCGACCCGGAGATCATTTTCAACTTGACCGTTTCTTTTCAAAATTACGGAAAATATGGAACCTGTCTGCTTCCAAATGCAGTTGACGTCGGTTGTGA | 91% | | |
| scaffold_11 | CsA | 6368 | 861628-867995 | CsA_DRJ1R | 249 | 867747-867995 | AATTGACGGCAAGCACGGCATGTGCCGGCATGTGCAGTCCCCAGAGGTAATCGCAACTCGTGTTGTCAACGCTGAGTCGTGACTTGGCGACTCACTAGGCTGGATGCCAAGGATTCTGATAGCTGTCCAGCGGCGTTAATGCACCGTCGTACCCGACCCGGAGATCATTTTAAACTTGACCGTTTCTTTTCAAAATTACGGAAAATATGGAACCTGTCTGCTTCCAAATGCAGTTGACATCAGATGTGA | | | |
| scaffold_49 | CsB | 6626 | 22030-28655 | CsB_DRJ1L | 110 | 22030-22139 | AGACTTGCTACGGTTTAAGAGCTGCTGTCAGAGCTGATTCGTGATAGACCAACACGATGTACGCTTCCTGCGGGAACAGATAAGCACCCGGGCCACCCAGTTGATGCAGT | 76% | | |
| scaffold_49 | CsB | 6626 | 22030-28655 | CsB_DRJ1R | 112 | 28544-28655 | AGACTCGCTACCGTTCGGCAGCTGCTATACCAGAGCTGGTACGTGAGTAGTTGACACGATATACGCTTGCTTCGGGAACAGATGATCTCCCGGGACAGCCTGCCGATGCAGT | | | |
| scaffold_17 | CsE | 7990 | 1330025-1338014 | CsE_DRJ1L | 233 | 1330025-1330257 | GAGCTGTCTGGAGCCAACCAAGCAACGGAACGAAAAGAAACGAATGTCTATCATCGTTGAATTACAACCAGGGAACTTTTAAAAAAATCTTTCCTAGGATTGAACCACGGATTAGAACTATACATGCAGATCAAGCTCATGCGGATCGGTTCGTGTAAGGCACTCATGTGCATAGCTATTAGACCTCGCATGATCGAACAATCACGTTTTGAAGCGTGGGCACGATGCGGACC | 93% | | |
| scaffold_17 | CsE | 7990 | 1330025-1338014 | CsE_DRJ1R | 230 | 1337785-1338014 | GAGCTGTCTGGAGCCAACCAAACAACGAAACGAGAAGAAACCAATGTCTATCATCGTTTAATTACAACCAGGGAACTTTTTAAAAACTCGTTCCTAGGATTGAAGCACGGATTGGAGCTATTGCAGATCAAGCTCATGCGGATCGGTTCGTGTAAGGCACTCATGTGCATAGCTATTAGACCTCGCATGATCAAACAATCACGTTTTGAAGCGTGGGCACAACGCGGACC | | | |
| scaffold_131 | CsF | 8155 | 808380-816534 | CsF_DRJ1L | 259 | 808380-808638 | ACTGCCTGGTTCGTGTTTGTTCGCTAGCTTCTGCTCGCTTTGTTTCTAACCTAACCCAAGAATGTTCGCACGGACCCGTTATTGGTGCCCGATAACAGCTTTCTTTATACTCTTATGGTCCTGTTCTACAACCAGGTTCCAGCTTATTTCTATTTTTCAAATCCAGTATCGGCCCAGTTCACTGCTTCAAAAACAGTGAGCTTGTCAGACACGTAAACCATCAATTTTTTGTACAATCCACAATTTAAAAATAAAAAAG | 78% | | |
| scaffold_131 | CsF | 8155 | 808380-816534 | CsF_DRJ1R | 256 | 816279-816534 | ACTGCTGCGTTCGTGCTCATTTGCTAGCTTTCGCTCGCTTTGTCTGTACTTAGCCTGACAATGTTATCATGGACCTGTTATTGGTACTCGATGACAGCTTTCTTTATACTCTTATGATCCTGTTCGTCAACGAGGTTCAAAGTTGCTTCCATCTTTCAAATCTAGTATCAGCCCAGTTCGCTGTTCAAGAAATAGTGAGGTTGCCGGACTTGTAGACCATCAATCATTGGTATGCTACACAACTTAGAGATAAAAG | | | |

| | | | | | | | | | | |
|---|---|---|---|---|---|---|---|---|---|---|
| scaffold_10 | CsD | 8168 | 961052-969219 | CsD_DRJ1L | 711 | 961052-961762 | GAAAAGTTTTGGGAGAGTTGGCTGTTGAAGTTGCGAGTTGAACTTCAGTTTCTCCCATGTCAAAACTACGTTGAATGAAATTGAGTTGAGCTTAGACGACGTTC<br>GACGAAACGGCCAGAAGCTCAGAGAAGATTGTCAACGAGCATCCAGCACAGAGTCGAGTGAGCCTAGCCACGAAGTTCGTGCCGTACGAGATGAGTCGTCG<br>AAAAGCATAGGACGGCGATAGTACGTTATACCCCGTTGACGGATGGCAATGTCTCGTAAAGAAGTGAAAAGCTTCATGTTGAGCAAGTGGTGATCCGTCGATT<br>TGAACCGAGCATCACTCGCATGATAGCCTCTCGTAGTGTCACTTCTAGAAGAACCGCACGTCGCTTCTCAGCAACAATTTACAGGTTTGGCAGTTGTTACACC<br>GCAACTAAGCGACAATAACTGCATTTTGACATACGACAGGTTTGTATCGTTATCATTGCACGTGAAGTTTCATCCACCAATGGTGCTCGACCTGTGAATACATT<br>GGATAGCGTGACTTAGCGAGTACCTTCTTTTATCACGACACCAAACTGCGTCAACAGCGTCTACCGTCAGCTCCAATATGTGGATAGGAGCATGAGAGTTATC<br>TGCTGCTCGCATATACCGTCTTTTCACTGTCCCCGCCCCAGGTACACGCCAGATAAACTAACAATGTGCTCACGTACAGATTTCAAAAGTAGG | 98% | | |
| scaffold_10 | CsD | 8168 | 961052-969219 | CsD_DRJ1R | 711 | 968509-969219 | GAAAAGTTTTGGGAGAGTTGGCTGTTGAAGTTGCGAGTTGAACTTCAGTTTATCCCATGTCAAAACTACGTTGAATGAAATTGAGTTGAGCTTAGACGACGTTC<br>GACGAAACGGCCAGAAGCTCAGAGAAGATTGTCAACGAGCATCCAGCACAGAGTCGAGTGAACCTAGCCACAAAGTTCGTGCCGTACGAGATGAGTCGTCGA<br>AAAGCATAGGACGGCGATAGTACGTTATACCCCGTTGACGGATAGCAATGTCTCGTAAAGAAGTGAAAAGCTTCATGTTGAGCAAGTGGTGATCCGTCGATTT<br>GAACCGAGCATCACTCGCATGATAGCCTCTCGTAGTGTCACCTCTAGAAGAACCGCACGTCGCTTCTTAGCAACAATTTACAGGTTTGGCAGTTGTTACACCG<br>CAACTAAGCGACAATAACTGCATTTTGACTTACGACAGGTTTGTATCGTTATCATTGCACGTGAAGTTTCATCCACCAATGGTGCTCGACCTGTGAATACATTG<br>GATAGCGTGACTTAGCGAGTACCTTCTTTTATCACGACACCAAACTGCGTCAACAGCGTCTACCGTCAGCTCCAATATGTGGATAGGAGCATGAGAGTTATCT<br>GCTGCTCGCATATACGGTCTTTTCACTGTCCCTGCCTCAGGTACACACCAGATAGACCAACAATGTGCTCACGCAAAGATTTCAAAAGTAGG | | | |
| scaffold_14 | CsG2 | 8338 | 192247-200584 | CsG2_DRJ1L | 173 | 192247-192419 | GATCGACGGCAGCATCTTTTTCATCAACGAGTGACAATCGTCACCAGCCCCCGGCATTGTAGAATTGAAGAACGGATTCATAACCTTCTTGGTTGACAAACAG<br>AGCGTGTAGACGCAATAATCTATCGAGGGTCCCAAAGGCCACATACCGTACGATCTTTGAGACCGACGGA | 74% | | |
| scaffold_14 | CsG2 | 8338 | 192247-200584 | CsG2_DRJ1R | 173 | 200412-200584 | GATCGATGGCAGCATCTCTTCCATCAGCCAGCGACCAACATCACGAGCCTCCGGCATCCAAGAATTGAAGAACATTTGTATAACCTTCTTGATTGACCAGCGG<br>AGCGTGTGGATGCGATGATCGTTTGAGGGGTTCAAAGGTCGTATACTTAATAATTATTCAGACCCACGGA | | | |
| scaffold_14 | CsG | 8656 | 76017-84672 | CsG_DRJ1L | 372 | 76017-76388 | ACAGCATTTGACATTGAAAATGTTTCTCCGCGAGTGTGAGTAAGAGACGGTAAGATAATTTCTGTTATCTTCAGCTCTCACTTTTCACAATTGATCCCATGATAC<br>TGCAGAGTGCGGCCAGAATATGCTGCGCACTGTAAGCGTGGTGACAACTGTGTCCAGAGCAAGCTCCCACCATGTCCACTGTTTCGCGTAAACTTTTCTCGTA<br>GCAAAGACTTATTGCTATCAGTTACAGTGTCACACGGTGGACGCTTGGCACAGGTTGACAGGTTGGCCTGCTGAGGTTGGAGATATGCAACGGGAGCTGGTA<br>ACGCAACACACTCACTTCGATATACGCATGCTAGGGGAAAACAGTTAATATTCTGCTAATGC | 97% | | |
| scaffold_14 | CsG | 8656 | 76017-84672 | CsG_DRJ1R | 372 | 84301-84672 | ACAACATCTGACACTGAAAATGTTTCTCCGCGAGTGTGAGTAAGAGACGGTAAGATAACTTCTGTTATCTTCAGCTCTCAGTTTTCACAATTGATCCCATCATAC<br>TGCAGAGTGCGGCCAGAATATGCTGCGCACTGTAAGCGTGGTGACAACTGGGTCCAGAGCAAGCTCCCACCATGTACACTGTTTCGCGTAAACTTTTCTCGT<br>AGCAAAGACTCATTGCTATCAGTTACCGTGTCACACGGTGGACGCTTGGCACAGGTTGACAGGTTGGCCTGCTGAGGTTGGAGAGATGCTACGGGAGCTGGT<br>AATGCAACACACTCACTTCGATATACGCATGCTAGGGGAAAACAGTTAATATTCTGCTAATGC | | | |
| scaffold_22 | CsI | 8779 | 695663-704441 | CsI_DRJ1L | 204 | 695663-695866 | GACATCTCTATCAGAGGACATCTCTACCAATCCAGTCAAGCATTCCCAGCTCGAGGATTCCGATCTTCTTGAGTTGCGGTTGATGGATTAGATTAGTAGAGATG<br>TCCTCTGTTTAGAGATGTCCTCTCTACAGCCACCAGTTTTTTTTCTGTATTCAGTTTGCAAAAGTGAAATGTTGCAGACTAAGAGCGTAGACGAATCATG | 84% | | |
| scaffold_22 | CsI | 8779 | 695663-704441 | CsI_DRJ1R | 202 | 704240-704441 | GAAATCTCTACCACCGGACATCTCTACCAGTCCAGTCAAGCAATCCCAACCCAAGGATTCCAATCTTCTTGAGTTTCAATTGGTGGATTGGATGGTAGAGAGG<br>GCCTCTGGGTAAAGATGTCGACTCTACAGCCACCAGCTTTTTTCTGTATTCAGATTGCAAAAGTGAAATGGTAAAAACTAAGAGCGTAGGCGATTCATG | | | |
| scaffold_128 | CsI2 | 9042 | 110016-119057 | CsI2_DRJ1L | 174 | 110016-110189 | CGCCTGAAAACGGCCTTCATTTAATCTGGAAACGAACCTAAGACAGTTGGACTAACAAGAATCGTGGCTCAGCGACTACCTAATCCTTCAATGTAATCTAACCA<br>TGATTGGCAGTTTGTGATACAGCTTCCGCTGTTGTCAAAGTAAGCTAAACAGAAGATCGAGGGAAACAAT | 75% | | |
| scaffold_128 | CsI2 | 9042 | 110016-119057 | CsI2_DRJ1R | 177 | 118881-119057 | CGCCTGATAATGACATTCAGTTATTCCGGACACCAACCTCCTGCAGGTGGCCTCACAGGGCACGTGGCTCAGCGATTTCCTAACCCTTAAATGCAATCTCACT<br>ATGATTGGTACTTTGTAATACACCCCCCGCTGCTCTCAAGCCAAGTCGTTAAGCAGAAGAACGAGGGAAACAAT | | | |
| scaffold_38 | CsH | 9050 | 1398066-1407115 | CsH_DRJ1L | 731 | 1398066-1398796 | GCAAATGTTCAGTGCATAGCCTACCTGTCGTTGGACGGTGACGAAGTTGTGAACACCGTGCTATCATTTGAGCGAAGCGCTTTCGCATTTTTAATTTATTTTCC<br>AAAGTTGAATGCGTTAAGCCCCTTCAAATAGCCTTCAGATAGTTTTTAGCGGAGTTTTTCACAAGCTCAAGAATTGACGTTGGTTGCAAATGCTGCAACGGCAC<br>GTAAACTCGCCAGCTTACAATGACCTGATATCTCGCATTGCACAGCGCTTTGACTATAGTTTTTTGGTCGCTTATAGTTATTTTGGATGTCGTACGTATGCGTAT<br>AGTATGCATGGTATACCTCCCACATGTCTACTAATGTGCTTGTCCTCTTGAACGTATATGTTTTTTTTCTATGGATGTATGCAGGAGTGCGTTTTCGTTGTTTTCA<br>GTAATTTCGTTATCGTTATTCGTGTGGAGGATGCATTAAATACGTTCATGATATCTCAATGTGTGTGTGATAAGTTGCTTATTTTCATATGTGTGATCACTTTTAT<br>CAAGCAGTTCTTAATCGTAGAATAGTTAGACTAGATGCGGAAACTGGTTAGGCGTTACGATCGACACGATGTGCACATTGCCCTGAACACTGCTGATATTTTCT<br>TGTAAACGCGATTTTCATTATATTCGCGGCATACTATAGTATGCTAGATCAACTGTATCGACACGGTATCAAGTGTTCCCGGAGCAAGCGTGTAATCATGGT | 98% | | |
| scaffold_38 | CsH | 9050 | 1398066-1407115 | CsH_DRJ1R | 731 | 1406385-1407115 | GCAAATGTTCAGTGCATAGCCTACCTGTCGTTGGACGGTGACGAAGTTGTGAACACCGTGCTATCATTTGAGCGAAGCGCTTTCGCATTTTGAATTTATTTTCC<br>AAAGTTGAATGCGTTAAGCCCCTTCAAATAGCCTTCAGATAGTTTTTAGCGGAGTTTTTCACAAGCTCAAGAATTGACGTTGGTTGCAAATGCTGCAACGGCAC<br>GTAAACTCGCCAGCTTACAATGACCTGATATCTCGCATTGCACAGCGCTTTGACTATAGTTTTTTGGTCGCTTATAGTTATTTTGGATGTCGTATGTATGAGTAT<br>AGTATGCATGGTATACCTCCCACATGTCTACTAATGTGCTTGTCCTCTTGAACGTATATGTTTTTTTTCTATGGATGTATGCAGGAGTGCGTTTTCGTTGTTCTC<br>AGTAATTTCGTTATCGTTATTCGTGTGGAGGATGCATTAAATACGTTCATGATATCTCAATGTGTGTGTGATAAGTTGCTTATTTTCATATGTGTGATCACTTTTA<br>TCCAGCAGTTCTTACTCGTAGAATAGTTAGACTAGATGCGGAAACTGGTTAGGCGGTACGGTCGACACGATGTGCACATTGCTCTGAACACTGCTGATATTTT<br>CTTGTAAACGCGATTTTTATTATATTCGCGGCATATTATAGTATGCTAGATCAACTGTATCGACATGGTATCAAGTGTTCCCGGAGCAAGCGTGTAATCGTGGT | | | |
| scaffold_16 | CsX6 | 9213 | 504600-513812 | CsX6_DRJ1L | 234 | 504600-504833 | TAACTAGATTCTTACTTTGGTTCTGTGAGAACGTGACCCGGTCACGTTTGTCATTATTCGATCAAAACTTATTAGTCTTGCTTATTTTAATTCTTCACGGTTGAAA<br>GATTGTCCTCGAGATTCTGATTCCGAAGCATTAGAACGACTATGAATCGTCTACACTCTTAGTCTTCACCATTTCACTTTTGCGAACTGAATACAGAAAAAAAGC<br>TGGTTACTGTGGAGTGGACATCT | 89% | | |
| scaffold_16 | CsX6 | 9213 | 504600-513812 | CsX6_DRJ1R | 231 | 513582-513812 | TAATACATGCTTACTTTGTTTCTGTGAAAACATTACGTGGAGTTTGGTCATCATTCGACCAAAACTTAATAGTCTTGCTTATTTTAATTCTTCACCGTTGAAAGAT<br>TGTCCTCGAGATTCTGATTCCGAAGCATTAGAACGACTATGAATCGTCTACGCTCTTAGTCTTCACTATTTCACTTTTGCAAACTGAATACAGGAAGAAAGCTG<br>GTGGCTGTAGAGTGGACATCT | | | |
| scaffold_15 | CsJ | 9484 | 2621922-2631405 | CsJ_DRJ1L | 342 | 2621922-2622263 | TCCCTGCGAATCCACAGCAAGACTCCGTCACCGCTCCATCGTCGGCCACGCTGCTTGACATTGCTGAGCAGTTGAAAAGTTCTGCAGAAATGACGGCGTTAC<br>ACGGCATCATGCATGAGGGTACAAGGCTGCGACAAGCGACGACTGAGCTCAACCACGGTTTATTTTGTTCTAGCTTCAAGCGTCGCTGATGTACCTCTGAGC<br>GAAAAAAGATGCAGCAATGTTTTTACATTTGTACATTTTTTTTTAGTACCAGTAACCATTGATGTGTTGCTCGTTGTATACTGATGGATGGGAATCTGCTTTTGAG<br>TCGTCCCCCGAACATTTACGCATAACGCAAGC | 75% | | 91% |
| scaffold_15 | CsJ | 9484 | 2621922-2631405 | CsJ_DRJ1int | 246 | 2628392-2628637 | TTGACATTACTCAACATTTGAAACGTCCTATGGACCTAACGGTGTTGCTGGATCATGTATGCGGGTGCAAAGCTTTGACGAGCATTGACTGGGCTCACTTATG<br>GTTTTTTGTTGTAGCTTCAAGCGTCGCAGATGCATCATTAATTGCACAAAGGTGCATCAATATTTTCACATCTGTACATTTTTTTGATTACCGGTAACCATTGTTG<br>TGTTACTCGTTTTATTTGGACGGATGGGAATCCGCTT | | 73% | |
| scaffold_15 | CsJ | 9484 | 2621922-2631405 | CsJ_DRJ1R | 341 | 2631065-2631405 | TCTCTGCGAATCCACATCAAGACTACCTTCGCTACATCGTCGACCACACTGCTTGAAATCACTAAGTAGTTGAAACGTTCTGCAGAAGTGACGGCGTTACACG<br>GCATCATGCATGAGGGTACAGGGCTGCGACGAGCGACGACTGAGCTCAACCATGGTTTATTTTTGTTCTAGCTTCAAGCGTCGCAGATGTATCTCTGAGCGAA<br>AAAAGATCCAGCAATGTTTTTACATTTGTACATTTTTTCTTAGTACCGGTAACCATTAATGTGTTGCTCGTTCTGTACTGATGAATGAGAACCTGCTTTTGAGTCG<br>TCCCCCGAACATTTACGCATAACGCAAGC | | | |
| scaffold_35 | CsX8 | 9999 | 164467-174465 | CsX8_DRJ1L | 249 | 164467-164715 | TTGGTAGCTTGCTTAACGTCAGCATCCTTCTTGTTCGTGATCCGTACTGTTTTTGCCTTCATTTTCACATGAACATCATCTTTCTCTCCGTTCTTGCGATAGCGG<br>CAAACGAGGGATCACGTTTGAGTGGTTTGTCTGCTCGCGCATGTCATAGTTACAGCTATTCATGATCTCCAGCAGGACCTCTCCCATTGCACACTAACTAACAT<br>CGTGTGCTCGTGACATCGTGTAATAAAGAACAGTTGCTTC | 80% | | |
| scaffold_35 | CsX8 | 9999 | 164467-174465 | CsX8_DRJ1R | 252 | 174214-174465 | TTGTTGGCTTGTTGAATGTCAGCATCCTTCTTGTTCGTGATCAGAACTGTTTTCATCTTCACTTCAACTTTAACATCATATTCCTCTCTGTCCTTGCGATTGAAGC<br>AAACGATGGAGCTTGTTTGAGTGGTCCATCTGCTCGCGCGTATGTCATAGTGACTGATCTTCATGATCTCCGGTAGCACCTCTTCCACTGCTCACTGACGAAC<br>ATCGTGAGCTCGTGGCATCGTGGAATTGAAGAACAATTGCATC | | | |

| | | | | | | | | | | | | |
|---|---|---|---|---|---|---|---|---|---|---|---|---|
| scaffold_4391 | CsL | 10024 | 8079-18102 | CsL_DRJ1L | 660 | 8079-8738 | TTTAGAACCTTACGGACACAGTTGATCTGCTCCGTCTTGCCAAGTGCCTGCCATGGTAGCTTATGCTTGTCATCTGTCTGAGTTCGTTAACGCTTACCTGTTCTGCTTGGCGGCTTGTTCAATCTCAGCATCCCTCTTGTTCGTGATCAGAACGGTTTTTACCTTCATTTTAACATAAACATCATCTTCCTCTCCGTTCTTACGATTGAGGCAAACGATGGTTCGCGTTTGAATGGTTTGTCTGCTCGCGTATGTGATAGTTACTGATTTTCGTGATCACCGGCAGCACGTTTTTCATTGCTCACTGACAAACATCGTGAGCTCGTGGCATCGTGGAATTGAAGAACAATTGCATCACATGGTTTTGGTTGACAAACAACTCTGTCATAATAATATTATTTGTCGTGCATGAGTCCGACAGGTCTTGCACCGCACAATCTTCGACAGTCGTGCAATTTTTTGGCAACCTTCACATCCATTTCAATGTCGACATTACCATGTAAAGCTTGCTGAAGATCGCTCTTCTGCGGATCCAAGTCAGCGTCAACGCTATAGGATGAAGATTCGTCAGTAAACTCGTTTTAGAAGTACTGTAGGTCAAATCCGTGGTATATCAGAACTTTTCGAATACGAAGTTGTCCTCGTCACAAAAACATACGC | 93% | | | | |
| scaffold_4391 | CsL | 10024 | 8079-18102 | CsL_DRJ1R | 634 | 17469-18102 | TTTAGAAACTTACGGACACAGTTGATGTGCTCCGTCTTGCCAAGTGCCTGCCATGGTAGCTTATGCTTCTCATCTGTCTGAGCTCGTTGACGCTTCCTTGTCCTGCTTTACGGTTTGTTCAATGTCAGCATCCCTCTTGTTCGTGATCAGAACGGTTTTTACCTTCATTTTAACATAAACATCATCTTCCTCTCCGTTCTTACGATTGAGGCAAACGATGGTTCGCGTTTGAATGGTTTGTCTGCTCGCGTATGTGATAGTTACTGATTTTCGTGATCACCGGCAGCACGTTTTTCATTGCTCACTGACAAACATCGTGAGCTCGTGGCATCGTGGAATTGAAGAACAATTGCATCACATGGTTTTGGTTGACAAACAACTCTGTCATAATAATATTATTTGTCGTGCATGAGTCCGACAGTCGTGCAATTTTTTGGCAACCTTCACATCCATTTCAATGTCGACATTACCATGTAAAGCTTGCTGAAGATCGCTCTACCGCGGATCCAAGTCAGCGCCAACGCTATAGGATGAAGACTCGTCAGTAAACTCGTTTTAGAAGTACTGTAGGTCAAATCCGTGGTATATCAGAACTTTTCGAATACGAAGTTGTCCTCGTCACAGAAACATACGC | | | | | |
| scaffold_5 | CsN | 10943 | 175169-186111 | CsN_DRJ1L | 254 | 175169-175422 | CTCTCTATTTGCTAGATTCATTGGAAAGTTACGTGTCCCCATATATATTAGCCAAAGTCATACACATGGGAGGAGCGAACGTGAATTCGAGGCTCTTAAGAGATCCCCAGGTAAAAGTTCCAAAACAAGTTGACGTTGCACAATTTGGGTACGTGTAGTTTGCACACGGGATATGGAGCTTTCAAGTTGAGAATGAAACCAACGGGTCCGGATATTGTTTAGTTTTCAAGAGCACCGCGTAACGTAATCAAAGA | 78% | | | | |
| scaffold_5 | CsN | 10943 | 175169-186111 | CsN_DRJ1R | 258 | 185854-186111 | CTCTCAATTTGCCAGATTCATTGAGAAGTAATGTGTCTCCGGATTAACTAGCTGGAATATTACTAAGATGGATGGAGTGAACGTGAATTTGAGGCATTTTAGAGATCTCCATGTAAAAGTTAATAAGATAAGTTGACGTTGCACGGTTTGGGAACTGGTAGTTATGTACACGGTACATGAATCTTTCAAGTCAAGGATGAAACCAACGGGTCGGGATATTGTTTACTTCTGAAGAGCAGTGCGTAACGTACTCAAAGA | | | | | |
| scaffold_110 | CsX2 | 10806 | 248068-258873 | CsX2_DRJ1L | 123 | 248068-248190 | GAAATACTATTTCACGACCTTGTGATTCATAGATCGTTGACGTTGACTATACTGAATTGGAGATAGAATCGACAACCCATGCAATTGTCTCCTGCAAACTTCGAAATATGCAGGTTGAAGAGC | 74% | | | | |
| scaffold_110 | CsX2 | 10806 | 248068-258873 | CsX2_DRJ1R | 125 | 253752-253876 | GAAATACCATTTTCCACGGCCTAGCCATGGATAGATTGTTGAAGTTATCTATACTGAATTTGGAATAGAATCGACAGACCATGCAATTCTATTCTCCGAACGACGAACTGTAAAAGTTAAATAGC | | | | | |
| scaffold_110 | CsX2 | 10806 | 248068-258873 | CsX2_DRJ2L | 127 | 249564-249690 | TGTGCAGAACCGAAAGTGATTGAAAGTGAATCGAGTAATTAGAACGATTAAACATAAGCTAGTATGTGGGCAGGTTGTTTGTGTATTATTTTTTGATTTTCGAAATGATGAAATTATGCCATAAAAT | 72% | | | | |
| scaffold_110 | CsX2 | 10806 | 248068-258873 | CsX2_DRJ2R | 128 | 258746-258873 | TGAGCAGAGCCGAAAGCGAATGAAAGTATATCGAGTGACAATCGGCATTGGACGTGAGATAGTAGGTAGACAGATAGTGTATTTTTTTTTGTTTTGATTTTCGAATTGTTAAAATTGTGCAATGAAAT | | | | | |
| scaffold_5218 | CsP | 12113 | 15720-27832 | CsP_DRJ1L | 277 | 15720-15996 | TGCTTAGCTCTGAGTCAGATTGTAGTTCAATTACTGAGTGAAAGGTGGTTGTCTTGAATTTCCAATGGTGTTAGCTATTTGGAACCCACTGAAAAAAATAGCCACCTGTTGCGGTTGGTGTTTACATTGTTTCCCTCGTTCTTCTGCTTAGCTTGACTTGAAAGCAGCGGGGGGTGTATCACAAACTCCCAATCATAGTGAGATTACATTCAAGGGTTAGGAAATCACTGAGCCACGTGCCCTGTTAGGCCATCTGCACGCGGTTGGTGTCCGCGAT | 72% | | | | |
| scaffold_5218 | CsP | 12113 | 15720-27832 | CsP_DRJ1R | 274 | 27559-27832 | TGCTTAGCCAGACGTCGGCCTGTAGTACTATCACTGAACGAGACGGCGCTGTCTTGTGTTTCTCATGGTGCTACTTAACTGGAACTCGAGGAAAAGACAGCCACTTATATCGGTCGAAGTCAAACTGTTACCCTGGATCTCCTGTTTAGCTTACTCTGACAGCAGCGGGAGCTGTATCACAAACTGCTGATCATGGTTAGATCACACTGAAGGGTCAGGTATTCGCTGAGCCACCATTCTTGTTAGTCCAACTGCCCTAAGTTCGTGTCCGGAT | | | | | |
| scaffold_23 | CsM | 12197 | 1422227-1434423 | CsM_DRJ1L | 307 | 1422227-1422533 | AACATTTTCTAAAGTTCCCACGAATTTTTCGGGCAACTGGTACGTTCTCCAGCCATCGTACAGAACTGTTTAGGGATATTTTCACAGCGAGACGACGGAGTGAAATTCGATCGACAAGAAAGTATGTCTTTGAGTACGTTACGCGGTGCTCTCAAGAAGTAAACAATATCCCGACCCGTCGGTTTCATCTTTGACTTGAAAGCTGCATGTACCATGTGCAAACTACCCGTTCCCCAAATTGTGCAACGTCAACTTGTCTTGGTAACTTTTACCTGGGGATCTCTTAAGAGCCTCAAATTCACGTTCA | 87% | | | | |
| scaffold_23 | CsM | 12197 | 1422227-1434423 | CsM_DRJ1R | 307 | 1434117-1434423 | AACATTCTGTAAAGCTCCCGCGAGCTGTTCGGATGACTTCAACGTTGTCCAGCCATTGTACAGAACTGTTTAGGGATATTTTCATAGCAAGACGAACTAGTGGAATCCGCGCGACGAGAATGTACGTCTTTGATTACGTTACCCAGTGCTCTTGAGAACTAAACAATATCCCGACCCGTTGGTTTCATCCTTAACTTGAAAGCTCCATGTACCGTGTGCAAACTACCCGTTCCTCAAAATGTGCAACGTCAACTTGTCTTGTTAACTTTTACCTGGGGATCTCTTAAGAGCCTCAAATTCACGTTCA | | | | | |
| scaffold_50 | CsQ | 12543 | 290527-303069 | CsQ_DRJ1L | 437 | 290527-290963 | CAATAAGGTTGATTGAACAGACCAATGTTGTTGCAGGCAGCTTAAGGACTGCTTGTAAAATGATGATCCATGGGATCGTCGGTCTCAGTTTGACCAAGTTTTGATTTTAATAGCAATTGAAATAACGACGCATTGTAATCCGTCTTCGGACCATATAGTGCTGGCTCCATGTCCATGTCGGTAAAGAGGCTCACGTGCTAAGGAAAACATGCGTTTTACGAGATAAACCATTGTATTCATGTCTGGATTCTACGAAACCATTGAATTAATCGATCTGATTGTAATTCATACTTGGACCATAATACATGCAGAATAATCAAGTAGTAAAGGGGCTTGCGAGTCAAAAGACACGTGGCCTCTACCGTCGTGCAATTGAACGAACAAAACAGATTCTTACGTCTGTTCTGTGGCAACATGGCCTGCAGTTAGGTTATCAT | 88% | | 87% | | |
| scaffold_50 | CsQ | 12543 | 290527-303069 | CsQ_DRJ1int | 383 | 293120-293502 | CAGTAAGGTTGATTGAACAGACCAATGTTGCTGTCCAGGCAGCTTGAGGACTGCTTGTAAAATGATACTCCATGGGATCGTCGCCATCACATCGACCAAGTTTGGATTTCAATAGCAATTGACATAACGATACATTGTAATCCGTTTTTGGAACATATAGTGCTGGCTCCATGTCCTTGTTGCAAAAGAGGCTCACATGCTAAGGAAAACATGCGTTTTACGAGATAAACAATTGTATTCAAGTCGGGATACTACGAAACCATTGATATAATCTGATTGTAATTCATACTCGGACCATAATAGATGCAGAATAGTCAAGTAGTAAAGGGGCTTAGGTGTCAAAAAACACGTGTGCTCTACCGTGGTGGTATTGAACGAACAAA | | 79% | | | |
| scaffold_50 | CsQ | 12543 | 290527-303069 | CsQ_DRJ1R | 408 | 302662-303069 | CAGTAACGTTGATTGAACAGATCAGGGGTGCTGTTGCAGGCAGCTTAAGCAACTTCTTGTAAAATGACAATCCATTTTTAATAGCAATTGAAATAACGACGCATTGTAATCCGTTTTCGGACCATATAGTGCTGGCTCCATGTCCATATCGGAAAAGAGGCTCACGTGCTAAGGAAAACATGCGTTTTACGAGATAAACAATTGTATTCATGCCTGGATTCTACGAAACCATTGAATCAATCGATCTGATTGTAATTCATACTTGGACCATAATACGTGCAGAATAATCAAGTAGTAAAGAGGCTTGCGAGTCAAAAGACACGTAGGCTCTACCGTCGTGCAATTGAACGAACAAAACAGATTCTTACGTCTGCTCTGTGGCAACATGACCTGCAGTTAGGTTATCAT | | | | | |
| scaffold_149 | CsU | 15338 | 374074-389411 | CsU_DRJ1L | 511 | 374074-374584 | AACCGTGGTCATCTTTGTCGCCACCAAGATCTGTCTGTCTCAAAGTGACGCTGACCTCTCAATTAGCTGTTCGAGTTGTTGATGGAAGGATGTGCATTCGAAGTTCCACGCTCTGCAATGTATTATAAGCTCGACTGCGCCATTGGTGGAGGATGATATGGCAGGTCGAAAAGATGGTACCAAGTGACGTGTACTTTTATTTTAAACGACACCACAAGAGGTTATCGTGCGGGTGCTGGTCGGGTCGAACAATGTCCACAAACCACTACCTGTACGATGTGAAGCTTCCACTAGTTTATCACGCAGAGCTGCCGTTCCACGGGATTGCACGGCGCCATCGCTTGCGTTTACTCTATACGTGTACTCGTGCAGTACAAGCTTGATAGCTATGTTCATATGACCACATGCTTGGTGGGCAGTGAAAATATTGCAACAACGACCGCTCCATCTAACAATGTACAAACATGATTCATGTGCTATTGCGATAGTTTTGATGCAGATTGTTCGACTA | 96% | | | | |
| scaffold_149 | CsU | 15338 | 374074-389411 | CsU_DRJ1R | 511 | 388901-389411 | AACCGTGGTCATCTTTGTCGCCACCAAGCTCTGTCTGTCTCAAAGTGACGCTGACCTCTCAATTAGCTGTTCGAGTTGTTGATGGAAGGATGTGCATTCGAAGTTCCACGCTCTGCAATGTATTATAAGCTCGACTGCGTCATTGATGGAGGATGATATGGCAGGTCGAAAAGATGGTACCAAGTGACGTGTACTTTTATTTTAAACGACACCACGAGAGGTTATCGTGCGGGTGCTGGTCGGGTCGAACAATGTCCACAAACCACTACCTGTACGATGTGAAGCTTCCACTAGTTTATCACGCAGAGCTGCCGTTCCACGGGATTGCACGGCGCCATCGCTTGCGCTTACTCTATACGTGTACTCGTGCGGTACAAGCTTGATAGCTATGTTCATATGACCCCATGCTTGGTGGGCAGTGGAAATATTGCTATTACGACCGGTTCATCTAACACTGTACAAACATGATATATATGCTATTGATATAGTTTTGATACAGATTGTTCAACTA | | | | | |
| scaffold_28 | CsW | 15807 | 614005-629811 | CsW_DRJ1L | 223 | 614005-614227 | GATTCTGCTGAATGATTCTACAGGCCATCATCCGTTTAGGTTTGACACTGCGCTCGATGTCCCGTGGTTACCCCGGCTTACAGTACTTAGTTACTAACTACTCAACAGAATGCAAACTTATTTCGATGTTTAGAGTTCTAGCACGAGGAACAGCAGCAGGACGCGGGTAGGTGCTGTAACGGTGTAACGAATGCCTGCATTGCATTTTATGCCATAATCCATC | 86% | 90% (on 184nt) | 77% | 77% | |
| scaffold_28 | CsW | 15807 | 614005-629811 | CsW_DRJ1int-1 | 223 | 616606-616828 | GATTCTGCTGGATGAATCTACAGGTCACCATCCGTTTAGGTTCGACACTGTGCTTGATATCCTGTGGTTACCCTGGCTCACAGTAATTAGTTACTAACTAATCGACAGATTGAAAAAGTTTTTCGATGTTCAAAGTTCTAGCATGAGGAACAGCAGCAGAAAGCGGGTAGGTGCTGTAGCCGTGTAATGAATGCCTGCTATGCATTCTGTGCTATAATCCATC | | | | | 75% |
| scaffold_28 | CsW | 15807 | 614005-629811 | CsW_DRJ1int-2 | 206 | 619316-619521 | GATTCTGCTGGATGAATCTACAGTATGACACTGTGCTTGATGTCCTGTGGTTACCCCGGCTTACATTTCTTAGTTACTAAGTACTCAACAGAAAGAAAACCTATTTCAATGTTTAAAGTTCTAGCACGAAGAACAGCAGCAGGACGCGGGTAGGTGCTGTAACGGTGTAACGAATGGCTGCTATGCATTCTATGCTACAATCCATC | 76% | | | | |
| scaffold_28 | CsW | 15807 | 614005-629811 | CsW_DRJ1R | 223 | 629589-629811 | GATTCTGCTTAATGATTGTACATGTTATCATTCAATTGGGTTTGACCCCGTGTTCGATAACCTGTGGTTCTCCCGACTTACAGTCATTCGTTACAAACCACTCAATCAAATAAAAACTTATTTCGACGTTTAAAATTCTAGCACATGGAACAGCAGCAAGACGCGGGTAGACGCTGTAACCCTTTAGTGACTACCTGCCATGCATTCTCCGCCACGATCCATC | | | | | |
| scaffold_5890 | CsZ | 15871 | 134147-150017 | CsZ_DRJ1L | 99 | 134147-134245 | AAACAATCGCATCAAACGCTCTTGGTTGACGTACAATGTGTGTGTGGATGCAATGGTTCACCGTGGGTCCCAAAGATGGCATTTCGTACGATCTTCAAG | 77% | | | | |
| scaffold_5890 | CsZ | 15871 | 134147-150017 | CsZ_DRJ1R | 97 | 149921-150017 | AAACAGTTGCGTCAAAGACTTTTGGTTGACACTCAATGTGTGTGGATGCAATGGTCTACCGCAGGTCCCATAAGTCGCATACCGTACGTTCTTCGAG | | | | | |
| scaffold_5934 | CsX1 | 17335 | 19391-36725 | CsX1_DRJ1L | 214 | 19391-19604 | GTATACACCACCGTACCACTTGGAACGCAAGAACTCTGATGGATTGTGGCAATCTTCCAATAAAGTACCTCGAACTTGTGTTATGGATCTGTATGGACGTGGAACCGTTGCCGTGCAAGATCCTTATAGAACGTTTTATCCATCGACACGTGTGCGTGAGGACGTTCCTCTCAAAGTCGGTGCAATATCATCATGCGATAACGGTTGCTGCAAT | 76% | | | | |

| | | | | | | | | | | | |
|---|---|---|---|---|---|---|---|---|---|---|---|
| scaffold_5934 | CsX1 | 17335 | 19391-36725 | CsX1_DRJ1int1 | 216 | 21903-22118 | GTATGCACGTCACGGTACCACTTGGTCCGCAAGCATTATGATGGGTTGTGACAAAGTTCCAGTATTCTACCTCGAACATTTGTTGTCAATCTTTATGGACGTGG GCACGTGGTCGTGCAAAAGCCTTATGAAACGTTTTACTTATCGTAGAACGTGCGCGACGACGTTCGCCACGAAATCAGTGCAATATCATTGTGCGATAACAGT TGCTGCAAT | [illegible] | 72% | 88% | 78% |
| scaffold_5934 | CsX1 | 17335 | 19391-36725 | CsX1_DRJint2 | 216 | 27254-27469 | GTATACACGCCACGGTACCACTTGGGCCGTAAGCAGTATGATGGGTTGTGGCAAAGTTCCAGTCTCCTACCTTAAGTATTTGTTGTCAATATTTATGGACGTAG GCACGTGGCCGTGCAAAAGCCTTATAAAACGTTTTAGTCAATGGAGAGGTTTTCTAACGTCGTTTGTCTTGAAGCTGGTGCATTTATCAATGCGAGATGACAGT TGCTGCAA | 85% | | | |
| scaffold_5934 | CsX1 | 17335 | 19391-36725 | CsX1_DRJ1R | 216 | 36510-36725 | GTATACACGCCACGGTACCACTTGGCCCGTAAGCAGTATGATGGGTTGTGGCAAAGTTCCAGTCTCCTACCTTCAACATTTGTTGTCAATCTTTATGGACGTAG GCACGTAGCCATGCAAAAGCCTTATGAAACGTTTTATTCATCGGAGAGCGTGCGCCACGACGATCCTCTCAAAGTCGGTGCAATATCATCGTGCGATAACAGT TGCTGCAAT | | | | |
| scaffold_116 | CsT | 23217 | 7789-31005 | CsT_DRJ1L | 638 | 7789-8426 | GTACGCCGCGACGTCACAGTGTTCGACAGTCTAGCCTCCACACCAATTAGCTATTACGAAAGCACCGAAGTAGTTTTTAAGCACTGACTATGAATATTAGCCAT GACTTCACAGCACCGAACATCCAAAGGACGCTGCATATTAGTTTGTTCTTCGATGGGCGGTTGTTTGTAAATGCACTGCTTTCGAGACAGGAAAAGGAAGAGG ATTTCATTAGAAAAGGGTCATACAGATGCTAACTTCCAACGTCAGAATCATTAGTCATATTCTGAAGAGTTCTACAACTTTTGTAATTAATCAATTGAAAAATCGT ACAATATATTGATTTAATAATAAGAAATAACTTGGTTGATATCTTGGTTATTTATAGGTCATTCCAAAAATGCGAATAAAATAAAAATAAATTAGCAAGAAATTTTA ACTCGTAACACACTTTTTACGATCGGATCACTAATTAGTATTAACTCAAGCCTGTTAATTATTTATTTTTTTCCTCAAGATAAAGATTGACATTATAATTTCTGCGA TGGTCGTAAAAGCGCGCATATTTGCTGGTTTCAATTTACGACCCATGTTAACATTTAACACAACGCGACGTATTCGCAGCCTGGTACACTGAACTCTGAAAAGC ATTGTGA | 96% | | | |
| scaffold_116 | CsT | 23217 | 7789-31005 | CsT_DRJ1R | 640 | 16673-17312 | GTAAGCCGTGACGTCATAGTGTTCCACAGTCTAGCCTCCACACCACTTAGCCATTACGAAAGCACCCAAGTAGTTTTTAAGCACCGACTATGAATATTAGCCAT GACTTCACAGCACCGAATATCCAAAGGACGCTGCATATTAGTTTGTTCTTCGATGGGCGGTTGTTTGTAAATGCACTGCTTTCGAGACAGGAAAAGGAAGAGG ATTTCATTAGAAAAGGGTCATGCAGATGCTAACGTCCAACGTCAGAATCATTAGTCATATTCTGAAGAGTTCTACAACTTTTGTAATTAATCAGTTGAAAAATCG TACAATATATTGATTTAATAATAAGAAATAACTTGGTTGATATCTTGGTTATTTATAGGTCATTCCAAAAATGCGAATAAAATAAAAATAAATTAGCAAGAAATTTT AACTCGTAACACACTTTTTACGATCGTATCACTAATTAGTATTAACTCATGCCTGTTAATTATTTATGGTTTTTTCCTCAAGATAAAGATTGACATTATAATTTCTG CAATGTCCGTAAAAGCGCGCATATTTGCTGGTTTTAATTTACGACCCATGTTTACATTTAACACAACGCGACGTATTTGTATCCTGGTACACTGAACTCTGAAAA GCATTGTGA | | | | |
| scaffold_116 | CsT | 23217 | 7789-31005 | CsT_DRJ2L | 1093 | 15405-16497 | TACATACATGTAGCGCACAATCGCCTAATTAGATCATCGACATAGAAGCTGACATGGAATAAACGAACTCTATCATCTTGAAGGTTTGGGTTTTTCGAATACATC TAATTCGGTATCGAAGCAGCCAGAACGTCGAACATGTCCATACATAAACCGTGCATTAAACTTGGATGCTGAAAAAGTTGAGAGTGCTTCACTGCCGTACTGC GTATTGTTATATCGTCACGTATATTGTAATACCGGAGTACGACAGTAGCGTTTGCAGTGTCCAAGTCGAATGCACGGGAACCTAACATCATCATCGTTATAATT GAAAAACTCATATATATGCCAGACCATAGATGGCAGAAGCCGTACAGAATTTGAACTTAAAGTACACAAATTACAAGGAGACTTGATGTAATGTCATCACTACC TGAAATGAAAAAAATGGACTGTCCATTCCTATTTTTATCATTTCATCTGTGACATAGCCGTTGGCCGTATCTTGCTGGGTGTACACCTGAACGTGGCGGGCATG TATAAATCATTATTGTTGAAGCGCAGTTCGCTCAACTCAATAGTTGTTGGTTTCAGAGAACTACTTATTAGTGTAAATAAGTGGTAGAATTTTTTTACCGGTTAAA TTTCTTTGCCGTTATGAATTTTATTTCTCAGCGGGCAAGAAGTGAGGCCCGTGAACATTGATACCTATCTTATGCCAGAAGACAACTATGCCACTCTCCAGTAA TGATCACATATCCTACCTAGCTTTTTCGAAAACTGCTGCATCGTTTCTTGTCGGGTGATACAGTCGAGTGATCGTCACTATATTCTGTCATCTTTCAAGCCACAA CGTTGTCATTCTCTGCAAGACCTTTCTAACAATGAATAATGTCGAATGTACACTATGGATGAGGTGAATCCAAGGCTTTTGTGGTTTTGACCGAAGTGTTCCAC GAATGACGGTGACTCTAGGTCAGTACACTTGAATTATTTTGCAGAAATTTCACCGTATACAAAGCGCCTTAAAGAATTTAAAAAAGATTGCAGTGATGATATGTA TCATAGAAGTTTATTTGGTAAGCAATTGCGACAGCAATCTCCAGTGGAA | 89% | | | |
| scaffold_116 | CsT | 23217 | 7789-31005 | CsT_DRJ2R | 1132 | 29874-31005 | TACATACATGCAGTACACAATCGCCCAATTAGATCATCGACATAGAAGCTGACAAGGAATAAACCAACTCTATCAGCTTGAAGGTTTGGGTTCTTCGAATACAT GTGATTCGGTATCAAAGCAGTCAGAACGTCGAACATGTCCATGCATGAACCGTGCATTTAACTTGGATGCTGAAAAAGTTGAGAGTGCTTCACTGCCGTACTG CGTATTGTTATATCGTCACGTATATTGTGATACCGAAGTACGGCAGTAGCGTTTGCAGTGTCCACGTCGAATGCACGGGAACCTAACATCATCATCGTCATCAT TGAAAAACTCATATATATGCCAGACCATAGATGGCAGAAGCCTGTACAGAATTTGAAGTTAAAGTCCGCAAATTACAAGAAACAGACTTGACGTAATGTCATCA CTATCTGAAATGAATAAATGGACTGTCCATTGCTATTTTTATCATTTAATCTGTGACATAGCCGTTGGCCGTATCTTGCTGGGTACACCTGAACGTGGCGAGCA TGTATAAATCATTATTATTGAAACGCAGTTCGCGCAACTCAATAGTTGTTTGTTTCAGAGAACTACTTATTTATAGAGTATGATTGTGGCATTAAAAAATGCTGGT AGTGTAAATAAGTGGTAGAATTTTTTTACCGGTTAAATTTCTTTGCCGTTATGAATTTTATTTCTCAGCGGGCAAAAAGTGAGGCCCGTGAACATTGATACCTAT CTTATGCCAGAAGACAACTATGCCACTCTCCAGTAATGATCACATATCCTACCTAGCTTTTTCGAAAACTGCTGCATCGTTTTTTGTCGGGTGATACAGTCGAG TGATCGTCACTATATTCTGTCATCTTTCAAGCCACAACGTTGTCATTCTCTGCAAGACCTTTCTAACAATGAATAATGTCGAATGTACACTATGGATGAGGTGAA CCCAAGGCTTTTGTGGTTTTGACCGAAGTGTTCCACCAACGACGGCGACTCTTGGTCGTCGGTACACTTGGATTATTTTGCAGATTTTCTACCGCATACAAGGA ACCTCTAAGCATATTTGGAAAAAAGTGCAGTGACGATGGTACCGTGGAAGCTCATTCGGAAACAATTGCAACAGAAATCTACAGTGGAA | | | | |
