## Additional file 7 for "Conserved and specific genomic features of endogenous polydnaviruses revealed by whole genome sequencing of two ichneumonid wasps"

#### Additional File 7. DRJ analysis

**A.** DNA motifs found in the direct repeated sequences flanking the IV segments inserted in wasp genomes. Analysis was performed using the DNAMINDA2 webserver (http://bmbl.sdstate.edu/DMINDA2/annotate.php); the input dataset was composed of 99 DRJ sequences (right junctions of HdIV and CsIV segments). A total of 89 motifs were obtained; only those whose occurrence exceed 70% of the DRJs are reported.

| Motif | Length (nt) | Consensus logo | Consensus sequence | Nb of occurrence in DRJs (n=99 DRJ sequences) | Nb of DRJs containing at least one motif (/99) |
|---|---|---|---|---|---|
| Motif-62 | 6 | 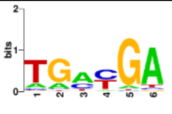 | TGAYGA | 1231 | 97 |
| Motif-56 | 6 | 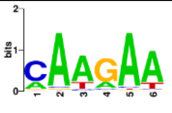 | CAAGAA | 638 | 97 |
| Motif-58 | 6 | 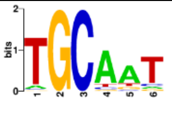 | TGCAAT | 380 | 90 |
| Motif-68 | 7 | 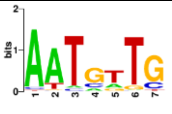 | AATGTTG | 281 | 82 |
| Motif-88 | 9 | 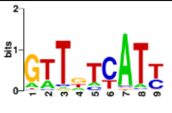 | GTTGTCATY | 205 | 76 |
| Motif-10 | 8 | 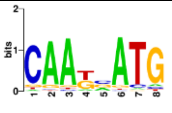 | CAATMATG | 197 | 75 |
| Motif-59 | 6 | 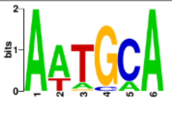 | AATGCA | 195 | 73 |

**B. Result of genome search using motifs predicted with DMINDA 2.0 webserver**

Occurrence rate of the motif in DRJ and whole genome sequences. Each motif was search among the 6 bp kmers present in the whole genome (201,969,604) and in the DRJs (33,930). The significance was evaluated using a Chi2 (taking into account the ratio of these motifs / all the other motifs in the DRJS and in the genome).

| Motif | DRJ | Whole genome | P-value |
|---|---|---|---|
| TC[G,A]TCA | 61 | 291,908 | 0.1015 |
| CAAGAA | 31 | 169,153 | 0.6959 |

#### C. Manual analysis of the regions containing an excision site

A CLUSTAL O(1.2.4) multiple sequence alignment was performed on the DRJs. The sequences used for alignment have 3 different origins: (i) DRJs from *H. didymator* genome; (ii) DRJs from *H. didymator* BAC clones; (iii) HdIV segments, e.g. PCR products (cloned in pGEM and Sanger sequenced) from encapsidated HdIV DNA template.

Color code:
DRJ1R (or right junction) underlined in dark grey; DRJ1L (or left junction) underlined in light grey
Nucleotides that differ between DRJ1R and DRJ1L are indicated in red (in blue, differences between genome and BAC sequences)
Regions in the segment sequence where potentially occurred a switch between right/left junctions are <u>underlined</u>.

##### <u>Segment Hd30</u>

Sequences used for alignment:
(i) DRJs from genome: Hd30_DRJ1R, Hd30_DRJ1L
(ii) DRJs from BAC clone # AB-06P08: right_AB, left_AB
(iii) HdIV segments: Contig_AB-20, Contig_AB-17, Contig_AB-16; plus sequence SH_AB previously sequenced

Alignment

```
left_AB             AGCCGACGTGATATATGCTGAAAACAGGTCTGACTCGCTACGAGGCTAGTTGGTTGGTCC
Hd30_DRJ1L          AGCCGACGTGATATATGCTGAAAACAGGTCTGACTCGCTACGAGGCTAGTTGGTTGGTCC
Contig_AB-17        AGTCAACATGGTGTATGTTCAGACCAAGTCTCCCTCGCTACGAGGCTAGTTGGTTGGTCC
Contig_AB-20        AGTCAACATGGTGTATGTTCAGACCAAGTCTCCCTCGCTACGAGGCTAGTTGGTTGGTCC
SH_AB               AGTCAACATGGTGTATGTTCAGACCAAGTCTCCCTCGCTACGAGGCTAGTTGGTTGGTCC
Contig_AB-16        AGTCAACATGGTGTATGTTCAGACCAAGTCTCCCTCGCTACGAGGCTAGTTGGTTGGTCC
right_AB            AGTCAACATGGTGTATGTTCAGACCAAGTCTCCCTCGCTACGAGGCTAGTTGGTTGGTCC
Hd30_DRJ1R          AGTCAACATGGTGTATGTTCAGACCAAGTCTCCCTCGCTACGAGGCTAGTTGGTTGGTCC
                    ** * ** ** * **** * * * ** ****  ***************************

left_AB             ATCAGTGCATTGCACGCTCGCTACGGGTTGGCACGCAGCATCGTGCCACAGCTGCAACTC
Hd30_DRJ1L          ATCAGTGCATTGCACGCTCGCTACGGGTTGGCACGCAGCATCGTGCCACTGCTGCAACTC
Contig_AB-17        ATCAGTGCATTGCACGCTCGCTACGGGTTGGCACGCAGCATCGTGCCACTGCTGCAACTC
Contig_AB-20        ATCCGTGCATTGCACGCTCGCTGCGGGTTGGCACCCAGCATCGTGCCACTACTGCGACTC
SH_AB               ATCCGTGCATTGCACGCTCGCTGCGGGTTGGCACCCAGCATCGTGCCACTACTGCGACTC
Contig_AB-16        ATCCGTGCATTGCACGCTCGCTGCGGGTTGGCACCCAACATCGTGCCACTACTGCGACTC
right_AB            ATCCGTGCATTGCACGCTCGCTGCGGGTTGGCACCCAGCATCGTGCCACTACTGCGACTC
Hd30_DRJ1R          ATCCGTGCATTGCACGCTCGCTGCGGGTTGGCACCCAGCATCGTGCCACTACTGCGACTC
                    *** ****************** *********** ** ***********  **** ****

left_AB             CAGCACCTTACAAACTGCAGCGATAATACACTTCGTGCTGTGGAGCACGGTGTCAACGGTGTTC
Hd30_DRJ1L          CAGCACCTTACAAACTGCAGCGATAATACACTTCGTGCTGTGGAGCACGGTGTCAACGGTGTTC
Contig_AB-17        CAACACCTTACAAACTGCAGCGATAATACACTTCGTGCTGTGGAGCACGGTGTCAACGGTGTTC
Contig_AB-20        CGGCACCTTACGAGCTGCAGCGATAAGACACTTCGTGCTGTGGAGCACGGTGTCAACGGTGTTC
SH_AB               CGGCACCTTACGAGCTGCAGCGATAAGACACTTCGTGCTGTGGAGCACGGTGTCAACGGTGTTC
Contig_AB-16        CGGCACCTTACAAACTGCAGCGATAATACACTTCGTGCTGTGGAGCACGGTGTCAACGGTGTTC
right_AB            CGGCACCTTACGAGCTGCAGCGATAAGACACTTCGTGCTGTGGAGCACGGTGTCAACGGTGTTC
Hd30_DRJ1R          CGGCACCTTACGAGCTGCAGCGATAAGACACTTCGTGCTGTGGAGCACGGTGTCAACGGTGTTC
                    *  ******** * ************ *************************************
```

##### <u>Segment Hd12</u>

Sequences used for alignment:
(i) DRJs from genome: Hd12_DRJ1R, Hd12_DRJ1L
(ii) DRJs from BAC clone # BG-42L09+BL-56G14): left_jct_BGBL1, right_jct_BGBL1
(iii) HdIV segments: Contig_BGBL1-11, Contig_BGBL1-16, Contig_BGBL1-19

Alignment

```
left_jct_BGBL1      --AGTACGGTAATTGCCGCCAGTCTTTGGCGTTATACTGTTTGCGGCTTATCTGGAGTTG
Hd12_DRJ1L          ATAGTACGATAATTGCCGCCAGTCTTTGGCGTACTACTGTTTACGGCTTATCTGGAGTTG
Contig_BGBL1-19     --AGAACGCTTATCACCACCAATCTTTGGCGTCATACTGTTTGCGGCTTATCTGCAGTTG
Contig_BGBL1-11     --AGAACGCTTATCACCACCAATCTTTGGCGTCATACTGTTTGCGGCTTATCTGCAGTTG
Contig_BGBL1-16     --AGAACGCTTATCACCACCAATCTTTGGCGTCATACTGTTTGCGGCTTATCTGCAGTTG
right_jct_BGBL1     --AGAACGCTTATCACCACCAATCTTTGGCGTCATACTGTTTGCGGCTTATCTGCAGTTG
Hd12_DRJ1R          ATAGAACGCTTATCACCACCAATCTTTGGCGTCATACTGTTTGCGGCTTATCTGCAGTTG
                      ** *** * **  ** *** **********  ******** *********** *****

left_jct_BGBL1      TCACTCATGGGTCAAGTTAAGAAAAACCTCATTTAGATGTCTCATATTCCTGTATGGTCT
Hd12_DRJ1L          TCACTCATGAGTCAAGTTAAGAAAAACCTCATTTAGATGTCTCATATTCCTGTATGGTCT
Contig_BGBL1-19     GTACTTATGGATCAAGTTGAGGAAAACCTTATCCAGGCGTCTCATACCCTTGTATGGTCT
Contig_BGBL1-11     GTACTTATGGATCAAGTTGAGGAAAACCTTATCCAGGCGTCTCATACCCTTGTATGGTCT
Contig_BGBL1-16     GTACTTATGGATCAAGTTGAGGAAAACCTTATCCAGGCGTCTCATACCCTTGTATGGTCT
right_jct_BGBL1     GTACTTATGGATCAAGTTGAGGAAAACCTTATCCAGGCGTCTCATACCCTTGTATGGTCT
Hd12_DRJ1R          GTACTTATGGATCAAGTTGAGGAAAACCTTATCCAGGCGTCTCATACCCTTGTATGGTCT
                      *** ***  ******* ** ******* **  **  ********  * **********

left_jct_BGBL1      GCTCCGTACCTAAACGTCGCGGGTACACGTACTCAGAAAGGAATGCACACCCAGGCCTTG
Hd12_DRJ1L          GCTCCGTACCTAAACGTCGCGGGTACACGTACTCAGAAAGGAATGCACACCCAGGCCTTG
Contig_BGBL1-19     GCTCCGTAACTAAACGTCGCGGGTACACGTACTCAGAAAGGAATGCACACCCAGGCCTTG
Contig_BGBL1-11     GCTCCGTACCTAAACGTCGCGGGTACACGTACTCAGAAAGGAATGCACACCCAGGCCTTG
Contig_BGBL1-16     GCTCCGTACCTAAACGTCGCGGGTACACGTACTCAGAAAGGAATGCACACCCAGGCCTTG
right_jct_BGBL1     GCTCCGTAACTAAACATCGCAGGTACACATACTCAGAGAGTAATCCACACTCAGACCTTA
Hd12_DRJ1R          GCTCCGTAACTGAACATCGCAGGTACACATACTCAGAGAGTAATCCACACTCAGACCTTA
                    ******** ** *** **** ******* ******** ** *** ***** *** ****

left_jct_BGBL1      CTCAAGACATATGATGAGCATATTCGTAAGCATAGTTTGCGATATACAACGGAGCTTTGC
Hd12_DRJ1L          CTCAAGACATATGATGAGCATATTCGTAAGCATAGTTTGCGATATACAACGGAGCTTTGC
Contig_BGBL1-19     CTCAGGACATATGATGAGCATATTCGTAAGCACAGTTTGCGATATACAACGGAGCTTTGC
Contig_BGBL1-11     CTCAAGACATATGATGAGCATATTCGTAAACATAGTTTGCGATATACAACGGAGCTTTGC
Contig_BGBL1-16     CTCAAGACATATGATGAGCATATTCGTAAGCATAGTTTGCGATATACAACGGAGCTTTGC
right_jct_BGBL1     CTCAAGACATATGATGGGCATATTCATAAGCATAGTTTGTGATATATAACGGAGCTTTGC
Hd12_DRJ1R          CTCAAGACATATGATGGGCATATTCATAAGCATAGTTTGTGATATATAACGGAGCTTTGC
                    **** *********** ******** *** ** ****** ****** *************

left_jct_BGBL1      ATCCCATGTGCACATAACACCATCAGAGATGAACGGGGAGATCTCCAATGCTATAAGGCC
Hd12_DRJ1L          ATCCCATGTGCACATAACTCCATCAGAGATGAACGGGGAGATCTCCAATGCTATAAGGCC
Contig_BGBL1-19     ATCCCATGTGCACATAACACCATCAGAGATGAACGGGGAGATCTCCAATGCTATAAGGCC
Contig_BGBL1-11     ATCCCATGTGCACATAACACCATCAGAGATGAACGGGGAGATCTCCAATGCTATAAGGCC
Contig_BGBL1-16     ATTTCATGTGCACATAACACCATCAGAGATGAACGGGGAGATCTCCAATGCTATAAGGCC
right_jct_BGBL1     ATCCCATGTGCACATAACTCTAACAGAGATAAGTAGGGAGATCTCCAATGCTATAGGGCT
Hd12_DRJ1R          ATCCCATGTGCACATAACTCTAACAGAGATAAGTAGGGAGATCTCCAATGCTATAGGGCT
                    **  ************** * * ******* *   ******************** ***

left_jct_BGBL1      CTAGTGGTTCCTACAAAGCGACGGTTTTACGTATCGGTCAACTGCTCCAGCTGTCATAGG
Hd12_DRJ1L          CTAGTGGTTCCTACAAAGCGACGGTTTTACGTATCGGTCAACTGCTCCAGCTGTCATAGG
Contig_BGBL1-19     CTAGTGGTTCCTACAAAGCGACGGTTTTACGTATCGGTCAACTGCTCCAGCTGTCATAGG
Contig_BGBL1-11     CTAGTGGTTCCTACAAAGCGACGGTTTTACGTATCGGTCAACTGCTCCAGCTGTCATAGG
Contig_BGBL1-16     CTAGTGGTTCCTACAAAGCGACGGTTTTACGTATCGGTCAACTGCTCCAGCTGTCATAGG
right_jct_BGBL1     CGAGTGGTTCCTACAGAGCGGCGCTTTTTCTTATCGGTCAACTAATAAAGCTGTCTTAGA
Hd12_DRJ1R          CGAGTGGTTCCTACAGAGCGGCGCTTTTTCTTATCGGTCAACTAATAAAGCTGTCTTAGA
                    * ************* **** ** **** * ************  *  ******* ***

left_jct_BGBL1      TGTTTGTTGTTGCTCGGTCGTGCTTGCAGTAATCCATCCTGCTCGGCAGAGA
Hd12_DRJ1L          TGTTTGTTGTTGCTCGGTCGTGCTTGCAGTAATCCATCCTGCTCGGCAGAGA
Contig_BGBL1-19     TGTTTGTTGTTGCTCGGTCGTGCTTGCAGTAATCCATCCTGCTCGGCAGAGA
Contig_BGBL1-11     TGTTTGTTGTTGCTCGGTCGTGCTTGCAGTAATCCATCCTGCTCGGCAGAGA
Contig_BGBL1-16     TGTTTGTTGTTGCTCGGTCGTGCTTGCAGTAATCCATCCTGCTCGGCAGAGA
right_jct_BGBL1     TGTTTTT--TTGCTCGGTCGTGCTCGCACAAATCCATCCTGCTCGGCAAAGA
Hd12_DRJ1R          TGTTTTTT-TTGCTCGGTCGTGCTCGCACAAATCCATCCTGCTCGGCAGAGA
                    ***** *  *************** ***  ****************** ***
```

**Segment Hd16**

Sequences used for alignment:

(i) DRJs from genome: Hd16_DRJ1R, Hd16_DRJ1L

(ii) DRJs from BAC clone # BG-42L09+BL-56G14: left_jct_BGBL2, right_jct_BGBL2

(iii) HdIV segments: Contig_BGBL2-12, Contig_BGBL2-16, Contig_BGBL2-19, SH_BGBL2

### Alignment

```
left_jct_BGBL2      ------------------------------------------------------------
Hd16_DRJ1L          ------------------------------------------------------------
Contig_BGBL2-19     ATAGAGTTGTCATGAGCGTAACTACATTTAAACTTGAAGCACTGTAAGAAGGAACGGTCT
Contig_BGBL2-12     ATAGAGTTGTCATGAGCGTAACTACATTTAAACTTGAAGCACTGTAAGAAGGAACGGTCT
Contig_BGBL2-16     ATAGAGTTGTCATGAGCGTAACTACATTTAAACTTGAAGCACTGTAAGAAGGAACGGTCT
SH_BGBL2            ATAGAGTTGTCATGAGCGTAACTACATTTAAACTTGAAGCACTGTAAGAAGGAACGGTCT
right_jct_BGBL2     ATAGAGTTGTCATGAGCGTAACTACATTTAAACTTGAAGCACTGTAAGAAGGAACGGTCT
Hd16_DRJ1R          ------TTGTCATGAGCGTAACTACATTTAAACTTGAAGCACTGTAAGAAGGAACGGTCT


left_jct_BGBL2      ------------------------------ATAAAATTGTCATGAGAG-----GAACTAA
Hd16_DRJ1L          ------------------------------------TTGTCATGAGAG-----GAACTAA
Contig_BGBL2-19     CACAGTACCTTACCTACACAAGTCTCCATCATCCTACACTAATAAGGATAAGAAAAATAA
Contig_BGBL2-12     CACAGTACCTTACCTACACAAGTCTCCATCATCCTACACTAATAAGGATAAGAAAAATAA
Contig_BGBL2-16     CACAGTACCTTACCTACACAAGTCTCCATCATCCTACACTAATAAGGATAAGAAAAATAA
SH_BGBL2            CACAGTACCTTACCTACACAAGTCTCCATCATCCTACACTAATAAGGATAAGAAAAATAA
right_jct_BGBL2     CACAGTACCTTACCTACACAAGTCTCCATCATCCTACACTAATAAGGATAAGAAAAATAA
Hd16_DRJ1R          CACAGTACCTTACCTACACAAGTCTCCATCATCCTACACTAATAAGGATAAGAAAAATAA
                                                           * ** **        ** ***

left_jct_BGBL2      ATTTAAACTTGTACCAACGTAAGAGG------AAACGCTCTCACAGTACC-TTTAGCTGC
Hd16_DRJ1L          ATTTAAACTTGTACCAACGTAAGAGG------AAACGCTCTCACAGTACC-TTTAGCTGC
Contig_BGBL2-19     AAGTAAACACATTCAACACAAACTTTTATTTCAATCTAAGGAACAGTATATGCTGCACTA
Contig_BGBL2-12     AAGTAAACACATTCAACACAAACTTTTATTTCAATCTAAGGAACAGTATATGCTGCACTA
Contig_BGBL2-16     AAGTAAACACATTCAACACAAACTTTTATTTCAATCTAAGGAACAGTATATGCTGCACTA
SH_BGBL2            AAGTAAACACATTCAACACAAACTTTTATTTCAATCTAAGGAACAGTATATGCTGCACTA
right_jct_BGBL2     AAGTAAACACATTCAACACAAACTTTTATTTCAATCTAAGGAACAGTATATGCTGCACTA
Hd16_DRJ1R          AAGTAAACACATTCAACACAAACTTTTATTTCAATCTAAGGAACAGTATATGCTGCACTA
                    *  *****   * * *    **          ** *      ******     *

left_jct_BGBL2      AAGCTTTAGATTAGATTCAACTGCACCATGGCCGGAACTGTTGCGAGCCGAGTCAAGCAA
Hd16_DRJ1L          AAGCTTTAGATTAGATTCAACTGCACCATGGCCGGAACTGTTGCGAGCCGAGTCAAGCAA
Contig_BGBL2-19     TTAACACTACTCAGATTAAGCTGCACCATGGCCGGAACTGTTGCGAGCCGAGTCAAGCAA
Contig_BGBL2-12     TTAACACTACTCAGATTAAGCTGCACCATGGCCGGAACTGTTGCGAGCCGAGTCAAGCAA
Contig_BGBL2-16     TTAACACTACTCAGATTAAGCTGCACCATGGCCGGAACTGTTGCGAGCCGAGTCAAGCAA
SH_BGBL2            TTAACACTACTCAGATTAAGCTGCACCATGGCCGGAACTGTTGCGAGCCGAGTCAAGCAA
right_jct_BGBL2     TTAACACTACTCAGATTAAGCTGCACCATGGCCGGAACTGTTGCGAGCCGAGTCAAGCAA
Hd16_DRJ1R          TTAACACTACTCAGATTAAGCTGCACCATGGCCGGAACTGTTGCGAGCCGAGTCAAGCAA
                              * ***** * ****************************************

left_jct_BGBL2      CACTTCTGTGTCCGTACGTTCAGCCATAGAATAATATTCCAAGACGTCTGGATTGTTTTG
Hd16_DRJ1L          CACTTCTGTGTCCGTACGTTCAGCCATAGAATAATATTCCAAGACGTCTGGATTGTTTTG
Contig_BGBL2-19     CACTTCTGTGTCCGTACGTTCAGCCATAGAATAATATTCCAAGACGTCTGGATTGTTTTG
Contig_BGBL2-12     CACTTGTGTGTCCGTACGTTCAGTCATGGAATAATATTCCAAGACGTCTGGATTGTTTTG
Contig_BGBL2-16     CACTTGTGTGTCCGTACGTTCAGTCATGGAATAATATTCCAAGACGTCTGGATTGTTTTG
SH_BGBL2            CACTTGTGTGTCCGTACGTTCAGCCATGGAATAATATTCCAAGACGTCTGGATTGTTTTG
right_jct_BGBL2     CACTTGTGTGTCCGTACGTTCAGCCATGGAATAATATTCCAAGACGTCTGGATTGTTTTG
Hd16_DRJ1R          CACTTGTGTGTCCGTACGTTCAGCCATGGAATAATATTCCAAGACGTCTGGATTGTTTTG
                    ***** ***************** *** ********************************

left_jct_BGBL2      ATGCACATTTGCAATGATCTCAGTGAACATCTCTTTAGATTCTTTAAGAAGAATCGCGGG
Hd16_DRJ1L          ATGCACATTTGCAATGATCTCAGTGAACATCTCTTTAGATTCTTTAAGAAGAATCGCGGG
Contig_BGBL2-19     ATGCACATTTGCAATGATCTCAGTGAACATCTCTTTAGATTCTTTAAGAAGAATCGCGGG
Contig_BGBL2-12     ATGCACATTTGCAATGATCTCAGTGAACATCTCTTTAGATTCTTTAAGAAGAATCGCGGG
Contig_BGBL2-16     ATGCACATTTGCAATGATCTCAGTGAACATCTCTTTAGATTCTTTAAGAAGAATCGCGGG
SH_BGBL2            ATGCACATTTGCAATGATCTCAGTGAACATCTCTTTAGATTCTTTAAGAAGAATCGCGGG
right_jct_BGBL2     ATGCACATTTGCAATGATCTCAGTGAACATCTCTTTAGATTCTTTAAGAAGAATCGCGGG
Hd16_DRJ1R          ATGCACATTTGCAATGATCTCAGTGAACATCTCTTTAGATTCTTTAAGAAGAATCGCGGG
                    ************************************************************

left_jct_BGBL2      GATCATATATTTAGTCAACCATGCGGAGACGTGCGCCGTACAGAAATGATGCCAGTGGAG
Hd16_DRJ1L          GATCATATATTTAGTCAACCATGCGGAGACGTGCGCCGTACAGAAATGATGCCAGTGGAG
Contig_BGBL2-19     GATCATATATTTAGTCAACCATGCGGAGACGTGCGCCGTACAGAAATGATGCCAGTGGAG
Contig_BGBL2-12     GATCATATATTTAGTCAACCATGCGGAGACGTGCGCCGTACAGAAATGATGCCAGTGGAG
Contig_BGBL2-16     GATCATATATTTAGTCAACCATGCGGAGACGTGCGCCGTACAGAAATGATGCCAGTGGAG
SH_BGBL2            GATCATATATTTAGTCAACCATGCGGAGACGTGCGCCGTACAGAAATGATGCCAGTGGAG
right_jct_BGBL2     GATCATATATTTAGTCAACCATGCGGAGACGTGCGCCGTACAGAAATGATGCCAGTGGAG
Hd16_DRJ1R          GATCATATATTTAGTCAACCATGCGGAGACGTGCGCCGTACAGAAATGATGCCAGTGGAG
                    ************************************************************

left_jct_BGBL2      TTCGTTCTTACACTCACGTCCAGGTGAAACCACTTTATTGGGTTTTTTCTCGCAGGAGCA
Hd16_DRJ1L          TTCGTTCTTACACTCACGTCCAGGTGAAACCACTTTATTGGGTTTTTTCTCGCAGGAGCA
Contig_BGBL2-19     TTCGTTCTTACACTCACGTCCAGGTGAAACCACTTTATTGGGTTTTTTCTCGCAGGAGCA
Contig_BGBL2-12     TTCGTTCTTGCACTCACGTCCAGGTGAAACCACTTTATTGGGTTTTTTCTCGCAGGAGCA
```

```
Contig_BGBL2-16     TTCGTTCTTGCACTCACGTCCAGGTGAAACCACTTTATTGGGTTTTTTCTCGCAGGAGCA
SH_BGBL2            TTCGTTCTTGCACTCACGTCCAGGTGAAACCACTTTATTGGGTTTTTTCTCGCAGGAGCA
right_jct_BGBL2     TTCGTTCTTGCACTCACGTCCAGGTGAAACCACTTTATTGGGTTTTTTCTCGCAGGAGCA
Hd16_DRJ1R          TTCGTTCTTGCACTCACGTCCAGGTGAAACCACTTTATTGGGTTTTTTCTCGCAGGAGCA
                    ********* **************************************************

left_jct_BGBL2      GGTGTCGCTGGCGACCGCCTTGCACTTCTGGAAGCGAATCTTCTTCACGAACTGTAGTAT
Hd16_DRJ1L          GGTGTCGCTGGCGACCGCCTTGCACTTCTGGAAGCGAATCTTCTTCACGAACTGTAGTAT
Contig_BGBL2-19     GGTGTCGCTGGCGACCGCCTTGCACTTCTGGAAGCGAATCTTCTTCACGAACTGTAGTAT
Contig_BGBL2-12     GGTGTCGCTGGCGACCGCCTTGCACTTCTGGAAGCGAATGTTCTTCACGAACTGTAGTAT
Contig_BGBL2-16     GGTGTCGCTGGCGACCGCCTTGCACTTCTGGAAGCGAATGTTCTTCACGAACTGTAGTAT
SH_BGBL2            GGTGTCGCTGGCGACCGCCTTGCACTTCTGGAAGCGAATGTTCTTCACGAACTGTAGTAT
right_jct_BGBL2     GGTGTCGCTGGCGACCGCCTTGCACTTCTGGAAGCGAATGTTCTTCACGAACTGTAGTAT
Hd16_DRJ1R          GGTGTCGCTGGCGACCGCCTTGCACTTCTGGAAGCGAATGTTCTTCACGAACTGTAGTAT
                    *************************************** ********************

left_jct_BGBL2      TACGGGCAGCCTCACAAAATCTCTCTCACCCGGCGGCCGCTCCCAAAATAGAGGCATGAG
Hd16_DRJ1L          TACGGGCAGCCTCACAAAATCTCTCTCACCCGGCGGCCGCTCCCAAAATAGAGGCATGAG
Contig_BGBL2-19     TACGGGCAGCCTCACAAAATCTCTCTCACCCGGCGGCCGCTCCCAAAATAGAGGCATGAG
Contig_BGBL2-12     TACGGGCAGCCTCACAAAATCTCTCTCACCCGGCGGCCGCTCCCAAAATAGAGGCATGAG
Contig_BGBL2-16     TATGGGCAGCCTCACAAAATCTCTCTCACCCGGCGGCCGCTCCCAAAATAGAGGCGTGAG
SH_BGBL2            TACGGGCAGCCTCACAAAATCTCTCTCACCCGGCGGCCGCTCCCAAAATAGAGGCGTGAG
right_jct_BGBL2     TACGGGCAGCCTCACAAAATCTCTCTCACCCGGCGGCCGCTCCCAAAATAGAGGCGTGAG
Hd16_DRJ1R          TACGGGCAGCCTCACAAAATCTCTCTCACCCGGCGGCCGCTCCCAAAATAGAGGCGTGAG
                    ** **************************************************** ****

left_jct_BGBL2      CCAGTCCCAATTTATCAGCAGGCGATCTTCTTCTGGCCTTGTTGCATCGAAGCTGTAGCG
Hd16_DRJ1L          CCAGTCCCAATTTATCAGCAGGCGATCTTCTTCTGGCCTTGTTGCATCGAAGCTGTAGCG
Contig_BGBL2-19     CCAGTCCCAATTTATCAGCAGGCGATCTTCTCCTGGCCTTGTTGCATCGAAGCTGTAGCG
Contig_BGBL2-12     CCAGTCCCAATTTATCAGCAGGCGATCTTCTTCTGGCCTTGTTGCATCGAAGCTGTAGCG
Contig_BGBL2-16     CCAGTCCCAATTTATCAGCAGGCGATCTTCTTCTGGCCTTGTTGCATCGAAGCTGTAGCG
SH_BGBL2            CCAGTCCCAATTTATCAGCAGGCGATCTTCTTCTGGCCTTGTTGCATCGAAGCTGTAGCG
right_jct_BGBL2     CCAGTCCCAATTTATCAGCAGGCGATCTTCTTCTGGCCTTGTTGCATCGAAGCTGTAGCG
Hd16_DRJ1R          CCAGTCCCAATTTATCAGCAGGCGATCTTCTTCTGGCCTTGTTGCATCGAAGCTGTAGCG
                    ******************************* ****************************

left_jct_BGBL2      GACGGGGCATGCCTTCCCGTTCAGGAAAGTCAAGTCCAGGTTTCGGATCGATGTAAGAAA
Hd16_DRJ1L          GACGGGGCATGCCTTCCCGTTCAGGAAAGTCAAGTCCAGGTTTCGGATCGATGTAAGAAA
Contig_BGBL2-19     GACGGGGCATGCCTTCCCGTTCAGGAAAGTCAAGTCCAGGTTTCGGATCGATGTAAGAAA
Contig_BGBL2-12     GACGGGGCATGCCTTCCCGTTCAGGAAAGTCAAGTCCAGGTTTCGGATCGATGTAAGAAA
Contig_BGBL2-16     GACGGGGCATGCCTTCCCGTTCAGGAAAGTCAAGTCCAGGTTTCGGATCGATGTAAGAAA
SH_BGBL2            GACGGGGCATGCCTTCCCGTTCAGGAAAGTCAAGTCCAGGTTTCGGATCGATGTAAGAAA
right_jct_BGBL2     GACGGGGCATGCCTTCCCGTTCAGGAAAGTCAAGTCCAGGTTTCGGATCGATGTAAGAAA
Hd16_DRJ1R          GACGGGGCATGCCTTCCCGTTCAGGAAAGTCAAGTCCAGGTTTCGGATCGATGTAAGAAA
                    ************************************************************

left_jct_BGBL2      CTGTTGATATGCTACGTTGAAACGTATAAATTCGATGAAGCTGAAAGGGGCGCCATTCGG
Hd16_DRJ1L          CTGTTGATATGCTACGTTGAAACGTATAAATTCGATGAAGCTGAAAGGGGCGCCATTCGG
Contig_BGBL2-19     CTGTTGATATGCTACGTTGAAACGTATAAATTCGATGAAGCTGAAAGGGGCGCCATTCGG
Contig_BGBL2-12     CTGTTGATATGCTACGTTGAAACGTATAAATTCGATGAAGCTGAAAGGGGCGCCATTCGG
Contig_BGBL2-16     CTGTTGATATGCTACGTTGAAACGTATAAATTCGATGAAGCTGAAAGGGGCGCCATTCGG
SH_BGBL2            CTGTTGATATGCTACGTTGAAACGTATAAATTCGATGAAGCTGAAAGGGGCGCCATTCGG
right_jct_BGBL2     CTGTTGATATGCTACGTTGAAACGTATAAATTCGGCGAAGTTGAAAGGGGCGCCATTCGG
Hd16_DRJ1R          CTGTTGATATGCTACGTTGAAACGTATAAATTCGGCGAAGTTGAAAGGGGCGCCATTCGG
                    **********************************  **** *******************

left_jct_BGBL2      CATGGCAGGCCCGCGGAACGTGAAGGTGTCCACCGGGTAAGTATGTGTTCTGTCGTGATC
Hd16_DRJ1L          CATGGCAGGCCCGCGGAACGTGAAGGTGTCCACCGGGTAAGTATGTGTTCTGTCGTGATC
Contig_BGBL2-19     CATGGCAGGCCCGCGGAACGTGAAGGTGTCCACCGGGTAAGTATGTGTTCTGTCGTGATC
Contig_BGBL2-12     CATGGCAGGCCCGCGGAACGTGAAGGTGTCCACCGGGTAAGTATGTGTTCTGTCGTGATC
Contig_BGBL2-16     CATGGCAGGCCCGCGGAACGTGAAGGTGTCCACCGGGTAAGTATGTGTTCTGTCGTGATC
SH_BGBL2            CATGGCAGGCCCGCGGAACGTGAAGGTGTCCACCGGGTAAGTATGTGTTCTGTCGTGATC
right_jct_BGBL2     CATGGCAGGCCCGCGGGACGTGAAGGTGTCCACCGGGTAAGTAAGTGTTCTGTCGTGATC
Hd16_DRJ1R          CATGGCAGGCCCGCGGGACGTGAAGGTGTCCACCGGGTAAGTAAGTGTTCTGTCGTGATC
                    **************** ************************** ****************

left_jct_BGBL2      TTCGTTGAACGGCATGTTGAGTTCTGGATATCACCTGCGTCTGGACGATGTTCCTTTGTG
Hd16_DRJ1L          TTCGTTGAACGGCATGTTGAGTTCTGGATATCACCTGCGTCTGGACGATGTTCCTTTGTG
Contig_BGBL2-19     TTCGTTGAACGGCATGTTGAGTTCTGGATATCACCTGCGTCTGGACGATGTTCCTTTGTG
Contig_BGBL2-12     TTCGTTGAACGGCATGTTGAGTTCTGGATATCACCTGCGTCTGGACGATGTTCCTTTGTG
Contig_BGBL2-16     TTCGTTGAACGGCATGTTGAGTTCTGGATATCACCTGCGTCTGGACGATGTTCCTTTGTG
SH_BGBL2            TTCGTTGAACGGCATGTTGAGTTCTGGATATCACCTGCGTCTGGACGATGTTCCTTTGTG
right_jct_BGBL2     TTCGTGGAACGGCATGTTGAGTTCTGGATATCACCTGCGTCTGGATGATGTTCCTTCGTG
Hd16_DRJ1R          TTCGTGGAACGGCATGTTGAGTTCTGGATATCACCTGCGTCTGGATGATGTTCCTTCGTG
                    ***** *************************************** ********** ***

left_jct_BGBL2      AATGTTCACGGTTGAGTGTCTAGCCGTCGACTGCGCGCAAATCCATTTATACCACTCCAA
```

```
Hd16_DRJ1L          AATGTTCACGGTTGAGTGTCTAGCCGTCGACTGCGCGCAAATCCATTTATACCACTCCAA
Contig_BGBL2-19     AATGTTCACGGTTGAGTGTCTAGCCGTCGACTGCGCGCAAATCCATTTATACCACTCCAA
Contig_BGBL2-12     AATGTTCACGGTTGAGTGTCTAGCCGTCGACTGCGCGCAAATCCATTTATACCACTCCAA
Contig_BGBL2-16     AATGTTCACGGTTGAGTGTCTAGCCGTCGACTGCGCGCAAATCCATTTATACCACTCCAA
SH_BGBL2            AATGTTCACGGTTGAGTGTCTAGCCGTCGACTGCGCGCAAATCCATTTATACCACTCCAA
right_jct_BGBL2     AATGTTCACGGTTGAGTGTCTAGCCGTCGACTGCGCGCAAATCCATTTATACCACTCCAG
Hd16_DRJ1R          AATGTTCACGGTTGAGTGTCTAGCCGTCGACTGCGCGCAAATCCATTTATACCACTCCAG
                    *********************************************************** 
```

```
left_jct_BGBL2      TAGCTTATCAT
Hd16_DRJ1L          TAGCTTATCAT
Contig_BGBL2-19     TAGCTTATCAT
Contig_BGBL2-12     TAGCTTATCAT
Contig_BGBL2-16     TAGCTTATCAT
SH_BGBL2            TAGCTTATCAT
right_jct_BGBL2     TAATTTATCAT
Hd16_DRJ1R          TAATTTATCAT
                    **  *******
```

#### **Segment Hd29 (present in BAC clone # BR-08O01)**

Sequences used for alignment:

(i) DRJs from genome: Hd29_DRJ1R, Hd29_DRJ1L

(ii) DRJs from BAC clone # BR-08O01: left_jct_BR, right_jct_BR

(iii) HdIV segments: Contig_BR-1, Contig_BR-4, Contig_BR-7, Contig_BR-10, SH_BR

Alignment

```
left_jct_BR         -----------------------------------TAGCGATCGCAGTGCGTCGCTGCAT
Hd29_DRJ1L          ATTTCGTGGCAGGCTTAGTT-TTGATGGAGGGACGTAGCGATCGCAGTGCGTCGCTGCAT
Contig_BR-10        -----------------------------------TGGCCACCGCTGTGCGTTGTTGAAC
Contig_BR-1         -----------------------------------TGGCCACCGCTGTGCGTTGTTGAAC
Contig_BR-7         -----------------------------------TGGCCACCGCTGTGCGTTGTTGAAC
SH_BR               -----------------------------------TGGCCACCGCTGTGCGTTGTTGAAC
Contig_BR-4         -----------------------------------TGGCCACCGCTGTGCGTTGTTGAAC
right_jct_BR        -----------------------------------TGGCCACCGCTGTGCGTTGTTGAAC
Hd29_DRJ1R          ATTTCGTGGCAAGCGACTTTTCATCGTAGAACGGATGGCCACCGCTGTGCGTTGTTGAAC
                                                       * ** * *** ****** * ** * 
```

```
left_jct_BR         CAATGATGCAGCTTTGTGTTCTTCTCGAACGGTGAACATTCGTGCAACCAACCTGACACG
Hd29_DRJ1L          CAATGATGCAGCTTTGTGTCCTTCTCGAACGGTGAACATTCGTGCAACCAACCTGACACG
Contig_BR-10        CAATGATCCAACTTTGTGTTCTTCAGGAACGGTGAACATTCGTGCAATCAACTTGACACG
Contig_BR-1         CAATGATCCAACTTTGTGTTCTTCAGGAACGGTGAACATTCGTGCAATCAACTTGACACG
Contig_BR-7         CAATGATCCAACTTTGTGTTCTTCAGGAACGGTGAACATTCGTGCAATCAACTTGACACG
SH_BR               CAATGATCCAACTTTGTGTTCTTCAGGAACGGTGAACATTCGTGCAATCAACTTGACACG
Contig_BR-4         CAATGATCCAACTTTGTGTTCTTCAGGAACGGTGAACATTCGTGCAATCAACTTGACACG
right_jct_BR        CAATGATCCAACTTTGTGTTCTTCAGGAACGGTGAACATTCGTGCAATCAACTTGACACG
Hd29_DRJ1R          CAATGATCCAACTTTGTGTTCTTCAGGAACGGTGAACATTCGTGCAATCAACTTGACACG
                    ******* ** ******** ****  ********************* **** *******
```

```
left_jct_BR         TCGCCATTCGGAGAGAGGTTCGATAGACCATGAGGAATATCCAAACGTCATTCATTGTGC
Hd29_DRJ1L          TCGCCATTCGGAGAGAGGTTCGATAGACCATGAGGAATATCCAAACGTCATTCATTGTGC
Contig_BR-10        TCGTCATGCGGCAAGAAATTCGATAGACCATGAGGAATATCGTAACGTCATTCATCGTGC
Contig_BR-1         TCGTCATGCGGCAAGAAATTCGATAGACCATGAGGAATATCGTAACGTCATTCATCGTGC
Contig_BR-7         TCGTCATGCGGCAAGAAATTCGATAGACCATGAGGAATATCGTAACGTCATTCATCGTGC
SH_BR               TCGTCATGCGGCAAGAAATTCGATAGACCATGAGGAATATCGTAACGTCATTCATCGTGC
Contig_BR-4         TCGTCATGCGGCAAGAAATTCGATAGACCATGAGGAATATCGTAACGTCATTCATCGTGC
right_jct_BR        TCGTCATGCGGCAAGAAATTCGATAGACCATGAGGAATATCGTAACGTCATTCATCGTGC
Hd29_DRJ1R          TCGTCATGCGGCAAGAAACTCGATAGACCATGAGGAATATCGTAACGTCATTCATCGTGC
                    *** *** ***  ***   **********************  ************ ****
```

```
left_jct_BR         GCACT--TCTAGGCTTTTGCAAT---CGAATCATTGCAGATACGCATTTTGAAGCAACGT
Hd29_DRJ1L          GCACT--TCTAGGCTATTGCAAT---CGAATCATTGCAGATACGCATTTTGAAGCAACGC
Contig_BR-10        ACGCATTCCTAGGACAAGTCTCTGCGCTCGAATCGTTGCGGATGCATTTGGAAGCAACGT
Contig_BR-1         ACGCATTCCTAGGACAAGTCTCTGCGCTCGAATCGTTGCGGATGCATTTGGAAGCAATGT
Contig_BR-7         ACGCATTCCTAGGACAAGTCTCTGCGCTCGAATCGTTGCGGATGCATTTGGAAGCAATGT
SH_BR               ACGCATTCCTAGGACAAGTCTCTGCGCTCGAATCGTTGCGGATGCATTTGGAAGCAATGT
Contig_BR-4         ACGCATTCCTAGGACAAGTCTCTGCGCTCGAATCGTTGCGGATGCATTTGGAAGCAATGT
right_jct_BR        ACGCATTCCTAGGACAAGTCTCTGCGCTCGAATCGTTGCGGATGCATTTGGAAGCAATGT
Hd29_DRJ1R          ACGCATTCCTAGGACAAGTCTCTGCGCTCGAATCGTTGCGGATGCATTTGGAAGCAATGT
                     * * *              *     *              * ****** ******* * 
```

```
left_jct_BR         GGTTCAAATGATGTTGGGGAAAAGTGTGCAAACATTTATTAGATTGACCGAATGTTCATT
```

```
Hd29_DRJ1L        GGTTCAAATGATGTTGGGGAAAAGTGTGCAAACATTTATTAGATTGACCGAATGTTCATT
Contig_BR-10      GGTTCAAATGATGTTGGGGAAAAGTGTGCAAACATTTATTAGATTGACCGAATGTTCATT
Contig_BR-1       GTCTCCAATTATGTTGGGAAAAAGTGTCCAAAAATTTATTAGATTGACCGAATGTTAATT
Contig_BR-7       GTCTCCAATTATGTTGGGAAAAAGTGTCCAAAAATTTATTAGATTGACCGAATGTTAATT
SH_BR             GTCTCCAATTATGTTGGGAAAAAGTGTCCAAAAATTTATTAGATTGACCGAATGTTAATT
Contig_BR-4       GTCTCCAATTATGTTGGGAAAAAGTGTCCAAAAATTTATTAGATTGACCGAATGTTAATT
right_jct_BR      GTCTCCAATTATGTTGGGAAAAAGTGTCCAAAAATTTATTAGATTGACCGAATGTTAATT
Hd29_DRJ1R        GTCTCCAATTATGTTGGGAAAAAGTGTCCAAAAATTTATTAGATTGACCGAATGTTAATT
                  *  ** *** ******** ******** **** *********************** ***
```

```
left_jct_BR       TGAACCTCTTGAAAAACTCACAGCTCGAGTTTAAGGAGAGTTTGACGGGATAGACGTGAC
Hd29_DRJ1L        TGAACCTCTTGAAAAACTCACAGCTCGAGGTTG---AGAGTTTGACGGGATAGACGTGAC
Contig_BR-10      TGAACCTCTTGAAAAACTCACAGCTCGAGTTTAAGGAGAGTTTGACGGGATAGACGTGAC
Contig_BR-1       TGGAGCTCTTGAGAAATTCACAGCTCGAGGTTAACGAGAGTTTGAAGGGATAAGCGTCAC
Contig_BR-7       TGGAGCTCTTGAGAAATTCACAGCTCGAGGTTAACGAGAGTTTGAAGGGATAAGCGTCAC
SH_BR             TGGAGCTCTTGAGAAATTCACAGCTCGAGGTTAACGAGAGTTTGAAGGGATAAGCGTCAC
Contig_BR-4       TGGAGCTCTTGAGAAATTCACAGCTCGAGGTTAACGAGAGTTTGAAGGGATAAGCGTCAC
right_jct_BR      TGGAGCTCTTGAGAAATTCACAGCTCGAGGTTAACGAGAGTTTGAAGGGATAAGCGTCAC
Hd29_DRJ1R        TGAAGCTCTTGAGAAATTCACAGCTCGAGGTTAACGAGAGTGTGAAGGGATAAGCGTCAC
                  ** * ******* *** ************ **    ***** *** ******  *** **
```

```
left_jct_BR       GTTCTTCATGGAAGGTGTGGCTCGCAAACGTCGCATGCAGCTTGCGAGTTCCAAATGAAA
Hd29_DRJ1L        GTTCTTCATGGAAGGTGTGGCTCGCAAACGTCGCATGCAGCTTGCGAGTTCCAAATGAAA
Contig_BR-10      GTTCTTCATGGAAGGTGTGGCTCGCAAACGTCGCATGCAGCTTGCGAGTTCCAAATGAAA
Contig_BR-1       GTTCTTCATGGGCGGTGCGGCTCGCACACGTCGCATGGAGCTTGCGAGTTCCAAATGAAA
Contig_BR-7       GTTCTTCATGGACGGTGCGGCTCGCACACGTCGCATGCAGCTTGCGAGTTCCAAATGAAA
SH_BR             GTTCTTCATGGACGGTGCGGCTCGCAAACGTCGCATGCAGCTTGCGAGTTCCAAATGAAA
Contig_BR-4       GTTCTTCATGGACGGTGCGGCTCGCACACGTCGCATGGAGCTTGCGAGTTCCAAATAAAA
right_jct_BR      GTTCTTCATGGACGGTGCGGCTCGCACACGTCGCATGGAGCTTGCGAGTTCCAAATAAAA
Hd29_DRJ1R        GTTCTTCATGGACGGTGCGGCTCGCACACGTCGCATGGAGCTTGCGAGTTCCAAATAAAA
                  ***********  **** ******** ********** ****************** ***
```

```
left_jct_BR       ACAACTGCGCGGGGTATTGTTTATTTCTGGAGAACACCG
Hd29_DRJ1L        ACAACTGCGCGGGGTATTGTTTATTTCTGGAGAACACCG
Contig_BR-10      ACAACTGCGCGGGGTATTGTTTATTTCTGGAGAACACCG
Contig_BR-1       ACAACTGCGCGGGGTATTGTTTATTTCTGGAGAACACCG
Contig_BR-7       ACAACTGCGCGGGGTATTGTTTATTTCTGGAGAACACCG
SH_BR             ACAACTGCGCGGGGTATTGTTTATTTCTGGAGAACACCG
Contig_BR-4       C-AACTGCGTGGGGTATTGTTTATCTCTTGAGAGCACCG no switch ????
right_jct_BR      C-AACTGCGTGGGGTATTGTTTATTTCTTGAGAGCACCG
Hd29_DRJ1R        C-AACTGCGTGGGGTATTGTTTATTTCTTGAGAGCACCG
                    ******* ************** *** **** *****
```

#### Segment Hd22

Sequences used for alignment:

(i) DRJs from genome: Hd22_DRJ1R, Hd22_DRJ1L

(ii) DRJs from BAC clone # CN-29M18: left_junction, right_junction

(iii) HdIV segments: Contig_CN-31, Contig_CN-33, Contig_CN-36, Contig_CN-39

Alignment

```
left_junction       CAATAGGGTGAATTCAAACGCTGGTCCAAATGGCAGGAGCTTGGCAGGACAGATGACATC
Hd22_DRJ1L          ----------AATTCAAACGCTGGTCCCAATGGCAGGAGCTTGGCAGGACAGATGACATC
Contig_CN-31        CAATAGGGTGAATTCAAACGCTGGTCCAAATGGCAGGAGCTTGGCAGGACAGATGACATC
Contig_CN-33        CAATAGGGTGAATTCAAACGCTGGTCCAAATGGCAGGAGCTTGGCAGGACAGATGACATC
Contig_CN-39        CAATAGGGTGAATTCAAACGCTGGTCCAAATGGCAGGAGCTTGGCAGGACAGATGACATC
Contig_CN-36        CAATAGGGTGAATTCAAACGCTGGTCCAAATGGCAGGAGCTTGACAAGGCAGATGATATC
right_junction      CATTGAGAAAAACTCAAGCGCCGATAGCAATGGCAGGAGCTTGACAAGGCAGATGATATC
Hd22_DRJ1R          ----------AACTCAAGCGCCGATAGCAATGGCAGGAGCTTGACAAGGCAGATGATATC
                              ** **** *** * *   *************** ** * ******* ***
```

```
left_junction       GAGTGAATGCGTAGGAGTCTGAAACAGCGCGTCGTGAACAATATCGTGACCATTGATCGC
Hd22_DRJ1L          GAGTGAATGCGTAGGAGTCTGAAACAGCGCGTCGTGAACAATATCGTGACCATTGATCGC
Contig_CN-31        GAGTGAATGCGTAGGAGTCTGAAACAGCGCGTCGTGAACAATATCGTGACCATTGATCGC
Contig_CN-33        GAGTGAATGCGTAGGAGTCTGAAACAGCGCGTCGTGAACAATATCGTGACCATTGATCGC
Contig_CN-39        GAGTGAATGCGTAGGAGTCTGAAACAGCGCGTCGTGAACAATATCGTGACCATTGATCGC
Contig_CN-36        GAGTGAGTGCGTAGGTCTCTGGAACCGCGGGTCGTGAACAATATCGTGACCATTGATCGC
right_junction      GAGTGAGTGCGTAGGTCTCTGGAACCGCGGGTCGTGAACAATATCGTGACCATTGATCGC
Hd22_DRJ1R          GAGTGAGTGCGTAGGTCTCTGGAACCGCGGGTCGTGAACAATATCGTGACCATTGATCGC
                    ****** ********  **** *** *** ******************************
```

```
left_junction       CATAGCGACTTGCCGCATCTCAGGATCGTGATCTAAAA--------CGTTTCACGTGA
Hd22_DRJ1L          CATAGCGACTTGCCGCATCTCAGGATCGTGATCTAAAA--------CGTTTCACGTGA
Contig_CN-31        CACAGCGACTTGGCGACTTGGCACACGTCAGCGTCAAGTTCCACAAAATCTGACGCGA
Contig_CN-33        CACAGCGACTTGGCGACTTGGCACACGTCAGCGTCAAGTTCCACAAAATCTGACGCGA
Contig_CN-39        CATAGCGACTTGGCGACTTGGCACACGTCAGCGTCAAGTTCCACAAAATCTGACGCGA
Contig_CN-36        CACAGCGACTTGGCGACTTGGCACACGTCAGCGTCAAGTTCCACAAAATCTGACGCGA
right_junction      CACAGCGACTTGGCGACTTGGCACACGTCAGCGTCAAGTTCCACAAAATCTGACGCGA
Hd22_DRJ1R          CACAGCGACTTGGCGACTTGGCACACGTCAGCGTCAAGTTCCACAAAATCTGACGCGA
                    ** ********* **  *      *        * **           * * *** **
```

#### **Segment Hd19**

Sequences used for alignment:
(i) DRJs from genome: Hd19_DRJ1R, Hd19_DRJ1L
(ii) DRJs from BAC clone # CP-38F08+CQ-40J21: CPCQ_right, CPCQ_left
(iii) HdIV segments: jct_CPCQ-28, jct_CPCQ-27, jct_CPCQ-23, jct_CPCQ-22

Alignment

```
CPCQ_left           TTTTTGCTGGTTCATGTTTCAGAGCAGCCGGTCTATGTGATACACGCTGCTTAGCCTGGT
Hd19_DRJ1L          TTTTTGCTGGTTCATGTTTCAGAGCAGCCGGTCTATGTGATACACGCTGCTTAGCCTGGT
jct_CPCQ-28         TTTTTGCTGCTTCATGTCTCAGAGCAGCTTGTCTACATGATACACGCTGATTTGTC-TGT
jct_CPCQ-27         TTTTTGCTGCTTCATGTCTCAGAGCAGCTTGTCTACATGATACACGCTGATTTGTC-TGT
jct_CPCQ-23         TTTTTGCTGCTTCATGTCTCAGAGCAGCTTGTCTACATGATACACGCTGATTTGTC-TGT
jct_CPCQ-22         TTTTTGCTGCTTCATGTCTCAGAGCAGCTTGTCTACATGATACACGCTGATTTGTC-TGT
CPCQ_right          TTTTTGCTGCTTCATGTCTCAGAGCAGCTTGTCTACATGATACACGCTGATTTGTC-TGT
Hd19_DRJ1R          TTTTTGCTGCTTCATGTCTCAGAGCAGCTTGTCTACATGATACACGCTGATTTGTC-TGT
                    ********* ******* **********  *****  ************ ** * *  **
```

```
CPCQ_left           GACGGCACGATGGTTCCCCTCATGCTGCACGAGCTCTGTGAGCTTT
Hd19_DRJ1L          GACGGCACGATGGTTCCCCTCATGCTGCACGAGCTCTGTGAGCTTT
jct_CPCQ-28         GACGGCACGATGGTTCCCCTCATGCTGCACGAGCTCTGTGAGCTTT
jct_CPCQ-27         GACGGCACGATGGTTCCCCTCATGCTGCACGAGCTCTGTGAGCTTT
jct_CPCQ-23         GACGGCACGATGGTTCCCCTCATGCTGCACGAGCTCTGTGAGCTTT
jct_CPCQ-22         GACGGCACGATAATTCGCCTCCTGCTGCACGAGCTCTGTGAGCTTT
CPCQ_right          GACGGCACGATAATTCGCCTCCTGCTGCACAAGCTTTGTGAGCTTT
Hd19_DRJ1R          GACGGCACGATAATTCGCCTCATGCTGCACAAGCTTTGTGAGCTTT
                    ***********  *** **** ******** **** **********
```

#### **Segments Hd28**

Sequences used for alignment:
(i) DRJs from genome: Hd28_DRJ1R, Hd28_DRJ1L, Hd24_DRJ1R (reverse), Hd24_DRJ1L (reverse)
(ii) DRJs from BAC clone # CR-47M17: CR_left, CR_right
(iii) HdIV segments: Contig_CR-05, Contig_CR-06, Contig_CR-09, Contig_CR-10

Alignment

```
CR_left             GCCTTTGCAGCTGAGCT-GCATGAGGTTTTCATGTCCTCGATAACTGGATTCCATTGTCA
Hd28_DRJ1L          GCCTTTGCAGCTGAGCT-GCATGAGGTTTTCATGTCCTCGATAACTGGATTCCATTGTCA
Contig_CR-05        GCCTTTGGAGCTAAACTTGCACGAGATTTCGGTGTCCGTGATAACTGAATGGCATTGTCA
Contig_CR-06        GCCTTTGGAGCTAAACTTGCACGAGATTTCGGTGTCCGTGATAACTGAATGGCATTGTCA
CR_right            GCCTTTGGAGCTAAACTTGCACGAGATTTCGGTGTCCGTGATAACTGAATGGCATTGTCA
Hd28_DRJ1R          GCCTTTGGAGCTAAACTTGCACGAGATTTCGGTGTCCGTGATAACTGAATGGCATTGTCA
                    ******* **** * ** *** *** ***   * ***  ******** **  ********
```

```
CR_left             GGCGTAAGCATTATGACGGCGGTGGTGTTAGGCGTTGCTTGGGAAGGTTGGTGAGCCACG
Hd28_DRJ1L          GGCGTAAGCATTATGACGGCGGTGGTGTTAGGCGTTGCTTGGGAAGGTTGGTGAGCCACG
Contig_CR-05        GGCGTAATATTTATGACGGC---------------------------------GGTGATG
Contig_CR-06        GGCGTAATATTTATGACGGC---------------------------------GGTGATG
CR_right            GGCGTAATATTTATGACGGC---------------------------------GGTGATG
Hd28_DRJ1R          GGCGTAATATTTATGACGGC---------------------------------GGTGATG
                    *******   **********                                  *  * *
```

```
CR_left             TCGACGTAGATTCAGGCGTGAAACGGAGAGCTGAAAGGTATAGTAGTCGTTCGACCAACT
Hd28_DRJ1L          TCGACGTAGATTCAGGCGTGAAACGGAGAGCTGAAAGGTATAGTAGTCGTTCGACCAACT
Contig_CR-05        TTAACGTAGGTTCAGGCGTTAGACTGAGAGCCGGAAGGTATGGTAGTCGTTCGACCGTCT
Contig_CR-06        TTAACGTAGGTTCAGGCGTTAGACTGAGAGCCGGAAGGTATGGTAGTCGTTCGACCGTCT
CR_right            TTAACGTAGGTTCAGGCGTTAGACTGAGAGCCGGAAGGTATGGTAGTCGTTCGACCGTCT
Hd28_DRJ1R          TTAACGTAGGTTCAGGCGTTAGACTGAGAGCCGGAAGGTATGGTAGTCGTTCGACCGTCT
                    *  ****** ********* * ** ****** * ******* **************  **

CR_left             CGGAATCACACCGACGAAAACTATTTGTCCATTGCTCAGGCTATGAGGACTTGAGGCAAT
Hd28_DRJ1L          CGGAATCACACCGACGAAAACTATTTGTCCATTGCTCAGGCTATGAGGACTTGAGGCAAT
Contig_CR-05        CCGAATCACGCTGACGAGAACTCTTCGTTCATCGCTTAAGCTCTGAGGACTTGACGCAAT
Contig_CR-06        CCGAATCACGCTGACGAGAACTCTTCGTTCATCGCTTAAGCTCTGAGGACTTGACGCAAT
CR_right            CCGAATCACGCTGACGAGAACTCTTCGTTCATCGCTTAAGCTCTGAGGACTTGACGCAAT
Hd28_DRJ1R          CCGAATCACGCTGACGAGAACTCTTCGTTCATCGCTTAAGCTCTGAGGACTTGACGCAAT
                    * ******* * ***** **** ** ** *** *** * ***  ********** *****

CR_left             CAAGTTCCGGACGCAAAGT----------TTTGTTTCCGGCCATAGATCACAAGTAAGAA
Hd28_DRJ1L          CAAGTTCCGGACGCAAAGT----------TTTGTTTCCGGCCATAGATCACAAGTAAGAA
Contig_CR-05        CAAGTTCTGGATGATAAACGCCAGACGCAAAGTTTTGTTTCCATAGATCACAAGTAAGAA
Contig_CR-06        CAAGTTCTGGATGATAAACGCCAGACGCAAAGTTTTGTTTCCATAGATCACAAGTAAGAA
CR_right            CAAGTTCTGGATGATAAACGCCAGACGCAAAGTTTTGTTTCCATAGATCACAAGTAAGAA
Hd28_DRJ1R          CAAGTTCTGGATGATAAACGCCAGACGCAAAGTTTTGTTTCCATAGATCACAAGTAAGAA
                    ******* *** *  **                 **    ********************

CR_left             CAGTTCTGGCAGACTGGACAGTCCACCGATACCCTCACAAGACTGTAAA-ACCTCTGGAC
Hd28_DRJ1L          CAGTTCTGGCAGACTGGACAGTCCACCGATACCCTCACAAGACTGTAAA-ACCTCTGGAC
Contig_CR-05        CAGTTCTTACGGACCTGACAGTCCAACATTACTCTCACGACACTGTAAA-ACCTATGGAC
Contig_CR-06        CAGTTCTTACGGACCTGACAGTTCAACATTACTCTCACGACACTGTAAA-ACCTATGGAC
CR_right            CAGTTCTTACGGACCTGACAGTCCAACATTACTCTCACGACACTGTAAA-ACCTATGGAC
Hd28_DRJ1R          CAGTTCTTACGGACCTGACAGTCCAACATTACTCTCACGACACTGTAAA-ACCTATGGAC
                    *******  * ***  ****** ** *  *** ***** ********** **** *****

CR_left             GTACTGCGCACGTACCCAAGTCCACGAAACACTCGTGCCACATCCATTTGATGTTGGCTT
Hd28_DRJ1L          GTACTGCGCACGTACCCAAGTCCACGAAACACTCGTGCCACATCCATTTGATGTTGGCTT
Contig_CR-05        GTACTTCGCACGTACCCGAGTCCACGAAACACTCGTGCTACATCCATTTGACGTTGGCCT
Contig_CR-06        GTACTTCGCACGTACCCGAGTCCACGAAACACTCGTGCTACATCCATTTGACGTTGGCCT
CR_right            GTACTTCGCACGTACCCGAGTCCACGAAACACTCGTGCTACATCCATTTGACGTTGGCCT
Hd28_DRJ1R          GTACTTCGCACGTACCCGAGTCCACGAAACACTCGTGCTACATCCATTTGACGTTGGCCT
                    ***** *********** ******************** ************ ****** *

CR_left             CTGCCAGGGCATGTCGCTAT--AGATTCCCAGCGTGTGCCCGTGTCCCAGAAAAAGCTTT
Hd28_DRJ1L          CTGCCAGGGCATGTCGCTAT--AGATTCCCAGCGTGTGCCCGTGTCCCAGAAAAAGCTTT
Contig_CR-05        TCT----GACAAGGCCCACTATAGAGTCCCAGCGTGTGCCCGTGTCCCGGAAGGAGCTTT
Contig_CR-06        TCT----GACAAGGCCCACTATAGAGTCCCAGCGTGTGCCCGTGTCCCGGAAGGAGCTTT
CR_right            TCT----GACAAGGCCCACTATAGAGTCCCAGCGTGTGCCCGTGTCCCGGAAGGAGCTTT
Hd28_DRJ1R          TCT----GACAAGGCCCACTATAGAGTCCCAGCGTGTGCCCGTGTCCCGGAAGGAGCTTT
                           * ** *****  *  *** ********************** ***  ******

CR_left             GTTCACAACCGCGACTTTTGACTGTCACCTTTGTTGTTACAGTTGCGTTACGTAACTACT
Hd28_DRJ1L          GTTCACAACCGCGACTTTTGACTGTCACCTTTGTTGTTACAGTTGCGTTACGTAACTACT
Contig_CR-05        T-ACTCCCAACCGACTTTTTAGTGTGACCTTTGTCGTTACAGTTGCGTTACGTAACTACT
Contig_CR-06        T-ACTCCCAACCGACTTTTTAGTGTGACCTTTGTCGTTACAGTTGCGTTACGTAACTACT
CR_right            T-ACTCCCAACCGACTTTTTAGTGTGACCTTTGTCGTTACAGTTGCGTTACGTAACTACT
Hd28_DRJ1R          T-ACTCCCAACCGACTTTTTAGTGTGACCTTTGTCGTTACAGTTGCGTTACGTAACTACT
                       * *     ******** * *** ******** *************************

CR_left             ACTGTGGAACTTCTCACTGCCACAGTCACAACAGCCTATCGACAACTGTTCTTTCAGTAT
Hd28_DRJ1L          ACTGTGGAACTTCTCACTGCCACAGTCACAACAGCCTATCGACAACTGTTCTTTCAGTAT
Contig_CR-05        ACTGTGGAACTTTTCACTACCACAGTTACAACAGCTTATCGGCAACAGTCCCTTCAGTAT
Contig_CR-06        ACTGTGGAACTTTTCACTACCACAGTTACAACAGCTTATCGGCAACAGTTCCTTCAGTAT
CR_right            ACTGTGGAACTTTTCACTACCACAGTTACAACAGCTTATCGGCAACAGTTCCTTCAGTAT
Hd28_DRJ1R          ACTGTGGAACTTTTCACTACCACAGTTACAACAGCTTATCGGCAACAGTTCCTTCAGTAT
                    ************ ***** ******* *******  ***** **** **** ********

CR_left             CAAGAAGTCATTATGCAAGTAGAGAATGGCCGGGGTTTTAAGTTGCATGAAA-GCACTCAA
Hd28_DRJ1L          CAAGAAGTCATTATGCAAGTAGAGAATGGCCGGGGTTTTAAGTTGCATGAAA-GCACTCAA
Contig_CR-05        CAAGAAGTAATTATTCAAGTGGAGAATGGCCGGGGTTTTAAGTTGCATGAAA-GCACTCAA
Contig_CR-06        CAAGAAGTAATTATTCAAGTGGAGAATGGCCGGGGTCGG-AGTTGCATGAAA-GCACTCAA
CR_right            CAAGAAGTAATTATTCAAGTGGAGAATGGCCGGGGTCGG-AGTTGCATGAAAAGCACGGAA
```

```
Hd28_DRJ1R          CAAGAAGTAATTATTCAAGTGGAGAATGGCCGGGGTCGG-AGTTGCATGAAAAGCACGGAA
                    ******** ***** ***** ***************    ************ ****  **

CR_left             TTGTCTACAATGCAGACGAG
Hd28_DRJ1L          TTGTCTACAATGCAGACGAG
Contig_CR-05        TTGTCTACAATGCAGACGAG
Contig_CR-06        TTGTCTACAATGCAGACGAG
CR_right            GTGTGTACAATACAGACGGG
Hd28_DRJ1R          GTGTGTACAATACAGACGGG
                     *** ****** ****** *
```
