## Additional file 8 for "Conserved and specific genomic features of endogenous polydnaviruses revealed by whole genome sequencing of two ichneumonid wasps"

<table>
<tr><th colspan="9">Genes in synteny blocks Figure 6C, a</th></tr>
<tr><th>Hd scaffold</th><th>Position in scaffold</th><th>Hd gene ID</th><th>Orthogroup #</th><th>Best Blast hit</th><th></th><th>Cs gene</th><th>Orthogroup #</th><th>Position in scaffold</th></tr>
<tr><td>scaffold144</td><td>1380719..1397417</td><td>HD005010</td><td>OG0000722</td><td>gi|817190146|ref|XP_012270430.1|uncharacterized protein LOC105694386 isoform X2 [Orussus abietii</td><td>Expect = 0.000e+0</td><td>gene.scaffold_16.g181.t1</td><td>OG0000722</td><td>2082674..2097848</td></tr>
<tr><td>scaffold144</td><td>1402033..1404758</td><td>HD005011</td><td>OG0008876</td><td>gi|1000730346|ref|XP_015587582.1 dynein assembly factor 4, axonemal-like [Cephus cinctus]</td><td>Expect = 8.300e-150</td><td>gene.scaffold_16.g180.t1</td><td>OG0008876</td><td>2075777..2079078</td></tr>
<tr><td>scaffold144</td><td>1405145..1406140</td><td>HD005012</td><td>OG0008079</td><td>gi|340719499|ref|XP_003398190.1|vascular endothelial growth factor A-A [Bombus terrestris]</td><td>Expect = 7.800e-47</td><td>gene.scaffold_16.g179.t1</td><td>OG0008079</td><td>2074775..2075425</td></tr>
<tr><td>scaffold144</td><td>1410042..1410578</td><td>HD005013</td><td>OG0009360</td><td>gi|987906890|ref|XP_015428945.1|PREDICTED: uncharacterized protein LOC107185710 [Dufourea nov</td><td>Expect = 2.700e-41</td><td>gene.scaffold_16.g178.t1</td><td>OG0009360</td><td>2064292..2070214</td></tr>
<tr><td>scaffold144</td><td>1412742..1423051</td><td>HD005014</td><td>OG0007034</td><td>gi|755949684|ref|XP_011301181.1|PREDICTED: nucleoprotein TPR [Fopius arisanus]</td><td>Expect = 0.000e+0</td><td>gene.scaffold_16.g177.t1</td><td>OG0007034</td><td>2058216..2067490</td></tr>
<tr><td>scaffold144</td><td>1423611..1429572</td><td>HD005015</td><td>OG0004177</td><td>gi|1000729872|ref|XP_015587356.1 protein virilizer isoform X1 [Cephus cinctus]</td><td>Expect = 0.000e+0</td><td>NA</td><td></td><td></td></tr>
<tr><td>scaffold144</td><td>1429964..1432653</td><td>HD005016</td><td>OG0007546</td><td>gi|1070214570|ref|XP_018376608.1 PREDICTED: pentatricopeptide repeat-containing protein 1, mitoch</td><td>Expect = 7.200e-127</td><td>NA</td><td></td><td></td></tr>
<tr><td>scaffold144</td><td>1434554..1435954</td><td>HD005017</td><td>OG0009748</td><td>gi|1070154465|ref|XP_018340726.1 PREDICTED: RISC-loading complex subunit TARBP2-like [Trachymyr</td><td>Expect = 2.500e-30</td><td>NA</td><td></td><td></td></tr>
<tr><td>scaffold144</td><td>1438643..1440697</td><td>HD005018</td><td>OG0003141</td><td>gi|645023101|ref|XP_008202437.1|PREDICTED: neuralized-like protein 2 [Nasonia vitripennis]</td><td>Expect = 8.100e-114</td><td>gene.scaffold_16.g36.t1</td><td>OG0003141</td><td>254028..255633</td></tr>
<tr><td>scaffold144</td><td>1442194..1445201</td><td>HD005019</td><td>OG0003140</td><td>gi|1317987018|ref|XP_023287837.1 neuronal acetylcholine receptor subunit alpha-5 isoform X1 [Orussu</td><td>Expect = 2.500e-59</td><td></td><td></td><td></td></tr>
<tr><td>scaffold144</td><td>1446370..1448826</td><td>HD005020</td><td>OG0009176</td><td>gi|939631532|ref|XP_008554100.2|PREDICTED: neuronal acetylcholine receptor subunit alpha-5-like [N</td><td>Expect = 7.200e-80</td><td>gene.scaffold_16.g37.t1</td><td>OG0009176</td><td>257530..261137</td></tr>
<tr><td>scaffold144</td><td>1450485..1451362</td><td>HD005021</td><td>OG0007822</td><td>Cys2_Hd20</td><td></td><td></td><td></td><td></td></tr>
<tr><td>scaffold144</td><td>1454299..1455511</td><td>HD005022</td><td>OG0007822</td><td>Cys1_Hd20</td><td></td><td></td><td></td><td></td></tr>
<tr><td>scaffold144</td><td>1457007..1457513</td><td>HD005023</td><td>OG0001009</td><td>gi|805787897|ref|XP_012141035.1|PREDICTED: uncharacterized protein LOC100876911 isoform X2 [M</td><td>Expect = 2.000e-20</td><td>gene.scaffold_16.g38.t1</td><td>OG0001009</td><td>270567..274688</td></tr>
<tr><td>scaffold144</td><td>1464373..1467934</td><td>HD005024</td><td>OG0001009</td><td>gi|795056915|ref|XP_011872016.1|PREDICTED: uncharacterized protein LOC105564328 isoform X1 [Vo</td><td>Expect = 2.100e-124</td><td></td><td></td><td></td></tr>
<tr><td>scaffold144</td><td>1472127..1477975</td><td>HD005025</td><td>OG0006050</td><td>gi|1000745701|ref|XP_015595595.1 cysteine and histidine-rich protein 1 homolog isoform X5 [Cephus c</td><td>Expect = 2.400e-190</td><td>gene.scaffold_16.g39.t1</td><td>OG0006050</td><td>278822..284602</td></tr>
<tr><td>scaffold144</td><td>1479354..1480103</td><td>HD005026</td><td>orphan</td><td>gi|759078894|ref|XP_011349245.1|polyadenylate-binding protein 1 [Ooceraea biroi]</td><td>Expect = 2.100e-23</td><td>gene.scaffold_16.g40.t1</td><td>orphan</td><td>287471..288253</td></tr>
<tr><td></td><td></td><td></td><td></td><td></td><td></td><td>gene.scaffold_16.g42.t1</td><td>orphan</td><td>331489..359465</td></tr>
<tr><td>scaffold144</td><td>1508038..1591917</td><td>HD005027</td><td>OG0001516</td><td>gi|755949182|ref|XP_011301029.1|PREDICTED: regulating synaptic membrane exocytosis protein 1 iso</td><td>Expect = 0.000e+0</td><td>gene.scaffold_16.g43.t1</td><td>OG0001516</td><td>385633..394202</td></tr>
<tr><td>scaffold144</td><td>1601947..1606850</td><td>HD005028</td><td>orphan</td><td>gi|998501744|ref|XP_015510887.1|PREDICTED: uncharacterized protein LOC107217756 [Neodiprion le</td><td>Expect = 2.200e-36</td><td>gene.scaffold_16.g44.t1</td><td>OG0001516</td><td>402325..405981</td></tr>
<tr><td>scaffold144</td><td></td><td></td><td></td><td></td><td></td><td>gene.scaffold_16.g45.t1</td><td>OG0001516</td><td>413940..415699</td></tr>
<tr><td>scaffold144</td><td>1607444..1607878</td><td>HD005029</td><td>OG0007431</td><td>gi|815909475|ref|XP_003489119.2|chromosome transmission fidelity protein 8 homolog [Bombus imp</td><td>Expect = 9.300e-30</td><td>gene.scaffold_16.g46.t1</td><td>OG0007431</td><td>427568..428032</td></tr>
<tr><td>scaffold144</td><td>1609469..1618501</td><td>HD005030</td><td>OG0001137</td><td>gi|970888608|ref|XP_015109471.1|PREDICTED: uncharacterized protein LOC107036203 [Diachasma all</td><td>Expect = 2.600e-122</td><td>gene.scaffold_16.g47.t1</td><td>OG0001137</td><td>431251..442478</td></tr>
</table>

<table>
<tr><th colspan="9">Genes in synteny blocks Figure 6C, b</th></tr>
<tr><th>Hd scaffold</th><th>Position in scaffold</th><th>Hd gene ID</th><th>Orthogroup #</th><th>Best Blast hit</th><th></th><th>Cs gene</th><th>Orthogroup #</th><th>Position in scaffold</th></tr>
<tr><td>scaffold351</td><td>2570811..2574280</td><td>HD010552</td><td>OG0006536</td><td>gi|970890219|ref|XP_015110360.1|PREDICTED: GPI mannosyltransferase 4 [Diachasma alloeum]</td><td>Expect = 6.400e-248</td><td>gene.scaffold_16.g210.t1</td><td>OG0006536</td><td>2319748..2322545</td></tr>
<tr><td>scaffold351</td><td>2575688..2584750</td><td>HD010553</td><td>OG0000820</td><td>gi|1000729971|ref|XP_015587409.1 histone deacetylase 6 isoform X3 [Cephus cinctus]</td><td>Expect = 0.000e+0</td><td>gene.scaffold_16.g211.t1</td><td>OG0000820</td><td>2323661..2331643</td></tr>
<tr><td>scaffold351</td><td>2585486..2590674</td><td>HD010554</td><td>OG0008529</td><td>gi|1000729989|ref|XP_015587415.1 mitochondrial intermediate peptidase isoform X1 [Cephus cinctus]</td><td>Expect = 1.300e-309</td><td>gene.scaffold_16.g212.t1</td><td>OG0008529</td><td>2333001..2336567</td></tr>
<tr><td>scaffold351</td><td>2592030..2601204</td><td>HD010555</td><td>OG0005613</td><td>gi|755986949|ref|XP_011311356.1|PREDICTED: cullin-4A [Fopius arisanus]</td><td>Expect = 0.000e+0</td><td>gene.scaffold_16.g213.t1</td><td>OG0005613</td><td>2338026..2342787</td></tr>
<tr><td>scaffold351</td><td>2603266..2606649</td><td>HD010556</td><td>OG0007486</td><td>gi|1000718247|ref|XP_015598324.1 protein crooked neck [Cephus cinctus]</td><td>Expect = 1.700e-240</td><td>gene.scaffold_16.g214.t1</td><td>OG0007486</td><td>2344137..2346984</td></tr>
<tr><td></td><td></td><td></td><td></td><td></td><td></td><td>gene.scaffold_16.g216.t1</td><td>orphan</td><td>2352827..2353396</td></tr>
<tr><td>scaffold351</td><td>2614497..2669283</td><td>HD010557</td><td>OG0000315</td><td>gi|1059231176|ref|XP_017760895.1 PREDICTED: nuclear factor 1 X-type-like isoform X7 [Eufriesea mexi</td><td>Expect = 0.000e+0</td><td>gene.scaffold_16.g218.t1</td><td>OG0000315</td><td>2360992..2364340</td></tr>
<tr><td></td><td></td><td></td><td></td><td></td><td></td><td>gene.scaffold_16.g219.t1</td><td>OG0000315</td><td>2373432..2374064</td></tr>
<tr><td></td><td></td><td></td><td></td><td></td><td></td><td>gene.scaffold_16.g220.t1</td><td>OG0000315</td><td>2387249..2388404</td></tr>
<tr><td>scaffold351</td><td>2678768..2680678</td><td>HD010558</td><td>OG0007969</td><td>gi|755943768|ref|XP_011299232.1|PREDICTED: actin-related protein 2/3 complex subunit 4 [Fopius ari</td><td>Expect = 5.700e-81</td><td>gene.scaffold_16.g221.t1</td><td>OG0007969</td><td>2416191..2419469</td></tr>
<tr><td>scaffold351</td><td>2683737..2685050</td><td>HD010559</td><td>orphan</td><td>N1_Hd18</td><td></td><td>gene.scaffold_16.g223.t1</td><td>OG0010684</td><td>2424942..2425688</td></tr>
<tr><td></td><td></td><td></td><td></td><td></td><td></td><td>gene.scaffold_16.g564.t1</td><td>orphan</td><td>2426081..2426764</td></tr>
<tr><td></td><td></td><td></td><td></td><td></td><td></td><td>gene.scaffold_16.g224.t1</td><td>orphan</td><td>2427951..2428691</td></tr>
<tr><td>scaffold351</td><td>2688803..2689573</td><td>HD010560</td><td>OG0005107</td><td>gi|972178531|ref|XP_015191350.1|PREDICTED: transmembrane protein 45B-like [Polistes dominula]</td><td>Expect = 3.600e-95</td><td>gene.scaffold_16.g225.t1</td><td>OG0005107</td><td>2430738..2431508</td></tr>
<tr><td>scaffold351</td><td>2690793..2699707</td><td>HD010561</td><td>OG0003436</td><td>gi|820860113|ref|XP_003697297.2|PREDICTED: LOW QUALITY PROTEIN: enhancer of mRNA-decapping</td><td>Expect = 0.000e+0</td><td>NA</td><td></td><td></td></tr>
<tr><td>scaffold351</td><td>2700580..2701786</td><td>HD010562</td><td>OG0007575</td><td>gi|1227092722|gb|OXU18804.1| hypothetical protein TSAR_005200 [Trichomalopsis sarcophagae]</td><td>Expect = 3.400e-82</td><td>NA</td><td></td><td></td></tr>
<tr><td>scaffold351</td><td>2706953..2708212</td><td>HD010563</td><td>OG0009633</td><td>gi|1580172942|ref|XP_028048451. uncharacterized protein LOC114255006 [Monomorium pharaonis]</td><td>Expect = 1.100e-29</td><td>NA</td><td></td><td></td></tr>
<tr><td>scaffold351</td><td>2712014..2713607</td><td>HD010564</td><td>OG0000118</td><td>gi|795032310|ref|XP_011863894.1|PREDICTED: speckle-type POZ protein A-like [Vollenhovia emeryi]</td><td>Expect = 6.600e-20</td><td>NA</td><td></td><td></td></tr>
<tr><td>scaffold351</td><td>2714791..2716848</td><td>HD010565</td><td>OG0005178</td><td>gi|1000730046|ref|XP_015587445.1 actin-related protein 10 [Cephus cinctus]</td><td>Expect = 2.800e-165</td><td>gene.scaffold_16.g124.t1</td><td>OG0005178</td><td>1439439..1441701</td></tr>
<tr><td>scaffold351</td><td>2717065..2719279</td><td>HD010566</td><td>OG0008078</td><td>gi|817195095|ref|XP_012273078.1|eukaryotic translation initiation factor 3 subunit F [Orussus abietinu</td><td>Expect = 4.500e-139</td><td>gene.scaffold_16.g125.t1</td><td>OG0008078</td><td>1441946..1443651</td></tr>
<tr><td>scaffold351</td><td>2720235..2740116</td><td>HD010567</td><td>OG0000165</td><td>gi|915666462|gb|KOC68367.1| Phosphatidylinositol 4-phosphate 5-kinase type-1 alpha [Habropod</td><td>Expect = 0.000e+0</td><td>gene.scaffold_16.g126.t1</td><td>OG0000165</td><td>1444610..1449264</td></tr>
<tr><td>scaffold351</td><td>2744556..2745842</td><td>HD010568</td><td>OG0000280</td><td>gi|998499361|ref|XP_015509568.1|PREDICTED: venom peptide isomerase heavy chain-like [Neodiprion</td><td>Expect = 5.200e-119</td><td>gene.scaffold_16.g129.t1</td><td>OG0000280</td><td>1468965..1476057</td></tr>
<tr><td>scaffold351</td><td>2746325..2748699</td><td>HD010569</td><td>OG0010371</td><td>no hit</td><td></td><td></td><td></td><td></td></tr>
<tr><td>scaffold351</td><td>2749640..2750960</td><td>HD010570</td><td>OG0010371</td><td>no hit</td><td></td><td></td><td></td><td></td></tr>
<tr><td>scaffold351</td><td>2754546..2756977</td><td>HD010571</td><td>OG0000280</td><td>gi|755977086|ref|XP_011308165.1|PREDICTED: trypsin-like [Fopius arisanus]</td><td>Expect = 1.600e-80</td><td>gene.scaffold_16.g130.t1</td><td>OG0000280</td><td>1479694..1481353</td></tr>
</table>

| | | | | | | | | |
|---|---|---|---|---|---|---|---|---|
| scaffold351 | 2757727..2759955 | HD010572 | OG0000280 | gi\|755977086\|ref\|XP_011308165.1\|PREDICTED: trypsin-like [Fopius arisanus] | Expect = 2.500e-81 | gene.scaffold_16.g131.t1 | OG0000280 | 1482054..1483447 |
| scaffold351 | 2760364..2762511 | HD010573 | OG0000280 | gi\|755977083\|ref\|XP_011308164.1\|PREDICTED: trypsin II-P29-like [Fopius arisanus] | Expect = 4.300e-75 | gene.scaffold_16.g132.t1 | OG0000280 | 1484135..1488423 |
| scaffold351 | 2765019..2770447 | HD010574 | OG0008378 | gi\|755977080\|ref\|XP_011308163.1\|PREDICTED: carboxypeptidase B-like [Fopius arisanus] | Expect = 8.000e-215 | gene.scaffold_16.g133.t1 | OG0008378 | 1489809..1493726 |

**Genes in synteny blocks Figure 6C, c**

| Hd scaffold | Position in scaffold | Hd gene ID | Orthogroup # | Best Blast hit | | Cs gene | Orthogroup # | Position in scaffold |
|---|---|---|---|---|---|---|---|---|
| | | NA | | | | *gene.scaffold_5890.g33.* | *OG0000028* | *162272..162676* |
| | | NA | | | | *gene.scaffold_5890.g32.* | *OG0000021* | *161726..162136* |
| | | NA | | | | *gene.scaffold_5890.g31.* | *OG0000024* | *156325..156699* |
| | | NA | | | | *gene.scaffold_5890.g30.* | *orphan* | *151548..155209* |
| | | NA | | | | *gene.scaffold_5890.g29.* | *OG0000024* | *151509..151883* |
| | | NA | | | | *gene.scaffold_5890.g28.* | *OG0002056* | *146321..147118* |
| | | NA | | | | *///* | | |
| | | NA | | | | *gene.scaffold_5890.g59* | *OG0002056* | *134843..135562* |
| scaffold351 | 2270991..2271302 | HD010503 | OG0000028 | gi\|170058994\|ref\|XP_001865168.1\|histone 1 [Culex quinquefasciatus] | Expect = 3.900e-48 | | | |
| scaffold351 | 2273424..2278678 | HD010504 | OG0003636 | gi\|1000742598\|ref\|XP_015593950.1 equilibrative nucleoside transporter 4 [Cephus cinctus] | Expect = 1.500e-267 | gene.scaffold_5890.g22.t | OG0003636 | 128010..132737 |
| scaffold351 | 2280628..2284708 | HD010505 | OG0001788 | gi\|954555894\|ref\|XP_014601533.1\|PREDICTED: serine/threonine-protein kinase ICK [Polistes canadens | Expect = 3.200e-256 | gene.scaffold_5890.g20.t | OG0001788 | 120284..125583 |
| scaffold351 | 2288342..2289300 | HD010506 | orphan | no hit | | gene.scaffold_5890.g18.t | OG0000067 | 111355..111939 |
| scaffold351 | 2292636..2299704 | HD010507 | OG0001161 | gi\|970889125\|ref\|XP_015109758.1\|PREDICTED: DENN domain-containing protein 1A-like isoform X1 [D | Expect = 0.000e+0 | gene.scaffold_5890.g17.t | OG0001161 | 101367..109432 |
| scaffold351 | 2300822..2301232 | HD010508 | OG0000021 | gi\|170053440\|ref\|XP_001862674.1\|histone H3 type 2 [Culex quinquefasciatus] | Expect = 3.000e-64 | gene.scaffold_5890.g16.t | OG0000021 | 97954..100617 |
| scaffold351 | 2302839..2303213 | HD010509 | OG0000024 | gi\|815814502\|ref\|XP_012228276.1\|PREDICTED: histone H2A-like [Linepithema humile] | Expect = 1.700e-58 | | | |
| scaffold351 | 2304346..2305863 | HD010510 | OG0008873 | gi\|1000757495\|ref\|XP_015601701.1 activator of basal transcription 1 isoform X2 [Cephus cinctus] | Expect = 1.900e-83 | | | |
| scaffold351 | 2306413..2308908 | HD010511 | OG0004415 | gi\|307203421\|gb\|EFN82496.1\| mRNA-decapping enzyme 2 [Harpegnathos saltator] | Expect = 1.000e-159 | gene.scaffold_5890.g15.t | OG0004415 | 95411..97479 |
| scaffold351 | 2310086..2311637 | HD010512 | OG0007967 | gi\|954580551\|ref\|XP_014614868.1\|PREDICTED: heme oxygenase 1 [Polistes canadensis] | Expect = 3.300e-93 | | | |
| scaffold351 | 2312957..2318568 | HD010513 | OG0006585 | gi\|1000715934\|ref\|XP_015586052.1 peroxisomal multifunctional enzyme type 2 isoform X1 [Cephus cin | Expect = 2.200e-259 | gene.scaffold_5890.g14.t | OG0006585 | 86991..95435 |
| scaffold351 | 2320285..2328116 | HD010514 | OG0002518 | gi\|1000757483\|ref\|XP_015601693.1 hypoxia up-regulated protein 1 isoform X1 [Cephus cinctus] | Expect = 1.000e-304 | gene.scaffold_5890.g13.t | OG0002518 | 79568..85710 |
| scaffold351 | 2329743..2330387 | HD010515 | orphan | Rep5_Hd17 | | | | |
| scaffold351 | /// | | | | | | | |
| scaffold351 | 2335459..2336166 | HD010519 | orphan | Rep1_Hd17 | | | | |
| scaffold351 | 2336786..2337193 | HD010520 | orphan | rep partial_Hd17 | | | | |
| scaffold351 | 2337418..2340007 | HD010521 | OG0002835 | gi\|808129402\|ref\|XP_012167532.1\|WW domain-binding protein 2 isoform X1 [Bombus terrestris] | Expect = 2.300e-101 | gene.scaffold_5890.g12.t | OG0002835 | 76069..78711 |
| scaffold351 | 2345118..2346993 | HD010522 | OG0006513 | gi\|1227095170\|gb\|OXU20342.1\| hypothetical protein TSAR_005800 [Trichomalopsis sarcophagae] | Expect = 1.500e-104 | gene.scaffold_5890.g11.t | OG0006513 | 69897..70791 |
| scaffold351 | 2349858..2352634 | HD010523 | OG0007554 | gi\|1059880711\|ref\|XP_017795737.1\|PREDICTED: ubiquitin domain-containing protein UBFD1-like [Habro | Expect = 2.500e-105 | gene.scaffold_5890.g10.t | OG0007554 | 63578..65852 |
| scaffold351 | 2353611..2362101 | HD010524 | OG0003626 | gi\|970912543\|ref\|XP_015122543.1\|PREDICTED: polycomb protein suz12-B [Diachasma alloeum] | Expect = 0.000e+0 | gene.scaffold_5890.g9.t1 | OG0003626 | 55732..62546 |
| scaffold351 | 2363619..2368282 | HD010525 | OG0004797 | gi\|1000730147\|ref\|XP_015587495.1 methyltransferase-like protein 22 [Cephus cinctus] | Expect = 5.700e-92 | gene.scaffold_5890.g8.t1 | OG0004797 | 51570..54218 |
| scaffold351 | 2368680..2372470 | HD010526 | OG0006517 | gi\|1476364581\|gb\|AXY94695.1\| disulfide-isomerase [Habrobracon hebetor] | Expect = 1.100e-196 | gene.scaffold_5890.g7.t1 | OG0006517 | 44960..49478 |

**Genes in synteny blocks Figure 7C, a**

| Hd scaffold | Position in scaffold | Hd gene ID | Orthogroup # | Best Blast hit | | Cs gene | Orthogroup # | Position in scaffold |
|---|---|---|---|---|---|---|---|---|
| Scaffold 91 | 339180-342661 | HD016092 | OG0004987 | gi\|780657091\|ref\|XP_011691874.1\|PREDICTED: solute carrier organic anion transporter family membe | Expect = 1.400e-275 | gene.scaffold_14.g96.t1 | OG0004987 | 672138..680445 |
| Scaffold 91 | 370839-379409 | HD016093 | OG0004824 | gi\|769843355\|ref\|XP_011632893.1\|solute carrier organic anion transporter family member 5A1 isoforn | Expect = 0.000e+0 | gene.scaffold_14.g97.t1 | OG0004824 | 687109..693641 |
| Scaffold 91 | 386420-391295 | HD016094 | OG0004411 | gi\|755943227\|ref\|XP_011299047.1\|PREDICTED: G-protein coupled receptor Mth isoform X2 [Fopius ari | Expect = 9.200e-139 | gene.scaffold_14.g98.t1 | orphan | 691335..692107 |
| Scaffold 91 | 392450-394130 | HD016095 | OG0001577 | gi\|1492480063\|ref\|XP_026674315.1 probable RNA methyltransferase CG11342 isoform X1 [Ceratina cal | Expect = 5.800e-72 | gene.scaffold_14.g99.t1 | OG0001577 | 701142..708415 |
| Scaffold 91 | 396109-397563 | HD016096 | OG0001965 | gi\|749787677\|ref\|XP_011147350.1\|myosin-2 essential light chain isoform X2 [Harpegnathos saltator] | Expect = 3.300e-77 | gene.scaffold_14.g100.t1 | OG0001965 | 709518..710732 |
| Scaffold 91 | 398929-404614 | HD016097 | OG0004676 | gi\|972210109\|ref\|XP_015187432.1\|PREDICTED: WD repeat, SAM and U-box domain-containing protein | Expect = 2.500e-298 | gene.scaffold_14.g101.t1 | OG0004676 | 713039..718640 |
| Scaffold 91 | 406627-407543 | HD016098 | OG0007680 | gi\|817183740\|ref\|XP_012278017.1\|GATA zinc finger domain-containing protein 1 isoform X2 [Orussus a | Expect = 1.600e-105 | gene.scaffold_14.g102.t1 | OG0007680 | 720734..721681 |
| Scaffold 91 | 408455-411290 | HD016099 | OG0005425 | gi\|48142301\|ref\|XP_397320.1\| AP-3 complex subunit sigma-2 isoform X1 [Apis mellifera] | Expect = 8.100e-92 | gene.scaffold_14.g103.t1 | OG0005425 | 722419..724763 |
| Scaffold 91 | 412851-415250 | HD016100 | OG0001553 | gi\|383857701\|ref\|XP_003704342.1 PREDICTED: beta-1,3-galactosyltransferase 1-like isoform X2 [Mega | Expect = 3.200e-143 | NA | | |
| Scaffold 91 | 418638-419902 | HD016101 | OG0008604 | gi\|345486223\|ref\|XP_003425426.1 PREDICTED: G patch domain-containing protein 4-like [Nasonia vitri | Expect = 1.400e-47 | NA | | |
| Scaffold 91 | 420807-422918 | HD016102 | OG0004778 | gi\|970907664\|ref\|XP_015119881.1\|PREDICTED: type 2 phosphatidylinositol 4,5-bisphosphate 4-phosph | Expect = 7.800e-127 | NA | | |
| Scaffold 91 | 425105..426659 | HD016103 | orphan | no hit | | NA | | |
| Scaffold 91 | 427639..428218 | HD016104 | orphan | no hit | | NA | | |
| Scaffold 91 | 429397..431144 | HD016105 | orphan | no hit | | NA | | |

<table>
<tr><td>Scaffold 91</td><td>433475..434375</td><td>HD016106</td><td>orphan</td><td>no hit</td><td></td><td>NA</td><td></td><td></td></tr>
<tr><td>Scaffold 91</td><td>435700..436543</td><td>HD016107</td><td>orphan</td><td>no hit</td><td></td><td>NA</td><td></td><td></td></tr>
<tr><td>Scaffold 91</td><td>437892..438750</td><td>HD016108</td><td>orphan</td><td>no hit</td><td></td><td>NA</td><td></td><td></td></tr>
<tr><td>Scaffold 91</td><td>440012..441810</td><td>HD016109</td><td>orphan</td><td>no hit</td><td></td><td>NA</td><td></td><td></td></tr>
<tr><td>Scaffold 91</td><td>443803..449675</td><td>HD016110</td><td>orphan</td><td>no hit</td><td></td><td>NA</td><td></td><td></td></tr>
<tr><td>Scaffold 91</td><td>450623..452672</td><td>HD016111</td><td>OG0005467</td><td>gi|987911334|ref|XP_015431390.1|PREDICTED: DNA repair protein XRCC1 [Dufourea novaeangliae]</td><td>Expect = 5.400e-86</td><td>NA</td><td></td><td></td></tr>
<tr><td>Scaffold 91</td><td>453857..454591</td><td>HD016112</td><td>orphan</td><td>IVSP_U1</td><td></td><td>NA</td><td></td><td></td></tr>
<tr><td>Scaffold 91</td><td>///</td><td></td><td></td><td></td><td></td><td>NA</td><td></td><td></td></tr>
<tr><td>Scaffold 91</td><td>467172..467876</td><td>HD016611</td><td>orphan</td><td>IVSP_U25</td><td></td><td>NA</td><td></td><td></td></tr>
<tr><td>Scaffold 91</td><td>467893..468866</td><td>HD016610</td><td>OG0008504</td><td>no hit</td><td></td><td>NA</td><td></td><td></td></tr>
<tr><td>Scaffold 91</td><td>470135..471607</td><td>HD016121</td><td>orphan</td><td>N-gene1_Hd15</td><td></td><td>NA</td><td></td><td></td></tr>
<tr><td>Scaffold 91</td><td>474019..474459</td><td>HD016122</td><td>orphan</td><td>no hit</td><td></td><td>NA</td><td></td><td></td></tr>
<tr><td>Scaffold 91</td><td>475549..482783</td><td>HD016123</td><td>OG0008504</td><td>gi|1000724119|ref|XP_015610535.1 uncharacterized protein LOC107275188 isoform X2 [Cephus cinctus</td><td>Expect = 2.200e-140</td><td>gene.scaffold_50.g34.t1.</td><td>OG0008504</td><td>228808..236437</td></tr>
<tr><td>Scaffold 91</td><td>485760..486020</td><td>HD016616</td><td>orphan</td><td>U1_Hd33</td><td></td><td></td><td></td><td></td></tr>
<tr><td>Scaffold 91</td><td>487963..488427</td><td>HD016613</td><td>orphan</td><td>U2_Hd33</td><td></td><td></td><td></td><td></td></tr>
<tr><td>Scaffold 91</td><td>487593..496279</td><td>HD016124</td><td>OG0008877</td><td>gi|1573754945|ref|XP_028042060.1 inter-alpha-trypsin inhibitor heavy chain H4-like [Bombyx mandarin</td><td>Expect = 1.400e-43</td><td>gene.scaffold_50.g35.t1</td><td>OG0008877</td><td>239447..245088</td></tr>
<tr><td>Scaffold 91</td><td>496775..497693</td><td>HD016125</td><td>OG0008877</td><td>gi|1185578680|ref|XP_020720113.1 uncharacterized protein LOC100648520 [Bombus terrestris]</td><td>Expect = 2.200e-27</td><td></td><td></td><td></td></tr>
<tr><td>Scaffold 91</td><td>498816..502616</td><td>HD016126</td><td>OG0007035</td><td>gi|817052951|ref|XP_012259318.1|tetratricopeptide repeat protein 27 [Athalia rosae]</td><td>Expect = 4.100e-262</td><td>gene.scaffold_50.g36.t1</td><td>OG0007035</td><td>245526..250163</td></tr>
<tr><td>Scaffold 91</td><td>503255..504795</td><td>HD016127</td><td>orphan</td><td>no hit</td><td></td><td>gene.scaffold_50.g37.t1</td><td>orphan</td><td>250869..252609</td></tr>
<tr><td>Scaffold 91</td><td>506500..510737</td><td>HD016128</td><td>OG0008077</td><td>gi|970916129|ref|XP_015124492.1|PREDICTED: uncharacterized protein LOC107046395 [Diachasma all</td><td>Expect = 3.700e-255</td><td>gene.scaffold_50.g38.t1</td><td>OG0008077</td><td>253612..257408</td></tr>
<tr><td>Scaffold 91</td><td>512122..516356</td><td>HD016129</td><td>OG0004493</td><td>gi|1061111877|ref|XP_017881969.1 uridine-cytidine kinase-like 1 [Ceratina calcarata]</td><td>Expect = 1.600e-268</td><td>gene.scaffold_50.g39.t1</td><td>OG0004493</td><td>258014..261162</td></tr>
<tr><td>Scaffold 91</td><td>517628..518212</td><td>HD016130</td><td>OG0002787</td><td>gi|665791319|ref|XP_008543211.1|PREDICTED: inosine triphosphate pyrophosphatase [Microplitis den</td><td>Expect = 2.700e-82</td><td>gene.scaffold_50.g40.t1</td><td>OG0002787</td><td>262054..262638</td></tr>
<tr><td>Scaffold 91</td><td>521509..534951</td><td>HD016131</td><td>OG0003558</td><td>gi|665791317|ref|XP_008543209.1|PREDICTED: uncharacterized protein LOC103568231 [Microplitis de</td><td>Expect = 0.000e+0</td><td>NA</td><td></td><td></td></tr>
<tr><td>Scaffold 91</td><td>536918..537427</td><td>HD016132</td><td>OG0000328</td><td>Vank1_Hd24</td><td></td><td>NA</td><td></td><td></td></tr>
<tr><td>Scaffold 91</td><td>538200..539270</td><td>HD016133</td><td>OG0010755</td><td>Vinx1_Hd24</td><td></td><td>NA</td><td></td><td></td></tr>
<tr><td>Scaffold 91</td><td>541304..542734</td><td>HD016134</td><td>orphan</td><td>Ngene-4</td><td></td><td>NA</td><td></td><td></td></tr>
<tr><td>Scaffold 91</td><td>///</td><td></td><td></td><td></td><td></td><td>NA</td><td></td><td></td></tr>
<tr><td>Scaffold 91</td><td>567663..567944</td><td>HD016614</td><td>orphan</td><td>Ngene-3</td><td></td><td>NA</td><td></td><td></td></tr>
<tr><td>Scaffold 91</td><td>573680..574969</td><td>HD016150</td><td>orphan</td><td>N1_Hd29</td><td></td><td>NA</td><td></td><td></td></tr>
<tr><td>Scaffold 91</td><td>579785..581438</td><td>HD016151</td><td>OG0001925</td><td>gi|1000751647|ref|XP_015598681.1 protein l(2)37Cc [Cephus cinctus] >gi|1000751649|ref|XP_015598(</td><td>Expect = 7.900e-133</td><td>NA</td><td></td><td></td></tr>
<tr><td>Scaffold 91</td><td>583666..596697</td><td>HD016152</td><td>OG0000557</td><td>gi|970889107|ref|XP_015109747.1|PREDICTED: myosin-VIIa-like [Diachasma alloeum]</td><td>Expect = 0.000e+0</td><td>NA</td><td></td><td></td></tr>
<tr><td>Scaffold 91</td><td>733490..737430</td><td>HD016153</td><td>OG0002720</td><td>gi|1061140064|ref|XP_017893504.1 transcription factor Sox-2 isoform X1 [Ceratina calcarata]</td><td>Expect = 3.700e-147</td><td>NA</td><td></td><td></td></tr>
</table>

**Genes in synteny blocks Figure 7C, b**

<table>
<tr><th>Hd scaffold</th><th>Position in scaffold</th><th>Hd gene ID</th><th>Orthogroup #</th><th>Best Blast hit</th><th></th><th>Cs gene</th><th>Orthogroup #</th><th>Position in scaffold</th></tr>
<tr><td>Scaffold 1275</td><td>5695132..5698339</td><td>HD001703</td><td>OG0002287</td><td>gi|1615708828|gb|TGZ38293.1| Uncharacterized protein DBV15_00393, partial [Temnothorax longi</td><td>Expect = 1.200e-89</td><td>gene.scaffold_134.g10.t1</td><td>OG0002287</td><td>124421..126230</td></tr>
<tr><td>Scaffold 1275</td><td>5698820..5699218</td><td>HD001704</td><td>OG0007747</td><td>gi|817198610|ref|XP_012274957.1|proteasome assembly chaperone 4 isoform X1 [Orussus abietinus]</td><td>Expect = 8.200e-35</td><td>gene.scaffold_134.g11.t1</td><td>OG0007747</td><td>128625..129023</td></tr>
<tr><td>Scaffold 1275</td><td>5700119..5709232</td><td>HD001705</td><td>OG0007746</td><td>gi|1000763707|ref|XP_015604930.1 midasin [Cephus cinctus]</td><td>Expect = 0.000e+0</td><td>gene.scaffold_134.g12.t1</td><td>OG0007746</td><td>129973..143060</td></tr>
<tr><td>Scaffold 1275</td><td>5715296..5722376</td><td>HD001706</td><td>OG0005758</td><td>gi|1000763721|ref|XP_015604938.1 uncharacterized protein LOC107272369 isoform X1 [Cephus cinctus</td><td>Expect = 6.200e-245</td><td>gene.scaffold_134.g13.t1</td><td>OG0005758</td><td>142507..148607</td></tr>
<tr><td>Scaffold 1275</td><td>5745780..5748027</td><td>HD001707</td><td>orphan</td><td>no hit</td><td></td><td>NA</td><td></td><td></td></tr>
<tr><td>Scaffold 1275</td><td>5789342..5804341</td><td>HD001708</td><td>OG0010542</td><td>gi|1000763707|ref|XP_015604930.1 midasin [Cephus cinctus]</td><td>Expect = 0.000e+0</td><td>gene.scaffold_6042.g1.t1</td><td>OG0010542</td><td>7898..20048</td></tr>
<tr><td>Scaffold 1275</td><td>5804928..5806155</td><td>HD001709</td><td>OG0008637</td><td>gi|817050671|ref|XP_012269284.1|28S ribosomal protein S15, mitochondrial [Athalia rosae]</td><td>Expect = 3.900e-66</td><td>gene.scaffold_6042.g3.t1</td><td>OG0008637</td><td>20431..21692</td></tr>
<tr><td>Scaffold 1275</td><td>5806461..5808045</td><td>HD001710</td><td>OG0008638</td><td>gi|1000725354|ref|XP_015585242.1 protein AATF [Cephus cinctus]</td><td>Expect = 1.900e-132</td><td>gene.scaffold_6042.g4.t1</td><td>OG0008638</td><td>22007..23884</td></tr>
<tr><td>Scaffold 1275</td><td>5809242..5809907</td><td>HD016565</td><td>orphan</td><td>Rep1_Hd6</td><td></td><td></td><td></td><td></td></tr>
<tr><td>Scaffold 1275</td><td>///</td><td></td><td></td><td></td><td></td><td>gene.scaffold_6042.g5.t1</td><td>OG0000017</td><td>38407..41846</td></tr>
<tr><td>Scaffold 1275</td><td>5818232..5818829</td><td>HD016570</td><td>orphan</td><td>U1.2_Hd6</td><td></td><td></td><td></td><td></td></tr>
<tr><td>Scaffold 1275</td><td>5853342..5855008</td><td>HD001713</td><td>OG0000907</td><td>gi|954550203|ref|XP_014598460.1|PREDICTED: fibrillin-1-like [Polistes canadensis]</td><td>Expect = 2.700e-10</td><td></td><td></td><td></td></tr>
<tr><td>Scaffold 1275</td><td>5859163..5862434</td><td>HD001714</td><td>OG0000759</td><td>gi|766921548|ref|XP_011505297.1|PREDICTED: major royal jelly protein 1-like isoform X1 [Ceratosolen</td><td>Expect = 8.700e-75</td><td>gene.scaffold_6042.g6.t1</td><td>OG0000759</td><td>56163..59912</td></tr>
<tr><td>Scaffold 1275</td><td>5868688..5896004</td><td>HD001715</td><td>OG0000907</td><td>gi|1477752320|ref|XP_006561358.2 fibrillin-2 isoform X1 [Apis mellifera]</td><td>Expect = 1.000e-135</td><td>gene.scaffold_6042.g7.t1</td><td>OG0000907</td><td>67573..82296</td></tr>
<tr><td></td><td></td><td></td><td></td><td></td><td></td><td>gene.scaffold_6042.g8.t1</td><td>OG0000907</td><td>84612..89146</td></tr>
<tr><td>Scaffold 1275</td><td>5899594..5902437</td><td>HD001716</td><td>OG0008827</td><td>gi|925679321|gb|KOX76257.1| hypothetical protein WN51_11586 [Melipona quadrifasciata]</td><td>Expect = 1.300e-41</td><td>gene.scaffold_6042.g9.t1</td><td>OG0008827</td><td>95154..98396</td></tr>
<tr><td>Scaffold 1275</td><td>5907509..5931751</td><td>HD001717</td><td>OG0000907</td><td>gi|951521230|ref|XP_014484382.1|PREDICTED: uncharacterized protein LOC106749449 isoform X1 [Di</td><td>Expect = 1.400e-276</td><td>gene.scaffold_6042.g11.t</td><td>OG0000907</td><td>102320..114213</td></tr>
<tr><td>Scaffold 1275</td><td>5934850..5939667</td><td>HD001718</td><td>OG0003835</td><td>gi|936686298|ref|XP_014221120.1|protein henna isoform X1 [Trichogramma pretiosum]</td><td>Expect = 8.800e-249</td><td>gene.scaffold_6042.g12.t</td><td>OG0003835</td><td>131205..134233</td></tr>
</table>

| | | | | | | | | |
|---|---|---|---|---|---|---|---|---|
| Scaffold 1275 | 5941301..5941799 | HD001719 | orphan | no hit | | NA | | |
| Scaffold 1275 | 5941600..5943451 | HD001720 | orphan | GlyPro1_Hd2 | | NA | | |
| Scaffold 1275 | /// | | | | | NA | | |
| Scaffold 1275 | 5952553..5953461 | HD016569 | orphan | SerThr_Hd2 | | NA | | |
| Scaffold 1275 | 5954908..5957857 | HD001723 | OG0008292 | gi\|970914653\|ref\|XP_015123688.1\|PREDICTED: vacuolar fusion protein MON1 homolog A [Diachasma a | Expect = 1.500e-249 | NA | | |
| Scaffold 1275 | 5958966..5960611 | HD001724 | OG0008989 | gi\|1000737514\|ref\|XP_015591270.1 pre-mRNA-splicing factor 38 isoform X1 [Cephus cinctus] | Expect = 7.300e-111 | NA | | |
| Scaffold 1275 | 5963322..5967084 | HD001725 | OG0006130 | gi\|665789861\|ref\|XP_008560590.1 PREDICTED: lipoma HMGIC fusion partner-like 3 protein [Microplitis | Expect = 1.500e-148 | NA | | |
| Scaffold 1275 | 5970045..5989460 | HD001726 | OG0003941 | gi\|1000737447\|ref\|XP_015591233.1 paired box pox-neuro protein isoform X2 [Cephus cinctus] | Expect = 3.600e-166 | NA | | |
| Scaffold 1275 | 6031309..6031662 | HD001727 | orphan | no hit | | NA | | |
| Scaffold 1275 | 6039464..6041203 | HD001728 | OG0008922 | no hit | | NA | | |
| Scaffold 1275 | 6047894..6053795 | HD001729 | OG0004544 | gi\|970914023\|ref\|XP_015123344.1\|PREDICTED: neither inactivation nor afterpotential protein G-like [D | Expect = 1.100e-201 | NA | | |
| Scaffold 1275 | 6055912..6056223 | HD001730 | orphan | no hit | | gene.scaffold_12.g37.t1 | orphan | 313226..314796 |
| Scaffold 1275 | 6065301..6068372 | HD001731 | orphan | U1_Hd7 | | gene.scaffold_12.g36.t1 | orphan | 269465..272474 |
| Scaffold 1275 | 6163755..6209923 | HD001732 | OG0001182 | gi\|665784719\|ref\|XP_008543429.1\|PREDICTED: zinc finger protein 608-like [Microplitis demolitor] | Expect = 8.000e-240 | gene.scaffold_12.g35.t1 | OG0001182 | 266509..281996 |
| Scaffold 1275 | 6215883..6220739 | HD001733 | OG0009154 | gi\|665785226\|ref\|XP_008546042.1\|PREDICTED: focadhesin [Microplitis demolitor] | Expect = 3.500e-300 | gene.scaffold_12.g34.t1 | OG0009154 | 254503..262293 |
| Scaffold 1275 | 6223570..6225534 | HD001734 | OG0005749 | gi\|759044886\|ref\|XP_011331377.1\|uncharacterized protein C1orf43 homolog [Ooceraea biroi] | Expect = 6.200e-121 | gene.scaffold_12.g33.t1 | OG0005749 | 246626..248241 |
| Scaffold 1275 | 6233178..6236130 | HD001735 | orphan | gi\|815899824\|ref\|XP_012236112.1\|RNA polymerase II elongation factor Ell [Bombus impatiens] | Expect = 2.200e-20 | | | |
| Scaffold 1275 | 6237301..6258275 | HD001736 | OG0001096 | gi\|1000725348\|ref\|XP_015585239.1 NGFI-A-binding protein homolog isoform X1 [Cephus cinctus] | Expect = 1.200e-181 | gene.scaffold_12.g32.t1 | OG0001096 | 233201..241710 |
| Scaffold 1275 | 6262652..6270854 | HD001737 | OG0001213 | gi\|817221046\|ref\|XP_012286061.1\|girdin isoform X1 [Orussus abietinus] | Expect = 4.100e-256 | gene.scaffold_12.g30.t1 | OG0001213 | 210669..216809 |
| Scaffold 1275 | 6273273..6277538 | HD001738 | OG0000750 | gi\|970914033\|ref\|XP_015123349.1\|PREDICTED: E3 ubiquitin-protein ligase RNF126 [Diachasma alloeum | Expect = 5.100e-91 | NA | | |
| Scaffold 1275 | 6295663..6305891 | HD001739 | OG0001026 | gi\|805820120\|ref\|XP_003707425.2\|PREDICTED: monocarboxylate transporter 10-like isoform X2 [Mega | Expect = 4.400e-278 | NA | | |
| Scaffold 1275 | 6326868..6327170 | HD001740 | orphan | no hit | | NA | | |
| Scaffold 1275 | 6327762..6333610 | HD001741 | OG0001259 | gi\|755965602\|ref\|XP_011305227.1\|PREDICTED: uncharacterized protein LOC105267814 [Fopius arisanu | Expect = 3.100e-204 | NA | | |
| Scaffold 1275 | 6336069..6336941 | HD001742 | OG0008975 | gi\|826447337\|ref\|XP_012531547.1\|cytochrome c oxidase assembly factor 5 [Monomorium pharaonis] | Expect = 9.100e-32 | NA | | |
| Scaffold 1275 | 6337655..6338948 | HD001743 | OG0007279 | gi\|970893534\|ref\|XP_015112156.1\|PREDICTED: lysM and putative peptidoglycan-binding domain-conta | Expect = 1.000e-66 | NA | | |
| Scaffold 1275 | /// | | | | | | | |
| Scaffold 1275 | 6498766..6500773 | HD001752 | OG0001027 | gi\|1059871709\|ref\|XP_017790766.1 PREDICTED: uncharacterized protein LOC108572935 [Habropoda la | Expect = 1.300e-295 | gene.scaffold_57.g37.t1 | OG0001027 | 701460..703459 |
| Scaffold 1275 | 6525619..6535728 | HD001753 | OG0008714 | gi\|1000769997\|ref\|XP_015608184.1 zwei Ig domain protein zig-8 isoform X1 [Cephus cinctus] | Expect = 1.300e-115 | gene.scaffold_57.g35.t1 | OG0008714 | 673695..676530 |
| Scaffold 1275 | 6576893..6577922 | HD001754 | orphan | no hit | | gene.scaffold_57.g34.t1 | OG0008714 | 667397..668577 |
| Scaffold 1275 | 6589791..6590075 | HD001755 | orphan | no hit | | | | |
| Scaffold 1275 | 6609826..6610452 | HD001756 | OG0010758 | no hit | | | | |
| Scaffold 1275 | 6654784..6656225 | HD001757 | OG0010662 | no hit | | | | |
| Scaffold 1275 | 6674792..6680997 | HD001758 | OG0010538 | gi\|817211154\|ref\|XP_012281702.1\|uncharacterized protein LOC105700447 [Orussus abietinus] | Expect = 4.400e-177 | gene.scaffold_57.g33.t1 | OG0010538 | 549863..558077 |
| Scaffold 1275 | 6687225..6691596 | HD001759 | OG0000577 | gi\|665815849\|ref\|XP_008556602.1\|PREDICTED: acylcarnitine hydrolase-like [Microplitis demolitor] | Expect = 1.100e-123 | gene.scaffold_57.g32.t1 | OG0000577 | 527847..544092 |
| Scaffold 1275 | 6694467..6698473 | HD001760 | OG0000577 | gi\|665815849\|ref\|XP_008556602.1\|PREDICTED: acylcarnitine hydrolase-like [Microplitis demolitor] | Expect = 2.300e-118 | | | |
| Scaffold 1275 | 6713912..6715341 | HD001761 | OG0010178 | gi\|954544039\|ref\|XP_014610010.1\|PREDICTED: UPF0602 protein C4orf47 homolog [Polistes canadensis | Expect = 2.700e-76 | gene.scaffold_57.g31.t1 | OG0010178 | 512556..513765 |
| Scaffold 1275 | 6755517..6755963 | HD001762 | orphan | no hit | | gene.scaffold_57.g29.t1 | orphan | 480378..482423 |
| Scaffold 1275 | 6788558..6789189 | HD001763 | OG0000376 | gi\|998503073\|ref\|XP_015511626.1\|PREDICTED: uncharacterized protein LOC107218308 [Neodiprion le | Expect = 2.500e-25 | gene.scaffold_57.g28.t1 | OG0000376 | 445783..447418 |
| Scaffold 127548 | | | | | | gene.scaffold_57.g27.t1 | orphan | 394294..400573 |
| Scaffold 1275 | 6832835..6833713 | HD001764 | orphan | IVSP_U29 | | gene.scaffold_57.g26.t1. | orphan | 391820..393241 |
| Scaffold 1275 | /// | | | | | /// | | |
| Scaffold 1275 | 6845548..6848646 | HD001769 | orphan | IVSP_U34 | | gene.scaffold_57.g24.t1. | orphan | 383305..386526 |
| Scaffold 1275 | 6856068..6858621 | HD001770 | OG0000376 | gi\|755932577\|ref\|XP_011314368.1\|PREDICTED: lachesin-like [Fopius arisanus] | Expect = 2.700e-25 | gene.scaffold_57.g22.t1 | OG0000376 | 370164..371046 |
| Scaffold 1275 | 6861330..6863003 | HD001771 | OG0001625 | gi\|1070588568\|ref\|XP_018406776.1 PREDICTED: uncharacterized protein LOC108771307 [Cyphomyrme | Expect = 1.500e-186 | | | |

**Genes in synteny blocks Figure 7C, c**

| Hd scaffold | Position in scaffold | Hd gene ID | Orthogroup # | Best Blast hit | | Cs gene | Orthogroup # | Position in scaffold |
|---|---|---|---|---|---|---|---|---|
| Scaffold 1275 | 10649612..1065171( | HD002066 | OG0004297 | gi\|815768838\|ref\|XP_012233244.1\|PREDICTED: caspase-3 isoform X1 [Linepithema humile] | Expect = 2.800e-70 | gene.scaffold_12.g65.t1 | OG0004297 | 668906..679965 |
| Scaffold 1275 | 10653145..1065624( | HD002067 | OG0009466 | gi\|972201540\|ref\|XP_015182858.1\|PREDICTED: sodium channel protein Nach-like isoform X1 [Polistes | Expect = 6.900e-140 | gene.scaffold_12.g66.t1 | OG0009466 | 681861..685155 |
| Scaffold 1275 | 10696314..1070282 | HD002068 | OG0002519 | gi\|1389785382\|ref\|XP_024946756.1 uncharacterized protein LOC107273655 isoform X4 [Cephus cinctus | Expect = 1.000e-157 | gene.scaffold_12.g67.t1 | OG0002519 | 733521..738350 |
| Scaffold 1275 | 10713573..1071834 | HD002069 | OG0002011 | gi\|817182955\|ref\|XP_012273795.1\|coiled-coil and C2 domain-containing protein 1-like isoform X2 [Oru | Expect = 2.300e-247 | gene.scaffold_12.g68.t1 | OG0002011 | 752453..758163 |
| Scaffold 1275 | 10718988..1072124 | HD002070 | OG0008538 | gi\|817182522\|ref\|XP_012271486.1\|V-type proton ATPase subunit d [Orussus abietinus] | Expect = 3.600e-199 | gene.scaffold_12.g70.t1 | OG0008538 | 758815..760937 |
| Scaffold 1275 | 10721836..1072315 | HD002071 | OG0004841 | gi\|817182518\|ref\|XP_012271468.1\|probable tRNA(His) guanylyltransferase isoform X1 [Orussus abietir | Expect = 2.500e-102 | gene.scaffold_12.g71.t1 | OG0004841 | 761672..762420 |

| | | | | | | | | | |
|---|---|---|---|---|---|---|---|---|---|
| Scaffold 1275 | 10723636..1072892 | HD002072 | OG0002227 | gi\|1000768710\|ref\|XP_015607512.1 | probable methyltransferase TARBP1 isoform X1 [Cephus cinctus] | Expect = 8.500e-229 | gene.scaffold_6122.g8.t1 | OG0002227 | 84619..90498 |
| Scaffold 1275 | 10729352..1073001 | HD002073 | OG0006556 | gi\|970884085\|ref\|XP_015126663.1\| | PREDICTED: ATP synthase subunit g, mitochondrial [Diachasma allo | Expect = 9.800e-33 | gene.scaffold_6122.g9.t1 | OG0006556 | 90282..90940 |
| Scaffold 1275 | 10730306..1073170 | HD002074 | OG0008346 | gi\|817057993\|ref\|XP_012269928.1\| | zinc finger protein 330 homolog [Athalia rosae] | Expect = 2.700e-140 | gene.scaffold_6122.g10.t | OG0008346 | 91228..92932 |
| Scaffold 1275 | 10732736..1073820 | HD002075 | OG0006263 | gi\|665809833\|ref\|XP_008553302.1\| | PREDICTED: arrestin domain-containing protein 2 [Microplitis demc | Expect = 3.600e-183 | gene.scaffold_6122.g11.t | OG0006263 | 93959..96475 |
| Scaffold 1275 | 10740693..1074533 | HD002076 | OG0007281 | gi\|1389758539\|ref\|XP_024943813.1 | synaptotagmin-4 isoform X2 [Cephus cinctus] | Expect = 4.500e-179 | gene.scaffold_6122.g12.t | OG0007281 | 101239..105178 |
| Scaffold 1275 | 10749875..1075318 | HD002077 | OG0007282 | gi\|970904570\|ref\|XP_015118179.1\| | PREDICTED: dolichol kinase isoform X1 [Diachasma alloeum] | Expect = 2.900e-175 | gene.scaffold_6122.g14.t | OG0007282 | 110423..112817 |
| Scaffold 1275 | 10755559..1075930 | HD002078 | OG0005355 | gi\|936697175\|ref\|XP_014226574.1\| | glucose-6-phosphate isomerase [Trichogramma pretiosum] | Expect = 5.700e-287 | gene.scaffold_6122.g15.t | OG0005355 | 116428..120182 |
| Scaffold 1275 | 10761570..1076280 | HD002079 | orphan | IVSP_U15 | | | | | |
| Scaffold 1275 | /// | | | | | | | | |
| Scaffold 1275 | 10785523..1078700 | HD002089 | orphan | IVSP_N-2 | | | | | |
| Scaffold 1275 | 10788460..1079089 | HD002090 | OG0005583 | gi\|1000723328\|ref\|XP_015610175.1 | probable cysteine--tRNA ligase, mitochondrial [Cephus cinctus] | Expect = 7.000e-170 | gene.scaffold_6122.g28.t | OG0005583 | 158304..160353 |
| Scaffold 1275 | 10790972..1079279 | HD002091 | OG0008727 | gi\|1000723356\|ref\|XP_015610186.1 | ribosome production factor 2 homolog [Cephus cinctus] | Expect = 5.600e-130 | gene.scaffold_6122.g29.t | OG0008727 | 160438..162016 |
| Scaffold 1275 | 10794756..1080633 | HD002092 | OG0000882 | gi\|820843698\|ref\|XP_012340057.1\| | PREDICTED: protein polybromo-1 isoform X4 [Apis florea] | Expect = 0.000e+0 | gene.scaffold_6122.g30.t | OG0000882 | 163578..176041 |
| Scaffold 1275 | 10808631..1080918 | HD002093 | OG0004828 | gi\|795054949\|ref\|XP_011871354.1\| | PREDICTED: succinate dehydrogenase assembly factor 4, mitochond | Expect = 4.300e-21 | gene.scaffold_6122.g31.t | OG0004828 | 178896..179652 |
| Scaffold 1275 | 10809957..1081106 | HD002094 | OG0006539 | gi\|925679306\|gb\|KOX76242.1\| | GPI mannosyltransferase 2 [Melipona quadrifasciata] | Expect = 2.200e-60 | | | |
| Scaffold 1275 | 10816751..1084960 | HD002095 | OG0003603 | gi\|1000760603\|ref\|XP_015603308.1 | angiopoietin-2 [Cephus cinctus] >gi\|1389766659\|ref\|XP_0249446C | Expect = 4.400e-295 | gene.scaffold_6122.g32.t | OG0003603 | 208607..221477 |
| Scaffold 1275 | 10837195..1085690 | HD002096 | OG0009828 | gi\|1070171260\|ref\|XP_018346064.1 | PREDICTED: angiopoietin-4 isoform X2 [Trachymyrmex septentriona | Expect = 3.600e-220 | gene.scaffold_6122.g33.t | OG0009828 | 227997..231684 |
| Scaffold 1275 | 10859683..1086000 | HD002097 | orphan | no hit | | | | | |
| Scaffold 1275 | 10860001..1086053 | HD002098 | orphan | IVSP_U35 | | | gene.scaffold_6122.g34.t | orphan | 234803..235273 |
| Scaffold 1275 | 10861166..1086163 | HD002099 | orphan | IVSP_U36 | | | | | |
| Scaffold 1275 | 10862390..1086479 | HD002100 | OG0006143 | gi\|107268279\|ref\|XP_015596389.1\| | PREDICTED: serine/threonine-protein kinase rio2 [Cephus cinctus] | | gene.scaffold_6122.g37.t | OG0006143 | 241011..243008 |
| Scaffold 1275 | 10866093..1087121 | HD002101 | OG0007135 | gi\|749759480\|ref\|XP_011141208.1\| | sorbitol dehydrogenase [Harpegnathos saltator] | Expect = 5.900e-157 | gene.scaffold_6122.g38.t | OG0007135 | 243912..249324 |
| Scaffold 1275 | 10872479..1087731 | HD002102 | OG0007814 | gi\|1569271860\|gb\|QBB02001.1\| | putative UPF0528 protein CG10038 [Cotesia chilonis] | Expect = 1.200e-113 | gene.scaffold_6122.g39.t | OG0007814 | 250807..256468 |
| Scaffold 1275 | 10921663..1092207 | HD002103 | orphan | no hit | | | | | |
| Scaffold 1275 | 10970846..1103466 | HD002104 | OG0002894 | gi\|1494653095\|gb\|RLU26944.1\| | hypothetical protein DMN91_000743 [Ooceraea biroi] | Expect = 1.500e-225 | gene.scaffold_6122.g40.t | OG0002894 | 361174..391485 |
| Scaffold 1275 | 10983515..1098646 | HD002105 | orphan | no hit | | | NA | | |
| Scaffold 1275 | 11051421..1105729 | HD002106 | OG0000642 | gi\|972199364\|ref\|XP_015181683.1\| | PREDICTED: serine/threonine-protein kinase tricorner isoform X1 [F | Expect = 4.700e-266 | gene.scaffold_6095.g2.t1 | OG0000642 | 36848..43783 |
| Scaffold 1275 | 11089871..1109165 | HD002107 | orphan | no hit | | | gene.scaffold_6095.g3.t1 | orphan | 79134..81658 |
| Scaffold 1275 | 11110231..1111272 | HD002108 | OG0000107 | gi\|970917263\|ref\|XP_015125099.1\| | PREDICTED: putative RNA-binding protein EEED8.10 [Diachasma all | Expect = 8.200e-156 | gene.scaffold_6095.g5.t1 | OG0000107 | 105829..109141 |
| Scaffold 1275 | 11118585..1111968 | HD002109 | OG0001394 | gi\|1000769696\|ref\|XP_015608031.1 | dual specificity mitogen-activated protein kinase kinase dSOR1 isof | Expect = 1.300e-106 | gene.scaffold_6095.g6.t1 | OG0001394 | 113898..116393 |
| Scaffold 1275 | 11120053..1112313 | HD002110 | OG0000139 | gi\|156547818\|ref\|XP_001606363.1\| | PREDICTED: sodium/potassium-transporting ATPase subunit alpha- | Expect = 0.000e+0 | gene.scaffold_6095.g7.t1 | OG0000139 | 115279..118353 |
| Scaffold 1275 | 11123403..1112437 | HD002111 | OG0001394 | gi\|571503163\|ref\|XP_393416.2\| | dual specificity mitogen-activated protein kinase kinase dSOR1 [Api | Expect = 1.100e-91 | gene.scaffold_6095.g8.t1 | OG0001394 | 117396..119865 |
