## Additional file 9 for "Conserved and specific genomic features of endogenous polydnaviruses revealed by whole genome sequencing of two ichneumonid wasps"

**<u>Additional File 9.</u>** Sequencing libraries used for the assembly of ichneumonid genomes

A *Campoletis sonorensis* genome

| | **Library ID** | **Insert size (bp)** | **Read length (bp)** | **Data (Gb)** | **Sequence depth (X)** |
|---|---|---|---|---|---|
| Paired-ends | CS_WGS_001 | 250 | 150 | 19.18 | 74.05 |

B. *Hyposoter didymator* genome

| | **Insert size (bp)** | **Read length (bp)** | **Data (Gb)** | **Sequence depth (X)** | **Experiment ID (NCBI SRA)** |
|---|---|---|---|---|---|
| Paired-ends | 250 | 135_145 | 6.96 | 19.89 | SRX7136286 |
| | 500 | 100_110 | 1.52 | 4.33 | SRX7136287 |
| | 800 | 100 | 3.01 | 8.60 | SRX7136288 |
| Mate-pairs | 2,000 | 100 | 7.93 | 22.66 | SRX7136289 |
| | 5,000 | 100 | 5.20 | 14.85 | SRX7136290 |
